# Modeling of African population history using *f*-statistics can be highly biased and is not addressed by previously suggested SNP ascertainment schemes

**DOI:** 10.1101/2023.01.22.525077

**Authors:** Pavel Flegontov, Ulaş Işıldak, Robert Maier, Eren Yüncü, Piya Changmai, David Reich

## Abstract

*f*-statistics have emerged as a first line of analysis for making inferences about demographic history from genome-wide data. These statistics can provide strong evidence for either admixture or cladality, which can be robust to substantial rates of errors or missing data. *f*-statistics are guaranteed to be unbiased under “SNP ascertainment” (analyzing non-randomly chosen subsets of single nucleotide polymorphisms) only if it relies on a population that is an outgroup for all groups analyzed. However, ascertainment on a true outgroup that is not co-analyzed with other populations is often impractical and uncommon in the literature. In this study focused on practical rather than theoretical aspects of SNP ascertainment, we show that many non-outgroup ascertainment schemes lead to false rejection of true demographic histories, as well as to failure to reject incorrect models. But the bias introduced by common ascertainments such as the 1240K panel is mostly limited to situations when more than one sub-Saharan African and/or archaic human groups (Neanderthals and Denisovans) or non-human outgroups are co-modelled, for example, *f*_*4*_-statistics involving one non-African group, two African groups, and one archaic group. Analyzing panels of SNPs polymorphic in archaic humans, which has been suggested as a solution for the ascertainment problem, cannot fix all these problems since for some classes of *f*-statistics it is not a clean outgroup ascertainment, and in other cases it demonstrates relatively low power to reject incorrect demographic models since it provides a relatively small number of variants common in anatomically modern humans. And due to the paucity of high-coverage archaic genomes, archaic individuals used for ascertainment often act as sole representatives of the respective groups in an analysis, and we show that this approach is highly problematic. By carrying out large numbers of simulations of diverse demographic histories, we find that bias in inferences based on *f*-statistics introduced by non-outgroup ascertainment can be minimized if the derived allele frequency spectrum in the population used for ascertainment approaches the spectrum that existed at the root of all groups being co-analyzed. Ascertaining on sites with variants common in a diverse group of African individuals provides a good approximation to such a set of SNPs, addressing the great majority of biases and also retaining high statistical power for studying population history. Such a “pan-African” ascertainment, although not completely problem-free, allows unbiased exploration of demographic models for the widest set of archaic and modern human populations, as compared to the other ascertainment schemes we explored.

## Introduction

Archaeogenetics has achieved remarkable progress in the last decade (Skoglund and Mathieson 2018, Stoneking et al. 2023), with genome-wide data for thousands of ancient humans now being published each year. No region of the world is now inaccessible to archaeogenetic research, although isolation of enough authentic DNA from skeletons excavated in tropical and sub-tropical areas (Lipson et al. 2018) or from Pleistocene individuals (Hajdinjak et al. 2021) remains a challenge. For generating usable archaeogenetic data from Africa, targeted enrichment of human DNA on dedicated single nucleotide polymorphism (SNP) capture panels is almost always necessary. A majority of ancient DNA studies on African populations (Skoglund et al. 2017, van de Loosdrecht et al. 2018, Prendergast et al. 2019, Lipson et al. 2020, Wang et al. 2020, Sirak et al. 2021, Lipson et al. 2022) relied on a SNP capture panel usually termed “1240K” (Fu et al. 2015, Mathieson et al. 2015), and some studies on Upper Paleolithic humans relied on a supplementary panel (“1000K”, comprising transversion polymorphisms found in two Yoruba individuals and transversion polymorphisms in the Altai Neanderthal genome) or on its union with 1240K (Fu et al. 2015, Hajdinjak et al. 2021). The 1240K panel was constructed of the following elements: all SNPs on the Human Origins array (itself composed of 13 sub-panels, each ascertained as heterozygous in a single high-coverage human genome, Patterson et al. 2012), all SNPs on the Illumina 650Y array, all SNPs on the Affymetrix 50k XBA array, and smaller numbers of SNPs chosen for other purposes (Fu et al. 2015). The 1240K capture panel is now used routinely for analyzing thousands of ancient humans across the world (Skoglund and Mathieson 2018, Olalde and Posth 2020), and successor panels including the full set of 1240K sites are now available (Rohland et al. 2022).

Bergström *et al*. (2020), relying on high-quality genomic data for present-day humans, showed that *f*_*4*_-statistics including three sub-Saharan African groups and one non-African group, or four sub-Saharan African (hereafter “African”) groups can be biased when computed on common SNP panels such as Illumina MEGA, the panel used by Li *et al*. (2008), and the Affymetrix Human Origins array (Patterson et al. 2012). An influence of ascertainment on common population genetic analyses (*ADMIXTURE, F*_*ST*_) was also demonstrated. However, the bias in *f*_*4*_-statistics including archaic humans and apes was not explored.

Bergström *et al*. (2020) found that selecting approximately 1.3M SNPs polymorphic in the group composed of high-coverage archaic human genomes (the Altai and Vindija Neanderthals, the “Denisova 3” Denisovan) effectively eliminated the biases affecting *f*_*4*_-statistics calculated on anatomically modern humans (AMH) and including 3 or 4 sub-Saharan African groups. A similar approach (selecting ca. 814K transversion sites variable between the Altai Neanderthal and Denisovan) was proposed by Skoglund *et al*. (2017). A SNP capture reagent relying on this principle, the *myBaits Expert Human Affinities Kit* “Ancestral 850K” module, became available in 2021 from Daicel Arbor Biosciences (https://arborbiosci.com/genomics/targeted-sequencing/mybaits/mybaits-expert/mybaits-expert-human-affinities/). This module targets approximately 850K biallelic transversion SNPs (autosomal and X-chromosomal) ascertained as polymorphic in the group composed of high-coverage archaic human genomes: the Altai (Prüfer et al. 2014), Vindija (Prüfer et al. 2017), and Chagyrskaya Neanderthals (Mafessoni et al. 2020), as well as the “Denisova 3” Denisovan genome (Meyer et al. 2012). This set of variable sites was shown to yield nearly unbiased *F*_*ST*_ values for pairs composed of an African and a non-African group within the Simons Genome Diversity Panel (SGDP) dataset (Mallick et al. 2016), in contrast to the 1240K panel (see a technical note on manufacturer’s website: https://arborbiosci.com/wp-content/uploads/2021/03/Skoglund_Ancestral_850K_Panel_Design.pdf).

These recommendations are motivated by a theoretical property of *f*-statistics: if a SNP is the result of a single historical mutation and there has not been natural selection, the statistics are expected to be unbiased if SNPs are either unascertained or ascertained as polymorphic in a population that is an outgroup for all populations being analyzed (Patterson et al. 2012, Wang and Nielsen 2012), and the results in Bergström *et al*. (2020) and in the technical note published on the Daicel Arbor Biosciences product page are consistent with this theoretical property of outgroup ascertainment. The problematic case is non-outgroup ascertainment, that is ascertainment on a population that is co-analyzed with others. A series of papers explored non-outgroup ascertainment affecting measures of population divergence on simulated data and real data for humans and domestic animals (Nielsen and Signorovitch 2003, Nielsen 2004, Nielsen et al. 2004, Clark et al. 2005, Guillot and Foll 2009, Albrechtsen et al. 2010, Wang and Nielsen 2012, Lachance and Tishkoff 2013, McTavish and Hillis 2015, Malomane et al. 2018, Geibel et al. 2021). However, *D*- and *f*-statistics which have more robustness than other allele frequency-based statistics in many cases (Patterson et al. 2012), were not considered in those studies. Limited exploration of non-outgroup ascertainment schemes was performed on simulated data in publications introducing the *D*- and *f*-statistics, with the conclusion that biases are not noticeable in practice (Durand et al. 2011, Patterson et al. 2012).

The existing recommendations for a bias-free SNP enrichment panel also rely on the assumption that archaic humans are nearly perfect outgroups with respect to all AMH, and the low-level archaic admixture in non-Africans (Green et al. 2010, Reich et al. 2010) does not contribute major bias. However, evidence is accumulating that supports archaic admixture in Africans (Chen et al. 2020, Hubisz et al. 2020), and, according to some models, “super-archaic” ancestry (i.e., symmetrically related to Neanderthals, Denisovans, and AMH) may reach 19% in the common ancestor of AMH (Durvasula and Sankararaman 2020). Moreover, for outgroup ascertainment to be unbiased from the theoretical perspective, the outgroup (or a closely related population) should not be then co-analyzed with other populations (Patterson et al. 2012, Wang and Nielsen 2012), and the individuals used for ascertainment should not be used as sole representatives of the respective groups. However, given the paucity of high-coverage archaic genomes (Meyer et al. 2012, Prüfer et al. 2014, 2017, Mafessoni et al. 2020) and the usefulness of archaic or African outgroups for calculation of *f*_*4*_-and *D*-statistics and for testing of admixture models (Maier et al. 2022 preprint), these recommendations are often ignored in published *f*-statistic, *qpAdm, qpGraph*, and *TreeMix* analyses (e.g., Skolgund et al. 2017, Lipson et al. 2020, 2022, Hajdinjak et al. 2021, Kılınç et al. 2021, Yaka et al. 2021). For instance, archaic individuals are co-analyzed with anatomically modern humans on archaic-ascertained SNPs (Skoglund et al. 2017, Hajdinjak et al. 2021) or a Yoruba group is co-analyzed with non-Africans on Yoruba-ascertained SNPs (Kılınç et al. 2021, Yaka et al. 2021).

Since outgroup ascertainment that is “clean” from the theoretical point of view is rarely used in practice, and since the statistical power of outgroup ascertainment to reject incorrect models of population history was not investigated, it is reasonable to examine the performance of archaic ascertainment and common SNP panels such as 1240K in situations that are often encountered in practice. A technical development important for the work reported here is the *ADMIXTOOLS 2* package (Maier et al. 2022 preprint), which extends the functionality of the original *ADMIXTOOLS* package (Patterson et al. 2012), enabling bootstrap resampling for most tools and a rapid algorithm for finding optima in complex admixture graph topology spaces. The *ADMIXTOOLS 2* package also makes calculating millions of *f*_*4*_-statistics and fitting tens of thousands of admixture graphs to data a routine task. These developments, taken together, allow us to explore biased *f*-statistics more systematically and provide more informed guidelines for future studies.

## Results

### 1. Empirical analyses: exploration of the effect of ascertainment bias on real data

We assembled a set of diploid autosomal genotype calls for 352 individuals (Suppl. Table 1) sequenced at high coverage (Mallick et al. 2016; Fan et al. 2019), including mostly present-day individuals from the Simons Genome Diversity Project (SGDP), several high-coverage ancient genomes with diploid genotype calls (Lazaridis et al. 2014, Fu et al. 2014), and three archaic human genomes: the “Denisova 3” Denisovan (Meyer et al. 2012), Vindija (Prüfer et al. 2017) and Altai Neanderthals (Prüfer et al. 2014). Relying on this “SGDP+archaic” dataset, we explored a wide array of ascertainment schemes: 1) A/T and G/C SNPs (henceforth “AT/GC”) that are, unlike the other mutation classes, unaffected by biased gene conversion (Pouyet et al. 2018), and are also unaffected by deamination ancient DNA damage; 2) random thinning of the unascertained or “AT/GC” sets down to the size approximately equal to that of the 1240K SNP panel if missing data are not allowed on a given population set; 3) the 1240K panel (Fu et al. 2015); 4) the 1000K panel composed of 997,780 SNPs comprising all transversion polymorphisms found in two African (Yoruba) individuals sequenced to high coverage and transversion polymorphisms found in the Altai Neanderthal genome (Fu et al. 2015); 5) the union of the 1000K and 1240K panels termed 2200K (Hajdinjak et al. 2021); 6) various components of the 1240K panel (the sites included in the Illumina 650Y and/or Human Origins SNP arrays, sites included exclusively in one of them, and the remaining sites); 7) the largest Human Origins sub-panels – panel 4 ascertained as sites heterozygous in a single San individual, panel 5 ascertained as sites heterozygous in a single Yoruba individual, their union (panels 4+5), and panel 13 including sites where a randomly chosen San allele is derived relative to the Denisovan (Patterson et al. 2012); 8) all sites polymorphic in a group uniting three high-coverage archaic genomes: the “Denisova 3” Denisovan, the Altai and Vindija Neanderthals (this ascertainment scheme is similar to those proposed by Bergström *et al*. 2020 and in the technical note published on the Daicel Arbor Biosciences product page); 9) transversion sites variable in the group comprising these three high-coverage archaic genomes; 10) restricting to SNPs that have high minor allele frequency (MAF > 5%) in the whole “SGDP+archaic” dataset, i.e. high global MAF; 11) restricting to SNPs having high global MAF combined with taking A/T and G/C SNPs only; 12) restricting to SNPs that have > 5% MAF in a selected African or non-African continental meta-population, irrespective of their frequency in the other meta-populations (there are nine such meta-populations in our dataset, and thus nine different ascertainments, see Suppl. Table 1); 13) restricting to SNPs that have > 5% MAF in a selected continental meta-population, A/T and G/C SNPs only. For a list of SGDP-derived SNP sets explored in this study and their sizes in terms of groups, individuals, and SNPs see Suppl. Table 2.

To investigate the influence of ascertainment on the ranking of admixture graph models according to their fits to data, we analyzed real data, considering sets of five populations and generated all possible admixture graph topologies with two admixture events (32,745 distinct topologies with no fixed outgroup; we considered graphs of this complexity as it was unfeasible to work with exhaustive collections of more complex graphs). First, we tested three combinations of groups (Suppl. Fig. 1). Admixture graph residuals on all sites, on a random subset of them approximately equal to the size of the 1240K set, and on AT/GC sites, are tightly correlated (*R* for a linear model ∼1). Residuals of graph models including non-Africans only are also highly correlated on all sites and 1240K sites (*R* = 0.95-0.99, Suppl. Fig. 1). In contrast, the worst *f*_*4*_-statistic residuals (WR) for graphs including one archaic human, three African groups, and one African group with ca. 60% of non-African ancestry (Fan et al. 2019) are poorly correlated on all sites and 1240K sites (*R* = 0.31-0.35). Thus, admixture graph fit rankings are severely affected by the 1240K ascertainment if certain population combinations are involved. We considered the possibility that this case of poor correlation was driven by admixture graph topologies that were obviously inconsistent with the data – that is, topologies could be shown to be inconsistent with the data based on gold standard SNP sets without ascertainment bias. However, the lack of strong correlation for some combinations of populations is not just driven by graphs with poor fits to the data. For example, WRs of admixture graphs that are well-fitting the data (WR <2.5 SE) on a random subsample of 840,000 sites have worst residuals ranging from nearly 0 SE to about 10 SE (Suppl. Fig. 1) on ca. 845,000 sites included in the 1240K panel. Rejecting a model that fits the data on unascertained data runs the risk of rejecting the true model, as we show on simulated data in the next section. The converse problem also applies: some admixture graphs are well-fitting (WR <2.5 SE) on the 1240K sites but fit a random sample of sites poorly (WR >5 SE, Suppl. Fig. 1).

Next, we explored the same exhaustive set of admixture graph topologies including five groups and two admixture events on the wider collection of ascertainments listed above and on a wider collection of populations. Twelve combinations of five groups including up to two archaic humans, up to five African groups, and up to five non-African groups were tested. In Suppl. Fig. 2 we compare various ways of looking on the effects of ascertainment, using a population quintuplet “Denisovan, Khomani San, Mbuti, Dinka, Mursi” as an example. In Table 1 we focus on the fraction of topologies that are rejected under ascertainment (WR > 3 SE) but accepted on all sites (WR < 3 SE) as a metric appropriate for quantifying the most serious effects of ascertainment bias, namely the probability of rejecting the true model. In the supplementary materials, we also show alternative measures: a metric reflecting the statistical power of ascertainment, namely the fraction of topologies that are accepted under a given ascertainment (WR < 3 SE) but rejected on all sites (WR > 3 SE) (Suppl. Table 3), and *R*^*2*^ of a linear trend for admixture graph WR (Suppl. Figs. 3 and 4, Suppl. Table 4) or log-likelihood (LL) scores (Suppl. Figs. 3 and 4, Suppl. Table 5).

**Table 1.**
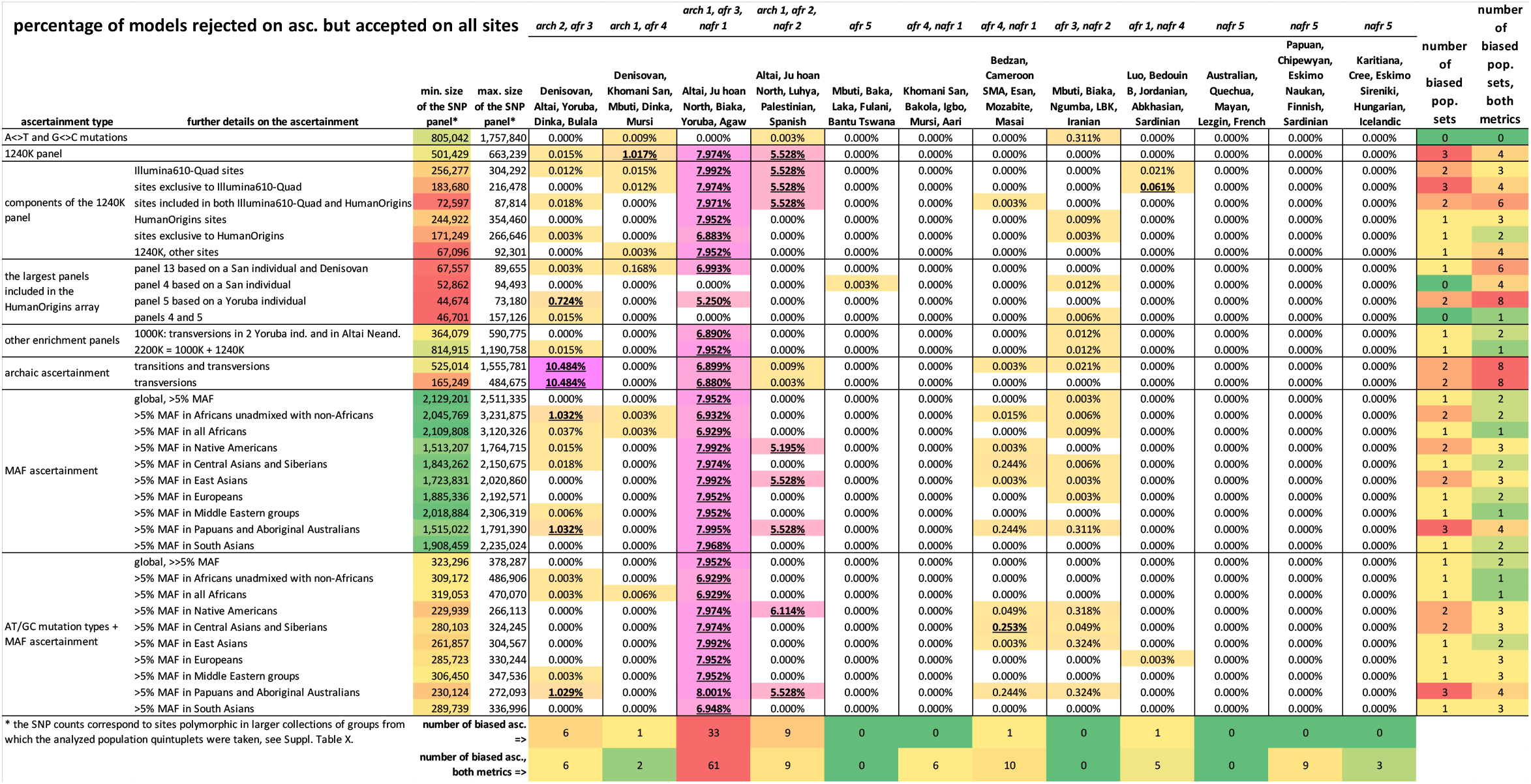
Performance of ascertainment schemes explored across 12 population quintuplets and assessed as the fraction of all possible admixture graph topologies that are rejected under ascertainment (WR > 3 SE) but accepted on all sites (WR < 3 SE). We also applied the binary classifier to determine if the ascertainment produces unbiased or biased results (the latter highlighted in bold and underlined text). The numbers of population quintuplets or ascertainment schemes affected by bias (according to the fraction of topologies that are rejected under ascertainment but accepted on all sites, or according to this metric and the fraction of topologies that are accepted under ascertainment but rejected on all sites) are shown in the two rightmost columns and in two bottom rows, respectively. The composition of the population sets is shown above the table in an abbreviated way: *arch*, archaic humans, followed by the number of archaic groups; *afr*, Africans; *nafr*, non-Africans or Africans with substantial non-African admixture (Fan et al. 2019).

Although we recognize that there can be no strict rule for classifying ascertainments into biased and unbiased ones since they form a continuum, for high-throughput analysis a classifier is useful. Moreover, fits of admixture graphs vary even in the absence of ascertainment bias, due to random site sampling effects (Fig. 1, Suppl. Fig. 2), as was shown in previous work (Maier et al. 2022 preprint). In this study, we considered a SNP set biased if a metric (such as the fraction of topologies rejected under ascertainment but accepted on all sites) was above (or below, as appropriate) the 2.5^th^ percentile of this metric’s distribution across 200 sets of randomly sampled SNPs equal to the size of the 1240K set for a given population combination.

**Fig. 1.**
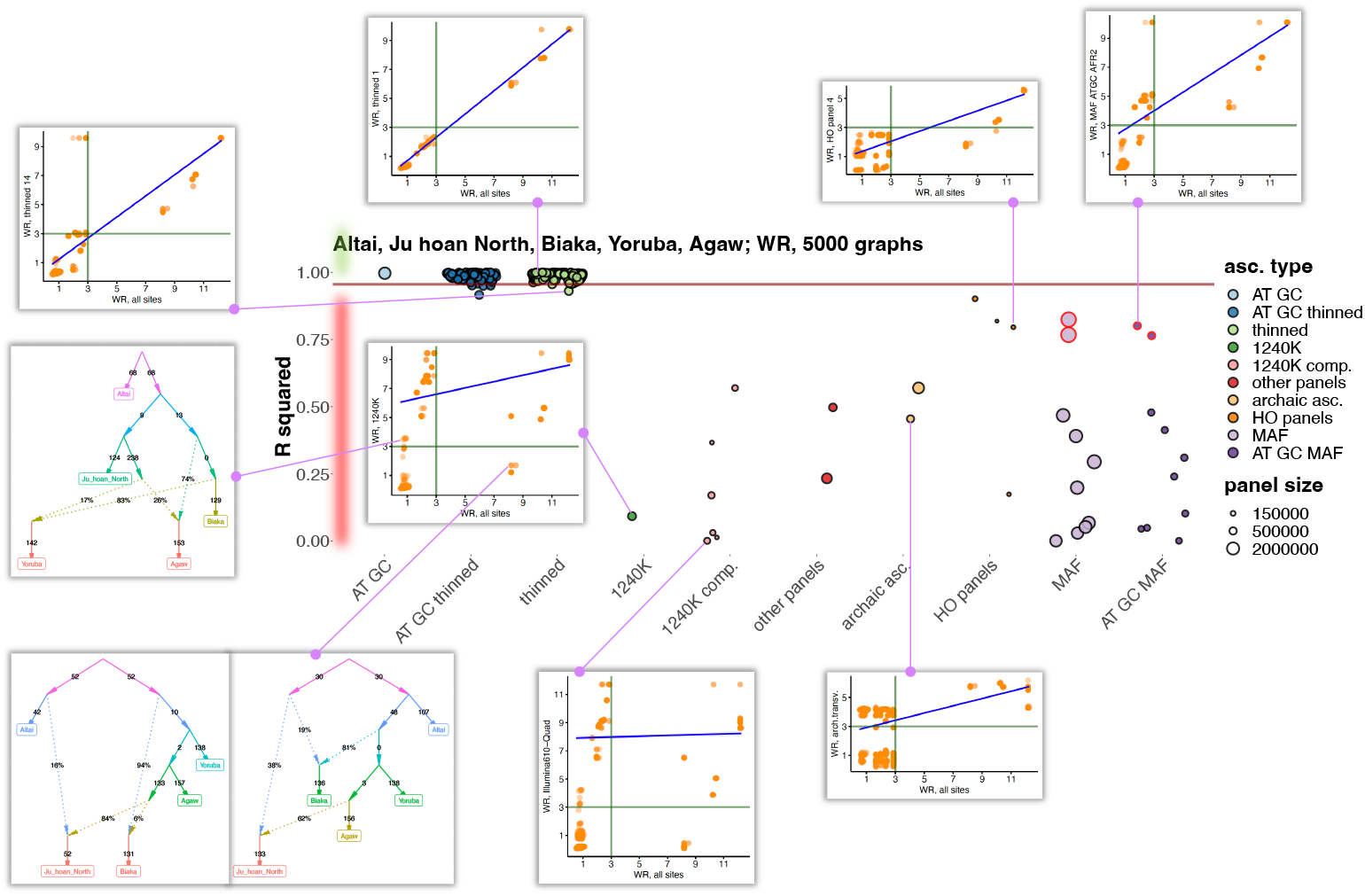
The effect of ascertainment bias on admixture graph fits illustrated on a population combination “Altai Neanderthal, Ju|’hoan North, Biaka, Yoruba, Agaw”. Five thousand best-fitting graphs (according to LL on all sites) of 32,745 possible graphs were selected, and correlation of WR was explored for graphs fitted on all sites and on ascertained sites. Results for ascertainment on variants common in Africans (either those having no detectable West Eurasian ancestry or all Africans in the SGDP dataset) are circled in red. Thirty eight site subsampling schemes were explored: 1) AT/GC mutation classes; 2) random thinning of the AT/GC dataset to the 1240K SNP count for a given combination of groups (no missing data allowed), results for 100 thinned replicates are shown; 3) random thinning of all sites to the 1240K SNP count, results for 100 thinned replicates are shown; 4) the 1240K enrichment panel; 5) major components of the 1240K panel: sites included in the Illumina 650Y and/or Human Origins SNP arrays, sites included exclusively in one of them, and remaining sites; 6) the 1000K and 2200K SNP panels; 7) restricting to sites polymorphic in a group composed of the three high-coverage archaic individuals (either all such sites or transversions only); 8) the largest Human Origins sub-panels (4, 5, 13) or their union (4+5); 9) restricting to common variants based on a global MAF threshold of 5% or on the same threshold in one of nine continental-scale groups; 10) the same procedure repeated on AT/GC sites. The size of the resulting SNP panels is coded by point size, and the ten broad ascertainment types are coded by color according to the legend. *R*^*2*^ values of a linear trend for admixture graph WRs are plotted (WR for the large collections of admixture graphs were compared on all sites and under a particular ascertainment). The 2.5^th^ WR percentile of all the thinned replicates combined, including those on all sites and AT/GC sites, is marked with the brown line. The area of the plot where ascertainments are considered biased according to this classifier is highlighted in red on the left. Scatterplots illustrating effects of selected ascertainment schemes on WR are shown beside the central plot and are connected to the respective data points (ascertainments) by magenta lines. Dots on these scatterplots correspond to distinct admixture graph topologies. Few examples of admixture graphs whose fits are affected by ascertainment are also shown beside the scatterplots.

Inspecting the key metric of ascertainment performance (the fraction of topologies that are rejected under ascertainment but accepted on all sites), we found only three site sampling schemes that were classified as unbiased for all the population quintuplets tested: the A/T and G/C mutation classes, Human Origins panel 4, and the union of Human Origins panels 4 and 5 (Table 1). However, due to the low number of sites in the latter two panels, the union of Human Origins panels 4 and 5, and especially panel 4, lack power to reject admixture graph models as compared to the 1240K panel and to the A/T and G/C mutation classes, as we show in Suppl. Table 3. Thus, the only ascertainment scheme that is problem-free according to both metrics is a random one: taking the A/T and G/C mutation classes.

Among the population quintuplets tested, “Altai Neanderthal, Ju|’hoan North, Biaka, Yoruba, Agaw” (Fig. 1, Suppl. Fig. 3a) and “Altai Neanderthal, Ju|’hoan North, Luhya, Palestinian, Spanish” (Suppl. Fig. 3b) are most susceptible to ascertainment bias (Table 1). A very similar quintuplet “Altai Neanderthal, ancient South African hunter-gatherers, Biaka, Yoruba, Agaw” is encountered within more complex admixture graph models that occupy a central place in Lipson *et al*. (2020, 2022) based on 1240K data (see an investigation of bias affecting the admixture graphs from these studies in Suppl. Text 1 and Suppl. Figs. 5 and 6). As explored below on real and simulated data, a class of *f*_*4*_-statistics that are strongly affected by non-random ascertainment underlies admixture graphs for both problematic population quintuplets: *f*_*4*_(African X, archaic; African Y, non-African). On the other hand, population sets including no archaic human were virtually unbiased (Table 1), but some ascertainment schemes showed limited power to reject admixture graph models in these cases (Suppl. Table 3).

Archaic ascertainment has been suggested in the literature (Skoglund et al. 2017, Bergström et al. 2020) as a way to reduce ascertainment bias, however this approach is guaranteed to work only if the outgroup or a related group is not included itself in admixture graphs or *f*-statistics, and if individuals used for ascertainment are not sole representatives of the respective groups in an analysis. Indeed, our practical-oriented analysis showed that archaic ascertainment is biased in the case of the most problematic population quintuplet “Altai Neanderthal, Ju|’hoan North, Biaka, Yoruba, Agaw” (Table 1); in fact, the archaic ascertainment approach is by far the most biased scheme for population sets including both Neanderthal and Denisovan individuals (Table 1, see also results on simulated data below), and in our analysis it also emerged as the scheme with the lowest statistical power to reject admixture graph models (Suppl. Table 3).

If we combine both key bias metrics (the fraction of topologies rejected under ascertainment but accepted on all sites, and the fraction of topologies accepted under ascertainment but rejected on all sites), the 1240K and archaic ascertainments are out-performed by many ascertainment schemes, and most notably by the following: 1) the union of Human Origins panels 4 and 5; 2) the 2200K panel, which combines various kinds of ascertainment such as the 1240K panel, ascertainment on two Yoruba individuals, and on the Altai Neanderthal (Fu et al. 2015); and 3) restricting to variants that are common in the African meta-population in SGDP (Suppl. Table 1), optionally followed by removal of all mutation classes except for A/T and G/C (Table 1). *R*^*2*^ of a linear trend for admixture graph WR is a metric that in some cases is informative in a way that the fractions of rejected/accepted topologies are not. As illustrated in Fig. 1, *R*^*2*^ may differ substantially across ascertainment schemes while the fractions of topologies rejected under ascertainment but accepted on all sites or *vice versa* stay nearly constant across most ascertainment schemes (Table 1, Suppl. Table 3). Considering *R*^*2*^ for admixture graph WR, restricting to variants that are common in a diverse set of African groups (we term this ascertainment scheme “pan-African” or “African MAF” for convenience) emerges as the least biased form of ascertainment (Suppl. Table 4). We note that conclusions of this sort are not quantitative since our collection of 12 population quintuplets, although diverse, is just a small sample from the vast set of all possible population combinations. However, exploring all possible combinations is infeasible, and we consider our approach to be useful as a practical guide for assessing the performance of ascertainment schemes when admixture graphs including archaic humans, Africans, and non-Africans are fitted to genetic data.

### 2. Simulation studies confirm the qualitative patterns from exploration of empirical data

A major limitation of our empirical analyses of ascertainment bias is that fitting a model with two admixture events is almost certainly inadequate for the histories relating various sets of populations being analyzed. Thus, it is almost certain that all fitted models will be wrong. When we fit wrong models, we have no guarantee that the (incorrect) admixture graph fit to the data will give the same signal of deviation for different SNP ascertainments. Different SNP ascertainments including random ascertainments will simply be sensitive to different aspects of the deviations between the wrong model and the true history. Thus, while the poor correlation between model fits on all sites and under different SNP ascertainment schemes for combinations of archaic humans, sub-Saharan Africans, and non-Africans is a potential signal of bias in analyses, it is valuable to analyze data where the truth is known, as is the case for simulations, to provide clear evidence that typical ascertainment schemes can cause false-positive inferences about history.

Using *msprime v.1.1.1* (Baumdicker et al. 2022), we simulated genetic data (a diploid genome composed of three 100 Mb chromosomes with recombination) that reproduce the *F*_*ST*_ values (Suppl. Fig. 7a) observed when comparing AMH groups, AMH and archaic humans, and AMH and chimpanzee (Fischer et al. 2006). Ten independent simulations were performed under the same parameters and under a topology (Fig. 2a) that in most important aspects conforms to commonly discussed models of the relationships between anatomically modern and archaic humans (Prüfer et al. 2014, Durvasula and Sankararaman 2020). We either simulated or omitted the Neanderthal gene flow to the ancestors of non-Africans (via an unsampled proxy group).

**Fig. 2.**
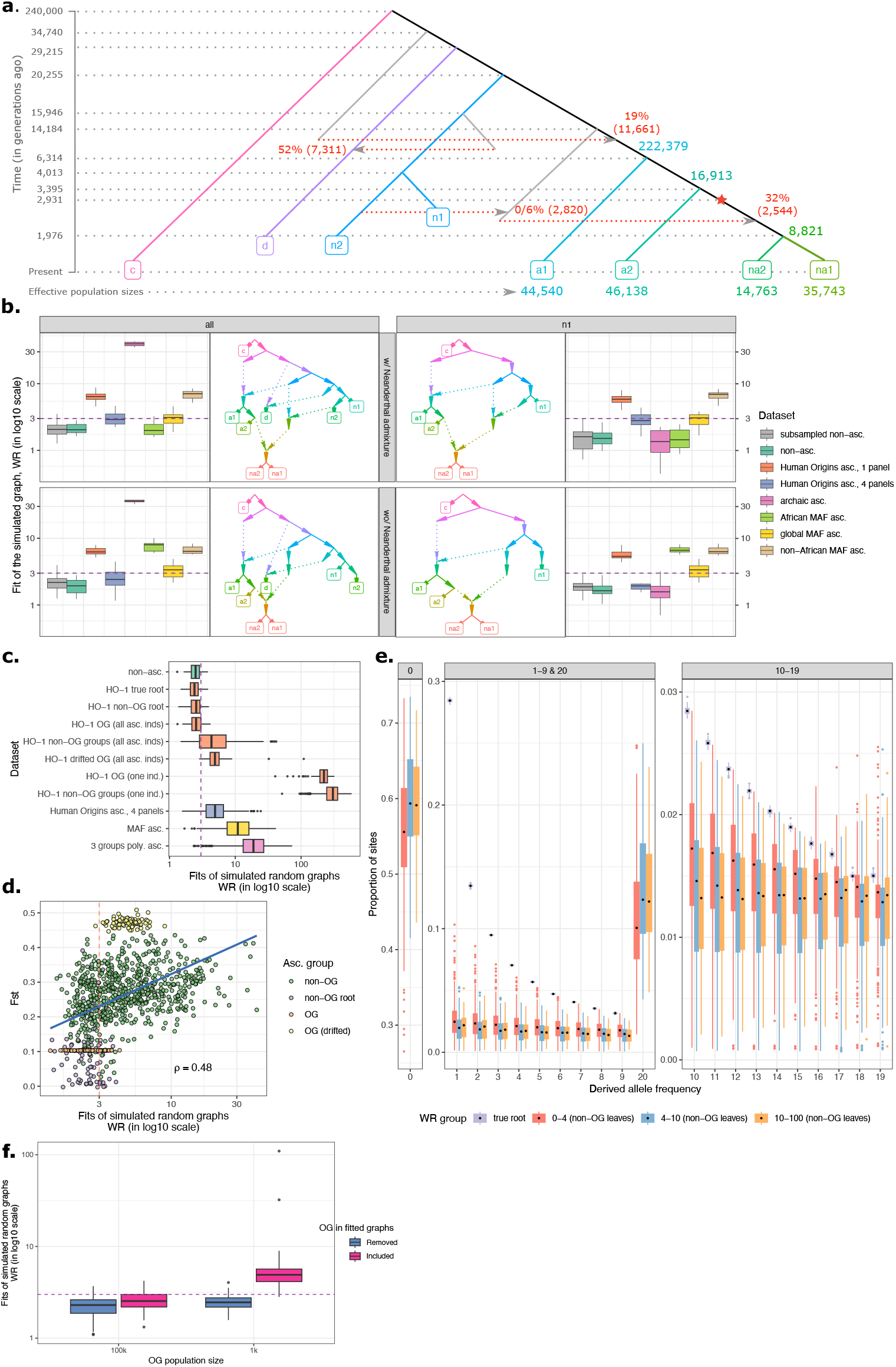
Exploring the influence of non-outgroup ascertainment on fits of admixture graphs on simulated data. Results are presented for two topologies (with or without the Neanderthal to non-African gene flow) and for eight types of SNP sets: 1) 10 sets of randomly selected variable sites matching the average size of the “Human Origins, one panel” set, 500K sites; 2) unascertained sites (on average 5.55M polymorphic sites without missing data); 3) Human Origins-like ascertainment, one panel based on the “a2” group (500K sites on average across simulation iterations); 4) Human Origins-like ascertainment, a union of four panels based on randomly selected individuals from four groups (“a1”, “a2”, “na1”, and “na2”, 1.34M sites on average); 5) archaic ascertainment (1.05M sites on average); 6) “African MAF ascertainment”, that is removal of sites with MAF < 5% in the union of “a1” and “a2” groups (1.85M sites on average); 7) similar MAF ascertainment on the union of “a1”, “a2”, “na1”, “na2” (1.62M sites on average); 8) similar MAF ascertainment on the union of “na1” and “na2” groups (1.48M sites on average). (**a**) The simulated topology, with dates of demographic events and sampling dates (in generations) shown on the y-axis or next to gene flows. The Neanderthal to non-African gene flow was simulated either at 0% (by omitting the “n2” to ghost AMH gene flow) or as shown in the figure. Effective population sizes of archaic groups are omitted for clarity. The out-of-Africa bottleneck is marked with a star. (**b**) Boxplots illustrating the effects of various ascertainment schemes on fits (WR) of the correct admixture graphs. The dashed line on the logarithmic scale marks a threshold often used in the literature for classifying models into fitting and non-fitting ones—3 standard errors—and the observation that common ascertainment schemes consistently produce much higher Z-scores than this threshold provides unambiguous evidence that ascertainment bias can profoundly compromise admixture graph fitting. The topologies fitted to the data are shown beside the boxplots. In the panels on the right, simple graphs including only one archaic lineage are fitted (“n1” used as an example, but very similar results were obtained for the “n2” and “d” groups). In the panels on the left, results for the full simulated model fitted to the data are shown. (**c**) Ascertainment bias was also explored across 80 simulated genetic histories in the form of random admixture graphs. WR of the correct admixture graph was used as a measure of bias. WRs for non-ascertained data and four ascertainment schemes are summarized with boxplots: 1) Human Origins-like ascertainment, one panel; 2) Human Origins-like ascertainment, four panels; 3) MAF-based ascertainment (restricting to common variants) in random sets of four populations; 4) ascertainment on sites polymorphic in random sets of three individuals (one individual sampled per population). Human Origins-like ascertainment on single individuals was performed on the true root or the root of non-outgroup populations, on non-outgroup populations, or on more or less drifted outgroups (having effective population sizes of 100,000 or 1,000 diploid individuals, respectively) co-modelled with the other populations (abbreviated as “OG”). Alternatively, the same individual that was used for ascertainment acted as the only representative of its group for model fitting. (**d**) correct admixture graphs under the Human Origins-like ascertainment (one panel) are guaranteed to be well-fitting (WR < ca. 4 SE) if *F*_*ST*_ between the whole population sample used for ascertainment vs. the sample at the root of the simulation is below 0.12. (**e**) Derived allele frequency spectra (derived allele count in a sample of 20 chromosomes vs. proportion of sites) across simulated root and non-outgroup populations grouped according to the level of ascertainment bias. The spectra were calculated for sites polymorphic in the root population sample of 20 chromosomes. Populations are binned by WR of the true graph fitted to sites heterozygous in a single individual randomly drawn from that population (single-panel Human Origins-like ascertainment). The boxplots summarize DAF across all simulated graphs. DAF bins are shown in three separate panels with different y-axis ranges: 0 derived alleles; 1 to 9 and 20 derived alleles; 10 to 19 derived alleles. (**f**) Illustration of the principle that outgroup ascertainment is expected to be unbiased only if an outgroup (abbreviated as “OG”) is not co-analyzed with the other populations. Human Origins-like ascertainment (one panel) was performed on more or less drifted outgroups (having effective population sizes of 100,000 or 1,000 diploid individuals, respectively) that were then either included in the fitted true admixture graphs or removed from them. WRs of these graphs are summarized with boxplots on the y-axis. The dashed horizontal line corresponds to WR = 3 SE.

We tested several non-outgroup ascertainment schemes: 1) ascertainment on heterozygous sites in a randomly selected individual from the “African 2” group (Fig. 2a, this ascertainment follows the scheme used for generating some of the 12 panels of sites comprising the Human Origins SNP array (Patterson et al. 2012)); 2) ascertainment on heterozygous sites in four randomly selected individuals, one per each “AMH” group (we consider the resulting SNP set to be qualitatively similar to the whole Human Origins SNP set); 3) archaic ascertainment (sites polymorphic in a group composed of one “Denisovan” individual and one individual per each “Neanderthal” group; the same individuals were subsequently used for calculating *f*-statistics); 4) “African MAF ascertainment”, that is restricting to sites with MAF > 5% in the union of two “African” groups; 5) similar MAF-based ascertainment on two “non-African” groups or 6) on all four “AMH” groups.

First, we fitted the correct admixture graph as often practiced in the literature (e.g., Lipson et al. 2020, 2022): including the outgroup, one “archaic” individual, and all “AMH” groups. Human Origins-like ascertainment (one panel) always leads to rejection of the correct model, both in the absence and in the presence of the Neanderthal gene flow to non-Africans, with WR ranging from 3.4 to 8.8 SE (Fig. 2b). Another form of Human Origins-like ascertainment (four panels) is less problematic but led to rejection of the correct model (WR > 3 SE) in 9 of 30 cases (in the presence of the Neanderthal gene flow to non-Africans), with WR up to 4.6 SE. Only the archaic and African MAF non-outgroup ascertainments (in the presence of the Neanderthal gene flow to non-Africans) did not lead to rejection of these simplified graph topologies, known to be correct since we simulated them. However, when the full simulated model (with the Neanderthal gene flow to non-Africans) including the outgroup and three “archaic” lineages is fitted to the data, all non-outgroup ascertainment schemes become problematic, except for African MAF ascertainment (Fig. 2b).

We also investigated effects of ascertainment on model ranking using the same approach as that applied to real data. All possible graph topologies with two admixture events (32,745) were fitted to population quintuplets of the following composition: “d or n1 or n2”, “a1”, “a2”, “na1”, “na2”. The fractions of topologies rejected/accepted under ascertainment but accepted/rejected on all sites (and the bias classifier) were then used to reveal simulation iterations and ascertained schemes that demonstrated biased model fits (Suppl. Table 6). When no Neanderthal/non-African gene flow was simulated, only the non-African MAF ascertainment emerged as problematic (at least half of simulation iterations for at least one population quintuplet were classified as affected by bias) according to the fraction of topologies rejected under ascertainment but accepted on all sites (Suppl. Table 6). When the Neanderthal to non-African gene flow was simulated, all ascertainment schemes, except for the Human Origins-like ascertainment (four panels), emerged as problematic according to the same metric (Suppl. Table 6). Summarizing these results on model ranking and on fits of the true model, we note that the Human Origins-like ascertainment (four panels) is relatively problem-free, unlike archaic ascertainment, MAF-based ascertainments, and Human Origins-like ascertainment (one panel), but it still led to rejection of the true model more often than on all sites or on random site subsamples (Fig. 2b).

Next, we explored non-outgroup ascertainment schemes that are similar to those presented in Fig. 2b but are based on randomly chosen groups (see Methods for details) and were applied to SNP sets resulting from simulated genetic histories in the form of random admixture graphs. Graphs of four complexity classes including 9 or 10 populations and 4 or 5 admixture events were simulated using *msprime v.1.1.1*. Only simulations where pairwise *F*_*ST*_ for groups were in the range characteristic for anatomically modern and archaic humans were selected for further analysis, resulting in 20 random topologies per graph complexity class, each including an outgroup (see examples of the simulated histories and *F*_*ST*_ distributions in Suppl. Fig. 8). Fits of the true admixture graph (WR) including an outgroup were compared on all sites and on ascertained SNP sets for each topology and ascertainment iteration (Fig. 2c). We note that our simulation setup generated groups sampled at different dates in the past (from 0 to ca. 40,000 generations) or, in other words, groups that have experienced widely different levels of genetic drift with respect to the root (Fig. 2d).

As illustrated by distributions of true admixture graph WRs in Fig. 2c, ‘blindly’ ascertaining on individuals or sets of groups randomly sampled across the graph almost guarantees rejecting the true historical model by a wide margin. Ascertainment on sites polymorphic in randomly composed sets of three individuals (one individual per group) and restricting to variants common (MAF > 5%) in randomly composed sets of four populations are two forms of ascertainment that are especially problematic (Fig. 2c). Human Origins-like ascertainments (one or four panels) often yield acceptable fits of the simulated graph (WR < 3 SE), although median WR of the true graphs equals 4.6 SE for these ascertainment schemes across all graph topologies and all (non-root and non-outgroup) populations used for ascertainment (Fig. 2c).

An illuminating result is that *F*_*ST*_ between the population used for Human Origins-like non-outgroup ascertainment and the root influences WR of the true graph: all ascertainments with the corresponding *F*_*ST*_ < 0.12 produce relatively unbiased fits of true graphs (WR < 4 SE, see Fig. 2d). In other words, ascertainment on heterozygous sites in a single individual taken from a population that is not an outgroup and is co-analyzed with other populations, but is genetically close to the root of the simulation, is relatively unbiased, unlike ascertainment on a single individual from a more drifted “present-day” population. We directly illustrate this effect by comparing results on outgroups co-analyzed with other populations that are more or less drifted with respect to the root (with effective population sizes differing by two orders of magnitude) (Fig. 2c,d). Co-analyzing the individual used for ascertainment with other groups does not produce biased results if that individual is a part of a wider population of 10 individuals. However, if that individual is the only representative of its group for model fitting, WRs are inflated drastically (Fig. 2c). We also illustrate the difference between true outgroup ascertainment, when an outgroup is not co-modelled with the other groups, and ascertainment on an outgroup that is included in the fitted model (Fig. 2f), which the kind of ascertainment shown in Fig. 2c and elsewhere. The former form of ascertainment is expected to be unbiased even for a highly drifted outgroup (Fig. 2f), while the latter is not (Patterson et al. 2012, Wang and Nielsen 2012).

Genetic distance from the root is a noisy proxy for the similarity of derived allele frequency (DAF) spectra of the root population and the population used for ascertainment. In Fig. 2e we show DAF spectra across all simulated populations used for Human-Origins-like non-outgroup ascertainment, but only for variants that are polymorphic in the respective root population sample of 10 diploid individuals. These results demonstrate that for the correct admixture graph to fit ascertained data well (WR < 4 SE, Fig. 2e), non-outgroup ascertainment should be based on a population where the derived allele frequency spectrum of sites that were polymorphic at the root is preserved relatively well (see a more extensive version of this plot in Suppl. Fig. 9). We note that some ascertainments may be unbiased with respect to the true graph but may have low power due to the paucity of sites with high MAF in “present-day” populations. Indeed, ascertainments on the root itself or on groups genetically close to the root (such as outgroups with a large effective size) are unbiased (Fig. 2c,d), but on average demonstrated lower power to reject incorrect admixture graph models as compared to single-panel Human Origins-like ascertainments on more drifted groups (Suppl. Fig. 10).

These results on randomized ascertainment schemes (not to be confused with random site sampling) and simulated histories in the form of random admixture graphs show that ascertainment on groups that are highly drifted with respect to the root of the groups being co-analyzed is problematic. Thus, if proper outgroup ascertainment is impractical (if an outgroup shares few polymorphisms with the other populations analyzed, or if an outgroup is needed for constraining the analysis), unascertained or randomly sampled sets of sites should be treated as a gold standard for admixture graph inference. The 1240K ascertainment is much more complex (Fu et al. 2015, Mathieson et al. 2015) than the ascertainment schemes explored on simulated data, but its effects are possibly intermediate between the effects of a MAF-based ascertainment (since all common SNP panels are more or less depleted for rare variants) and ascertainment on heterozygous sites in single individuals from several groups (since approximately half of the 1240K sites are derived from the Human Origins SNP array, Fu et al. 2015). Thus, we expect an accurate admixture graph including at least one archaic human, at least two African groups, and at least one non-African group (Fig. 2a) to fit the data poorly under the 1240K ascertainment.

Finally, we checked if non-outgroup ascertainment could bias the simplest cladality tests in the absence of gene flow. A tree of four groups conforming to the *f*_*4*_-statistic (A, B; C, O) was simulated using *msprime v.1.1.1*, with a tree depth of 4,000 or 40,000 generations (Fig. 3a). All the groups had a uniform effective population size of 100,000 diploid individuals, except for a 10x to 10,000x size reduction immediately after the A-B divergence (1,999 or 9,995 generations in the past). While the dramatic drop in the effective population size of group A yields a complex shape of the derived allele frequency spectrum in {A, B} (Martin and Amos 2021), two of three ascertainment schemes explored here (Human Origins-like ascertainment on one group and MAF-based ascertainment, but not removal of the derived end of the allele frequency spectrum; see Methods) increase the noise in the *f*_*4*_-statistic (A, B; C, O), but do not shift the statistic away from its expectation at 0 (Suppl. Fig. 11). These results confirm an observation by Patterson et al. (2012) that in the case of perfect trees SNP non-outgroup ascertainment does not lead to false rejection of cladality. However, as demonstrated in Fig. 2, non-outgroup ascertainment is generally problematic in the case of complex demographic histories with multiple admixture events.

### 3. An overview of *f*_*4*_-statistic biases caused by non-outgroup ascertainment

We explored various classes of *f*_*4*_-statistics exhaustively to obtain a “bird’s-eye view” on ascertainment biases that was previously difficult to do due to technical challenges in calculating millions of *f*-statistics (Bergström *et al*. 2020). Another motivation for this analysis was the fact that it is unfeasible to explore fits of large collections of admixture graphs on thousands of population sets, ascertainment schemes and random site subsamples. However, if an exhaustively sampled class of *f*-statistics is demonstrated to be unbiased, all admixture graph fits based on those statistics are expected to be unbiased too.

We used the standard deviation of residuals from a linear trend for *f*_*4*_-statistic Z-scores on all sites and under ascertainment (termed “residual SE” for brevity and expressed in the same units as Z-scores) as a single statistic for summarizing results, and we note that it reflects both bias introduced by ascertainment and variance generated by random site sampling. In Suppl. Table 7 and Suppl. Figs. 12 and 13 we show residual SE values for a collection of 27 exhaustively sampled *f*_*4*_-statistic classes and for the large collection of ascertainment schemes that was used for exploring effects of ascertainment on admixture graph fits in section 1. The *f*_*4*_-statistic classes explored can be described concisely as African(all SGDP populations)_*x*_;archaic_*y*_;chimpanzee_*1*_, African(unadmixed with West Eurasians)_*x*_;archaic_*y*_;Mediterranean/Middle Eastern (abbreviated as Med/ME)_*z*_, African(unadmixed with West Eurasians)_*x*_;East Asian_*y*_(y>0), American_*x*_;European_*y*_;Papuan_*z*._ Here, *x, y*, and *z* stand for the number of groups in the population quadruplet; thus, “African_*3*_;East Asian_*1*_” would mean three Africans and one East Asian. All possible distinct *f*_*4*_-statistics composed of those “ingredients” were considered.

The effect of ascertainment schemes varies dramatically across the classes of *f*_*4*_-statistics, but ascertainment schemes based on one or two African individuals (Human Origins sub-panels 4, 5, 13, 4 and 5 combined), on the three archaic individuals (either all sites or transversions only), and components of the 1240K panel such as Illumina650Y emerged as the worst-performing when results across all the *f*_*4*_-statistic classes were considered (Suppl. Table 7, Suppl. Fig. 12). Ascertainment schemes based on a global MAF threshold or on a MAF threshold in a single non-African continental meta-population, and the 1000K and 2200K panels are similar in their effects to the 1240K ascertainment (Suppl. Table 7, Suppl. Fig. 12). We recognize that there is a continuum between unbiased and biased ascertainment schemes, and that for nearly all schemes and *f*_*4*_-statistic classes a majority of statistics remain unaffected by ascertainment, but for describing our results in a concise way and for partially factoring out effects of SNP panel size, we applied the criterion similar to that employed above for admixture graph fits: residual SE for an *f*_*4*_-statistic class is higher than the 97.5^th^ percentile across 200 randomly thinned datasets matching the 1240K panel in size. According to this criterion, the 1240K ascertainment is problematic in the case of the following nine *f*_*4*_-statistic classes (Suppl. Table 7): 1) African_*4*_, 2) African_*3*_;Med/ME_*1*_, 3) African_*3*_;East Asian_*1*_, 4) African_*3*_;archaic_*1*_, 5) African_*3*_;chimpanzee_*1*_, 6) African_*2*_;archaic_*1*_;Med/ME_*1*_, 7) African_*2*_;archaic_*1*_;chimpanzee_*1*_, 8) archaic_*3*_;Med/ME_*1*_, 9) archaic_*3*_;African_*1*_, and unproblematic for the remaining 18 classes exhaustively explored in this analysis. Unlike all the other classes explored here (Suppl. Table 7, Suppl. Fig. 12), the class of statistics African_*2*_;archaic_*1*_;Med/ME_*1*_, specifically statistics of the form *f*_*4*_(African X, archaic; African Y, non-African), is substantially biased under all non-random types of ascertainments (Fig. 4a). The classes African_*3*_;X are problematic under all ascertainment schemes except for the pan-African ascertainment (Suppl. Table 7, see an example in Fig. 4b), and the class African_*4*_ is problematic under all ascertainment schemes except for the 1000K, 2200K, and pan-African ascertainment (Suppl. Table 7). Scatterplots underlying these residual SE estimates are also shown in Fig. 5 (for some of the most problematic classes highlighted above) and in Suppl. Figs. 14-16 (for all classes). Importantly, the classes of statistics most affected by ascertainment (African_*2*_;archaic_*1*_;Med/ME_*1*_, African_*2*_;archaic_*1*_;chimpanzee_*1*_, African_*3*_;X, and African_*4*_) are often relevant for fitting admixture graph models of African population history (see Suppl. Text 1). However, for most classes that were classified as problematic, except for African_*2*_;archaic_*1*_;Med/ME_*1*_, African_*2*_;archaic_*1*_;chimpanzee_*1*_, and African_*3*_;X, residuals remain below 1 SE for a great majority of *f*_*4*_-statistics (Suppl. Table 7 and Suppl. Fig. 12), and thus these statistics are probably not problematic in practice.

**Fig. 4.**
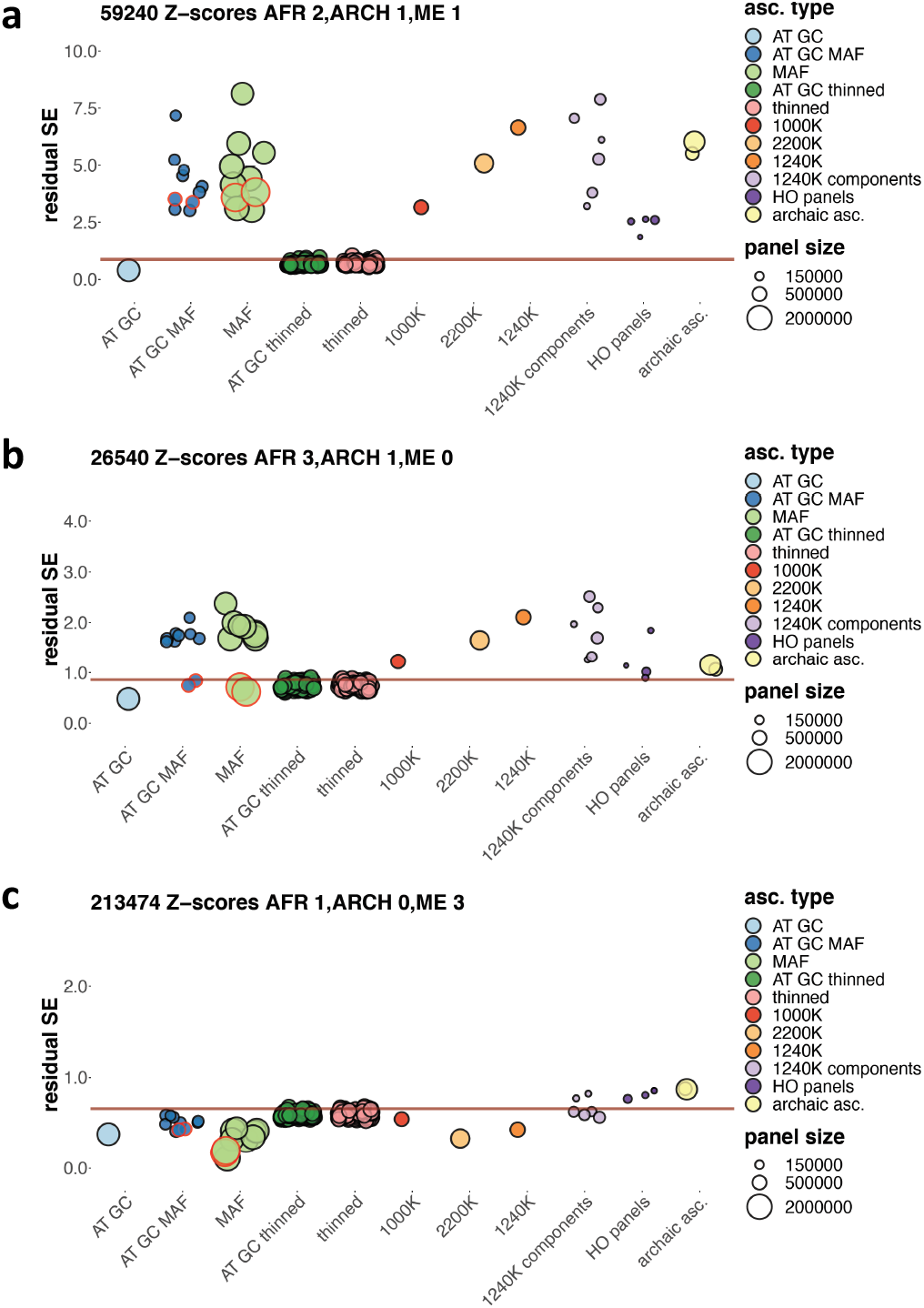
Variance in *f*_*4*_-statistic Z-scores resulting from ascertainment and random site subsampling expressed as standard deviations of residuals of a linear model (expressed in the same units as *f*_*4*_-statistic Z-scores). Results are shown for three classes of *f*_*4*_-statistics: *f*_*4*_(African X, archaic; African Y, Mediterranean/Middle Eastern), *f*_*4*_(African X, African Y; African Z, archaic) (expressed in another notation as African_*3*_;archaic_*1*_), and *f*_*4*_(African, Med/ME X; Med/ME Y, Med/ME Z) (expressed in another notation as African_*1*_;Mediterranean/Middle Eastern_*3*_). Results for ascertainment on variants common in Africans (either those having no detectable West Eurasian ancestry according to Fan et al. (2019) or on all Africans in the SGDP dataset) are circled in red. Residual SE values for *f*_*4*_-statistic Z-scores lying not far from 0 (absolute Z-scores on all sites < 15) are plotted. The 97.5% percentiles of all the thinned replicates combined, including those on all sites and AT/GC sites, are marked by the brown lines. Size of the SNP panels is coded by point size, and the broad ascertainment types are coded by color according to the legend. Thirty eight site subsampling schemes were explored: 1) AT/GC mutation classes; 2) AT/GC mutation classes and restricting to common variants based on a global MAF threshold of 5% or on the same threshold applied to one of nine continental-scale groups; 3) the same procedure repeated on all sites; 4) random thinning of the AT/GC dataset to the 1240K SNP count for a given combination of groups (no missing data allowed), results for 100 thinning replicates are shown; 5) random thinning of all sites to the 1240K SNP count, results for 100 thinning replicates are shown; 6) the 1000K, 1240K, and 2200K enrichment panels (2200K = 1240K + 1000K); 7) major components of the 1240K panel: sites included in the Illumina 650Y and/or Human Origins SNP arrays, sites included exclusively in one of them, and remaining sites; 8) the largest Human Origins sub-panels (4, 5, 13) or their union (4+5); 9) restricting to sites polymorphic in a group composed of the three high-coverage archaic individuals (either all such sites or transversions only).

**Fig. 5.**
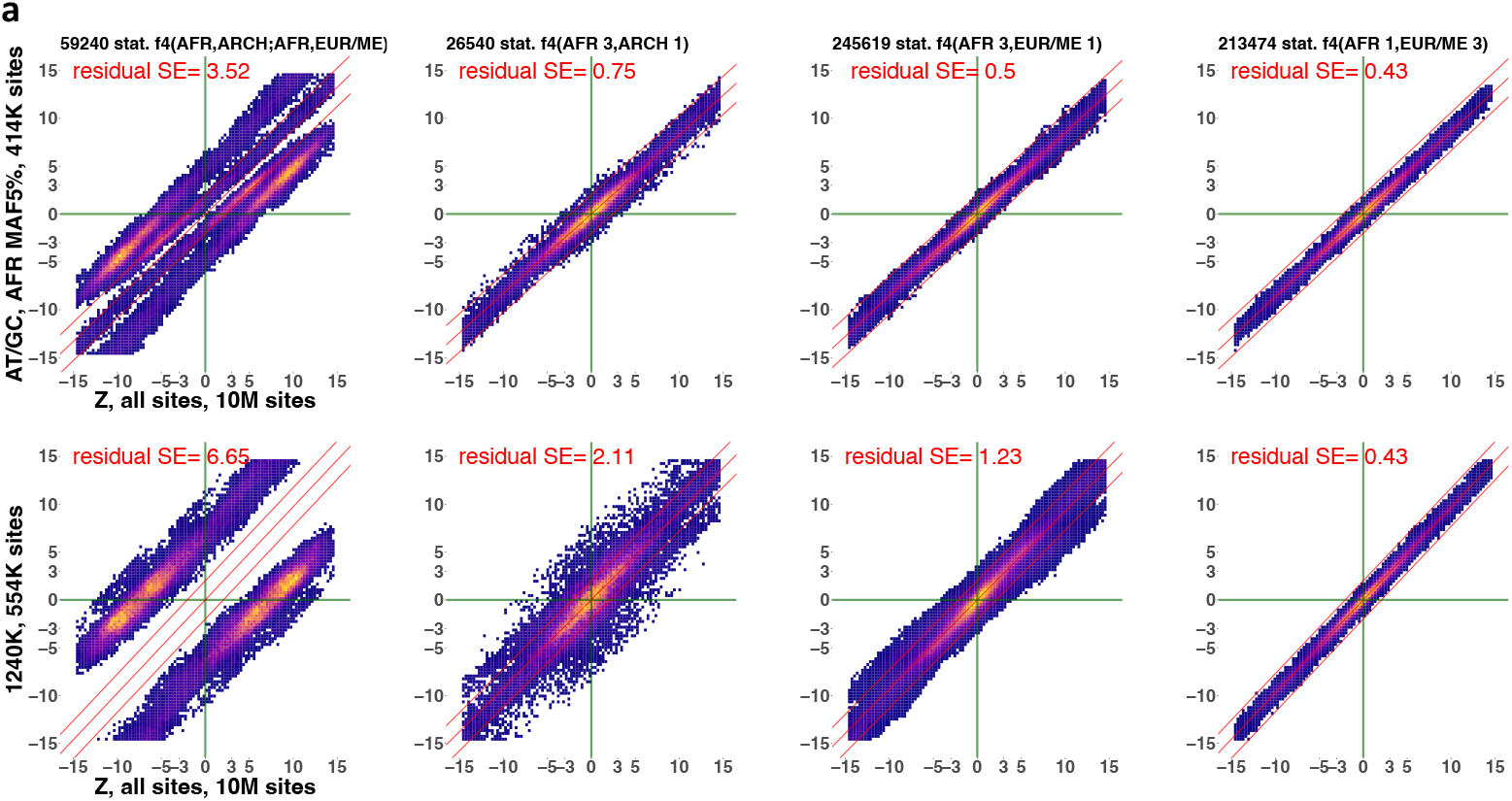
Scatterplots illustrating the effects of two ascertainment schemes on Z-scores of *f*_*4*_-statistics of four classes including African and/or archaic and/or Mediterranean/Middle Eastern groups. Only statistics of the form *f*_*4*_(African X, archaic; African Y, non-African) were considered in the class African_*2*_;archaic_*1*_;Med/ME_*1*_. The *f*_*4*_-statistic classes were selected to represent severe ascertainment bias (leftmost panels), moderate level of bias (two middle panels) and no bias (rightmost panels). The ascertainments selected are 1240K (the most widely used SNP enrichment panel) and the new “pan-African” scheme proposed in this study to mitigate ascertainment bias for nearly all *f*_*4*_-statistic classes. For results on other *f*_*4*_-statistic classes see Suppl. Fig. 14, and results for a wider range of ascertainment schemes are summarized in Suppl. Figs. 12 and 13. Class labels and numbers of statistics plotted are shown above the panels. Instead of individual points, heatmaps illustrating point density are shown. Z-scores on all sites (10 million sites, as indicated on the x-axes) are compared to Z-scores on ascertained datasets on the y-axes. Ascertainment types and site counts are shown on the y-axes. All plots include only statistics with absolute Z-scores below 15 on all sites. A linear model fitted to the data and lines representing ± 2 SE are shown in red. Residual SE values for those linear trends are shown in each plot in red.

Results very similar to those presented above were obtained with a different metric: *R*^*2*^ of a linear model for *f*_*4*_-statistics themselves (Suppl. Table 8), instead of residual SE of a linear model for *f*_*4*_-statistic Z-scores (Suppl. Table 7). For additional details on *f*_*4*_-statistic classes see Suppl. Text 2 (and Suppl. Figs. 13-16), and for a dissection of effects of ascertainment on few selected *f*_*4*_-statistics see Suppl. Text 3 (and Suppl. Tables 9-12).

In contrast, statistics including non-Africans only, or one or two African groups and non-Africans (see an example in Figs. 4c and 5), are unproblematic under the 1240K, 2200K, pan-African and other MAF-based ascertainments (but demonstrate increased variance due to paucity of sites with high MAF under some other ascertainment types such as Human Origins panels 4&5 and archaic ascertainment, Suppl. Table 7).

Pan-African ascertainment (restricting to variants common across 68 African individuals unadmixed with West Eurasians, or across 94 African individuals, Suppl. Table 1), emerged as the best-performing non-outgroup ascertainment scheme. Unlike the other ascertainment schemes explored in this study, this type of ascertainment demonstrates a bias only in the case of the (African X, archaic; African Y, non-African) class of *f*_*4*_-statistics (when only statistics with |Z| < 15 SE on all sites were considered, Suppl. Table 7, Suppl. Fig. 12). Another class of *f*_*4*_-statistics is biased under this ascertainment scheme when all statistics are considered: *f*_*4*_(non-African X, archaic; African, non-African Y) (Suppl. Fig. 12), and pan-African ascertainment is unbiased in the case of the other 25 classes of *f*_*4*_-statistics explored in this study (Suppl. Table 7, Suppl. Fig. 12), which also translates into downstream analyses such as fits of admixture graph models (Fig. 1, Table 1, Suppl. Fig. 3, Suppl. Tables 3-5).

## Discussion

*f*-statistics (Patterson et al. 2012) form a foundation for a range of methods (*qpWave, qpAdm, qpGraph*) that are used widely for studying population genetic history of humans and other species (see, for instance, Bergström et al. 2020b, Librado et al. 2021). Here, we focused on *f*_*4*_-statistics, which are used as standalone tests for cladality (Reich et al. 2009, Patterson et al. 2012) and underlie the *qpAdm* method for fitting admixture models (Haak et al. 2015, Harney et al. 2021). The other *f*-statistics (*f*_*2*_ and *f*_*3*_) can be defined as special cases of *f*_*4*_-statistics [*f*_*2*_(*A, B*) = *f*_*4*_(*A, B; A, B*) and *f*_*3*_(*A; B, C*) = *f*_*4*_(*A, B; A, C*)], and are subject to the same kinds of biases. The existence of bias in the case of non-outgroup ascertainment was recognized in a publication introducing a suite of methods relying on *f-*statistics (Patterson et al. 2012), but its effects on large collections of *f*_*4*_-statistics or on fits of diverse admixture graph models were not explored in that study and in subsequent studies. Since usage of archaic or African outgroups is often unavoidable for calculation of *f*_*4*_-and *D*-statistics and for construction of admixture graph or *qpAdm* models (e.g., Skolgund et al. 2017, Lipson et al. 2020, 2022, Hajdinjak et al. 2021, Kılınç et al. 2021, Yaka et al. 2021), unbiased ascertainment on an outgroup that is not co-analyzed with other populations (as illustrated on simulated data in Fig. 2f) is uncommon in practice. And frequently used SNP panels such as 1240K were built using very complex forms of non-outgroup ascertainment. Therefore, in this study we focused on practical rather than theoretical aspects of the ascertainment bias problem and considered forms of non-outgroup ascertainment that are common in the literature on archaeogenetics of humans, including ascertainment on a phylogenetic outgroup co-analyzed with other populations.

The present analysis showed that *f*_*4*_-statistics of specific types are affected by ascertainment bias. The most striking example we found is a class of statistics *f*_*4*_(African X, archaic; African Y, non-African). All statistics in this class are strongly biased in the same direction under the 1240K ascertainment (Suppl. Fig. 15) and under all other non-random ascertainment schemes explored on real (Suppl. Table 7, Suppl. Fig. 12d) and simulated data (Suppl. Fig. 7b). In contrast, all *f*_*4*_-statistic classes we explored including one or two African groups and non-Africans, or non-Africans only, turned out to be unbiased under the 1240K ascertainment (Suppl. Table 7, Suppl. Fig. 12). Thus, numerous studies relying on fitting *qpAdm* and/or admixture graph models including one African group and various non-Africans are probably minimally affected by ascertainment bias, as we also demonstrated on exhaustive collections of simple admixture graphs for few population quintuplets (Fig. 1, Table 1, Suppl. Fig. 1, Suppl. Tables 3-5). When these classes of methods are applied to African population history, the situation is different, however. As we demonstrated, the 1240K panel emerges as biased when fits of simple admixture graphs including five African groups or one or two archaic and three or four African groups are considered (Fig. 1, Table 1, Suppl. Fig. 1, Suppl. Tables 3-5). We also demonstrated that the 1240K ascertainment affects fits of more complex admixture graphs including in all cases chimpanzee and Altai Neanderthal, and also four or six African groups and one or two groups with substantial non-African ancestry (Supp. Text 1, Suppl. Figs. 5 and 6). We expect fits of many other admixture graphs for Africans beyond those tested in this study to be affected by the 1240K ascertainment since the *f*_*4*_-statistic classes African_*2*_;archaic_*1*_;non-African_*1*_, African_*2*_;archaic_*1*_;chimpanzee_*1*_, and African_*3*_;X are substantially biased under this ascertainment (Suppl. Table 7). These effects were reproduced on simulated data when accurate graphs including “chimpanzee”, one “archaic” lineage, and several “African” and “non-African” lineages were fitted to the data ascertained in various ways (Fig. 2b).

In line with theoretical expectations, *f*_*4*_-statistics including AMH groups only are largely unbiased under archaic ascertainment (Skoglund et al. 2017, Bergström et al. 2020, technical note published on the Daicel Arbor Biosciences product page). However, as compared to other SNP panels of similar size, archaic ascertainment increases variance in nearly all *f*_*4*_-statistic classes of the types non-African_*3*_;X and non-African_*4*_ (Suppl. Table 7, Suppl. Figs. 12-14). Increased variance in these cases can be explained by the low information content of an archaically ascertained panel: unlike the other non-random ascertainment schemes we tested, archaic ascertainment preserves most sites with nearly fixed ancestral variants and leads to just a moderate enrichment for common variants (DAF between 5% and 95%), especially if DAF is based on non-Africans (Suppl. Text 3, Suppl. Fig. 7d, Suppl. Table 10). Thus, the archaically ascertained panel includes a relatively small number of variants that are common in AMH and especially in non-Africans (Suppl. Table 10), and that increases the noise level. This elevated noise level in *f*-statistics under archaic ascertainment translates to reduced power to reject admixture graph models based on these *f*-statistics (Fig. 1, Table 1, Suppl. Fig. 1, Suppl. Tables 3-5). This effect was also reproduced on simulated data (Suppl. Fig. 10). If archaic humans are included in an *f*-statistic or an admixture graph, archaic ascertainment is no longer guaranteed to be unbiased (see Fig. 2 c,d), and indeed due to the existence of the Neanderthal to non-African gene flow it fails to fix the bias affecting the most problematic class of statistics *f*_*4*_(African X, archaic; African Y, non-African), as demonstrated on simulated data in Suppl. Figs. 7b and 17.

Many ascertainment schemes such as the 1240K, 2000K, Illumina 650Y panels and MAF-based ascertainment on non-Africans skew average DAF across populations in the quadruplet since these panels are enriched for derived variants common in non-Africans vs. Africans and in AMH vs. archaic humans (Suppl. Text 3, Suppl. Table 10). Overrepresentation of derived variants in certain groups of the quadruplet skews *f*_*4*_-statistics. We conclude that two ascertainment schemes most often used for studies of African population history (1240K and archaic ascertainment) are not optimal for various reasons: overrepresentation of derived variants common in non-Africans in the former case and a small number of variants common in AMH in the latter case.

We found that there exists a non-outgroup ascertainment scheme that is less biased than the other schemes we tested: restricting to variants that are common in a diverse collection of African groups. This scheme demonstrated a bias only in the case of the *f*_*4*_(African X, archaic; African Y, non-African) and *f*_*4*_(non-African X, archaic; African, non-African Y) classes of *f*_*4*_-statistics among the 27 classes investigated (Suppl. Table 7, Suppl. Figs. 12, 14, and 17). This scheme does not favor derived variants common in non-Africans and supplies many variants common in both Africans and non-Africans (Suppl. Table 10). While for many *f*_*4*_-statitic classes and admixture graphs, the difference in performance of the pan-African and archaic ascertainment schemes is small (Table 1, Suppl. Figs. 3, 4, and 12, Suppl. Tables 3-5 and 7), the pan-African scheme is applicable when Neanderthals and Denisovans are co-analyzed (Suppl. Figs. 3 and 4), while archaic ascertainment generates extreme shifts in *f*_*4*_-statistics in this case (Fig. 2b, Suppl. Fig. 18). The pan-African scheme is also effective for analyses focused on non-Africans, demonstrating no elevated noise level typical for archaic ascertainment (Suppl. Table 7, Suppl. Table 3). Thus, pan-African ascertainment is the most widely applicable scheme among those explored in this study. According to our results on collections of admixture graphs (Table 1) and on *f*_*4*_-statistic classes (Suppl. Table 7), a similar form of ascertainment, namely combining sites heterozygous in a single San and a single Yoruba individual (Human Origins panels 4 & 5) is also largely unbiased, with the exception of statistics of the form *f*_*4*_(African X, archaic; African Y, non-African). However, this ascertainment is also more noisy due to the low number of sites available.

As we demonstrated on simulated data, for a non-outgroup ascertainment to be unbiased it should be based on a population where the derived allele frequency spectrum of sites that were polymorphic at the root is preserved relatively well (Fig. 2e) (however, such an ascertainment usually has relatively low statistical power for rejection of incorrect admixture graph models, see Suppl. Fig. 10). We note that the group where ascertainment was performed was co-modelled with the other groups, as is often done in practice. In the light of these results, archaic ascertainment’s sub-optimal performance as a non-outgroup ascertainment is due to the fact that Denisovans and Neanderthals have had a low long-term effective population size (Mafessoni et al. 2020), and thus are highly drifted with respect to the root. Moreover, it is often unavoidable that individuals used for archaic ascertainment are used as sole representatives of the respective groups analyzed, and that is also problematic (Fig. 2c). Africans, in contrast, have had much higher effective population sizes (Mafessoni et al. 2020), and we propose that restricting to variants common in a diverse set of African genomes is much more reliable (than archaic ascertainment or ascertainment on a single African population or individual) for preserving the spectrum of variants that existed at the root of archaic and anatomically modern humans. At the same time, pan-African ascertainment supplies enough variants that are common in non-Africans, making it also relatively powerful statistically for analyses focused on non-Africans.

An enrichment approach is powerful for large-scale ancient DNA research in Africa due to DNA preservation issues in the hot climate (Skoglund et al. 2017). We did not test a range of allele frequency cut-offs or counts of individuals for pan-African ascertainment, and we do not propose a list of sites for a new DNA enrichment panel. However, our results imply that an effective approach for designing such a panel, which would also be useful for human archaeogenetic studies worldwide, would be to combine selection of the A/T and G/C mutation types with depletion of variants rare in Africa. Frequencies of alleles at A/T and G/C loci are not affected by biased gene conversion (its rate depends on population heterozygosity, Pouyet et al. 2018), and these loci are not hypermutable, and are not affected by deamination damage in ancient DNA. As we demonstrated, restricting to A/T and G/C sites does not bias *f*_*4*_-statistics (Suppl. Table 7, Suppl. Figs. 12 and 16, Suppl. Table 10) or admixture graph fits (Table 1, Suppl. Tables 3-5). Another reason for taking AT/GC sites only is simply reducing the number of sites since enrichment reagents have limited capacity, and this ascertainment scheme with a 5% MAF threshold yields about 1.6 million variable sites on the “SGDP+archaic” dataset.

## Methods

### 1. Simulating genetic data

#### 1.1 Simulating the relationships of AMH and archaic humans with msprime v.0.7.4

Twenty-two chromosomes matching the size of the human chromosomes in the *hg19* assembly were simulated with a flat recombination rate (2 × 10^−8^ per nt per generation) and a flat mutation rate, 1.25 × 10^−8^ per nt per generation (Scally & Durbin 2012). The standard coalescent simulation algorithm was used (Kelleher et al. 2016), and diploid genomes were assembled from these independently simulated 22 haploid chromosomes. Although this approach does not recapitulate the linkage disequilibrium pattern in real human genomes, it does not make a difference for simulating allele frequencies in deeply divergent groups since chromosome histories become quickly independent in the past (Nelson et al. 2020).

The following groups were simulated: chimpanzee (“Chimp”, one individual sampled at the end of the simulation), the Vindija Neanderthal (“Neanderthal”, one individual sampled 2,000 generations or 50,000 years in the past, considering a generation time of 25 years), the high-coverage Denisovan “Denisova 3” (“Denisovan”, one individual sampled 2,000 generations in the past), five African groups (10 individuals per group sampled at “present”) and three non-African groups (10 individuals per group sampled at “present”). Five classes of simulated topologies are shown in Suppl. Fig. 17b; for a full list of simulation parameters and their values see Suppl. Table 13. Only one simulation iteration was performed for each combination of parameters.

We applied archaic ascertainment to the simulated data: restricting to sites polymorphic in the group composed of two “archaic” individuals, “Denisovan” and “Neanderthal” (this scheme reproduces the archaic ascertainment applied to real data, the “SGDP+archaic” dataset, in Suppl. Figs. 17a and 18a). For calculating *f*-statistics on unascertained and ascertained SNP sets, the software package *ADMIXTOOLS 2* (Maier et al. 2022 preprint) was used. Since there was no missing data and all individuals were diploid, we first calculated all possible *f*^*2*^-statistics for 4 Mbp-sized genome blocks (with the “*maxmiss=0*”, “*adjust_pseudohaploid=FALSE*”, and “*minac2=FALSE*” settings), and then used them for calculating *f*_*4*_-statistics as linear combinations of *f*_*2*_-statistics. This protocol was used for generating the results shown in Suppl. Figs. 17c and 18c.

#### 1.2 Simulating the relationships of AMH and archaic humans with msprime v.1.1.1

More realistic simulations were performed with *msprime v.1.1.1* which allows accurate simulation of recombination and of multi-chromosome diploid genomes relying on the Wright-Fisher model (Nelson et al. 2020, Baumdicker et al. 2022). We simulated three chromosomes (each 100 Mb long) in a diploid genome by specifying a flat recombination rate (2 × 10^−8^ per nt per generation) along the chromosome and a much higher rate at the chromosome boundaries (log_e_2 or ∼0.693 per nt per generation, see https://tskit.dev/msprime/docs/stable/ancestry.html#multiple-chromosomes). A flat mutation rate, 1.25 × 10^−8^ per nt per generation (Scally & Durbin 2012), and the binary mutation model were used. To maintain the correct correlation between chromosomes, the discrete time Wright-Fischer model was used for 25 generations into the past, and deeper in the past the standard coalescent simulation algorithm was used (as recommended by Nelson et al. 2020).

The following groups were simulated: chimpanzee (“c”, one individual sampled at the end of the simulation), the Altai Neanderthal (“n1”, one individual sampled 3,790 generations in the past), the Vindija Neanderthal (“n2”, one individual sampled 1,700 generations in the past), the high-coverage Denisovan “Denisova 3” (“d”, one individual sampled 1,700 generations in the past), two African groups (“a1” and “a2”, 10 individuals per group sampled at “present”) and two non-African groups (“na1” and “na2”, 10 individuals per group sampled at “present”). The topology, dates and some effective population sizes are shown in Fig. 2a; for a full list of simulation parameters see Suppl. Table 13. Ten simulation iterations were performed for each combination of parameters, and two combinations were tested: with or without the Neanderthal to non-African gene flow.

Upon assessing genetic distances across the simulated groups using *F*_*ST*_, the following ascertainment schemes were applied:

1. restricting to sites that are heterozygous in a randomly selected individual from the “a2” group (this scheme simulates the generation of one Human Origins SNP panel, Patterson et al. 2012);
2. taking heterozygous sites from one randomly selected individual per “AMH” population (“a1”, “a2”, “na1”, “na2”) and merging these SNP sets (this scheme simulates the generation of the whole Human Origins SNP array, Patterson et al. 2012);
3. restricting to sites having high minor allele frequency (> 5%) in the union of “African” groups “a1” and “a2” (this scheme simulates the MAF ascertainment on Africans);
4. restricting to sites having high minor allele frequency (> 5%) in the union of “non-African” groups “na1” and “na2” (this scheme simulates the MAF ascertainment on a non-African continental meta-population);
5. restricting to sites having high minor allele frequency (> 5%) in the union of all “AMH” groups “a1”, “a2”, “na1”, and “na2” (this scheme simulates the global MAF ascertainment);
6. restricting to sites polymorphic in the group composed of three “archaic” individuals, “d”, “n1”, and “n2” (this scheme simulates the archaic ascertainment applied to the “SGDP+archaic” dataset throughout this study).

#### 1.3 Simulating random admixture graphs and simple trees with msprime v.1.1.1

Genetic histories in the form of random admixture graphs were simulated using the *msprime v.1.1.1* settings described above. We simulated admixture graphs of four complexity classes: the graphs included 8 or 9 non-outgroup populations, one outgroup (all sampled at leaves), and 4 or 5 pulse-like admixture events. Demographic events were separated by date intervals ranging randomly between 1,500 and 8,000 generations, with an upper bound on the tree depth at 40,000 generations. To be more precise, demographic events were not placed in time entirely randomly, but were tied to one or few other events of the same “topological depth” within the graph, as illustrated by examples of the simulated topologies in Suppl. Fig. 8. The same principle was applied to the sampling dates for non-root groups, which were tied to other demographic events such as divergence and admixture of other populations. The random graph topologies and simulated parameter sets were generated using the *random_sim* function from the *ADMIXTOOLS 2* package: https://uqrmaie1.github.io/admixtools/reference/random_sim.html Outgroups facilitate automated exploration of graph topology space. Outgroup branches diverged from the other populations at 40,000 generation in the past and had a large constant effective population size of 100,000 diploid individuals. Other effective population sizes were constant along each edge and were picked randomly from the range of 2,000-40,000 diploid individuals. Admixture proportions for all admixture events varied randomly between 10% and 40%. The root of the simulation and the root of all non-outgroup populations were sampled, and the other populations were sampled at branch tips exclusively. This setup generates groups sampled at widely different dates in the past (from 0 to ca. 40,000 generations) or, in other words, located at various genetic distances from the root (Fig. 2d). The outgroup population was sampled at the “present” of the simulation. Sample sizes for all populations were identical: 10 diploid individuals with no missing data.

For subsequent analyses we selected only simulations where pairwise *F*_*ST*_ for groups were in the range characteristic for anatomically modern and archaic humans (in each simulation there was at least one *F*_*ST*_ value below 0.15; see Suppl. Fig. 8). In this way, 20 random topologies were simulated per complexity class. Each topology was simulated only once, and 80 simulations were generated in total (see examples of the topologies and respective *F*_*ST*_ distributions in Suppl. Fig. 8). Another set of simulations was prepared with the same topologies and parameters, except for the effective population size on the outgroup branch which was set at 1,000 diploid individuals instead of 100,000.

The following ascertainment schemes were applied to the outcomes of these randomized simulations: 1) ascertainment on sites heterozygous in a single randomly selected individual (this Human Origins-like ascertainment was repeated for all simulated groups including the outgroup and root groups, generating 920 ascertained datasets); 2) unions of four such Human Origins-like SNP panels, with only one individual per group considered (10 random sets of four groups excluding the outgroup and root groups were explored per topology, generating 800 ascertained datasets); 3) ascertainment on sites polymorphic in a group composed of three randomly selected individuals, with only one individual per group considered (10 random sets of three groups excluding the outgroup and root groups were explored per topology, generating 800 ascertained datasets); and 4) MAF ascertainment, that is restricting to sites having MAF > 5% in random meta-groups (10 random sets of four groups excluding the outgroup and root groups were explored per topology, generating 800 ascertained datasets). Group sets used for each ascertainment were recorded. Genetic distances (*F*_*ST*_) were calculated for all populations (including the outgroup and the last common ancestor of all non-outgroup populations) vs. the root sample (Fig. 2d).

Alternatively, simple trees were simulated using the *msprime v.1.1.1* settings described above. A tree of four groups conforming to the *f*_*4*_-statistic (A, B; C, O) was simulated using *msprime v.1.1.1*, with a tree depth of 4,000 generations (Suppl. Fig. 11). All the groups had a uniform effective population size of 100,000 diploid individuals, except for a bottleneck happening immediately after the A-B divergence (at 1,999 generations in the past) and lasting until the end of the simulation. The following bottleneck classes were simulated: no bottleneck (control), 10x, 100x, 1,000x, and 10,000x reduction in effective population size. For each bottleneck class, 20 independent simulations were performed. All the samples were drawn at “present”: sample sizes were 25, 25, 25 and 10 for populations A, B, C and O, respectively (except for the 10,000x bottleneck class since group A included 10 individuals only in that case). Three ascertainment schemes were tested for the simulated trees: 1) ascertainment on sites heterozygous in a single randomly selected individual (this Human Origins-like ascertainment was repeated for all simulated groups, including group O); 2) restricting to sites having MAF > 5% (or 10%, or 2.5%) in the union of groups A and B composed of 50 diploid individuals; and 3) removal of sites with *derived* allele frequency > 95% (or 90%, or 97.5%) in the union of groups A and B. The latter ascertainment scheme was added since the ascertainment schemes we tested on real data deplete the derived end of the allele frequency spectrum more than the ancestral end (Suppl. Tables 10-12).

For calculating *f*-statistics and fitting admixture graphs to unascertained and ascertained SNP sets, the *ADMIXTOOLS 2* (Maier et al. 2022 preprint) software package was used. Since there was no missing data and all individuals were diploid, we first calculated all possible *f*_*2*_-statistics for 4 Mbp-sized genome blocks (with the “*maxmiss=0*”, “*adjust_pseudohaploid=FALSE*”, and “*minac2=FALSE*” settings) and then used them for calculating *f*_*4*_-statistics as linear combinations of *f*_*2*_-statistics or for fitting admixture graphs (with the “*numstart=100*” and “*diag=0.0001*” settings). This calculation protocol was used for generating the results shown in Figs. 2 and in Suppl. Figs. 7-11. When true admixture graphs were fitted to ascertained data, full population samples of 10 individuals were used by default, and in some cases, as indicated in the figure legends, the individual used for ascertainment was used as the only representative of the respective population. Outgroups were included in the fitted graphs; in other words, they were co-analyzed with the other groups. Outgroups were not co-modelled with the other populations in Fig. 2f only.

For assessing the power of ascertained simulated datasets to reject incorrect admixture graph models, we first generated a set of such incorrect graphs per each simulated topology. For that purpose, an algorithm for finding well-fitting topologies (*findGraphs* from the *ADMIXTOOLS 2* package) was started on non-ascertained data 300 times, seeded by random graphs containing either the simulated number of admixture events (*n*, 100 runs), or *n-1* events (100 runs), or *n+1* events (100 runs). For a list of settings for the *findGraphs* algorithm see Maier *et al*. (2022 preprint). Thousands of diverse graphs explored by *findGraphs* in the process of topology optimization were generated in this way for each simulated graph, and 100 poorly fitting graphs were randomly picked from a subset of these graphs having LL scores between 70 and 300. This subset of graphs was then fitted to all ascertained datasets derived from the same simulated admixture graph.

### 2. Constructing the set of real data

We used the *cteam-lite* dataset described in Mallick *et al*. 2016, composed of the full SGDP set (300 high-coverage genomes from present-day populations), the chimpanzee genome (pseudo-haploid genotype calls, see http://hgdownload.cse.ucsc.edu/goldenPath/panTro2/bigZips/), and the Altai Neanderthal, “Denisova 3” Denisovan, Ust’-Ishim, WHG Loschbour, and LBK Stuttgart ancient genomes (see SI section 3 in that study). We supplemented *cteam-lite* by 44 present-day African genomes sequenced using the SGDP protocols by Fan *et al*. (2019), the Vindija Neanderthal’s genome (Prüfer et al. 2017), and the genome of an ancient African forager individual I10871 sequenced by Lipson *et al*. (2020) (Suppl. Table 1). Sites polymorphic in this set of 352 individuals were extracted from the *cteam-lite* files of the “hetfa” format using the *cpoly* tool (Mallick et al. 2016): alleles were grouped into derived and ancestral (polarized) according to the chimpanzee genome; missing data and heterozygous sites were allowed. For each genome, we used individual base quality masks included in *cteam-lite* or constructed using the same protocol for other genomes (Vindija Neanderthal and Fan et al. 2019): minimum base quality was set by default at 1, as recommended in SI section 3 in Mallick *et al*. 2016, which discarded lowest-quality regions marked as “0”, “?”, or “N”. The individual I10871 was not included in most analyses in this study (except for the complex admixture graphs in Suppl. Text 1) due to its relatively high rate of deamination errors.

The resulting dataset prior to missing data removal and ascertainment includes 94,691,841 autosomal sites (Suppl. Table 2). To keep the polarity of alleles, all data manipulations and ascertainments were performed using *PLINK v.2.0 alpha* (Chang et al. 2015). For calculating *f*_*4*_-statistics, sets of continental-level meta-populations were selected (e.g., Africans and East Asians or Africans and archaic humans) and then *f*_*4*_-statistics were calculated for all possible combinations of populations in the resulting subset of the “SGDP+archaic” dataset, with no missing data (at the population level) allowed *within the selected subset*. This was done to avoid potential biases associated with data missing non-randomly across groups. Alternatively, *f*_*4*_-statistics were drawn randomly from a certain class of statistics, and no missing data (at the population level) were allowed in the resulting *population quadruplets*.

### 3. Influence of ascertainment on fits of admixture graphs to real data

First, we fitted all possible graphs including two admixture events (32,745 distinct topologies with no fixed outgroup) for three combinations of groups: 1) one archaic individual, three African groups, and one African group with substantial West Eurasian-related ancestry (Altai Neanderthal, Ju|’hoan North, Biaka, Yoruba, and Agaw, respectively);

2) five deeply divergent ancient and present-day non-African groups (Ust’-Ishim, Papuan, Onge, LBK Stuttgart, Even); and 3) five deeply divergent present-day non-African groups (Papuan, Onge, Palestinian, Even, Mala). These three sets of simple graphs were fitted to all sites, AT/GC sites, and 1240K sites (no missing data were allowed at the group level within these sets of five populations); 5,000 best-fitting models were selected according to the LL score on all sites and WRs of those models were compared across SNP sets (Suppl. Fig. 1).

Next, we explored the same exhaustive set of admixture graph topologies including five groups and two admixture events on the wider collection of ascertainments. Twelve combinations of five groups including up to two archaic humans, up to five African groups, and up to five non-African groups were tested. To ensure fair comparison across at least a subset of population combinations, as a starting point for generating ascertained site sets we used either 11,706,773 sites (with no missing data at the group level) polymorphic in a set of 48 archaic and African groups composed of a total of 97 individuals; or 10,051,585 such sites in a set of 59 archaic, African, European, and Middle Eastern groups composed of a total of 120 individuals; or 5,296,653 such sites in a set of 51 Papuan, Native American, European, Anatolian, and Caucasian groups composed of a total of 112 individuals (Suppl. Tables 1 and 2).

We examined the fits of these collections of admixture graphs from different perspectives. (1) We considered just 5,000 topologies that are best-fitting on the unascertained site set (Suppl. Figs. 1-3) or all 32,745 topologies tested (Suppl. Fig. 4). (2) We also considered alternative admixture graph fit metrics, LL or WR. LL as a fit metric (see left-hand panels in Suppl. Figs. 3 and 4) is more accurate than WR, but difficult to compare across population sets. Finally, instead of *R* or *R*^*2*^ of a linear trend as a measure of correlation of admixture graph fits (Fig. 1, Suppl. Figs. 1-4) we considered the fraction of all possible models of a certain complexity that are rejected under ascertainment (WR > 3 SE) but accepted on all sites (WR < 3 SE), or *vice versa*.

### 4. Automated inference of fitting admixture graphs on real data

The 12-population admixture graph published by Lipson et al. 2020 (and later used as a skeleton graph in Lipson et al. 2022) and simpler 7- and 10-population intermediate graphs presented in the former study were revisited by Maier *et al*. (2022 preprint), and thousands of alternative well-fitting graphs of the same complexity were found using the *find_graphs* function from the *ADMIXTOOLS 2* package (https://uqrmaie1.github.io/admixtools/articles/graphs.html). Maier *et al*. used the 1240K dataset only, and in the current study we re-fitted the admixture graphs found by the algorithm on the 1240K SNP panel to the AT/GC and unascertained datasets derived from “SGDP+archaic” and also repeated automated admixture graph inference on these two additional SNP sets. Advantages and pitfalls of automated admixture graph inference are described in detail in Maier *et al*., along with justifications for the specific protocol used in that study, and here we used protocols identical to those employed by Maier *et al*. We first calculated all possible *f*_*2*_-statistics for 4 Mbp-sized genome blocks (with the “*maxmiss=0*”, “*adjust_pseudohaploid=FALSE*”, and “*minac2=2*” settings, see Maier *et al*. 2022 for details on the settings) and then used them for fitting admixture graphs (with the “*numstart=100*” and “*diag=0.0001*” settings) and for automated admixture graph inference with the *find_graphs* function (see the Methods section in Maier *et al*. for a complete list of arguments for this function). Only one topology constraint was used at the graph space exploration step: chimpanzee was assigned as an outgroup.

## Acknowledgements

The authors are grateful to Mark Lipson and Nick Patterson for discussions. P.F., U.I., and P.C. were supported by the Czech Ministry of Education, Youth and Sports (program ERC CZ, project no. LL2103). P.F. and P.C. were supported by the Czech Science Foundation (project no. 21-27624S). P.F. was also supported by a subsidy from the Russian federal budget (project No. 075-15-2019-1879 “From paleogenetics to cultural anthropology: a comprehensive interdisciplinary study of the traditions of the peoples of transboundary regions: migration, intercultural interaction and worldview”). This research was funded by NIH grant HG012287, by the Allen Discovery Center program, a Paul G. Allen Frontiers Group advised program of the Paul G. Allen Family Foundation, by John Templeton Foundation grant 61220, by private gifts from Jean-Francois Clin to D.R. and P.F., and by the Howard Hughes Medical Institute. Computational resources for this work were supplied by the projects “e-Infrastruktura CZ” (e-INFRA CZ LM2018140) and “IT4Innovations National Supercomputing Center – LM2015070” supported by the Ministry of Education, Youth and Sports of the Czech Republic.

