## Supplementary Information for "Modeling of African population history using *f*-statistics can be highly biased and is not addressed by previously suggested SNP ascertainment schemes"

1. Effects of ascertainment on real data: fits of complex admixture graphs

Lipson *et al*. (2020) constructed a complex admixture graph for Africans, including the Altai Neanderthal and chimpanzee. The model was built manually on a dataset derived from the 1240K panel, and a final model was also confirmed to fit the union of the Human Origins sub-panels 4 and 5 or archaic-ascertained sites. We aimed at reassessing these results in the light of *f*-statistic biases and took advantage of a new algorithm for automated search of the admixture graph space, *findGraphs* (Maier et al. 2022 preprint). Since the final model in that study is rather complex (12 populations including chimpanzee as an outgroup and 12 admixture events, which results in a space of up to $1.22*10^{42}$ topologies), we focused on a simpler intermediate model shown in that study. The simpler model consists of 10 groups and 8 admixture events (Lipson et al. 2020, Fig. S3.25 in that study). The corresponding space of all possible graphs is still impressively large: up to $1.9*10^{26}$ topologies. To enable comparison of various ascertainment schemes on whole-genome shotgun data, the population composition of the dataset we worked on was slightly different from the dataset used by Lipson *et al.*: the ancient South African hunter-gatherer group was replaced by a related group, present-day Ju|'hoan North, and instead of the Shum Laka ancient group from Cameroon only one shotgun-sequenced individual from that group (I10871) was used.

We began by fitting the published 10-population model with 8 admixture events to three SNP sets without missing data at the group level: 1) 1240K, ca. 839,000 polymorphic sites; 2) AT/GC mutation types, ca. 3.8 million polymorphic sites; and 3) "all sites", ca. 25 million polymorphic sites. The published model had widely different fits on these three datasets (Suppl. Fig. 5a): WR of the published admixture graph model was 2.7 SE on the 1240K dataset, 4.8 SE on AT/GC sites, and 8.4 SE on all sites.

Next, we attempted to find alternative models in the same complexity class (i.e., 10 groups and 8 admixture events) and investigated admixture graph fits (LL and WR) across the three datasets for all distinct newly found topologies. The *findGraphs* topology search algorithm was started 10,000 times from random graphs with chimpanzee set as an outgroup, and that procedure was repeated for each SNP set. For simplicity, only one graph with the best LL was taken from each *findGraphs* run. The fits of the resulting collections of ca. 10,000 distinct topologies are summarized in Suppl. Fig. 5b. As shown in Fig. 1, Table 1, and Suppl. Fig. 1 for much simpler graphs, model fits on the 1240K panel and on all sites are correlated poorly (Pearson's *R* for LL ranging from 0.35 to 0.46), and the fits on AT/GC sites and on all sites are correlated better (*R* for LL ranging from 0.58 to 0.73, Suppl. Fig. 6a). Among thousands of topologies inferred on the 1240K dataset, 63% fit the 1240K data well (WR < 3 SE), but all of them do not fit the unascertained dataset (WR from 4.6 to 21 SE, Suppl. Fig. 6b). The converse analysis also reveals concerns: of 24 topologies found that fit the unascertained dataset relatively well (WR between 3 and 4 SE), 18 topologies fit the 1240K dataset worse (WR > 4 SE) despite its much smaller size, while 6 topologies fit both datasets with WR between 3 and 4 SE. These results are in line with those for the simpler graphs shown in Fig. 1 and Suppl. Fig. 1, where classes of topologies that fit the unascertained data and do not fit the ascertained data by a wide margin (and *vice versa*) are highlighted. Similar results were obtained for a smaller published intermediate graph (Lipson et al. 2020, Fig. S3.24) with 7 groups and 4 admixture events (Suppl. Fig. 6c,d).

*2. An overview of $f_4$-statistic biases caused by ascertainment*

The following group of $f_4$-statistics was explored exhaustively: statistics including Africans (unadmixed with West Eurasians according to Fan *et al*. 2019, see group annotations in Suppl. Table 1) and/or archaic humans and/or Mediterranean/Middle Eastern groups (abbreviated as "Med/ME"). This group of ca. 1.37 million statistics was subdivided into 14 classes depending on the composition of the population quadruplet: for instance, all statistics including 3 African groups and 1 archaic individual; or 2 African groups, 1 archaic individual and 1 Med/ME group, etc. The following notation is used for referring to these classes: African$_3$;archaic$_1$, etc. In the analyses below we focused on $f_4$-statistics that do not deviate from 0 too far (absolute Z-score on all sites is < 15 SE) since in practice it is often important if a statistic is consistent with 0 on a certain set of sites but deviates from 0 by > 3 SE on another dataset, and especially if it changes its sign.

Several $f_4$-statistic classes were affected by ascertainment, as illustrated by distributions of absolute Z-score differences (Suppl. Fig. 13) and by scatterplots of Z-scores (Suppl. Fig. 14). The unascertained dataset (all sites) was used as a baseline in all cases. By far the worst performing $f_4$-statistic class was African$_2$;archaic$_1$;Med/ME$_1$, and specifically statistics $f_4$(African X, archaic; African Y, Med/ME). All statistics belonging to this sub-class are biased in the case of the 1240K ascertainment since none of them lie on the diagonal (Fig. 5), and all 8 non-random ascertainment schemes explored, including archaic ascertainment schemes, demonstrated similar patterns (Suppl. Fig. 13). This observation is in line with the results on simulated data where statistics $f_4$("African 1", "archaic"; "African 2", "non-African") are affected by all types of ascertainments tested in the presence of an archaic to non-African gene flow (Suppl. Fig. 7b).

The following three $f_4$-statistic classes were also affected by bias, but were less problematic according to the estimated standard deviation of residuals from a linear model (termed "residual SE" for brevity and expressed in the same units as $f_4$-statistic Z-scores): 1) African$_3$;archaic$_1$, 2) African$_3$;Med/ME$_1$ (Fig. 5), 3) African$_4$ (Suppl. Figs. 13 and 14). Notably, removal of variants rare in Africans helped to mitigate the bias in the case of the latter three classes almost completely (Fig. 5, Suppl. Figs. 13 and 14). Archaic ascertainment helped to shift modes of the |Z-score difference| distributions towards 0 for all the classes except for $f_4$(African X, archaic; African Y, Med/ME), however long distribution tails, i.e. a high noise level, observed for 7 $f_4$-statistic classes such as African$_3$;archaic$_1$ and African$_4$ (Suppl. Figs. 13 and 14) suggest that this ascertainment scheme is performing worse in practice than the pan-African ascertainment.

Next, we demonstrated that among ca. 63,000 randomly selected statistics $f_4$(African X, archaic; African Y, non-African), all deviate from the diagonal by at least 2 SE under the 1240K ascertainment (Suppl. Fig. 15; all continental-scale populations of non-Africans in the SGDP dataset were tested, including such groups as Papuans and the Upper Paleolithic Ust'-Ishim individual). We also demonstrated on exhaustive collections of $f_4$-statistics that do not include archaic individuals or African groups only (statistics African$_x$;East Asian$_y$(y>0) pooled with American$_x$;European$_y$;Papuan$_z$) that the effect of the pan-African ascertainment is not distinguishable from that of random thinning (Suppl. Fig. 16). In contrast, the 1240K ascertainment, Human Origins panels 4, 5, 13, 4+5, and archaic ascertainment resulted in a fraction of $f_4$-statistic Z-scores deviating from the diagonal by > 2 SE (Suppl. Fig. 16).

According to our mass comparison of ascertainment schemes on 27 $f_4$-statistic classes (Suppl. Table 7), the pan-African ascertainment outperformed all other ascertainment schemes explored, including archaic ascertainment, and the same result was observed when exploring admixture graph model fits (Table 1, Suppl. Fig. 3, Suppl. Tables 3-5). For this comparison we disregarded $f_4$-statistics with |Z| > 15 SE on all sites, and thus all extreme outliers in the $f_4$(Neanderthal, X; Denisovan, Y) statistic classes that emerge due to the fact that the same pair of individuals, Neanderthal and Denisovan, were used both for ascertainment and for calculating *f*-statistics, as we show on simulated (Fig. 2b, Suppl. Fig. 18c) and real data (Suppl. Fig. 18a). These outliers (i.e., statistics changing their sign from highly positive to highly negative, Suppl. Fig. 18) serve as another argument in favor of pan-African ascertainment over archaic ascertainment (see, e.g., Suppl. Fig. 12a) since the former is applicable to a wider range of population sets. Due to the paucity of high-coverage archaic genomes (Meyer et al. 2012, Prüfer et al. 2014, Mafessoni et al. 2020), it is often unavoidable that the same individuals are used for both ascertainment and for calculating *f*-statistics or for fitting admixture graphs.

On simulated data we focused on two types of $f_4$-statistics that, as we found empirically, are most affected by ascertainment: $f_4$("African 1", "archaic individual"; "African 2", "non-African 1 or 2") (Suppl. Fig. 7b) and $f_4$("non-African 1", "archaic individual"; "African 1 or 2", "non-African 2") (Suppl. Fig. 7c). In the absence of Neanderthal gene flow to non-Africans, the Z-scores of the former statistics are not affected substantially by any ascertainment scheme we applied: Z-scores across nearly all simulation iterations, population combinations, and ascertainment schemes remain significantly negative (< -3 SE). However, in the presence of the Neanderthal gene flow the *Z*("African 1", "archaic individual"; "African 2", "non-African 1 or 2") are shifted from significantly positive to nearly 0, or from 0 to significantly negative under the following ascertainment schemes: Human Origins (one panel ascertained on an individuals from the "African 2" group), African MAF, and archaic (Suppl. Fig. 7b).

*3. Mechanisms of bias in selected $f_4$-statistics*

In order to understand mechanisms underlying the biases that we documented several individual $f_4$-statistics were analyzed in detail. We considered a statistic of the most biased class $f_4$(African X, archaic; African Y, non-African), specifically $f_4$(Altai Neanderthal, Biaka; Mbuti, Saharawi) (Saharawi is an African group with a low proportion of sub-Saharan African ancestry, Fan et al. 2019). In Suppl. Table 9 $f_4$-statistic values and Z-scores are shown for the unascertained site set and across nine ascertainment schemes.

In Suppl. Table 10 the statistic $f_4$(Altai Neanderthal, Biaka; Mbuti, Saharawi) is explored across the derived allele frequency (DAF) spectrum and on six site sets: 1) all sites; 2) AT/GC mutation classes; 3) 1240K; 4) global MAF > 5%; 5) MAF > 5% across Africans (unadmixed with non-Africans, Suppl. Table 1); and 6) ascertainment on the three archaic individuals (all sites). Alleles were classified as derived or ancestral according to the pseudohaploid chimpanzee genome, and DAF was defined either on the African meta-population (unadmixed with non-Africans) or on various non-African meta-populations. Since results were similar for those non-African meta-populations, results for the European meta-population only are shown in Suppl. Table 10. For simplicity, sites were stratified by DAF into three bins: nearly fixed ancestral (DAF <= 5%), intermediate frequency (DAF 5-95%), and nearly fixed derived (DAF >= 95%). No sites with missing allele frequencies were allowed at the group level in Altai Neanderthal, Biaka, Mbuti, Saharawi and at the meta-population level in Africans.

First, we consider results on all sites and on AT/GC mutation classes, which are nearly identical (Suppl. Table 10). As expected for a relatively complex demographic history with gene flows and bottlenecks (Martin and Amos 2020), the statistic $f_4$(Altai Neanderthal, Biaka; Mbuti, Saharawi) is highly variable across the three DAF bins (Suppl. Table 10). On sites with intermediate DAF in Africans, the statistic is highly positive, and on much more numerous sites with nearly fixed ancestral or derived alleles the statistic is mildly negative. Since sites with nearly fixed ancestral or derived variants predominate in the unascertained dataset (Suppl. Table 10), the statistic approaches 0 on all sites (Suppl. Table 9). All the types of non-random ascertainment explored here increase dramatically the proportion of sites with intermediate DAF: it reaches 7.8% for all sites or AT/GC sites and varies between 22.5% and 98.7% for non-randomly ascertained sites. The proportion of nearly fixed sites, especially those with nearly fixed derived alleles, drops dramatically under non-random ascertainment: from 28-31% to 0.1-8.4% (Suppl. Table 10). Since the statistic $f_4$(Altai Neanderthal, Biaka; Mbuti, Saharawi) is highly variable across the DAF spectrum, discarding a great majority of nearly fixed sites shifts its value. As we showed on simulated data, the fact that archaic human lineages do not represent a true outgroup for AMH makes the class of statistics $f_4$(African X, archaic; African Y, non-African) especially problematic, i.e., biased under all ascertainment types tested in this study (Figs. 2b and 4a).

But distortion of the DAF spectrum is not the only effect that non-random ascertainment schemes have. Within each DAF bin, ascertained sites and all sites show different $f_4$-statistic values (Suppl. Table 10). For instance, in the intermediate bin (DAF in Africans) the statistic $f_4$(Altai Neanderthal, Biaka; Mbuti, Saharawi) equals 0.0064 for the

1240K sites and 0.0021 for all sites (Z-scores = 14 and 6, respectively). The same is true for the nearly fixed bins and for DAF based on the European meta-population (Suppl. Table 10). Ascertainment on global MAF in the "SGDP+archaic" dataset (removal of variants that are rare across all 350 individuals, i.e., that have MAF < 5%) produces a comparable shift in the same direction (Suppl. Table 10). Among individuals in the SGDP dataset, 73% are non-Africans (Suppl. Table 1), and variants common in non-Africans are favored by this ascertainment scheme. In other words, the global MAF ascertainment does not shift average DAF across all four populations involved in the statistic in the same way: both average $DAF_{\textit{Altai-Biaka}}$ and especially average $DAF_{\textit{Mbuti-Saharawi}}$ in the intermediate DAF bin move in the negative direction under ascertainment, which reflects relative paucity of derived variants in the Neanderthal and relative excess of derived variants in Saharawi under this ascertainment (Suppl. Table 10).

As stated above, similar effects are observed under the 1240K ascertainment (Suppl. Table 10), and we argue that the same explanation holds for the global MAF and 1240K ascertainment schemes. Although ca. 59% of sites on the Human Origins array were ascertained on African individuals (San, Yoruba, and Mbuti), the 1240K panel is a complex construct where approximately half of sites are derived from the Illumina 650Y and Affymetrix 50k arrays (Fu et al. 2015), themselves products of complex ascertainment based mostly on Eurasian populations. Thus, derived variants common in non-Africans but rare in Africa and in archaic humans are overrepresented on the 1240K panel, which skews $f_4$-statistics within the DAF bins. In contrast, under archaic ascertainment average $DAF_{\textit{Altai-Biaka}}$ moves in the positive direction in the intermediate and "nearly fixed ancestral" bins, and those bins account for >90% of ascertained sites in this case (Suppl. Table 10). Archaic ascertainment by definition favors derived variants common in archaic humans but does not favor derived variants common in AMH, thus $DAF_{\textit{Altai-Biaka}}$ becomes highly positive in these bins, and the resulting $f_4$-statistic becomes highly negative (Suppl. Tables 9 and 10).

We also considered statistics that are among the most distant outliers in two other biased classes: $f_4$(Fulani, Ju|'hoan North; Igbo, Ogiek) of the $\text{African}_{\textit{4}}$ class, and $f_4$(Burmese, Dinka; Ju|'hoan North, Sengwer) of the $\text{African}_{\textit{3}}$;$\text{East Asian}_{\textit{1}}$ class. These two statistics show similar patterns: the bias in Z-scores detected under the 1240K and global MAF ascertainment is much smaller or non-existent under the pan-African and archaic ascertainments (Suppl. Table 9) despite the depletion of nearly fixed derived sites common for all these datasets (Suppl. Tables 11 and 12). In the case of the statistics $f_4$(Fulani, Ju|’hoan North; Igbo, Ogiek) and $f_4$(Burmese, Dinka; Ju|’hoan North, Sengwer), the 1240K and global MAF ascertainments also have very similar effects within DAF bins, especially within the "nearly fixed ancestral" bin where both statistics move in the negative direction (Suppl. Tables 11 and 12). A smaller effect with the same direction is observed in the intermediate DAF bin. Thus, the shift in statistics can be explained by unequal enrichment for derived variants across the four populations. Since the African MAF and archaic ascertainment schemes do not favor derived variants of non-African origin, the bias is much smaller or non-existent in those cases (Suppl. Table 9).

*4. Archaic ascertainment explored on simulated genetic data*

We simulated various scenarios of gene flow between archaic humans and AMH. Whole human genomes evolving via mutation, recombination, and drift were simulated using *msprime* v.0.7.4 (Kelleher et al. 2016). We simulated five different topologies that are not intended to cover the whole diversity of possible topologies, and different gene flow intensities were tested for all the topologies (see a complete list of parameters tested in Suppl. Figs. 17b and 18b). We also varied the time and intensity of the out-of-Africa bottleneck (Suppl. Figs. 17b and 18b). Each simulation included "chimpanzee", "Denisovan" and "Neanderthal" populations composed of one sampled diploid individual each, five paraphyletic "African" groups (10 diploid individuals sampled per group) with intra-African gene flows, and three monophyletic "non-African" groups (10 diploid individuals sampled per group) connected with gene flows too (see detailed illustrations of model parameters in Suppl. Figs. 17b and 18b and in Suppl. Table 13). The simulated African and non-African groups diverged 100 kya (given a generation time of 25 years). For all simulated histories, SNPs polymorphic in a group composed of the Neanderthal and Denisovan individuals (two individuals in total) were taken as the ascertained set. $f_4$-statistics including the same Neanderthal and Denisovan individuals, and those including at least one of these archaic individuals and chimpanzee were calculated for the simulated groups, and results for four simulated histories are shown in Suppl. Figs. 17c and 18c.

Under the simplest simulation scenario (labelled as "model 1 pArc:NA pnAfr:0%"), with no gene flow from the Neanderthal lineage to the common ancestor of non-Africans, several classes of statistics that can be described as $f_4$(Neanderthal, X; Denisovan, Y) change their sign under archaic ascertainment from highly positive to highly negative (Suppl. Fig. 18c). The bias here emerges due to the ascertainment procedure itself since the same pair of individuals, Neanderthal and Denisovan, were used both for ascertainment and for calculating *f*-statistics. Introducing a single archaic gene flow to non-Africans (3% of their ancestry derived from the Neanderthal branch at 70 kya) affects the patterns of outliers under archaic ascertainment (Suppl. Fig. 18c, "model 1 pArc:NA pnAfr:3%"), and it becomes more similar to that observed on real data (Suppl. Fig. 18a). $f_4$-statistics calculated on real data (AT/GC sites vs. archaic ascertainment) did not include Papuans and Australians since results for those populations would be affected by Denisovan admixture that was not simulated; they also do not include Africans having substantial non-African admixture (Suppl. Table 1), to make distinction between $f_4$-statistic classes clearer. Few differences from the pattern observed on real data are as follows: 1) statistics $f_4$(Neanderthal, African; chimpanzee, non-African) change their sign on simulated data but do not change their sign on real data; 2) all statistics $f_4$(Neanderthal, non-African; Denisovan, non-African) change their sign on simulated data, but some do not change their sign on real data (Suppl. Fig. 18a). Next, we simulated a model of archaic introgression proposed by Durvasula and Sankararaman (2020) with 4 relevant gene flows (labelled as "model 2"). Here we tested 7 combinations of super-archaic ancestry proportion in AMH and Neanderthal ancestry proportion in non-Africans, in addition to four settings for the out-of-Africa bottleneck (Suppl. Fig. 18b). For large proportions of super-archaic ancestry in AMH (19%, the upper bound found in the original study, or 25%), the results on simulated and real data differed for more than two classes of statistics (Suppl. Fig. 18c). For a small proportion of super-archaic ancestry in AMH (2%, the lower bound found in the original study), the results on simulated (Suppl. Fig. 18c) and real data (Suppl. Fig. 18a) differ for two classes of statistics: statistics $f_4$(Neanderthal, African; chimpanzee, non-African) change their sign on simulated data under archaic ascertainment but do not show this behavior on real data; and statistics $f_4$(Neanderthal, X; Denisovan, chimpanzee) and $f_4$(Neanderthal, chimpanzee; Denisovan, X) form two distant clusters on simulated data, but are clustered closely together on real data (Suppl. Fig. 18c, "model 2, PArc 2%, PnAfr 3%").

Next, we tested three alternative simulated graphs, outcomes of an automated admixture graph inference procedure on real data (AT/GC sites, see Methods for a description of the *findGraphs* protocol). The best-scoring topology on real data (labelled as "model 3") includes two gene flows from a divergent Neanderthal-related lineage (splits from the Neanderthal lineage *sensu stricto* at 350 kya in our simulations): a flow into the common ancestor of AMH at 250 kya and another flow into the common ancestor of non-Africans at 70 kya (Suppl. Fig. 18b). If both gene flow proportions remain low (3%), this topology reproduces the pattern of outliers observed on real data closely (Suppl. Fig. 18c, "model 3, PArc 3%, PnAfr 3%"): statistics $f_4$(Neanderthal, African; chimpanzee, non-African) do not change their sign, and statistics $f_4$(Neanderthal, X; Denisovan, chimpanzee) and $f_4$(Neanderthal, chimpanzee; Denisovan, X) cluster together. However, results for one class of statistics still disagree on simulated (Suppl. Fig. 18c) and real data (Suppl. Fig. 18a): many statistics $f_4$(X, Y; chimpanzee, Denisovan) change their sign on real data, but do not do that on simulated data. We also tested two other topologies that emerged among the best outcomes of the automated graph inference procedure on real data, labelled as "model 4" and "model 5" (Suppl. Fig. 18b), and patterns of outliers differ from those observed on real data for several classes of $f_4$-statistics.

Our observations on a narrow region of parameter space suggest that neither the simplest model of archaic introgression in AMH, i.e., the low-level pulse-like gene flow from Neanderthals to non-Africans, nor the more complex model proposed by Durvasula and Sankararaman (2020), are able to predict the behavior of $f_4$-statistics under archaic ascertainment. These results on simulated data also suggest that since archaic humans do not represent a true outgroup for AMH (due to various gene flows between archaic humans and AMH), archaic ascertainment should be used with caution for reconstructing population history of AMH.

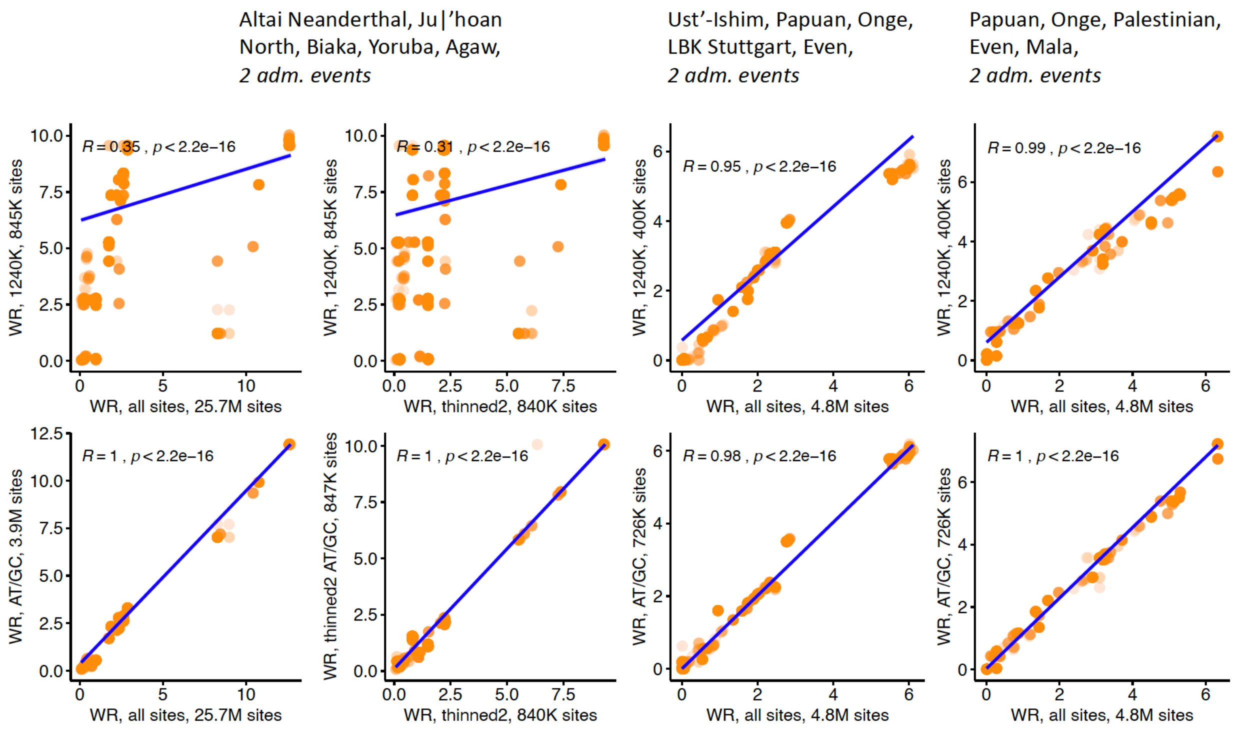

**Suppl. Fig. 1.** Scatterplots illustrating the effects of the 1240K ascertainment on LL and WR for exhaustive collections of simple admixture graphs. Five thousand best-fitting graphs (according to LL on all sites) of 32,745 possible graphs were selected for each combination of populations, and correlation of WRs was explored for graphs fitted on all sites and on ascertained datasets. Results are shown for three population combinations indicated in plot titles. On x-axes results for all sites or for a randomly thinned site set are shown. Results for the 1240K ascertainment are shown in the upper row on the y-axes, and results for AT/GC sites are shown in the lower row on the y-axes. Linear trends fitted to the plotted points are shown in blue, along with Pearson correlation coefficient.

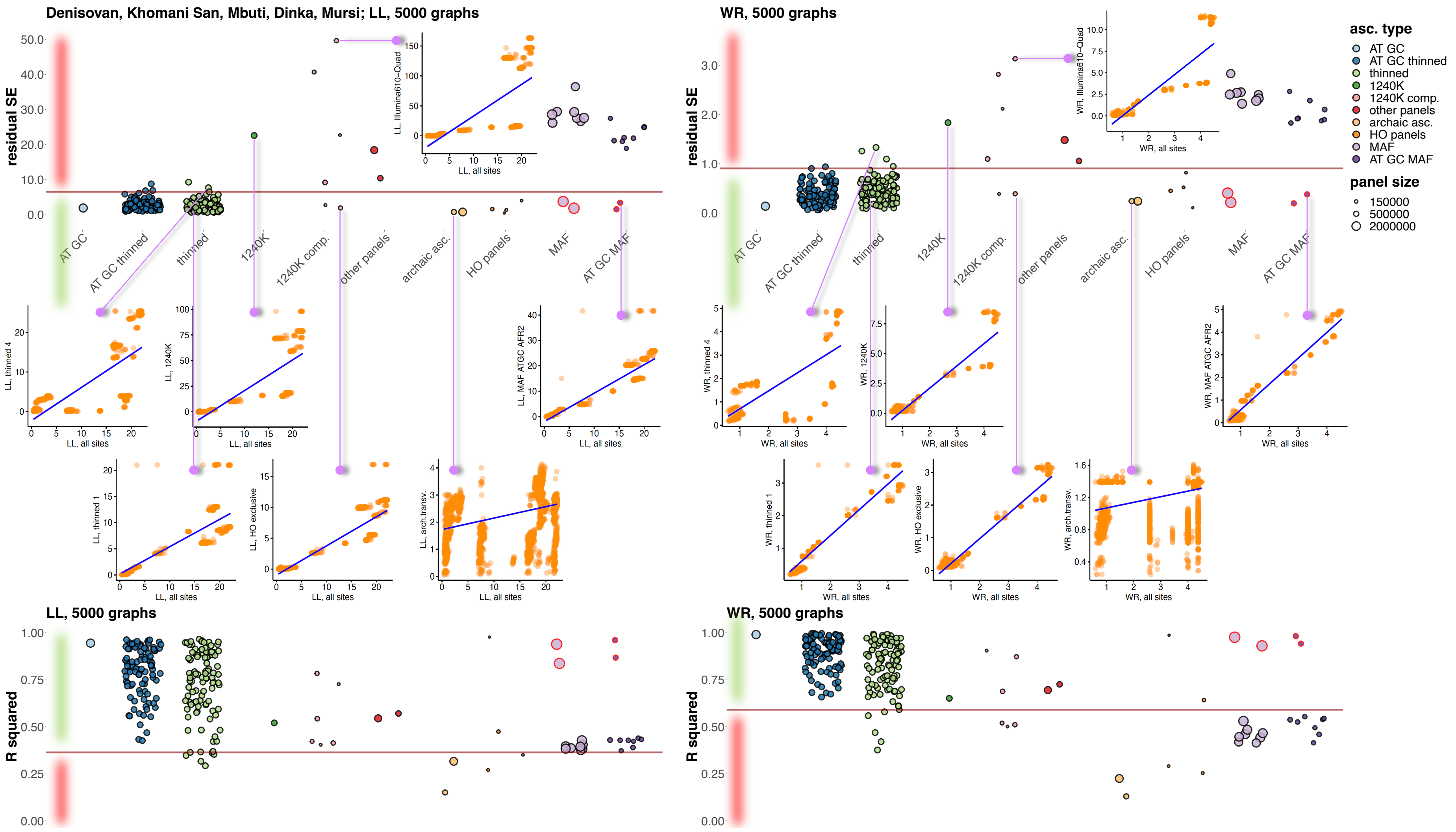

**Suppl. Fig. 2.** Two alternative approaches for visualizing the effect of ascertainment bias on admixture graph fits illustrated using one population combination, “Denisovan, Khomani San, Mbuti, Dinka, Mursi”. On top, residual SE of linear trends are shown for various ascertainment schemes, and at the bottom $R^2$ of linear trends are shown. Five thousand best-fitting graphs (according to LL on all sites) of 32,745 possible graphs were selected, and correlation of LL (left-hand panels) or WR (right-hand panels) was explored for graphs fitted on all sites and on ascertained datasets. Results for ascertainment on variants common in Africans (either those having no detectable West Eurasian ancestry or all Africans in the SGDP dataset) are circled in red. As a starting point for generating different ascertainments we used 11,706,773 sites (with no missing data at the group level) polymorphic in a set of 48 archaic and African groups composed of 97 individuals (Suppl. Table 1). Thirty eight site subsampling schemes were explored: 1) AT/GC mutation classes; 2) random thinning of the AT/GC dataset to the 1240K SNP count for a given combination of groups (no missing data allowed), results for 100 thinned replicates are shown; 3) random thinning of all sites to the 1240K SNP count, results for 100 thinned replicates are shown; 4) the 1240K SNP panel; 5) major components of the 1240K panel: sites included in the Illumina 650Y and/or Human Origins SNP arrays, sites included exclusively in one of them, and remaining sites; 6) the 1000K and 2200K SNP panels; 7) restricting to sites polymorphic in a group composed of the three high-coverage archaic individuals (either all such sites or transversions only); 8) the largest Human Origins sub-panels (4, 5, 13) or their union (4+5); 9) restricting to common variants based on a global MAF threshold of 5% or on the same threshold in one of nine continental-scale groups; 10) the same procedure repeated on AT/GC sites. The size of the resulting SNP panels is coded by point size, and ten broad ascertainment types are coded by color according to the legend. The 97.5$^{th}$ (in the case of residual SE) or 2.5$^{th}$ (in the case of $R^2$) LL or WR percentiles of all the thinned replicates combined, including those on all sites and AT/GC sites, are marked by brown lines. Areas of the plots where ascertainments are considered biased according to these thresholds are highlighted in red on the left-hand side of the plots. Scatterplots illustrating effects of selected ascertainment schemes on LL or WR are shown in the middle of the figure and are connected to the respective data points (ascertainments) by magenta lines. Each dot on these scatterplots corresponds to a distinct admixture graph topology.

a

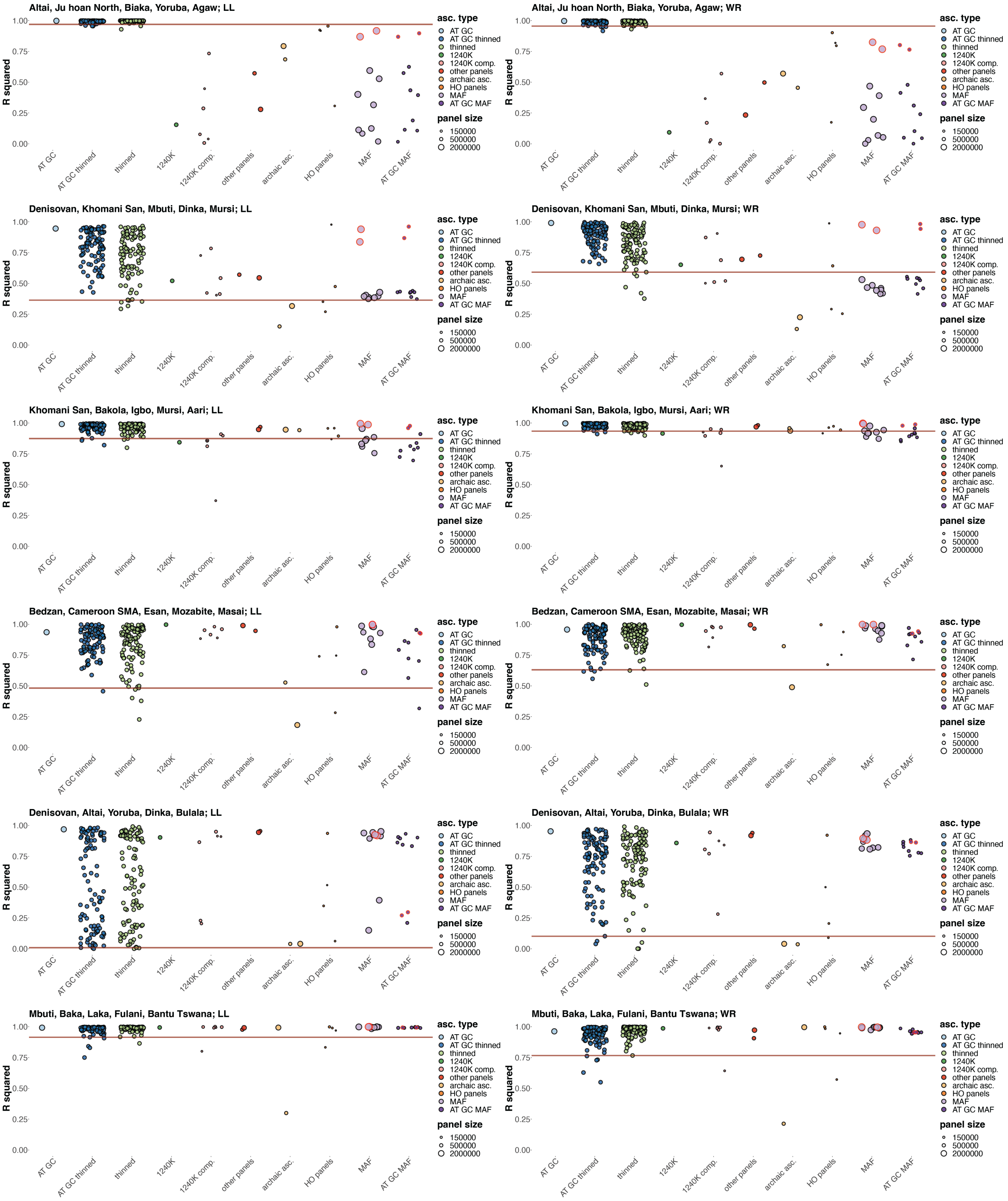
Altai, Ju hoan North, Biaka, Yoruba, Agaw; LL
Altai, Ju hoan North, Biaka, Yoruba, Agaw; WR
Denisovan, Khomani San, Mbuti, Dinka, Mursi; LL
Denisovan, Khomani San, Mbuti, Dinka, Mursi; WR
Khomani San, Bakola, Igbo, Mursi, Aari; LL
Khomani San, Bakola, Igbo, Mursi, Aari; WR
Bedzan, Cameroon SMA, Esan, Mozabite, Masai; LL
Bedzan, Cameroon SMA, Esan, Mozabite, Masai; WR
Denisovan, Altai, Yoruba, Dinka, Bulala; LL
Denisovan, Altai, Yoruba, Dinka, Bulala; WR
Mbuti, Baka, Laka, Fulani, Bantu Tswana; LL
Mbuti, Baka, Laka, Fulani, Bantu Tswana; WR
R squared
1.00
0.75
0.50
0.25
0.00
AT GC
AT GC thinned
thinned
1240K
1240K comp.
other panels
archaic asc.
HO panels
MAF
AT GC MAF
asc. type
AT GC
AT GC thinned
thinned
1240K
1240K comp.
other panels
archaic asc.
HO panels
MAF
AT GC MAF
panel size
150000
500000
2000000

**b**

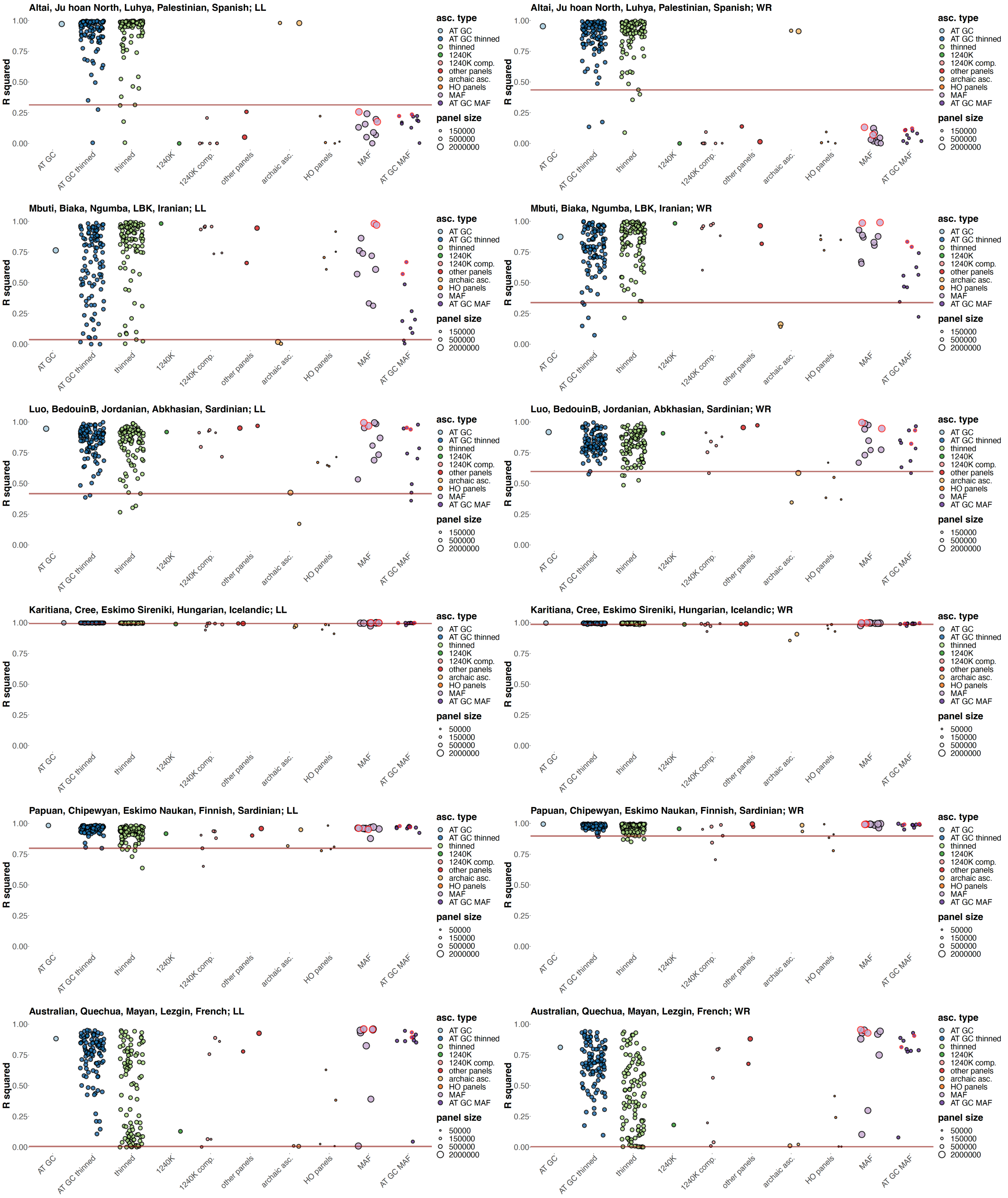
Altai, Ju hoan North, Luhya, Palestinian, Spanish; LL
Altai, Ju hoan North, Luhya, Palestinian, Spanish; WR
Mbuti, Biaka, Ngumba, LBK, Iranian; LL
Mbuti, Biaka, Ngumba, LBK, Iranian; WR
Luo, BedouinB, Jordanian, Abkhasian, Sardinian; LL
Luo, BedouinB, Jordanian, Abkhasian, Sardinian; WR
Karitiana, Cree, Eskimo Sireniki, Hungarian, Icelandic; LL
Karitiana, Cree, Eskimo Sireniki, Hungarian, Icelandic; WR
Papuan, Chipewyan, Eskimo Naukan, Finnish, Sardinian; LL
Papuan, Chipewyan, Eskimo Naukan, Finnish, Sardinian; WR
Australian, Quechua, Mayan, Lezgin, French; LL
Australian, Quechua, Mayan, Lezgin, French; WR
R squared
1.00
0.75
0.50
0.25
0.00
AT GC
AT GC thinned
thinned
1240K
1240K comp.
other panels
archaic asc.
HO panels
MAF
AT GC MAF
asc. type
AT GC
AT GC thinned
thinned
1240K
1240K comp.
other panels
archaic asc.
HO panels
MAF
AT GC MAF
panel size
50000
150000
500000
2000000

**Suppl. Fig. 3.** Variance in fits of a collection of simple admixture graphs (five groups and two admixture events) resulting from ascertainment or random site subsampling expressed as $R^2$ of linear trends. Five thousand best-fitting graphs (according to LL on all sites) of 32,745 graphs were selected for each combination of populations, and correlation of LL or WR was explored for graphs fitted on all sites and on ascertained datasets. Results are shown for twelve population combinations indicated in plot titles (panels **a, b**). Results for ascertainment on variants common in Africans (either those having no detectable West Eurasian ancestry or on all Africans in the SGDP dataset) are circled in red. As a starting point for generating different ascertainments, we used either 11,706,773 sites (with no missing data at the group level) polymorphic in a set of 48 archaic and African groups composed of 97 individuals, or 10,051,585 such sites in 59 archaic, African, European, and Middle Eastern groups composed of 120 individuals, or 5,296,653 such sites in 51 Papuan, Native American, European, Anatolian, and Caucasian groups composed of 112 individuals (Suppl. Table 1). Thirty eight site subsampling schemes were explored: 1) AT/GC mutation classes; 2) random thinning of the AT/GC dataset to the 1240K SNP count for a given combination of groups (no missing data allowed), results for 100 thinned replicates are shown; 3) random thinning of all sites to the 1240K SNP count, results for 100 thinned replicates are shown; 4) the 1240K SNP panel; 5) major components of the 1240K panel: sites included in the Illumina 650Y and/or Human Origins SNP arrays, sites included exclusively in one of them, and remaining sites; 6) the 1000K and 2200K SNP panels; 7) restricting to sites polymorphic in a group composed of the three high-coverage archaic individuals (either all such sites or transversions only); 8) the largest Human Origins sub-panels (4, 5, 13) or their union (4+5); 9) restricting to common variants based on a global MAF threshold of 5% or on the same threshold in one of nine continental-scale groups; 10) the same procedure repeated on AT/GC sites. The size of the resulting SNP panels is coded by point size, and ten broad ascertainment types are coded by color according to the legend. $R^2$ values for LL are plotted in the left-hand panels, and values for WR are plotted in the right-hand panels. The 2.5$^{th}$ LL or WR percentiles of all the thinned replicates combined, including those on all sites and AT/GC sites, are marked by brown lines.

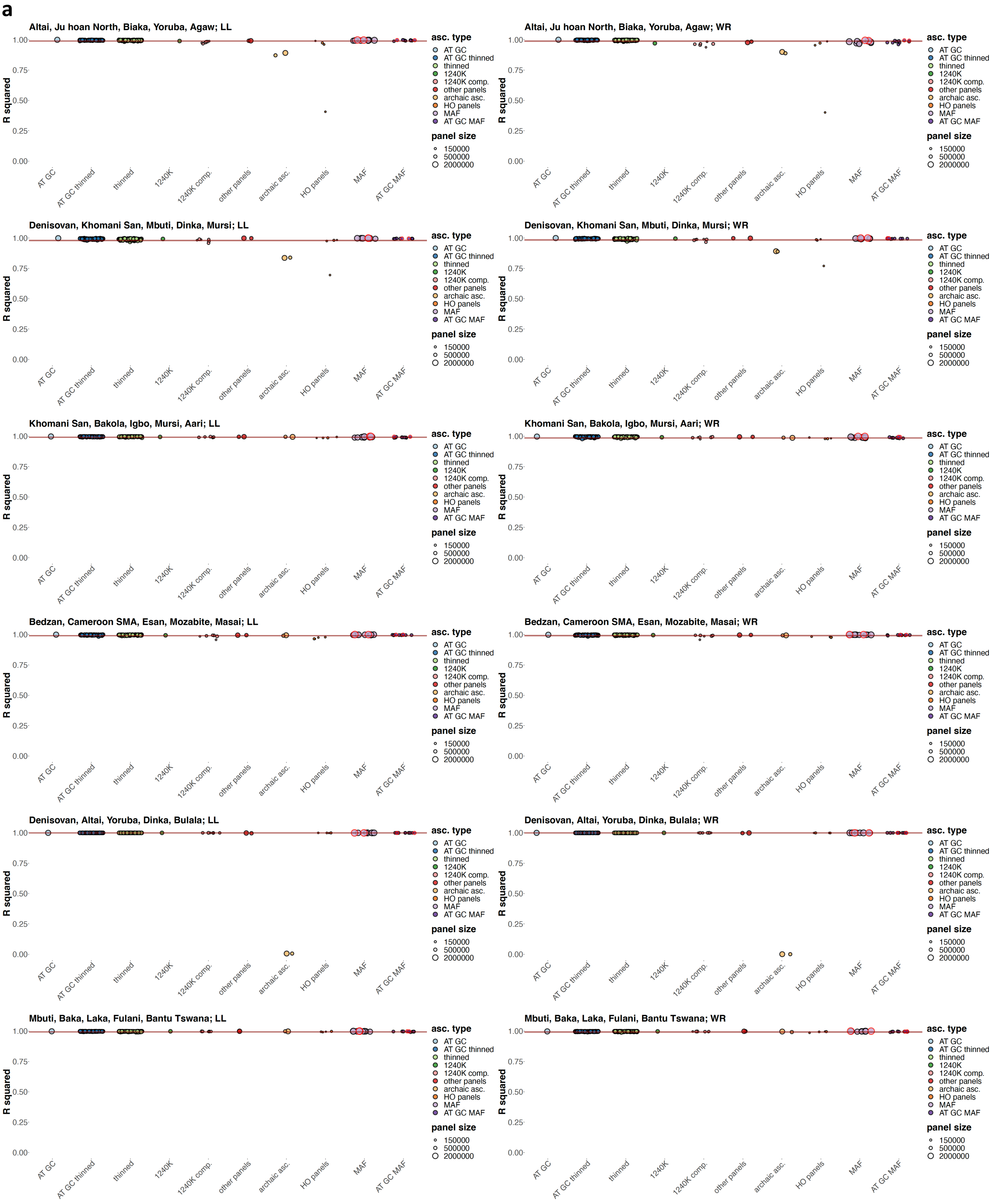
a
Altai, Ju hoan North, Biaka, Yoruba, Agaw; LL
Altai, Ju hoan North, Biaka, Yoruba, Agaw; WR
Denisovan, Khomani San, Mbuti, Dinka, Mursi; LL
Denisovan, Khomani San, Mbuti, Dinka, Mursi; WR
Khomani San, Bakola, Igbo, Mursi, Aari; LL
Khomani San, Bakola, Igbo, Mursi, Aari; WR
Bedzan, Cameroon SMA, Esan, Mozabite, Masai; LL
Bedzan, Cameroon SMA, Esan, Mozabite, Masai; WR
Denisovan, Altai, Yoruba, Dinka, Bulala; LL
Denisovan, Altai, Yoruba, Dinka, Bulala; WR
Mbuti, Baka, Laka, Fulani, Bantu Tswana; LL
Mbuti, Baka, Laka, Fulani, Bantu Tswana; WR
R squared
1.00
0.75
0.50
0.25
0.00
AT GC
AT GC thinned
thinned
1240K
1240K comp.
other panels
archaic asc.
HO panels
MAF
AT GC MAF
asc. type
AT GC
AT GC thinned
thinned
1240K
1240K comp.
other panels
archaic asc.
HO panels
MAF
AT GC MAF
panel size
150000
500000
2000000

**b**

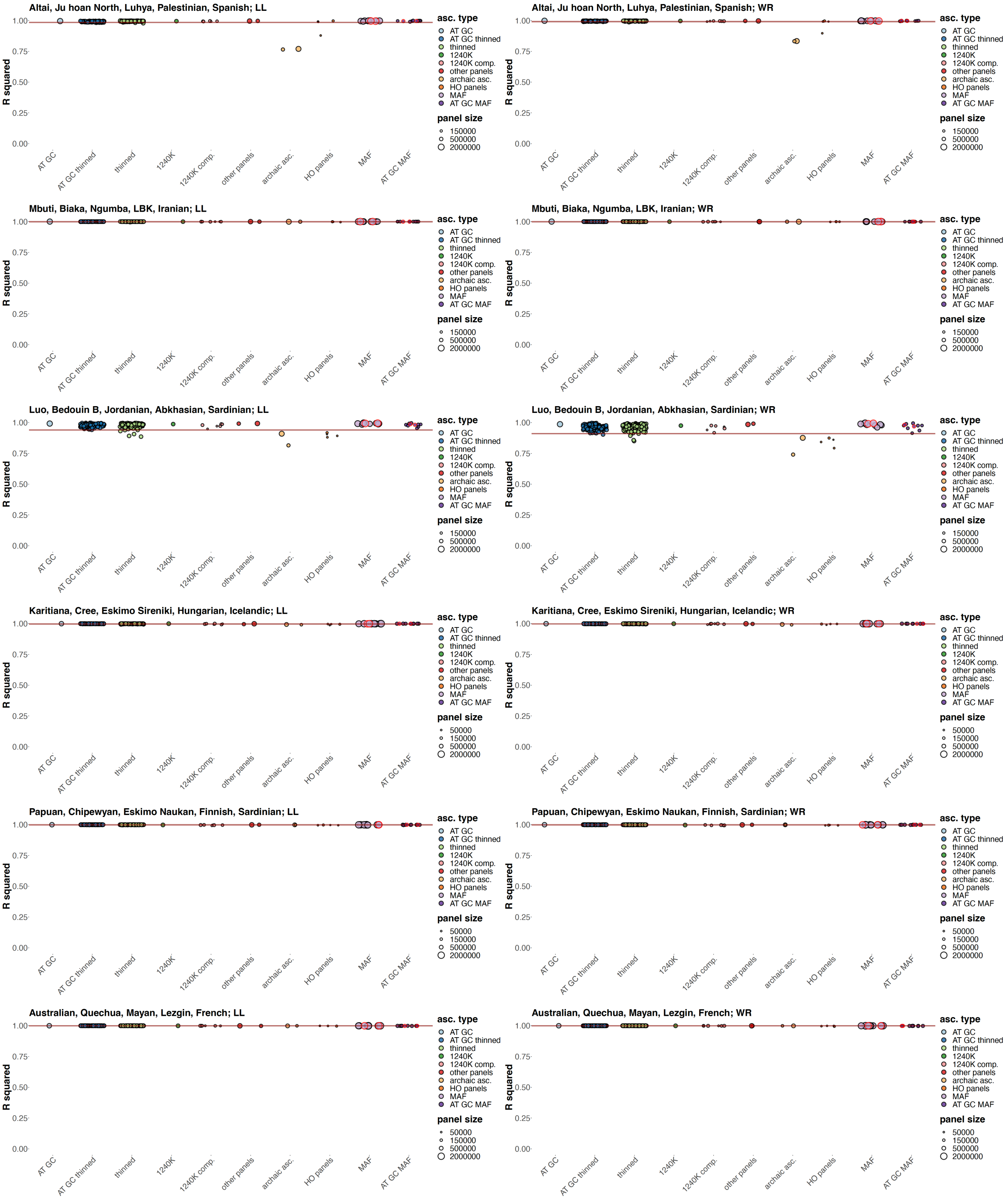
Altai, Ju hoan North, Luhya, Palestinian, Spanish; LL
Altai, Ju hoan North, Luhya, Palestinian, Spanish; WR
Mbuti, Biaka, Ngumba, LBK, Iranian; LL
Mbuti, Biaka, Ngumba, LBK, Iranian; WR
Luo, Bedouin B, Jordanian, Abkhasian, Sardinian; LL
Luo, Bedouin B, Jordanian, Abkhasian, Sardinian; WR
Karitiana, Cree, Eskimo Sireniki, Hungarian, Icelandic; LL
Karitiana, Cree, Eskimo Sireniki, Hungarian, Icelandic; WR
Papuan, Chipewyan, Eskimo Naukan, Finnish, Sardinian; LL
Papuan, Chipewyan, Eskimo Naukan, Finnish, Sardinian; WR
Australian, Quechua, Mayan, Lezgin, French; LL
Australian, Quechua, Mayan, Lezgin, French; WR
R squared
1.00
0.75
0.50
0.25
0.00
AT GC
AT GC thinned
thinned
1240K
1240K comp.
other panels
archaic asc.
HO panels
MAF
AT GC MAF
asc. type
panel size
50000
150000
500000
2000000

**Suppl. Fig. 4.** Variance in fits of an exhaustive collection of 32,745 simple admixture graphs (five groups and two admixture events) resulting from ascertainment or random site subsampling expressed as $R^2$ of linear trends. Correlation of LL or WR was explored for graphs fitted on all sites and on ascertained datasets. Results are shown for twelve population combinations indicated in plot titles (panels **a, b**). Results for ascertainment on variants common in Africans (either those having no detectable West Eurasian ancestry or on all Africans in the SGDP dataset) are circled in red. As a starting point for generating different ascertainments, we used either 11,706,773 sites (with no missing data at the group level) polymorphic in a set of 48 archaic and African groups composed of 97 individuals, or 10,051,585 such sites in 59 archaic, African, European, and Middle Eastern groups composed of 120 individuals, or 5,296,653 such sites in 51 Papuan, Native American, European, Anatolian, and Caucasian groups composed of 112 individuals (Suppl. Table 1). Thirty eight site subsampling schemes were explored: 1) AT/GC mutation classes; 2) random thinning of the AT/GC dataset to the 1240K SNP count for a given combination of groups (no missing data allowed), results for 100 thinned replicates are shown; 3) random thinning of all sites to the 1240K SNP count, results for 100 thinned replicates are shown; 4) the 1240K SNP panel; 5) major components of the 1240K panel: sites included in the Illumina 650Y and/or Human Origins SNP arrays, sites included exclusively in one of them, and remaining sites; 6) the 1000K and 2200K SNP panels; 7) restricting to sites polymorphic in a group composed of the three high-coverage archaic individuals (either all such sites or transversions only); 8) the largest Human Origins sub-panels (4, 5, 13) or their union (4+5); 9) restricting to common variants based on a global MAF threshold of 5% or on the same threshold in one of nine continental-scale groups; 10) the same procedure repeated on AT/GC sites. The size of the resulting SNP panels is coded by point size, and ten broad ascertainment types are coded by color according to the legend. $R^2$ values for LL are plotted in the left-hand panels, and values for WR are plotted in the right-hand panels. The 2.5$^{th}$ LL or WR percentiles of all the thinned replicates combined, including those on all sites and AT/GC sites, are marked by brown lines.

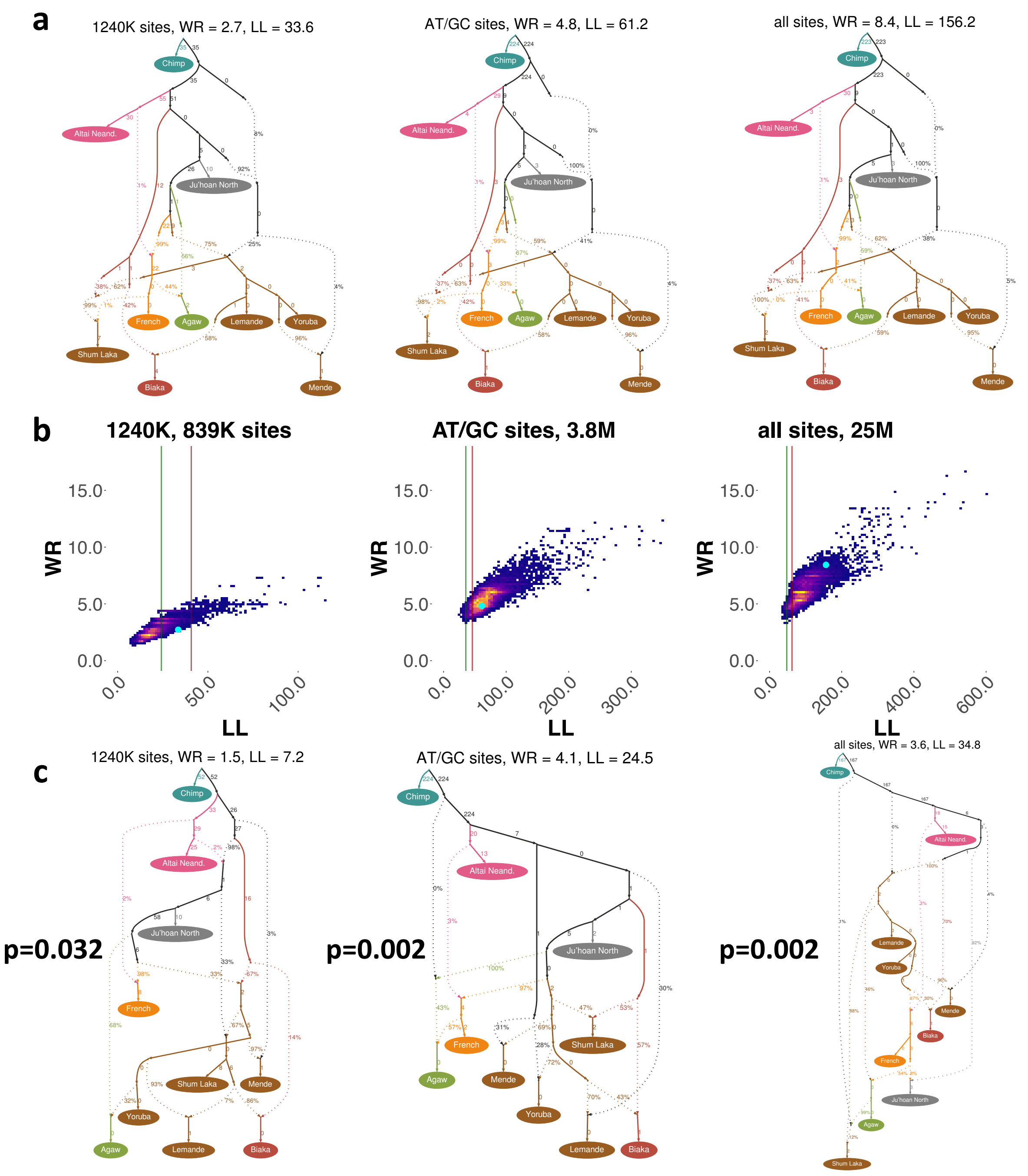

**Suppl. Fig. 5.** Results of a search for optimal admixture graph models on three datasets based on Lipson *et al*. (2020): 1240K, AT/GC mutation types, and all sites. (**a**) The published 10-population model (Lipson et al. 2020) with 8 admixture events fitted on the three datasets, with WR and LL values shown. Distinct populations are colored along with their ancestral lineages, and the cluster of West African populations is colored in brown. (**b**) Density plots illustrating fits of ca. 10,000 distinct topologies per dataset (found with *findGraphs*, Maier et al. 2022 preprint) in the LL vs. WR coordinates. Green vertical lines mark the median LL of the highest-ranking model fitted to bootstrap replicates of the dataset, and red lines mark 95$^{th}$ percentile of that distribution. Position of the published 10-population model in these coordinates is marked by the cyan dot. (**c**) Highest-ranking models (according to LL) with 8 admixture events found with *findGraphs* on each dataset. Populations and ancestral lineages are color-coded in the same way as in panel **a**. WR and LL values are shown above the plots. We also compared the fit (i.e., LL) of the highest-ranking newly found model with that of the published model on each dataset relying on a bootstrap resampling approach: comparison of two LL distributions on 500 resampled sets of SNP blocks (Maier et al. 2022 preprint). For all three SNP sets, the difference in LL between the highest-ranking model found by the automated search and the published model was statistically significant, with empirical two-tailed p-values ranging from $< 0.002$ to 0.032. In other words, it was shown for the three datasets explored that the published model fits significantly worse than the newly found highest-ranking models.

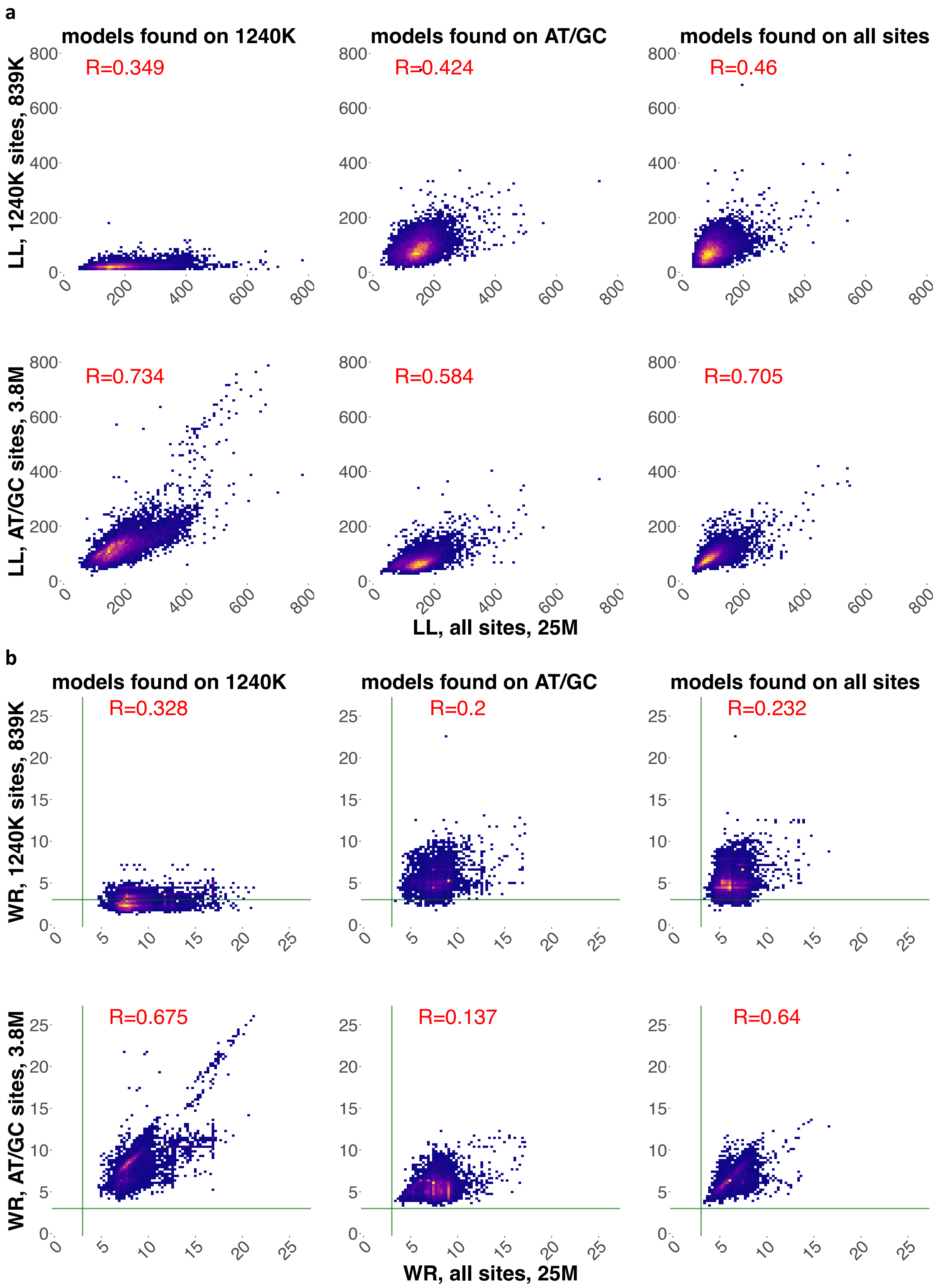
a
models found on 1240K
models found on AT/GC
models found on all sites
R=0.349
R=0.424
R=0.46
R=0.734
R=0.584
R=0.705
LL, 1240K sites, 839K
LL, AT/GC sites, 3.8M
LL, all sites, 25M
b
models found on 1240K
models found on AT/GC
models found on all sites
R=0.328
R=0.2
R=0.232
R=0.675
R=0.137
R=0.64
WR, 1240K sites, 839K
WR, AT/GC sites, 3.8M
WR, all sites, 25M

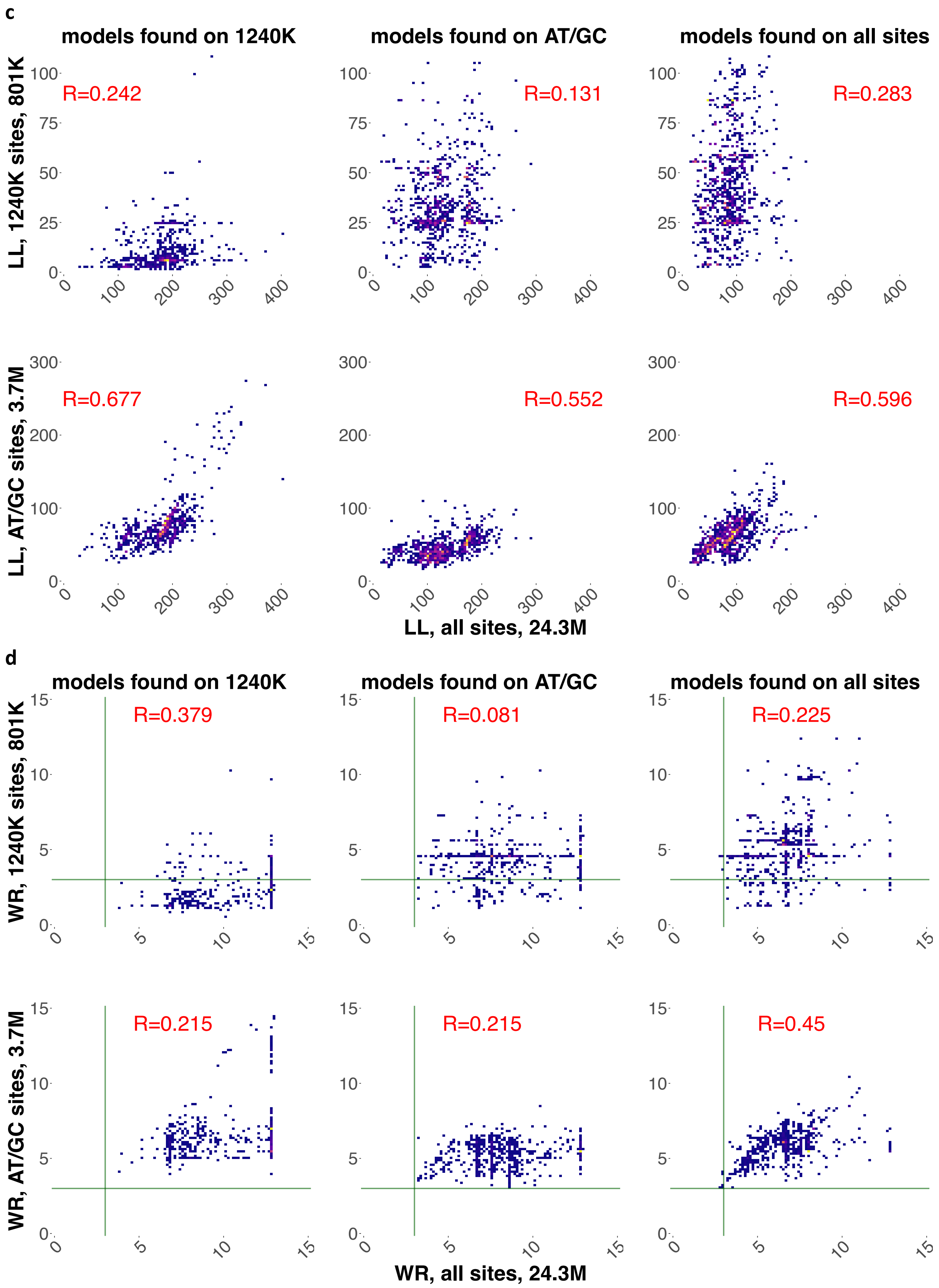

**Suppl. Fig. 6.** Density scatterplots illustrating the effects of the 1240K and AT/GC ascertainments on LL (**a, c**) or WR (**b, d**) of admixture graphs found using *findGraphs* on the set of groups from Lipson *et al*. (2020). (**a, b**) Between 9,927 and 9,990 unique topologies including 10 groups and 8 admixture events were found in 10,000 independent iterations of the *findGraphs* algorithm on three datasets (shown in three columns). (**c, d**) Between 779 and 971 unique topologies including 7 groups and 4 admixture events were found in 2,000 independent iterations of the *findGraphs* algorithm on the three datasets. On x-axes LL or WR for all sites are shown. LL or WR values for the 1240K ascertainment are shown in the upper row on the y-axes, and LL or WR for AT/GC sites are shown in the lower row. Pearson correlation coefficients are displayed beside each plot in red. The commonly used WR threshold for fitting models (3 SE) is marked with green vertical and horizontal lines.

**a**

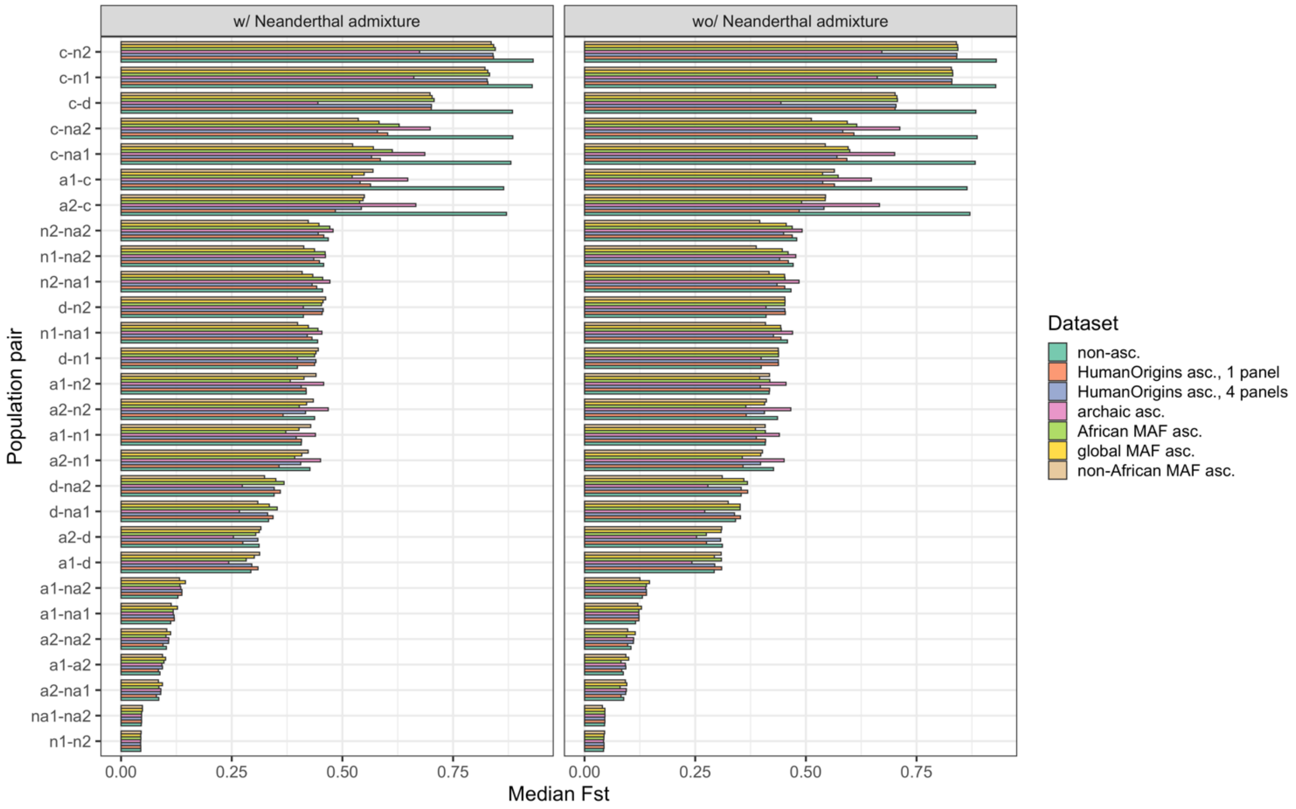
w/ Neanderthal admixture
wo/ Neanderthal admixture
Population pair
c-n2
c-n1
c-d
c-na2
c-na1
a1-c
a2-c
n2-na2
n1-na2
n2-na1
d-n2
n1-na1
d-n1
a1-n2
a2-n2
a1-n1
a2-n1
d-na2
d-na1
a2-d
a1-d
a1-na2
a1-na1
a2-na2
a1-a2
a2-na1
na1-na2
n1-n2
0.00
0.25
0.50
0.75
Median Fst
Dataset
non-asc.
HumanOrigins asc., 1 panel
HumanOrigins asc., 4 panels
archaic asc.
African MAF asc.
global MAF asc.
non-African MAF asc.

**b**

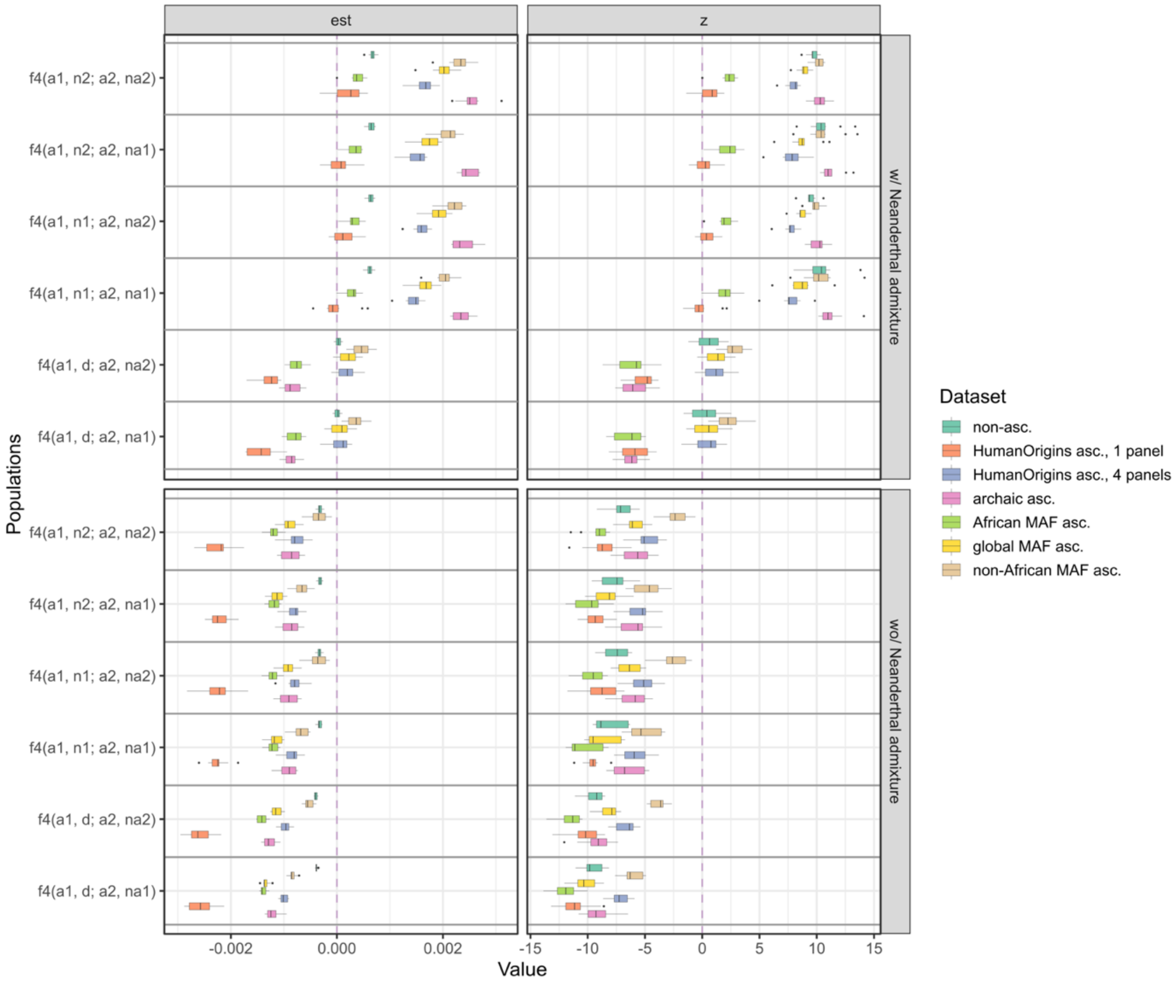
est
z
f4(a1, n2; a2, na2)
f4(a1, n2; a2, na1)
f4(a1, n1; a2, na2)
f4(a1, n1; a2, na1)
f4(a1, d; a2, na2)
f4(a1, d; a2, na1)
Populations
w/ Neanderthal admixture
wo/ Neanderthal admixture
-0.002
0.000
0.002
-15
-10
-5
0
5
10
15
Value
Dataset
non-asc.
HumanOrigins asc., 1 panel
HumanOrigins asc., 4 panels
archaic asc.
African MAF asc.
global MAF asc.
non-African MAF asc.

**c**

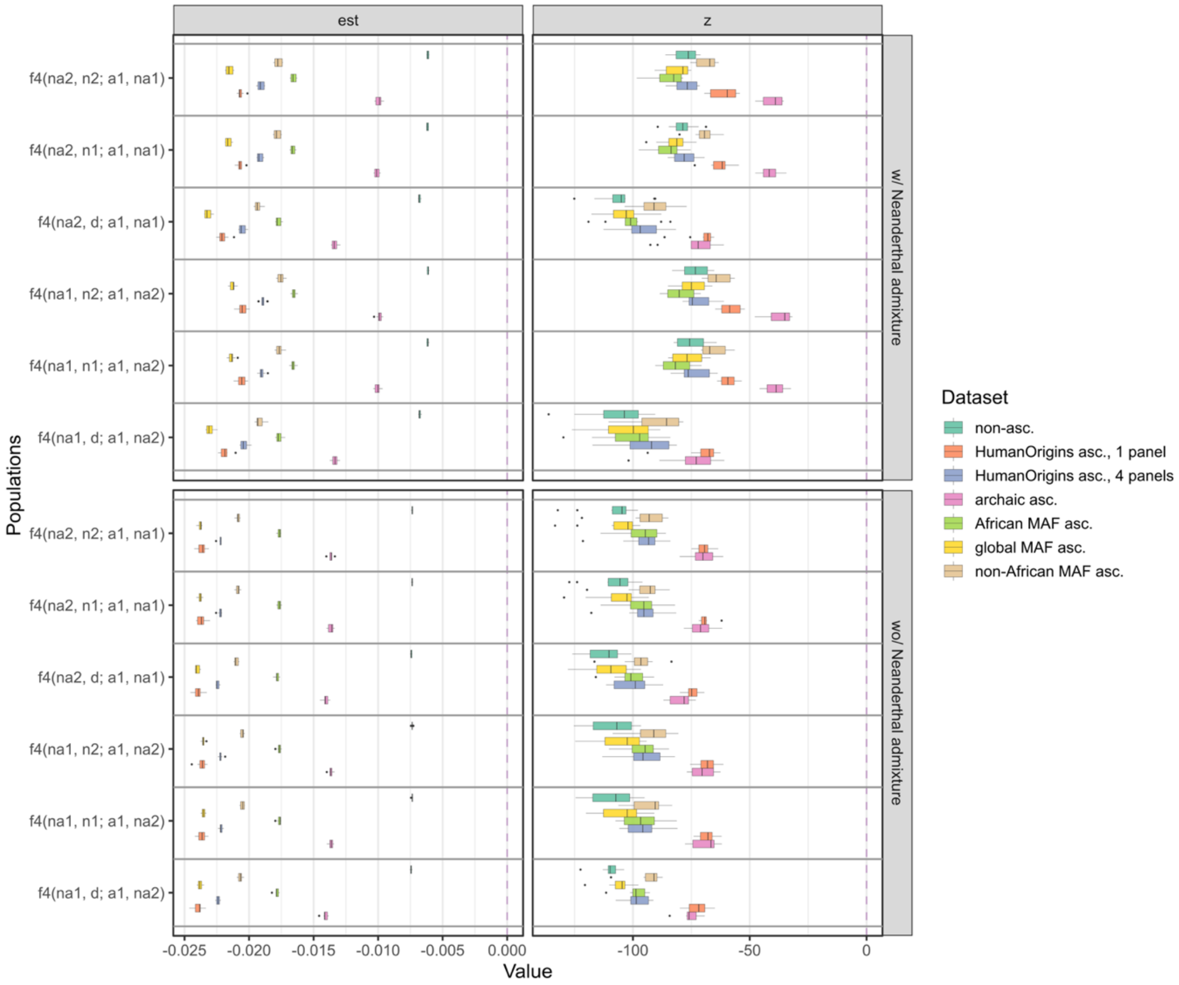

est
z
f4(na2, n2; a1, na1)
f4(na2, n1; a1, na1)
f4(na2, d; a1, na1)
f4(na1, n2; a1, na2)
f4(na1, n1; a1, na2)
f4(na1, d; a1, na2)
w/ Neanderthal admixture
wo/ Neanderthal admixture
Populations
-0.025
-0.020
-0.015
-0.010
-0.005
0.000
-100
-50
0
Value
Dataset
non-asc.
HumanOrigins asc., 1 panel
HumanOrigins asc., 4 panels
archaic asc.
African MAF asc.
global MAF asc.
non-African MAF asc.

**d**

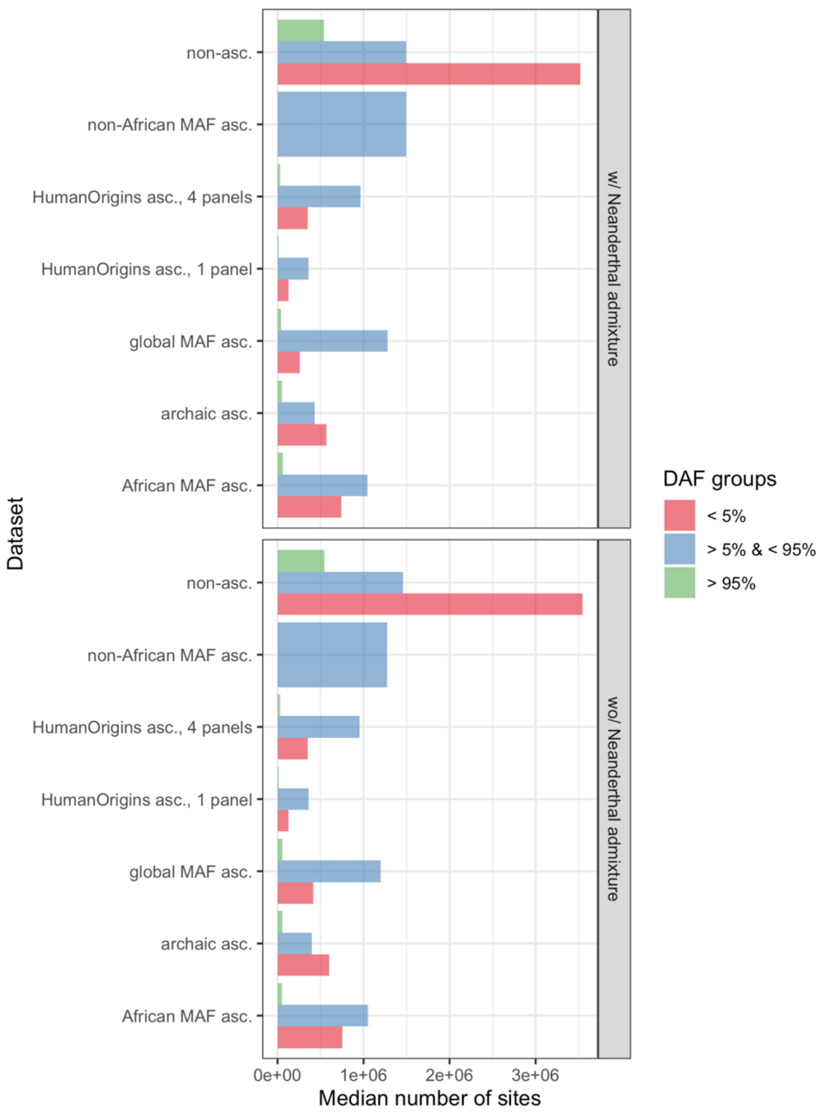

non-asc.
non-African MAF asc.
HumanOrigins asc., 4 panels
HumanOrigins asc., 1 panel
global MAF asc.
archaic asc.
African MAF asc.
w/ Neanderthal admixture
non-asc.
non-African MAF asc.
HumanOrigins asc., 4 panels
HumanOrigins asc., 1 panel
global MAF asc.
archaic asc.
African MAF asc.
wo/ Neanderthal admixture
Dataset
0e+00
1e+06
2e+06
3e+06
Median number of sites
DAF groups
< 5%
> 5% & < 95%
> 95%

e

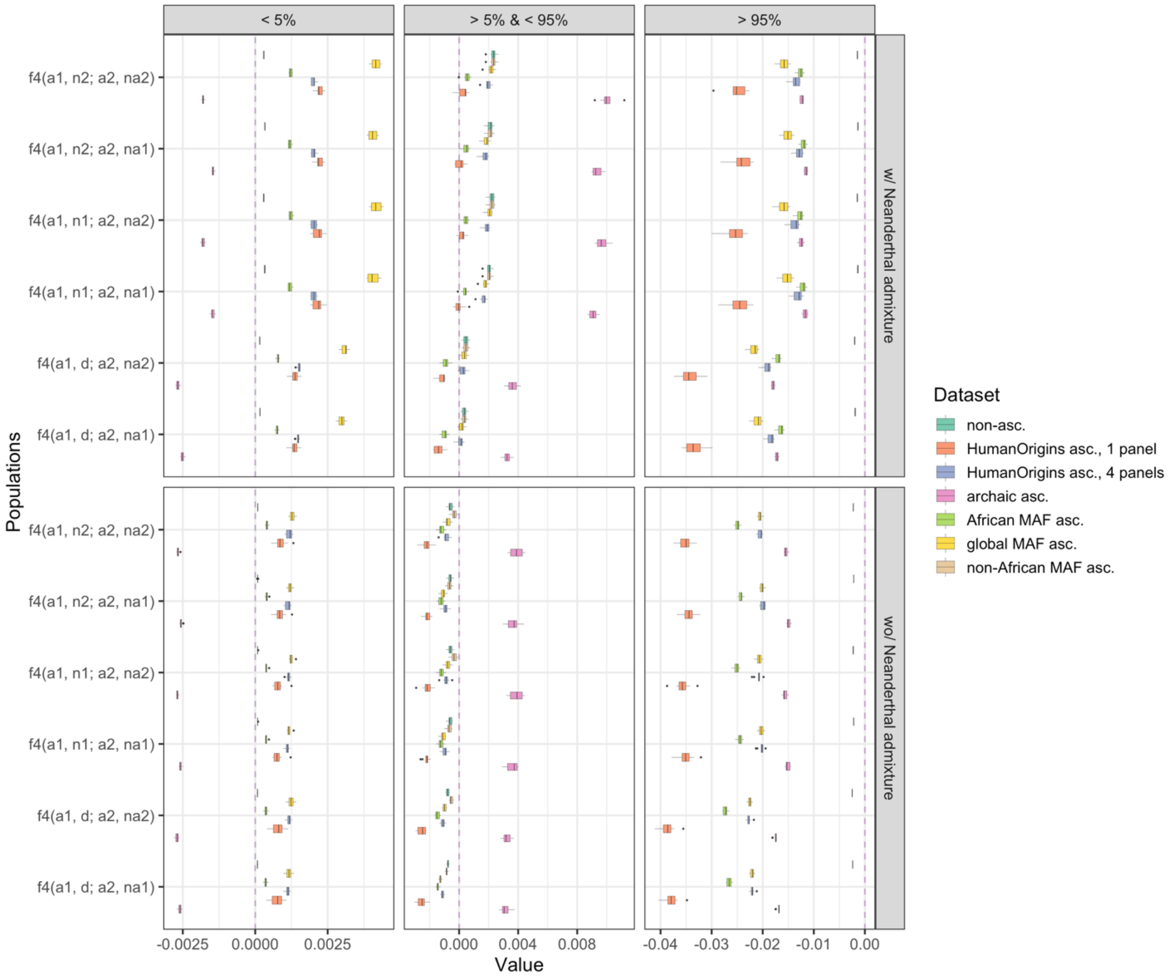

< 5%
> 5% & < 95%
> 95%
w/ Neanderthal admixture
wo/ Neanderthal admixture
f4(a1, n2; a2, na2)
f4(a1, n2; a2, na1)
f4(a1, n1; a2, na2)
f4(a1, n1; a2, na1)
f4(a1, d; a2, na2)
f4(a1, d; a2, na1)
Populations
-0.0025
0.0000
0.0025
0.000
0.004
0.008
-0.04
-0.03
-0.02
-0.01
0.00
Value
Dataset
non-asc.
HumanOrigins asc., 1 panel
HumanOrigins asc., 4 panels
archaic asc.
African MAF asc.
global MAF asc.
non-African MAF asc.

**f**

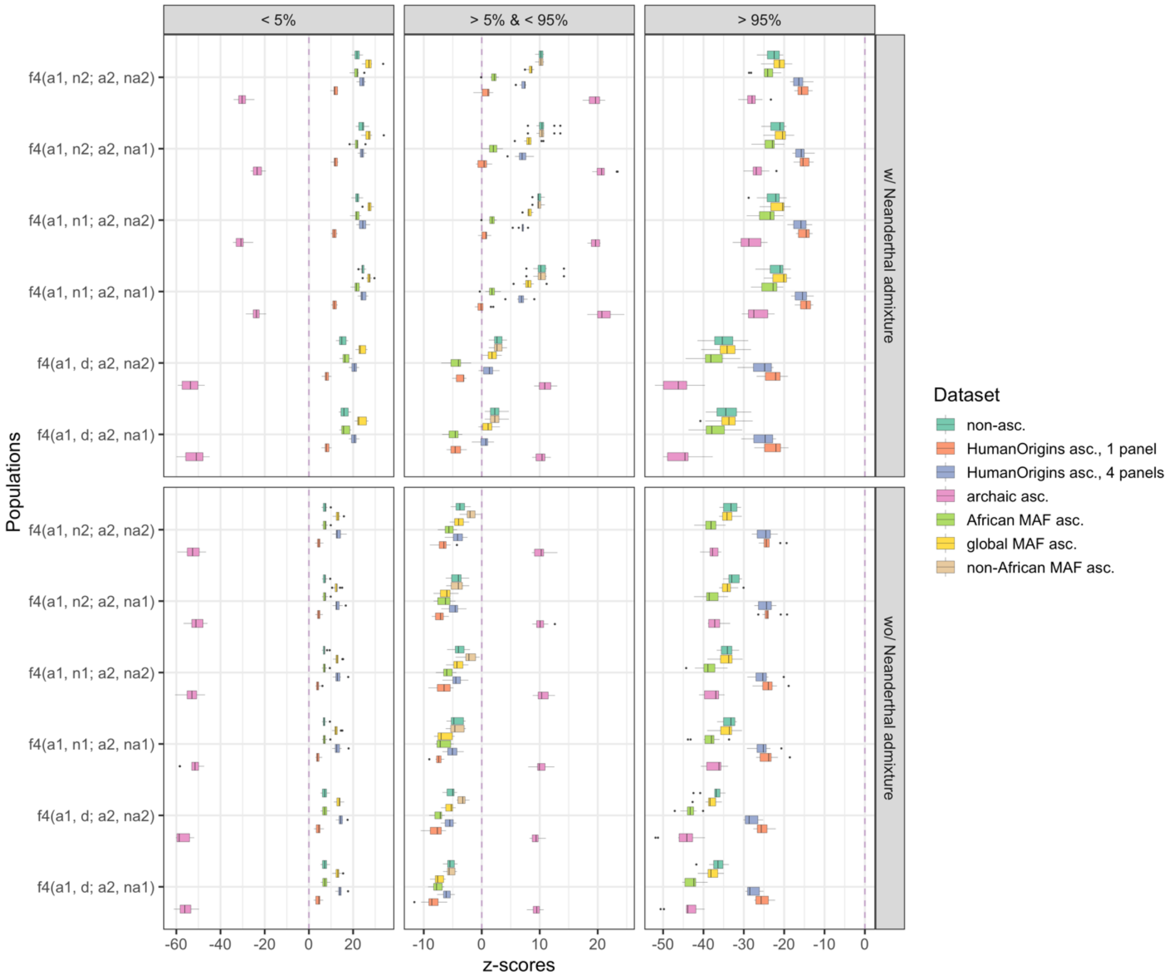

**Suppl. Fig. 7.** (**a**) Influence of ascertainment on $F_{ST}$ across all population pairs on data simulated with or without the "Neanderthal" admixture in "non-Africans". Median $F_{ST}$ values are shown across 10 simulation iterations. Results for seven types of SNP sets are presented: 1) unascertained sites (on average 5.55M polymorphic sites without missing data); 2) Human Origins-like ascertainment, one panel based on the "a2" group (500K sites on average across simulation iterations); 3) Human Origins-like ascertainment, four panes based on randomly selected individuals from four groups ("a1", "a2", "na1", and "na2", 1.34M sites on average); 4) archaic ascertainment (1.05M sites on average); 5) African MAF ascertainment, that is restricting to sites with MAF > 5% in the union of "a1" and "a2" groups (1.66M sites on average); 6) similar MAF ascertainment on the union of "a1", "a2", "na1", "na2" (2.75M sites on average); 7) similar MAF ascertainment on the union of "na1" and "na2" groups (1.04M sites on average). (**b**) Boxplots summarizing various $f_4$-statistics of the form $f_4$("African 1", "archaic"; "African 2", "non-African") for the two simulated topologies, on unascertained and ascertained data across 10 simulation runs. $f_4$-statistics are shown on the left (labelled "est") and their Z-scores are shown on the right. (**c**) Boxplots summarizing various $f_4$-statistics of the form $f_4$("non-African 1 or 2", "archaic"; "African 1", "non-African 2 or 1") for the two simulated topologies, on unascertained and ascertained data across 10 simulation runs. $f_4$-statistics are shown on the left (labelled "est") and their Z-scores are shown on the right. (**d**) Derived allele frequency (DAF) spectra for the unascertained and ascertained datasets. DAF was defined on the union of groups "na1" and "na2", and three allele frequency bins were defined: < 5%, >= 5 & <= 95, > 95%. Median site counts across 10 simulation iterations are presented. Similar results on real data are shown in Suppl. Table 10. (**e** and **f**) Boxplots summarizing DAF-stratified $f_4$-statistics of the form

$f_4$("African 1", "archaic"; "African 2", "non-African 1 or 2") (**e**) and their Z-scores (**f**) for the two simulated topologies, on unascertained and ascertained data.

**a**

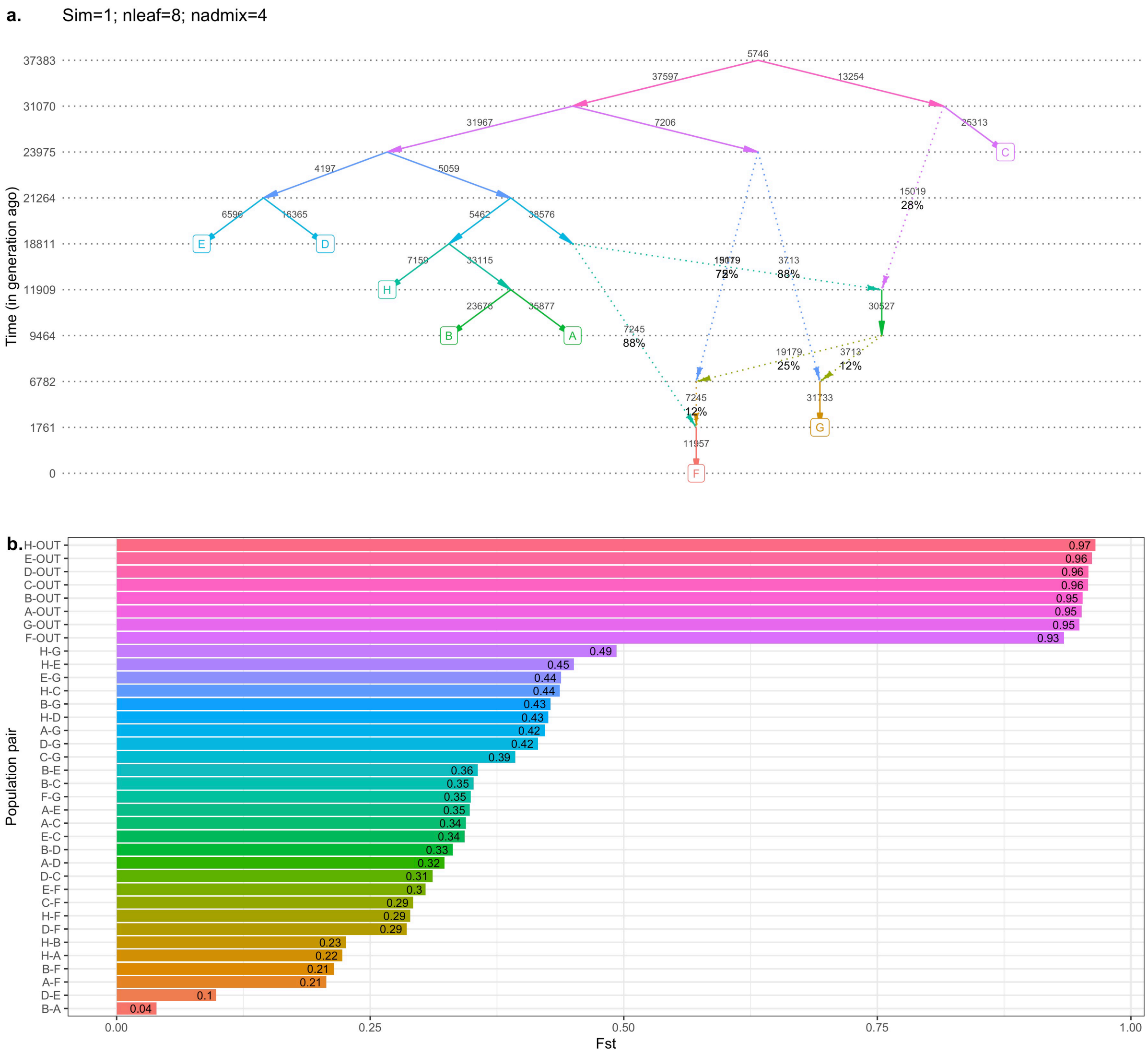
a.
Sim=1; nleaf=8; nadmix=4
Time (in generation ago)
37383
31070
23975
21264
18811
11909
9464
6782
1761
0
A
B
C
D
E
F
G
H
b.
Population pair
Fst
0.00
0.25
0.50
0.75
1.00
H-OUT 0.97
E-OUT 0.96
D-OUT 0.96
C-OUT 0.96
B-OUT 0.95
A-OUT 0.95
G-OUT 0.95
F-OUT 0.93
H-G 0.49
H-E 0.45
E-G 0.44
H-C 0.44
B-G 0.43
H-D 0.43
A-G 0.42
D-G 0.42
C-G 0.39
B-E 0.36
B-C 0.35
F-G 0.35
A-E 0.35
A-C 0.34
E-C 0.34
B-D 0.33
A-D 0.32
D-C 0.31
E-F 0.3
C-F 0.29
H-F 0.29
D-F 0.29
H-B 0.23
H-A 0.22
B-F 0.21
A-F 0.21
D-E 0.1
B-A 0.04

**b**

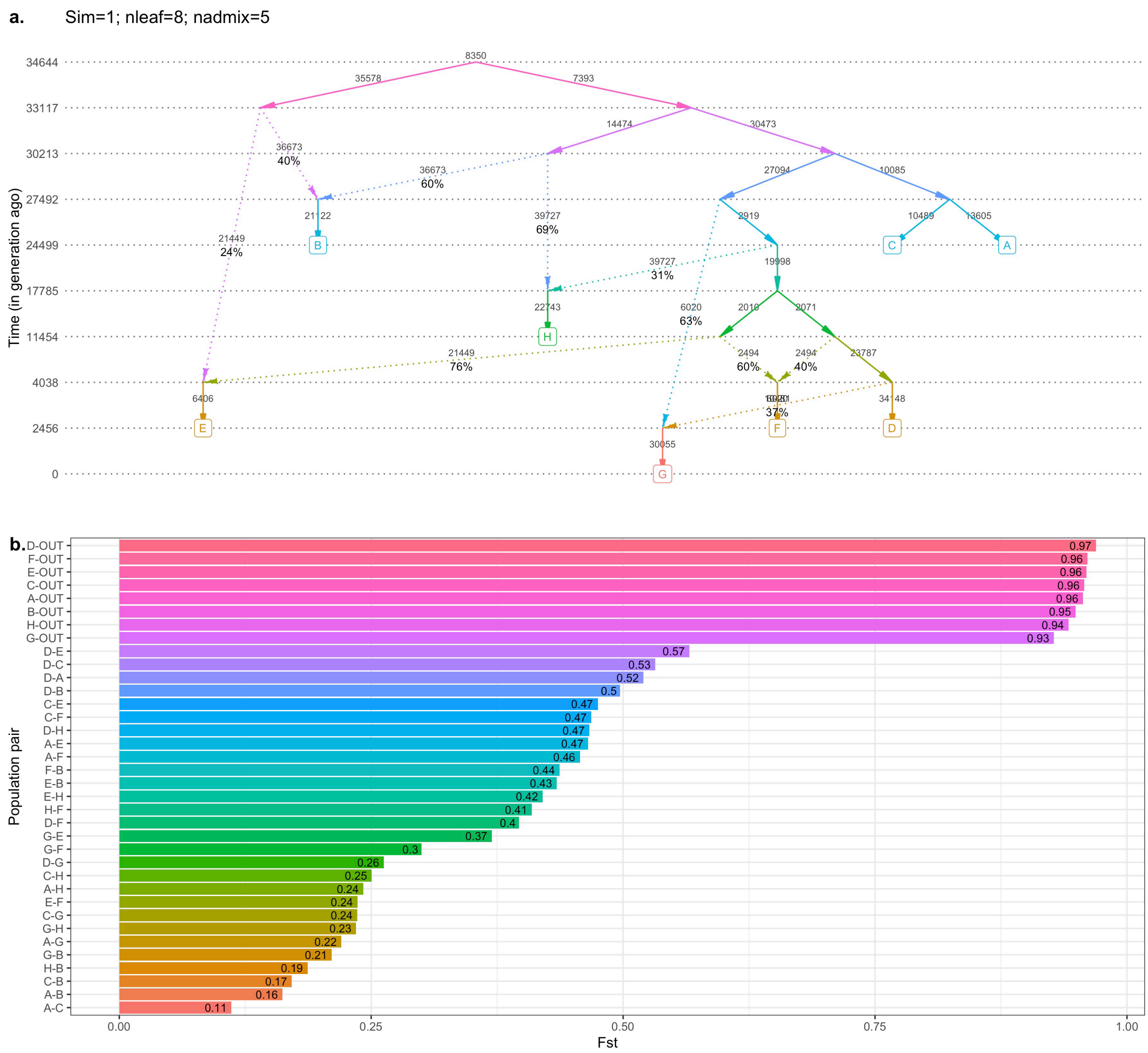
a.
Sim=1; nleaf=8; nadmix=5
Time (in generation ago)
34644
33117
30213
27492
24499
17785
11454
4038
2456
0
A
B
C
D
E
F
G
H
b.
Population pair
Fst
D-OUT 0.97
F-OUT 0.96
E-OUT 0.96
C-OUT 0.96
A-OUT 0.96
B-OUT 0.95
H-OUT 0.94
G-OUT 0.93
D-E 0.57
D-C 0.53
D-A 0.52
D-B 0.5
C-E 0.47
C-F 0.47
D-H 0.47
A-E 0.47
A-F 0.46
F-B 0.44
E-B 0.43
E-H 0.42
H-F 0.41
D-F 0.4
G-E 0.37
G-F 0.3
D-G 0.26
C-H 0.25
A-H 0.24
E-F 0.24
C-G 0.24
G-H 0.23
A-G 0.22
G-B 0.21
H-B 0.19
C-B 0.17
A-B 0.16
A-C 0.11
0.00
0.25
0.50
0.75
1.00

**c**

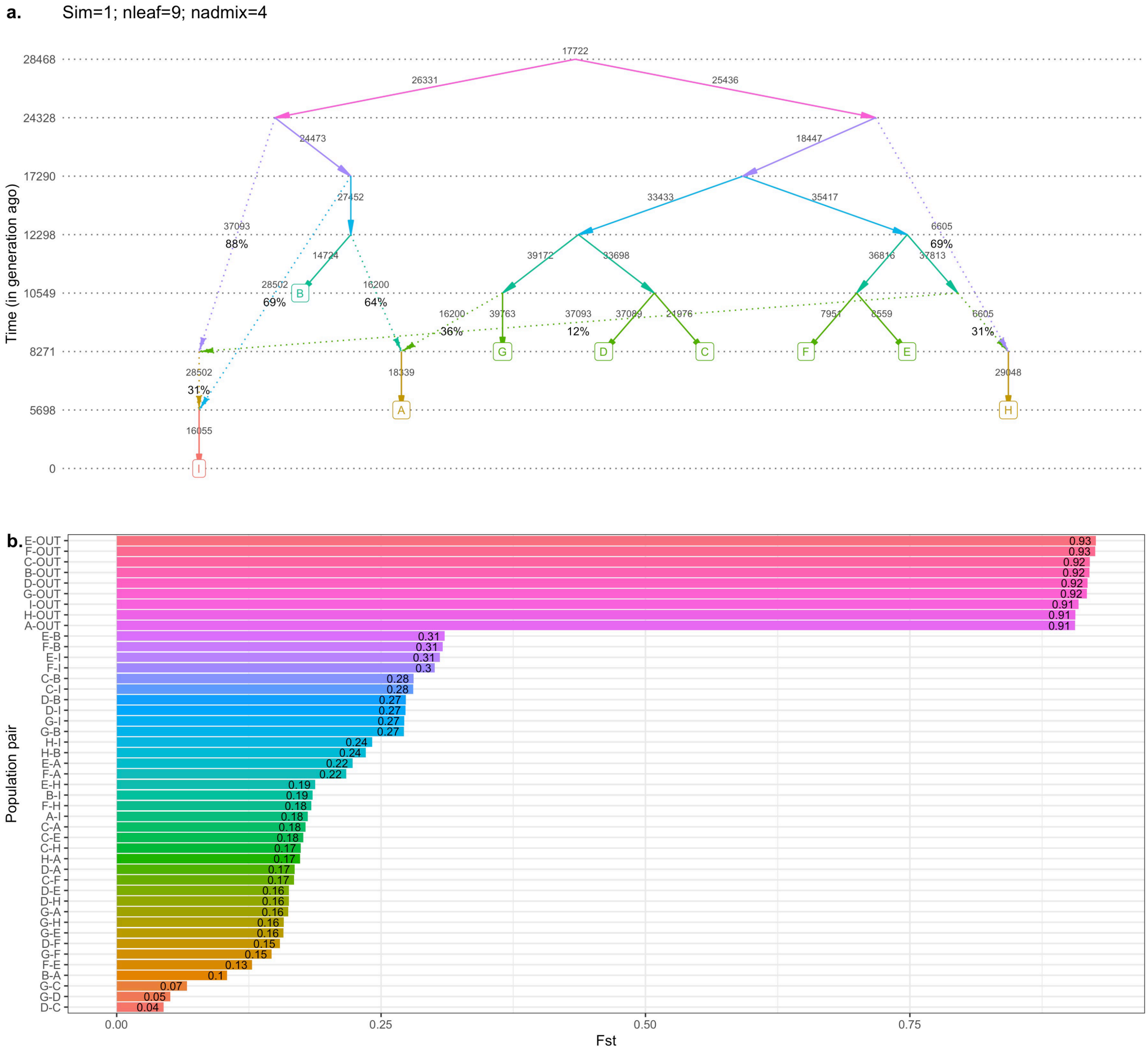
a.
Sim=1; nleaf=9; nadmix=4
Time (in generation ago)
28468
24328
17290
12298
10549
8271
5698
0
b.
Population pair
Fst
E-OUT
F-OUT
C-OUT
B-OUT
D-OUT
G-OUT
I-OUT
H-OUT
A-OUT
0.00
0.25
0.50
0.75

**d**

**a.** Sim=1; nleaf=9; nadmix=5

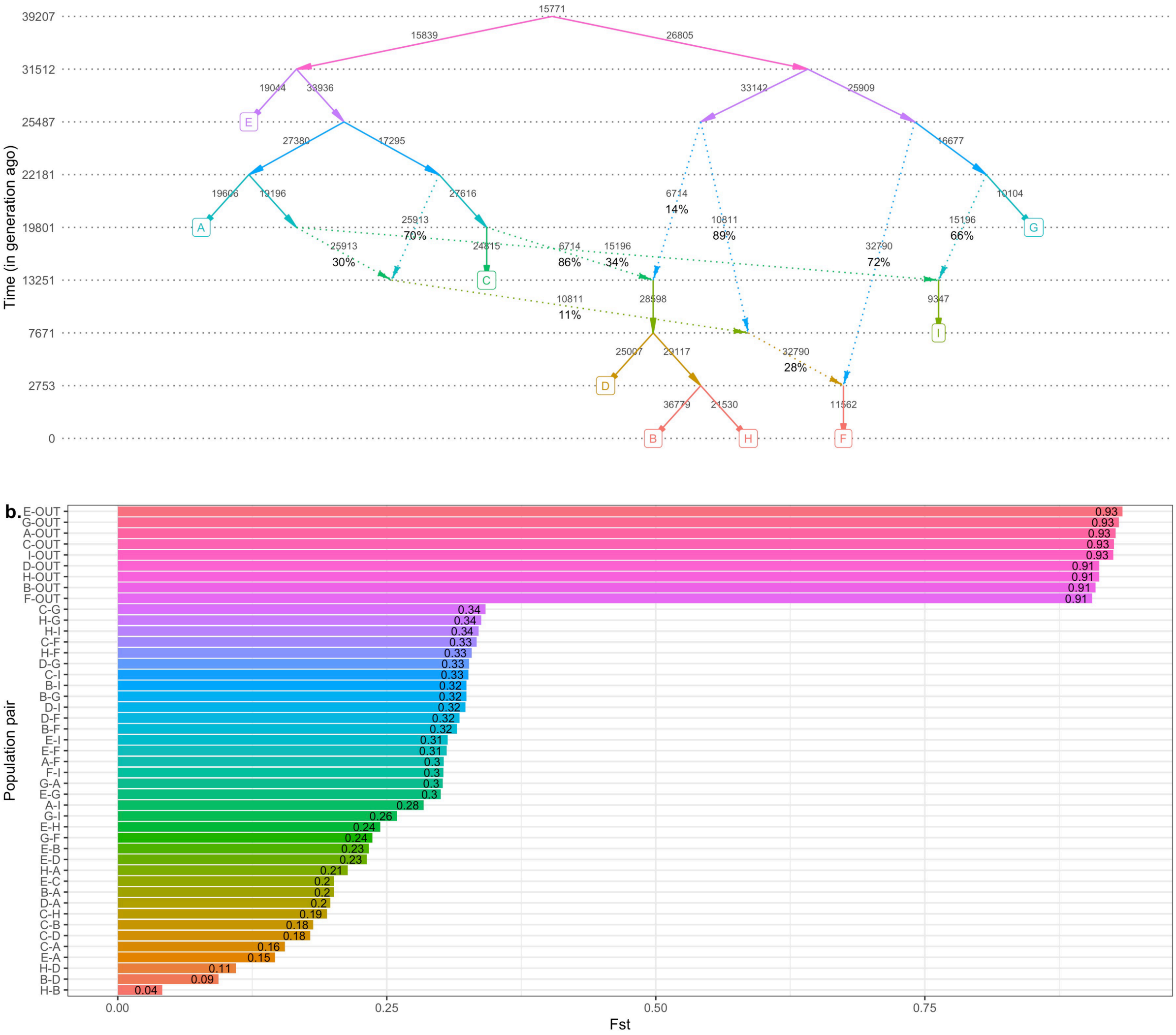

**Suppl. Fig. 8.** Examples of simulated genetic histories in the shape of random admixture graphs including 8 (**a**, **b**) or 9 (**c**, **d**) non-outgroup populations and 4 (**a**, **c**) or 5 (**b**, **d**) admixture events. One topology per graph complexity class is shown as an example, along with $F_{ST}$ values for all population pairs. Time in generations is shown on the y-axis (for visual clarity, points on this axis are not spaced proportionately). Effective population sizes (in diploid individuals) are shown beside each edge. Sampled populations are labelled by letters in squares. Outgroups are not visualized; their divergence was placed at 40,000 generations ago for all the simulated histories, and their effective population size was 100,000 or alternatively 1,000 diploid individuals (both series of simulations were prepared). The color gradients have no special meaning and are used for visual clarity only.

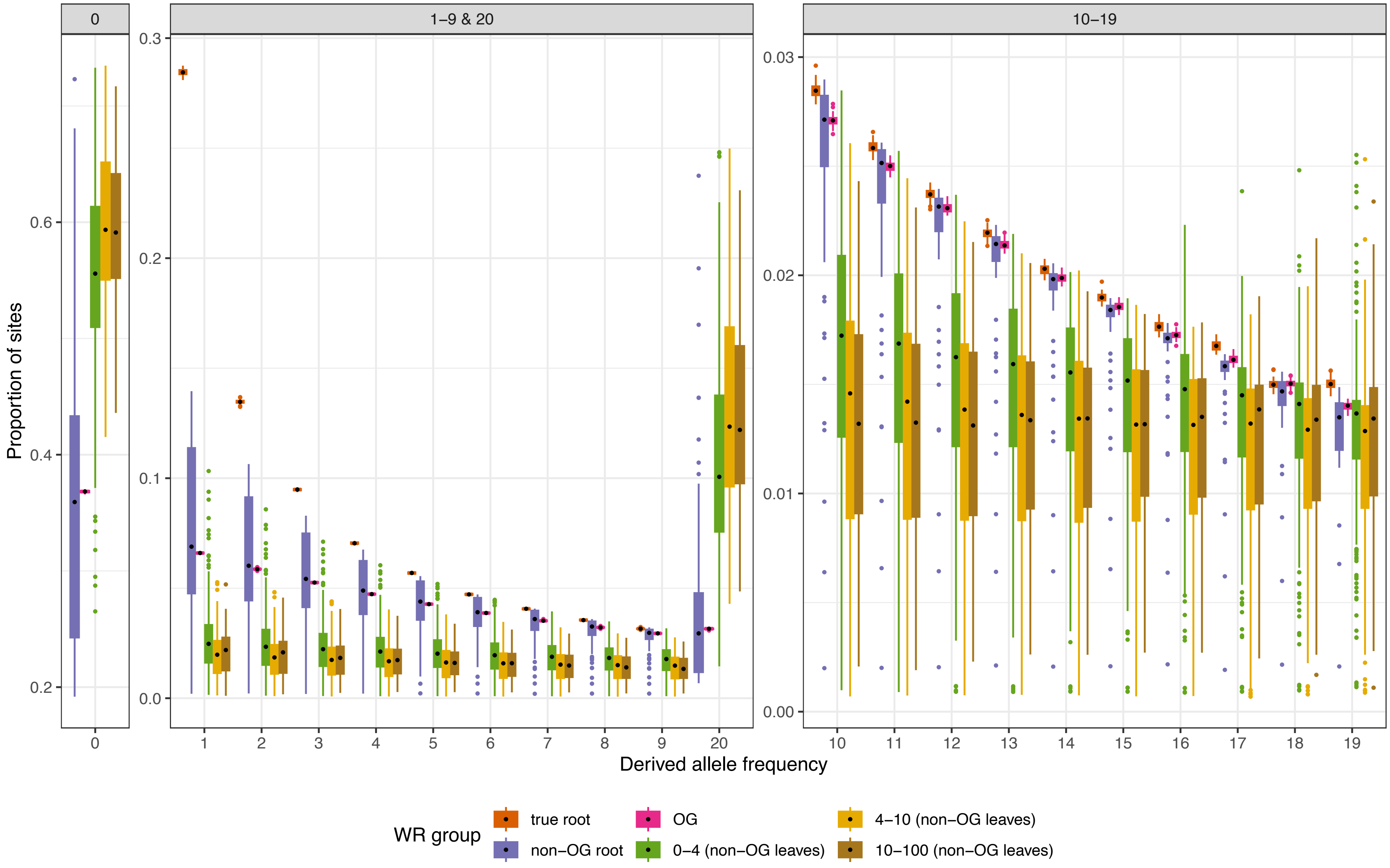

0
1–9 & 20
10–19
Proportion of sites
0.2
0.4
0.6
0.0
0.1
0.2
0.3
0.00
0.01
0.02
0.03
0
1
2
3
4
5
6
7
8
9
20
Derived allele frequency
WR group
true root
non–OG root
OG
0–4 (non–OG leaves)
4–10 (non–OG leaves)
10–100 (non–OG leaves)

**Suppl. Fig. 9.** Derived allele frequency spectra (derived allele count in a sample of 20 chromosomes vs. proportion of sites) across simulated root, outgroup, and non-outgroup populations grouped according to the level of ascertainment bias. We note that outgroups had an effective population size of 100,000 diploid individuals and were included in the fitted admixture graphs; in other words, they were co-analyzed with the other populations. The spectra were calculated for sites polymorphic in the root population sample of 20 chromosomes. Populations sampled at leaves are binned by WR of the true graph fitted to sites heterozygous in a single individual randomly drawn from that population (single-panel Human Origins-like ascertainment). The boxplots summarize DAF across all simulated graphs. DAF bins are shown in three separate panels with different y-axis ranges: 0 derived alleles; 1 to 9 and 20 derived alleles; 10 to 19 derived alleles.

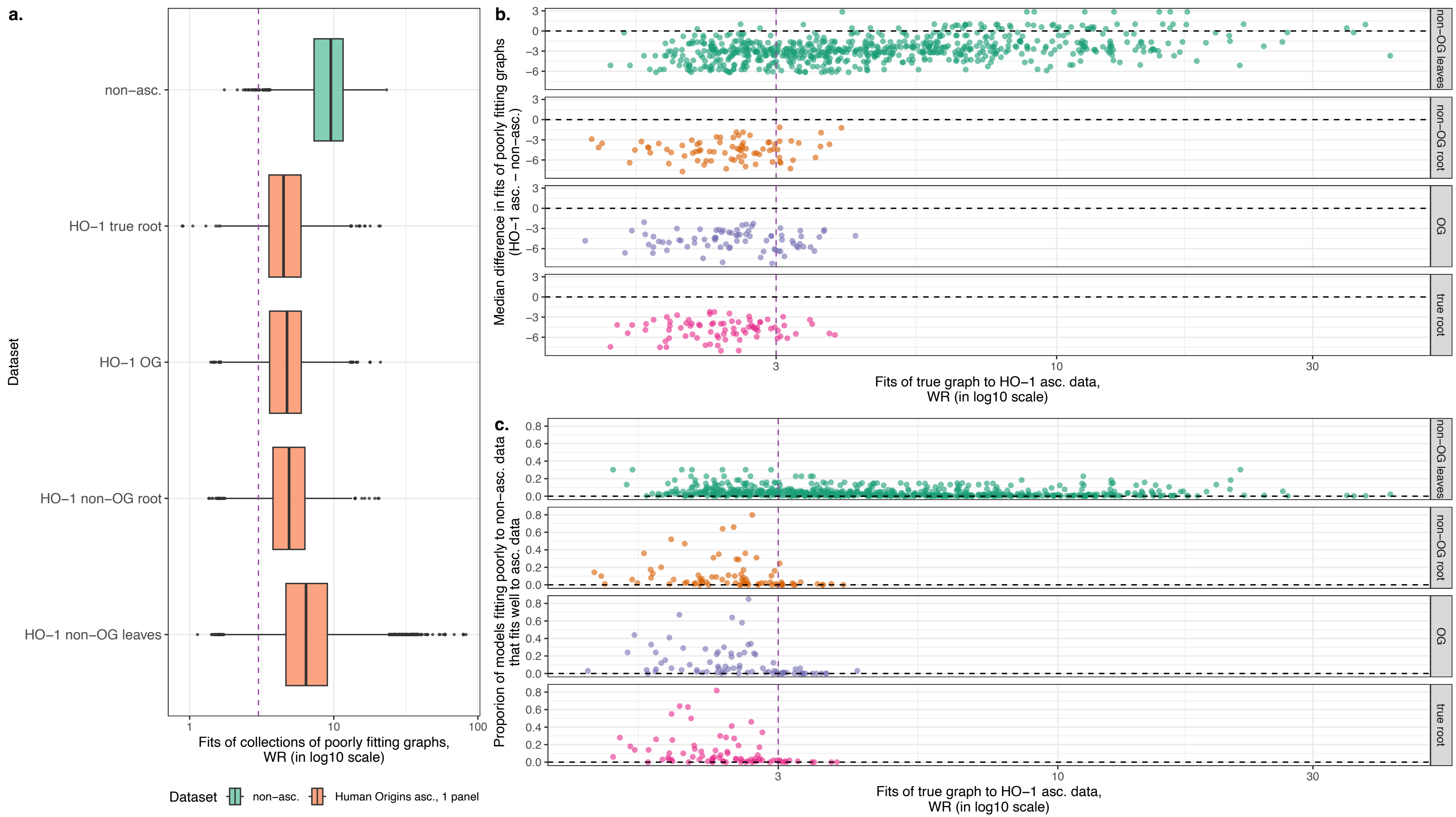

**Suppl. Fig. 10.** Assessing the power of ascertained SNP datasets to reject incorrect admixture graphs. A set of 100 topologically diverse poorly fitting graphs was generated for each simulated topology (see Methods) and fitted to both non-ascertained data and to all SNP sets ascertained using the single-panel Human Origins-like scheme. (**a**) Boxplots summarizing distributions of WRs of incorrect graphs fitted to datasets of several types: non-ascertained,

ascertained on the outgroup with an effective population size of 100,000 that was co-modelled with the other populations (“OG”), on the root population sample (“true root”), on the root of all non-outgroup populations (“non-OG root”), and on non-outgroup populations (“non-OG leaves”). Results for all simulated topologies were pooled. (**b**) Simulated datasets in the “bias” vs. “power” coordinates. WR of the true simulated graph serves as a measure of ascertainment bias (on the *x*-axis), and median difference in WRs of poorly fitting graphs fitted to ascertained and non-ascertained data (on the *y*-axis) serves as a measure of power for each ascertained SNP dataset (Human Origins-like, one panel). Results are shown separately for SNP sets ascertained on the outgroup with an effective population size of 100,000 that was co-modelled with the other populations (“OG”), on the root population sample (“true root”), on the root of all non-outgroup populations (“non-OG root”), and on non-outgroup populations (“non-OG leaves”). Each dot represents an ascertained SNP set. (**c**) This series of plots is similar to that in panel **b**, but another measure of statistical power is used: the proportion of graphs which fit poorly to non-ascertained data (WR > 3 SE) but fit well (WR < 3 SE) to ascertained data.

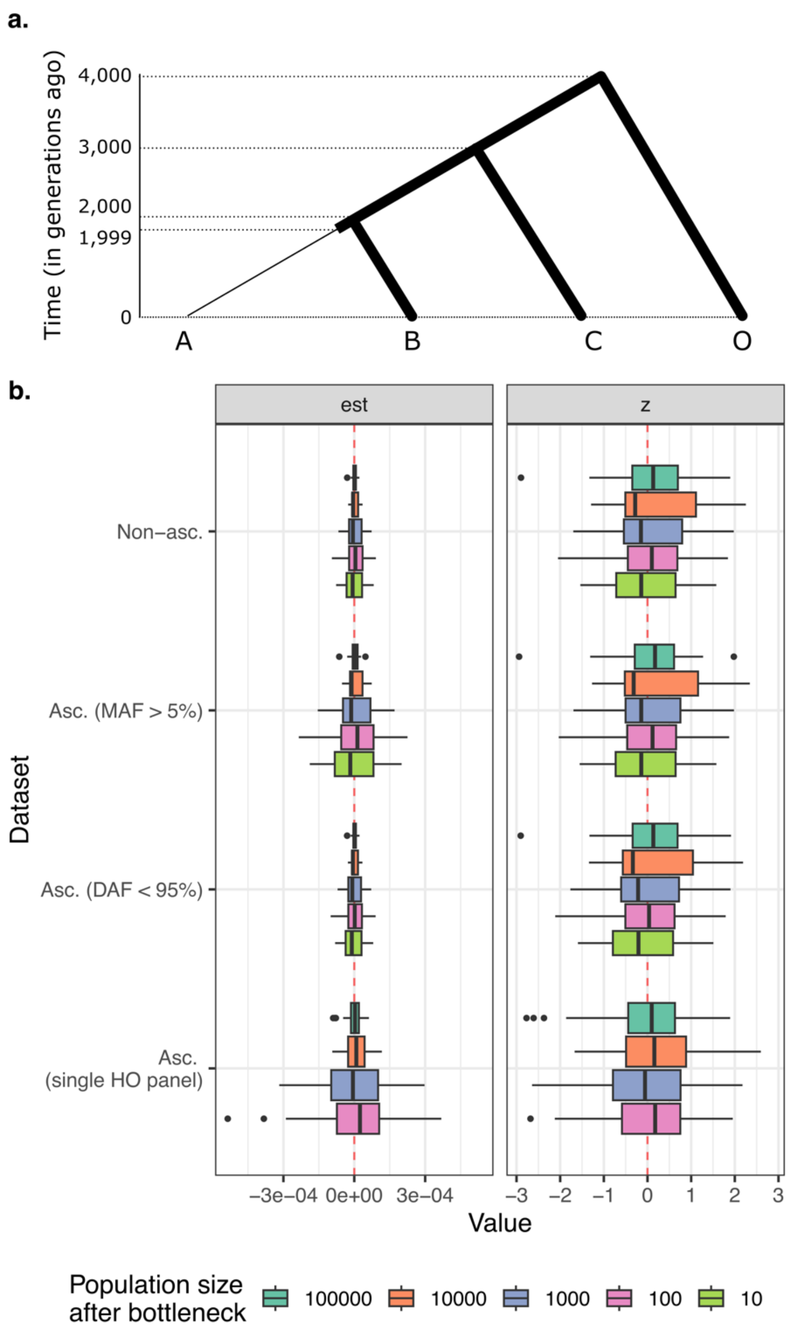

**Suppl. Fig. 11.** The influence of ascertainment on $f_4$-cladality tests in the case of a simple tree (O, (C, (A, B))) shown in (**a**). Effective population size was constant across the tree, except for a drop in group A's size at 1,999 generations in the past: from 100,000 to 10,000, 1,000, 100, or 10 diploid individuals. There was also a set of control simulations without any reduction in effective population size. In panel (**b**), $f_4$-statistics $f_4$(A, B; C, O) and their Z-scores on 20 non-ascertained datasets per bottleneck class and on datasets ascertained in various ways are summarized with boxplots. The ascertainment schemes are as follows: 1) keeping sites with MAF > 5% in the union of groups A and B; 2) keeping sites with *derived* allele frequency < 95% in the union of groups A and B; 3) ascertainment on sites heterozygous in a single randomly selected individual, repeated for all four simulated groups (abbreviated as "Asc. (single HO panel)"). Very similar results were obtained for other MAF (2.5%, 10%) and DAF cutoffs (90%, 97.5%), and they are not shown for brevity. In the case of the Human Origins-like ascertainment, no results are shown for simulations with the smallest effective size of group A since too few SNPs were available due to rapid fixation of variants in the population with a low effective size.

a

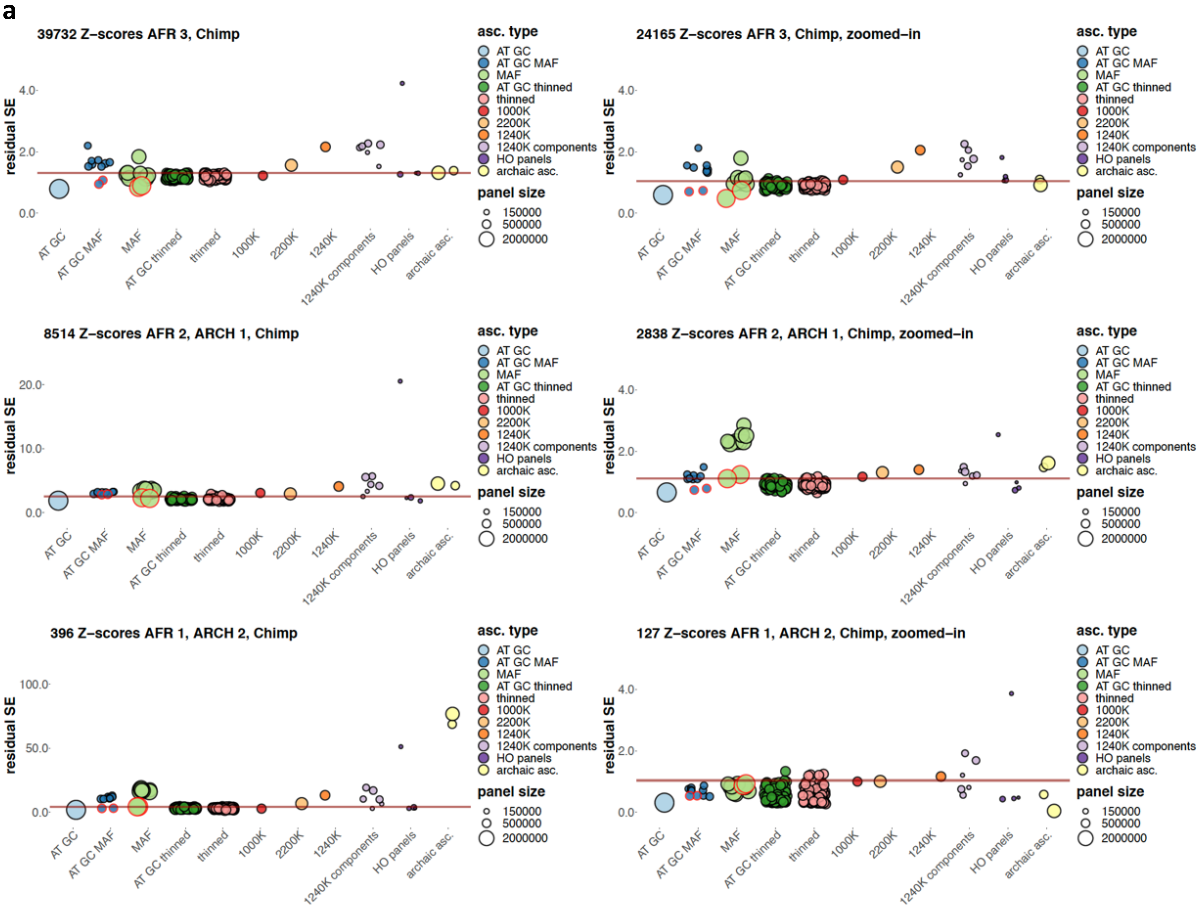
39732 Z–scores AFR 3, Chimp
24165 Z–scores AFR 3, Chimp, zoomed–in
8514 Z–scores AFR 2, ARCH 1, Chimp
2838 Z–scores AFR 2, ARCH 1, Chimp, zoomed–in
396 Z–scores AFR 1, ARCH 2, Chimp
127 Z–scores AFR 1, ARCH 2, Chimp, zoomed–in
residual SE
asc. type
AT GC
AT GC MAF
MAF
AT GC thinned
thinned
1000K
2200K
1240K
1240K components
HO panels
archaic asc.
panel size
150000
500000
2000000

**b**

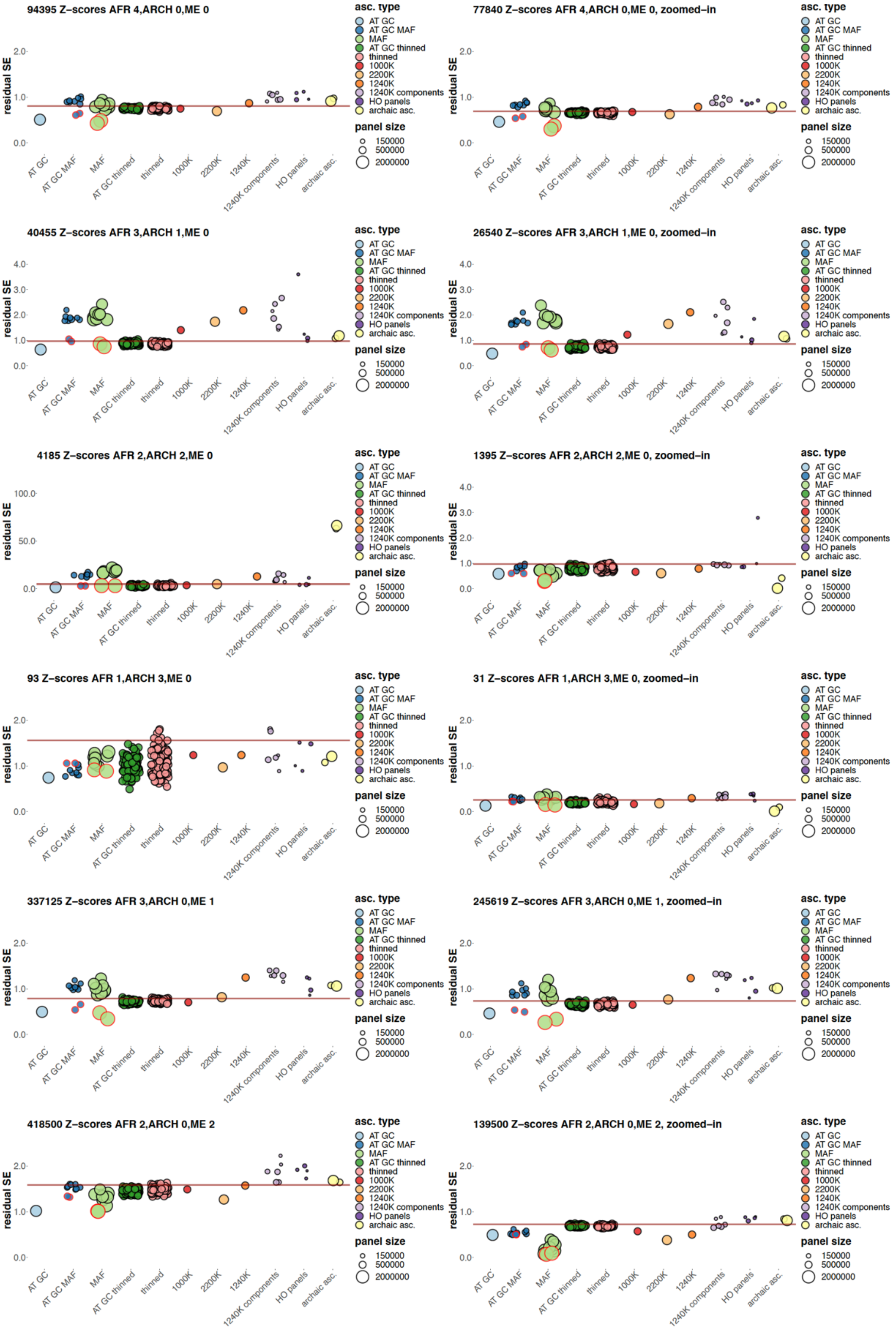
94395 Z–scores AFR 4,ARCH 0,ME 0
77840 Z–scores AFR 4,ARCH 0,ME 0, zoomed–in
40455 Z–scores AFR 3,ARCH 1,ME 0
26540 Z–scores AFR 3,ARCH 1,ME 0, zoomed–in
4185 Z–scores AFR 2,ARCH 2,ME 0
1395 Z–scores AFR 2,ARCH 2,ME 0, zoomed–in
93 Z–scores AFR 1,ARCH 3,ME 0
31 Z–scores AFR 1,ARCH 3,ME 0, zoomed–in
337125 Z–scores AFR 3,ARCH 0,ME 1
245619 Z–scores AFR 3,ARCH 0,ME 1, zoomed–in
418500 Z–scores AFR 2,ARCH 0,ME 2
139500 Z–scores AFR 2,ARCH 0,ME 2, zoomed–in
residual SE
AT GC
AT GC MAF
MAF
AT GC thinned
thinned
1000K
2200K
1240K
1240K components
HO panels
archaic asc.
asc. type
panel size
150000
500000
2000000

**c**

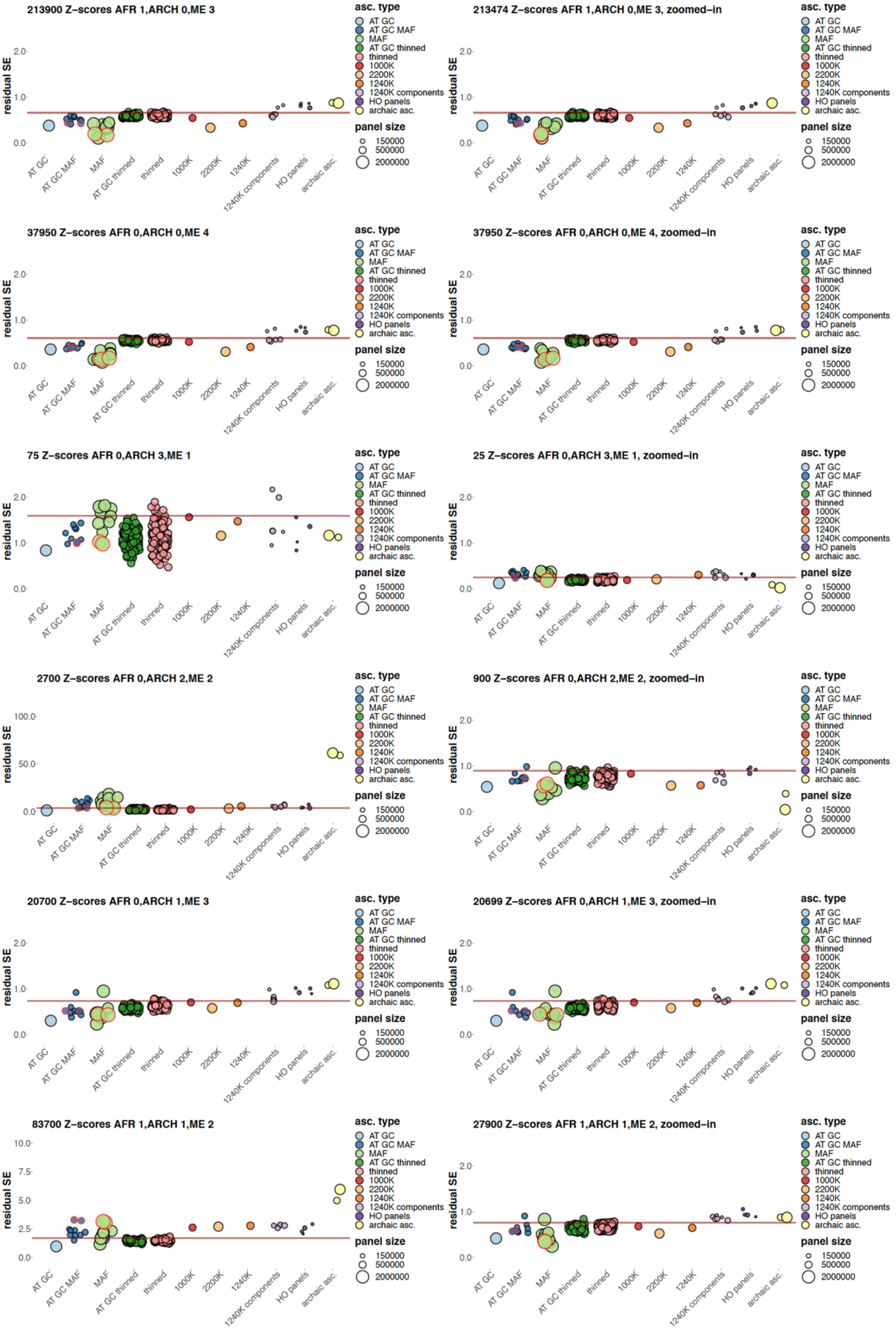
213900 Z–scores AFR 1,ARCH 0,ME 3
213474 Z–scores AFR 1,ARCH 0,ME 3, zoomed–in
37950 Z–scores AFR 0,ARCH 0,ME 4
37950 Z–scores AFR 0,ARCH 0,ME 4, zoomed–in
75 Z–scores AFR 0,ARCH 3,ME 1
25 Z–scores AFR 0,ARCH 3,ME 1, zoomed–in
2700 Z–scores AFR 0,ARCH 2,ME 2
900 Z–scores AFR 0,ARCH 2,ME 2, zoomed–in
20700 Z–scores AFR 0,ARCH 1,ME 3
20699 Z–scores AFR 0,ARCH 1,ME 3, zoomed–in
83700 Z–scores AFR 1,ARCH 1,ME 2
27900 Z–scores AFR 1,ARCH 1,ME 2, zoomed–in
residual SE
asc. type
AT GC
AT GC MAF
MAF
AT GC thinned
thinned
1000K
2200K
1240K
1240K components
HO panels
archaic asc.
panel size
150000
500000
2000000

**d**

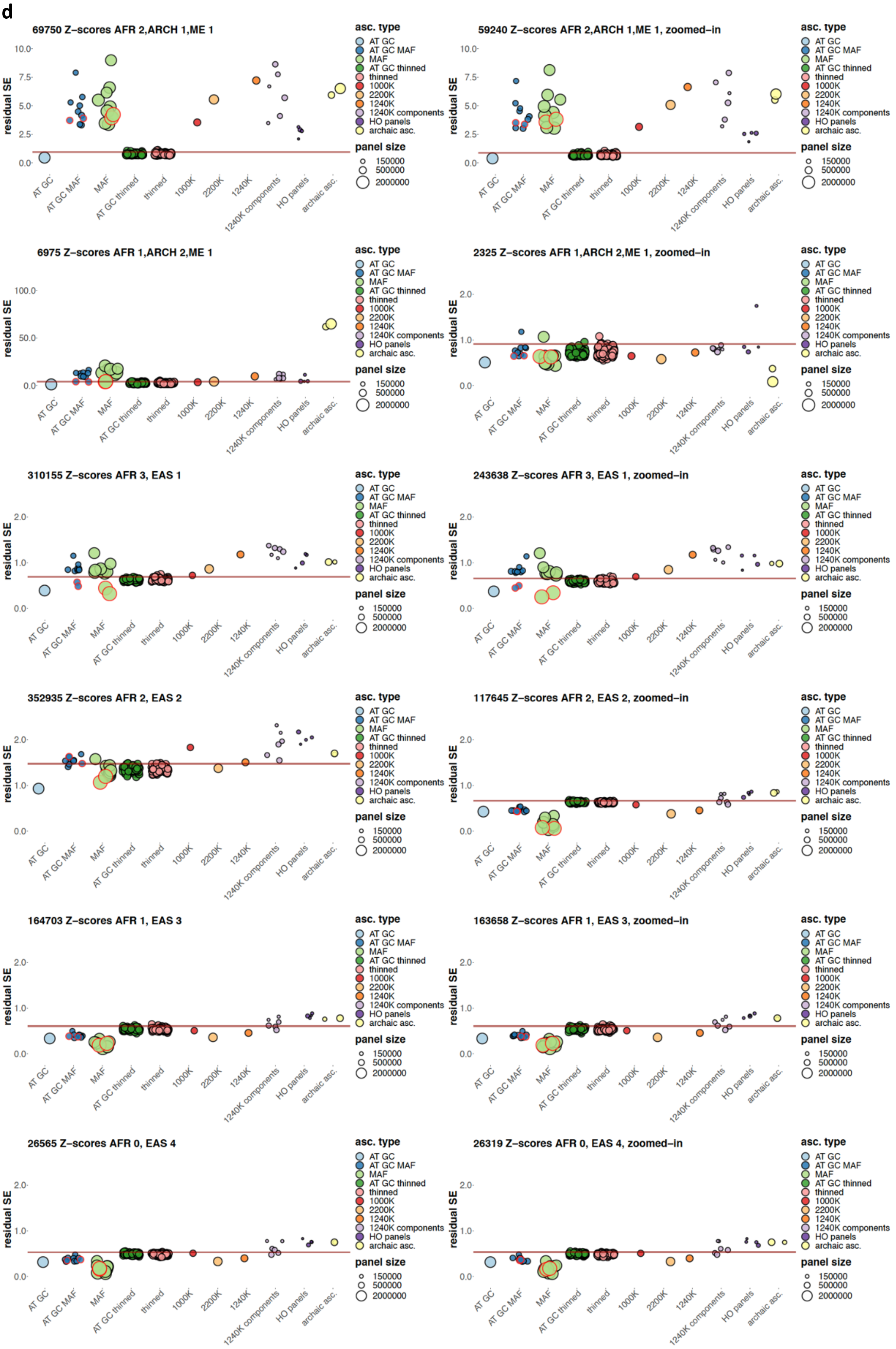
69750 Z–scores AFR 2,ARCH 1,ME 1
59240 Z–scores AFR 2,ARCH 1,ME 1, zoomed–in
6975 Z–scores AFR 1,ARCH 2,ME 1
2325 Z–scores AFR 1,ARCH 2,ME 1, zoomed–in
310155 Z–scores AFR 3, EAS 1
243638 Z–scores AFR 3, EAS 1, zoomed–in
352935 Z–scores AFR 2, EAS 2
117645 Z–scores AFR 2, EAS 2, zoomed–in
164703 Z–scores AFR 1, EAS 3
163658 Z–scores AFR 1, EAS 3, zoomed–in
26565 Z–scores AFR 0, EAS 4
26319 Z–scores AFR 0, EAS 4, zoomed–in
residual SE
AT GC
AT GC MAF
MAF
AT GC thinned
thinned
1000K
2200K
1240K
1240K components
HO panels
archaic asc.
asc. type
panel size
150000
500000
2000000

**e**

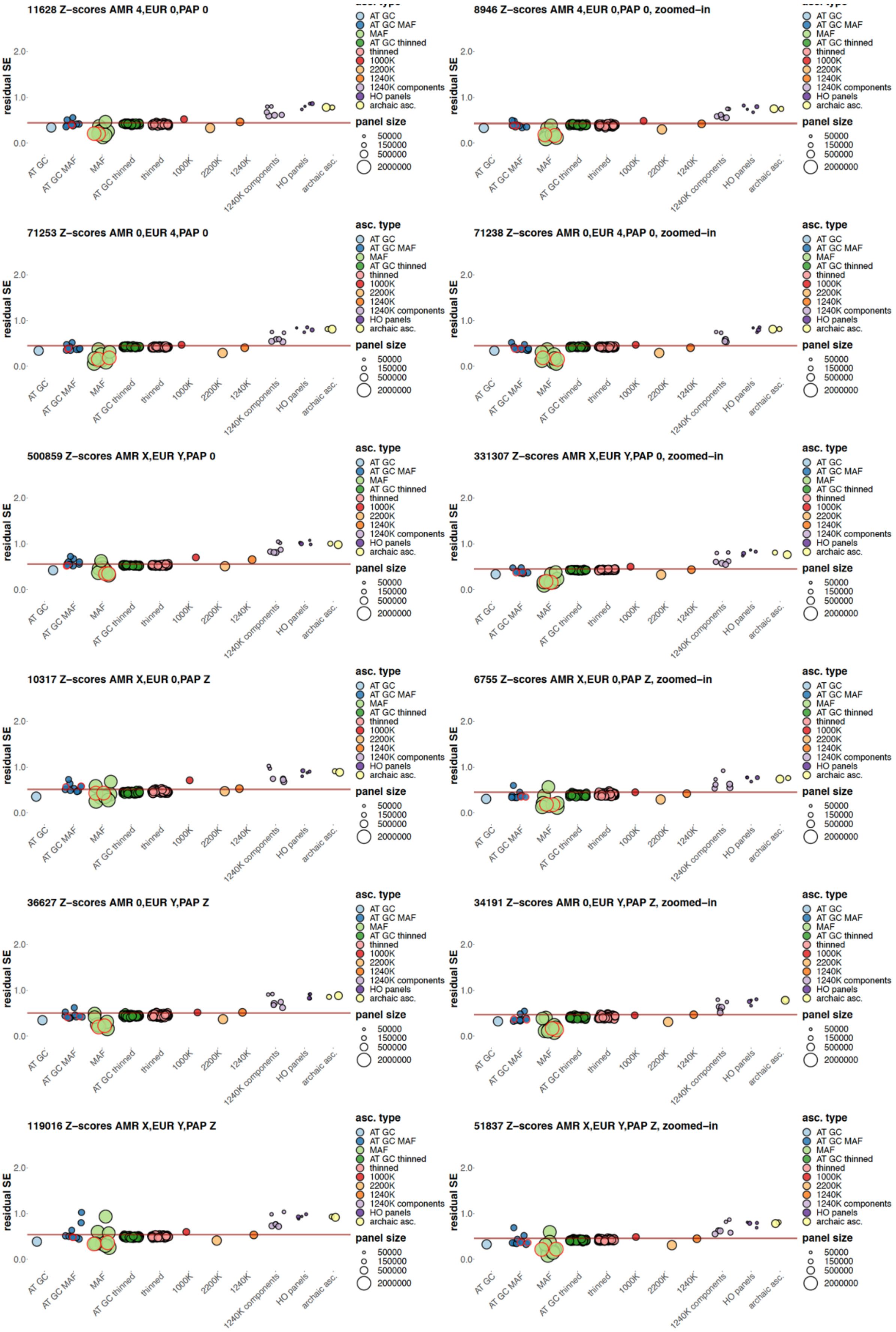
11628 Z–scores AMR 4,EUR 0,PAP 0
8946 Z–scores AMR 4,EUR 0,PAP 0, zoomed–in
71253 Z–scores AMR 0,EUR 4,PAP 0
71238 Z–scores AMR 0,EUR 4,PAP 0, zoomed–in
500859 Z–scores AMR X,EUR Y,PAP 0
331307 Z–scores AMR X,EUR Y,PAP 0, zoomed–in
10317 Z–scores AMR X,EUR 0,PAP Z
6755 Z–scores AMR X,EUR 0,PAP Z, zoomed–in
36627 Z–scores AMR 0,EUR Y,PAP Z
34191 Z–scores AMR 0,EUR Y,PAP Z, zoomed–in
119016 Z–scores AMR X,EUR Y,PAP Z
51837 Z–scores AMR X,EUR Y,PAP Z, zoomed–in
residual SE
0.0
1.0
2.0
AT GC
AT GC MAF
MAF
AT GC thinned
thinned
1000K
2200K
1240K
1240K components
HO panels
archaic asc.
asc. type
panel size
50000
150000
500000
2000000

**Suppl. Fig. 12.** Variance in $f_4$-statistic Z-scores resulting from ascertainment and random site subsampling expressed as standard deviations of residuals of a linear trend (expressed in the same units as $f_4$-statistic Z-scores). Results for ascertainment on variants common in Africans (either those having no detectable West Eurasian ancestry according to Fan et al. (2019) or on all Africans in the SGDP dataset) are circled in red. Results are shown for 27 classes of $f_4$-statistics indicated in plot titles in panels **a**-**e**. Only statistics of the form $f_4$(African X, archaic; African Y, non-African) were considered in the class African$_2$;archaic$_1$;Med/ME$_1$. Residual SE values for $f_4$-statistic Z-scores lying not far from 0 (absolute Z-scores on all sites < 15) are plotted in the right-hand panels, and residual SE values for all Z-scores are plotted in the left-hand panels. The 97.5% percentiles of all the thinned replicates combined, including those on all sites and AT/GC sites, are marked by the brown lines. Most y-axis scales are the same (from 0 to 2.5 SE), except for few exceptions in panels **a**, **b**, **c**, and **d**. Size of the resulting SNP panels is coded by point size, and ten broad ascertainment types are coded by color according to the legend. Thirty eight site subsampling schemes were explored: 1) AT/GC mutation classes; 2) AT/GC mutation classes and restricting to common variants based on a global MAF threshold of 5% or on the same threshold in one of nine continental-scale groups; 3) the same procedure repeated on all sites; 4) random thinning of the AT/GC dataset to the 1240K SNP count for a given combination of groups (no missing data allowed), results for 100 thinning replicates are shown; 5) random thinning of all sites to the 1240K SNP count, results for 100 thinning replicates are shown; 6) the 1000K and 2200K enrichment panels (2200K = 1240K + 1000K); 7) the 1240K panel; 8) major components of the 1240K panel: sites included in the Illumina 650Y and/or Human Origins SNP arrays, sites included exclusively in one of them, and remaining sites; 9) the largest Human Origins sub-panels (4, 5, 13) or their union (4+5); 10) restricting to sites polymorphic in a group composed of the three high-coverage archaic individuals (either all such sites or transversions only).

a

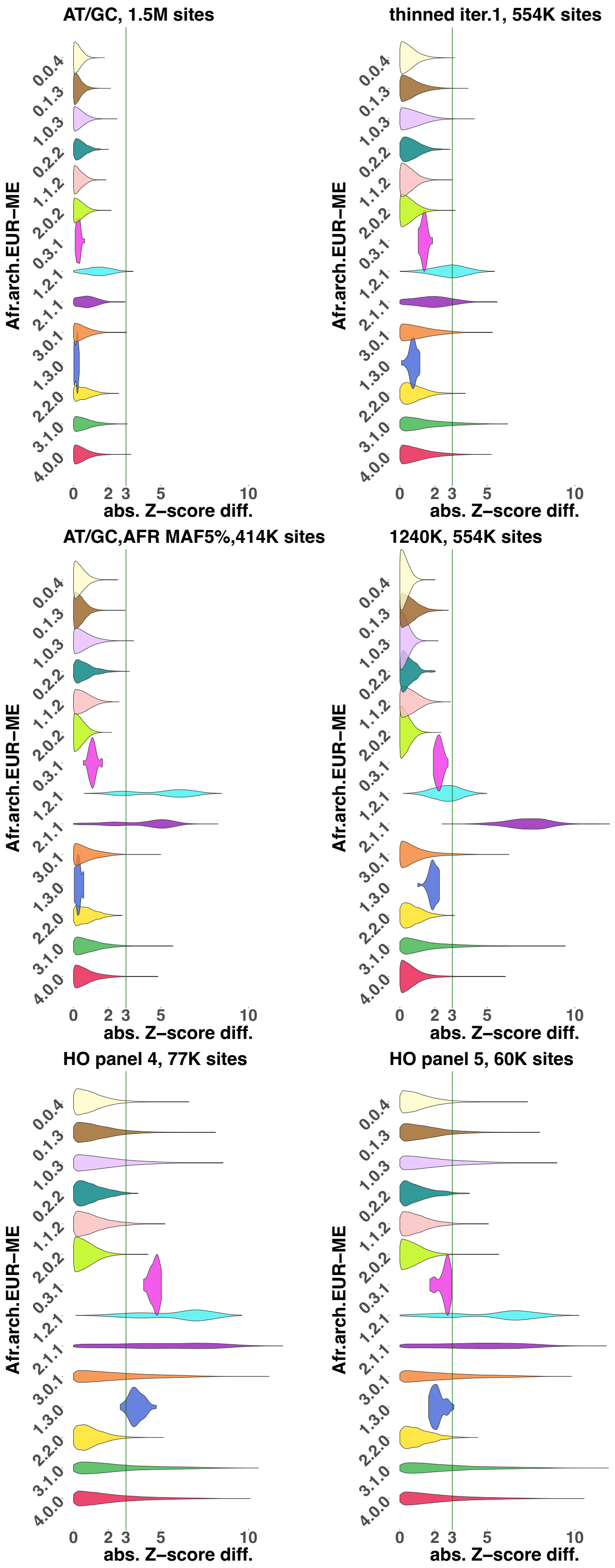
AT/GC, 1.5M sites
thinned iter.1, 554K sites
AT/GC,AFR MAF5%,414K sites
1240K, 554K sites
HO panel 4, 77K sites
HO panel 5, 60K sites
Afr.arch.EUR−ME
0.0.4
0.1.3
1.0.3
0.2.2
1.1.2
2.0.2
0.3.1
1.2.1
2.1.1
3.0.1
1.3.0
2.2.0
3.1.0
4.0.0
0
2
3
5
10
abs. Z−score diff.

**b**

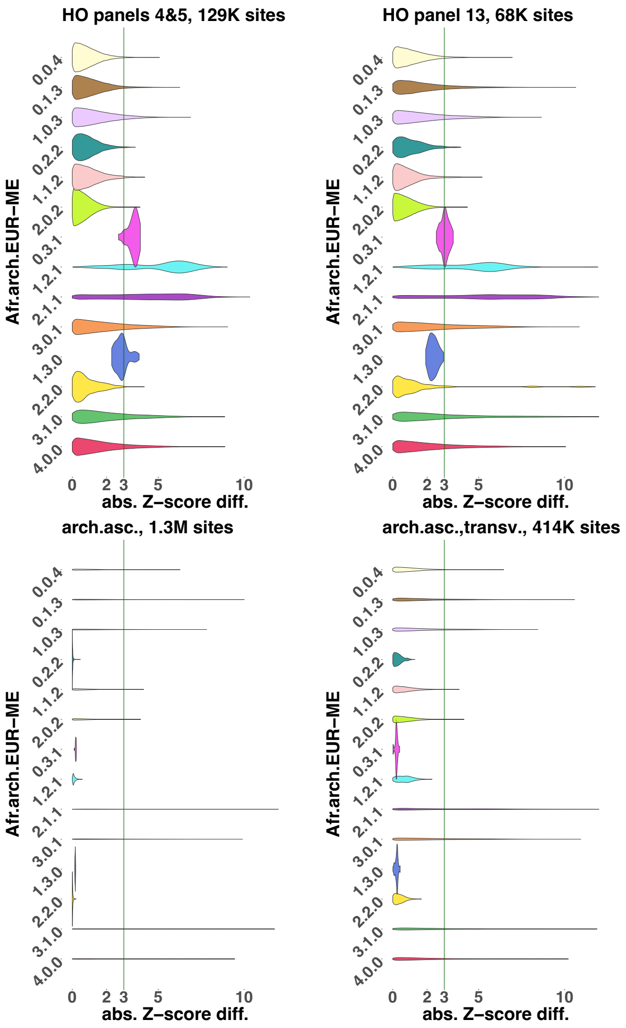

**Suppl. Fig. 13.** Violin plots illustrating the effects of 10 ascertainment schemes on Z-scores of 14 $f_4$-statistic classes that were explored exhaustively: statistics including African and/or archaic and or Mediterranean/Middle Eastern groups. Only statistics of the form $f_4$(African X, archaic; African Y, non-African) were considered in the class African$_2$;archaic$_1$;Med/ME$_1$. $f_4$-statistic classes are labelled on y-axes. Z-score difference was calculated as the Z-score on all sites (ca. 10 million sites) minus the Z-score on an ascertained dataset, and the distributions of absolute difference values are visualized. Ascertainment types and site counts are shown in plot titles. Only statistics with absolute Z-scores below 15 on all sites were considered for this analysis.

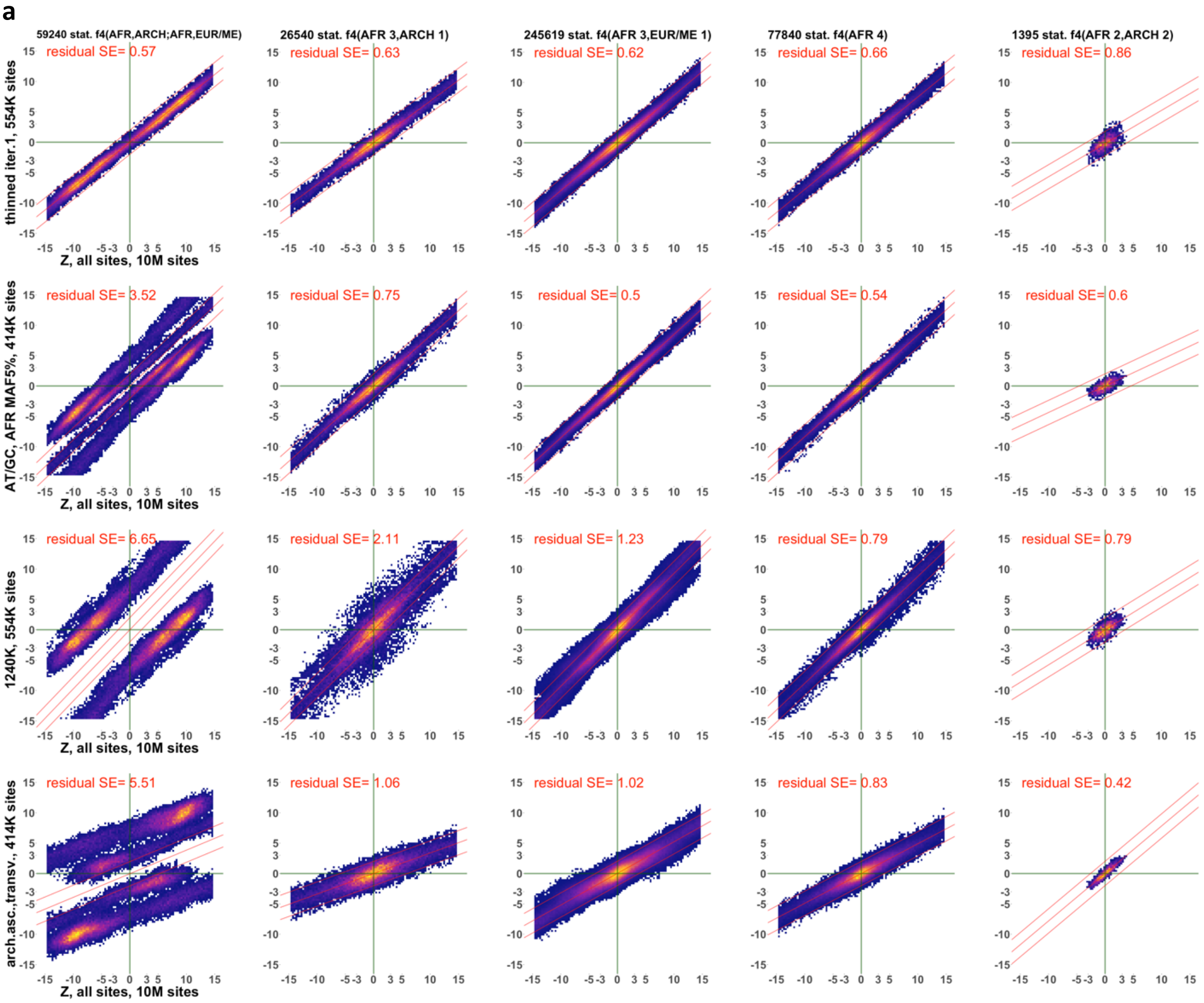
a
59240 stat. f4(AFR,ARCH;AFR,EUR/ME)
26540 stat. f4(AFR 3,ARCH 1)
245619 stat. f4(AFR 3,EUR/ME 1)
77840 stat. f4(AFR 4)
1395 stat. f4(AFR 2,ARCH 2)
thinned iter.1, 554K sites
AT/GC, AFR MAF5%, 414K sites
1240K, 554K sites
arch.asc.,transv., 414K sites
Z, all sites, 10M sites
residual SE= 0.57
residual SE= 0.63
residual SE= 0.62
residual SE= 0.66
residual SE= 0.86
residual SE= 3.52
residual SE= 0.75
residual SE= 0.5
residual SE= 0.54
residual SE= 0.6
residual SE= 6.65
residual SE= 2.11
residual SE= 1.23
residual SE= 0.79
residual SE= 0.79
residual SE= 5.51
residual SE= 1.06
residual SE= 1.02
residual SE= 0.83
residual SE= 0.42

**b**

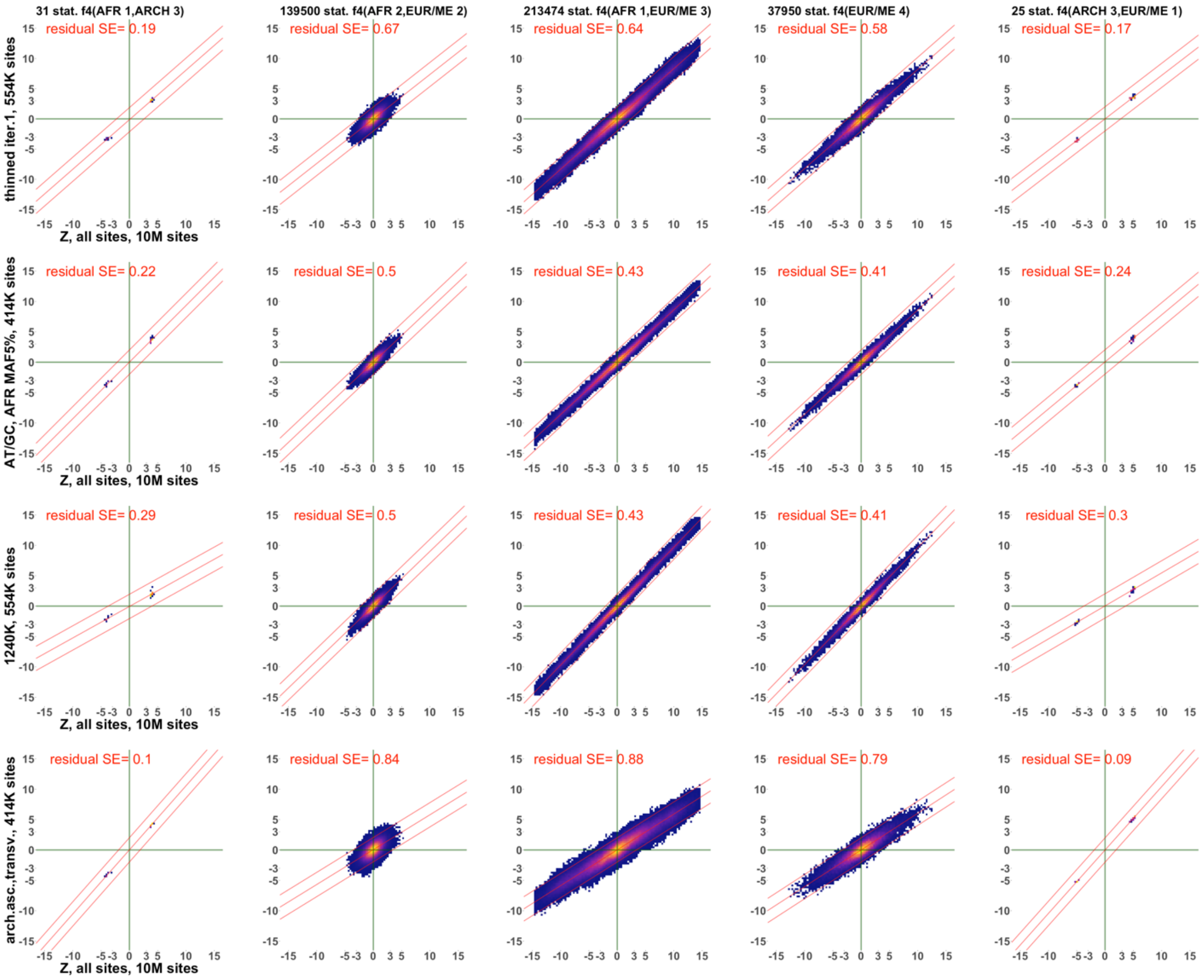
31 stat. f4(AFR 1,ARCH 3)
139500 stat. f4(AFR 2,EUR/ME 2)
213474 stat. f4(AFR 1,EUR/ME 3)
37950 stat. f4(EUR/ME 4)
25 stat. f4(ARCH 3,EUR/ME 1)
thinned iter.1, 554K sites
residual SE= 0.19
residual SE= 0.67
residual SE= 0.64
residual SE= 0.58
residual SE= 0.17
AT/GC, AFR MAF5%, 414K sites
residual SE= 0.22
residual SE= 0.5
residual SE= 0.43
residual SE= 0.41
residual SE= 0.24
1240K, 554K sites
residual SE= 0.29
residual SE= 0.5
residual SE= 0.43
residual SE= 0.41
residual SE= 0.3
arch.asc.,transv., 414K sites
residual SE= 0.1
residual SE= 0.84
residual SE= 0.88
residual SE= 0.79
residual SE= 0.09
Z, all sites, 10M sites

**c**

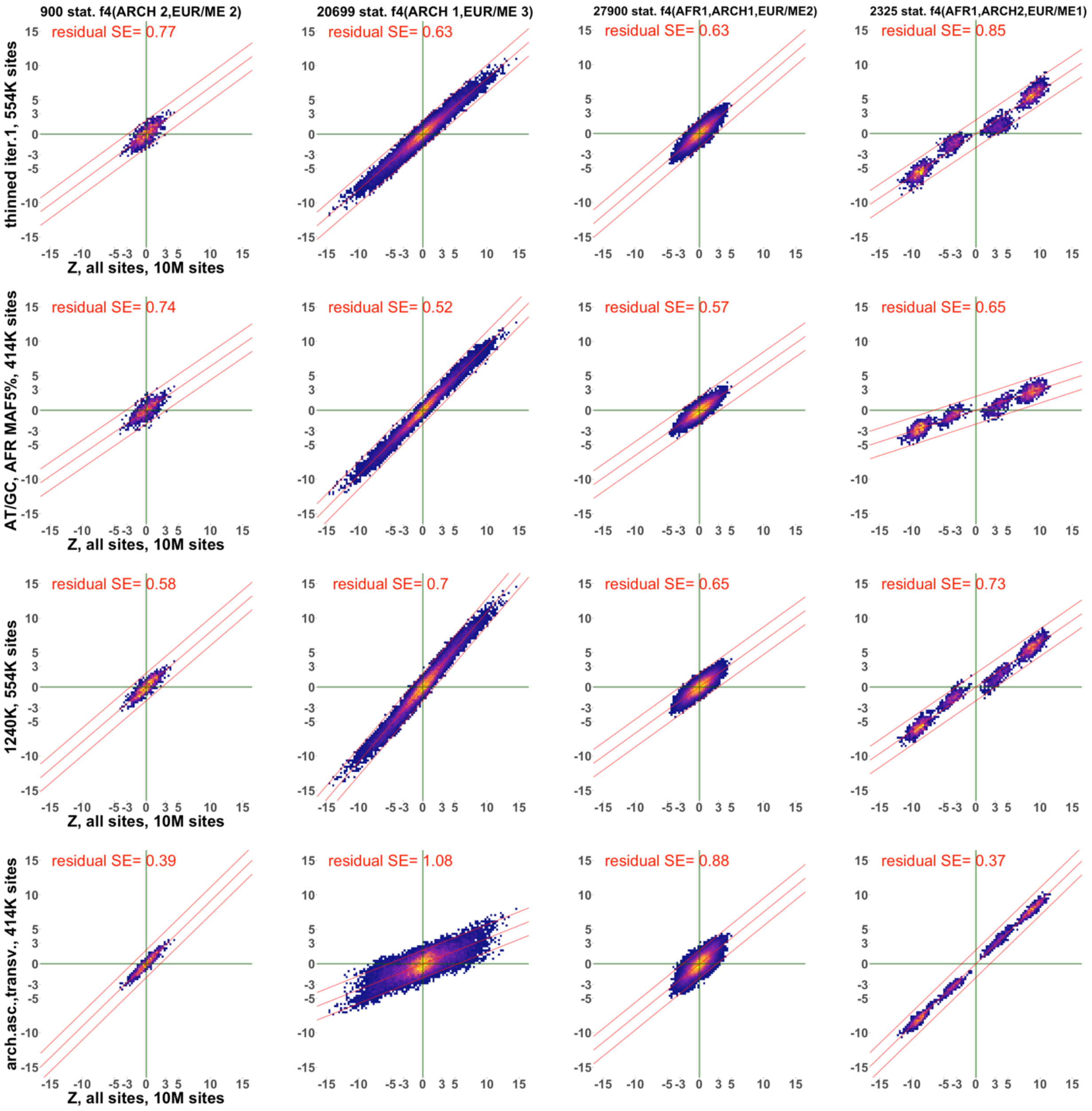

**Suppl. Fig. 14.** Scatterplots illustrating the effects of four ascertainment schemes on Z-scores of $f_4$-statistics of 14 classes including African and/or archaic and/or Mediterranean/Middle Eastern groups (abbreviated as Med/ME). Only statistics of the form $f_4$(African X, archaic; African Y, non-African) were considered in the class African$_2$;archaic$_1$;Med/ME$_1$. The class labels and numbers of statistics plotted are shown in the top row of each panel (**a**, **b**, and **c**). Instead of individual points, heatmaps illustrating point density are shown. Z-scores on all sites (ca. 10 million sites, as indicated on the x-axes) are compared to Z-scores on ascertained datasets on the y-axes. Ascertainment types and site counts are shown on the y-axes. All plots include only statistics with absolute Z-scores below 15 on all sites. A linear model fitted to the data and lines representing ± 2 SE are shown in red. Residual SE values for those linear trends are shown in each plot in red.

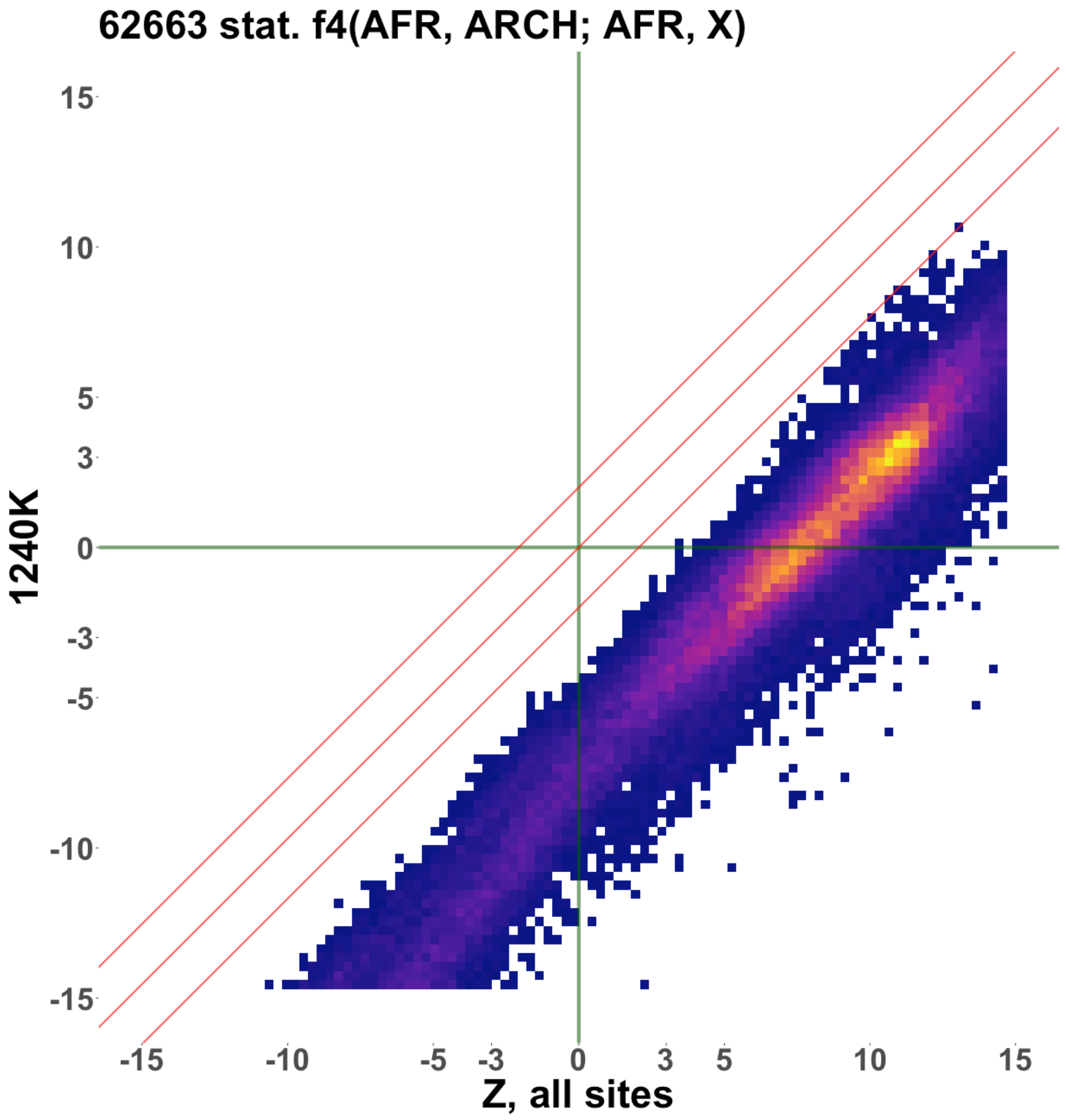

**Suppl. Fig. 15.** A scatterplot illustrating the effect of the 1240K ascertainment on Z-scores of $f_4$-statistics (African X, archaic; African Y, X), where X is any non-African except for the Mediterranean and Middle Eastern groups (results for those groups are shown in Suppl. Fig. 14). Instead of individual points, heatmaps illustrating point density are shown. Z-scores on all sites are compared to Z-scores on the 1240K dataset on the y-axis (site counts varied across population quadruplets). The plot includes only statistics with absolute Z-scores below 15 on all sites. A linear model fitted to the data and lines representing ± 2 SE are shown in red. The intercept of the linear trend was set at 0.

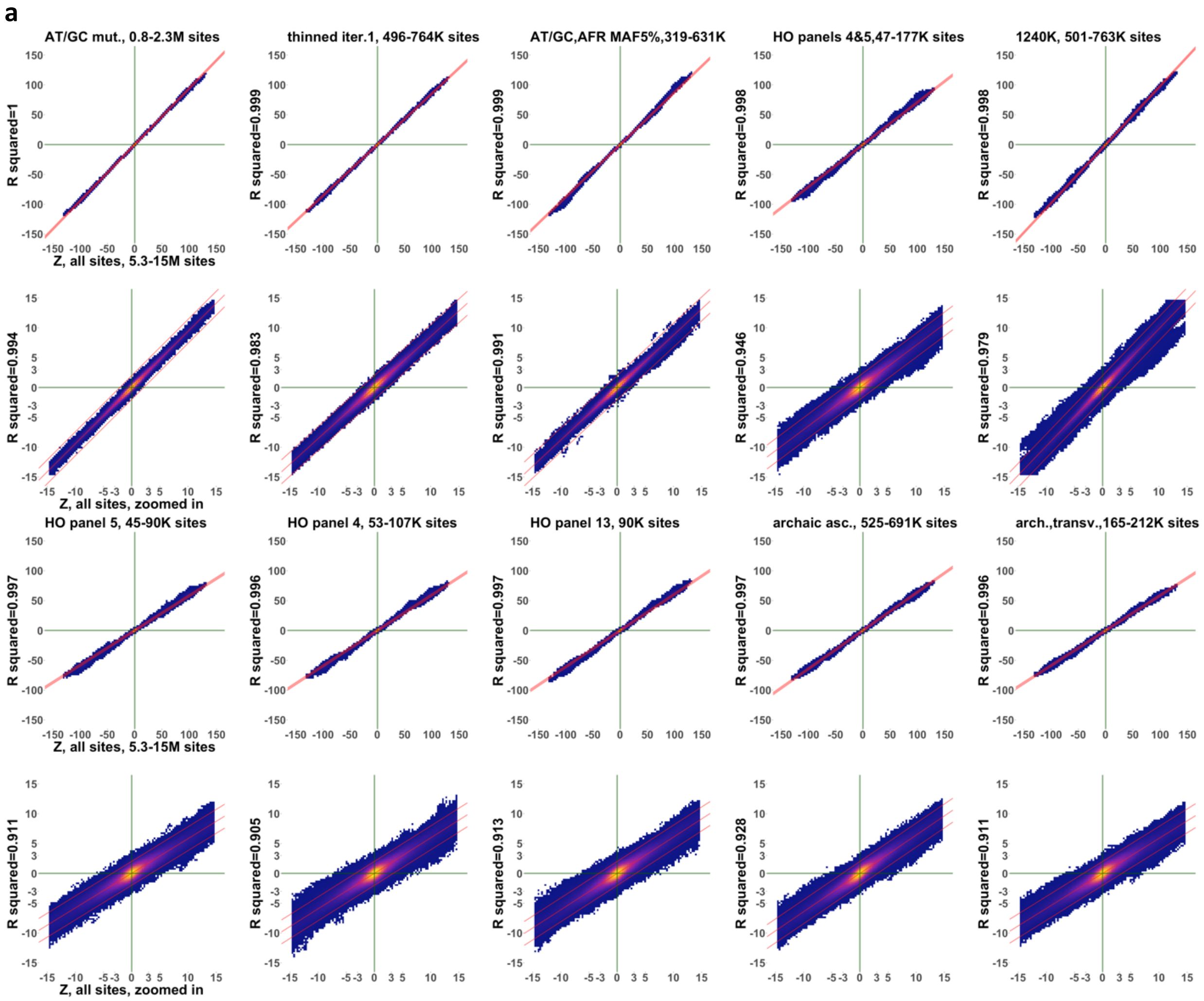
a
AT/GC mut., 0.8-2.3M sites
thinned iter.1, 496-764K sites
AT/GC,AFR MAF5%,319-631K
HO panels 4&5,47-177K sites
1240K, 501-763K sites
R squared=1
R squared=0.999
R squared=0.999
R squared=0.998
R squared=0.998
Z, all sites, 5.3-15M sites
R squared=0.994
R squared=0.983
R squared=0.991
R squared=0.946
R squared=0.979
Z, all sites, zoomed in
HO panel 5, 45-90K sites
HO panel 4, 53-107K sites
HO panel 13, 90K sites
archaic asc., 525-691K sites
arch.,transv.,165-212K sites
R squared=0.997
R squared=0.996
R squared=0.997
R squared=0.997
R squared=0.996
Z, all sites, 5.3-15M sites
R squared=0.911
R squared=0.905
R squared=0.913
R squared=0.928
R squared=0.911
Z, all sites, zoomed in

**b**

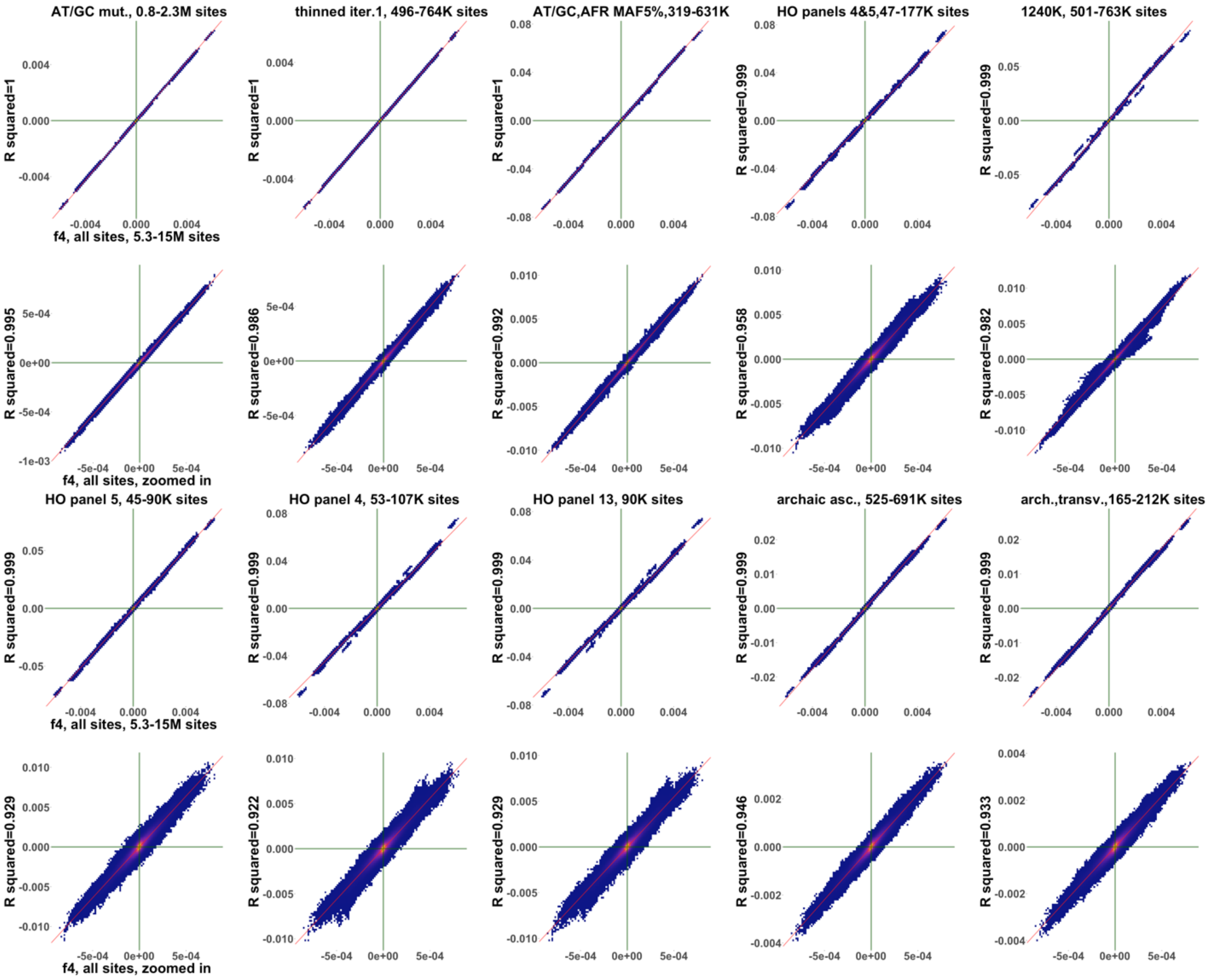

**Suppl. Fig. 16.** Scatterplots illustrating the effects of 10 ascertainment schemes on Z-scores (**a**) or $f_4$-statistics (**b**) of two classes: 854,358 distinct statistics including up to three African groups and/or up to four East Asian groups, pooled with 749,700 distinct statistics including American and/or European and/or Papuan groups. Instead of individual points, heatmaps illustrating point density are shown. Z-scores on all sites (from ca. 5.3 to 15 million sites, as indicated on the x-axes) are compared to Z-scores on ascertained datasets on the y-axes. Ascertainment types and site counts are shown in plot titles. The 2$^{nd}$ and 4$^{th}$ rows of plots in each panel represent close-up views on the origin of the plots: statistics with absolute Z-scores below 15 on all sites. Linear models fitted to the data and lines representing ± 2 SE are shown in red. $R^2$ values for those models are shown on the y-axes. First, linear models were fitted to the sets of Z-scores or $f_4$-statistics lying near the origin (having absolute Z-score below 15 on all sites), and then the same linear equations were re-fitted to complete sets of Z-scores or $f_4$-statistics. Since we show both $f_4$-statistics and Z-scores side by side here, $R^2$ was used instead of residual SE as a measure of correlation.

**Suppl. Fig. 17.**

**a**

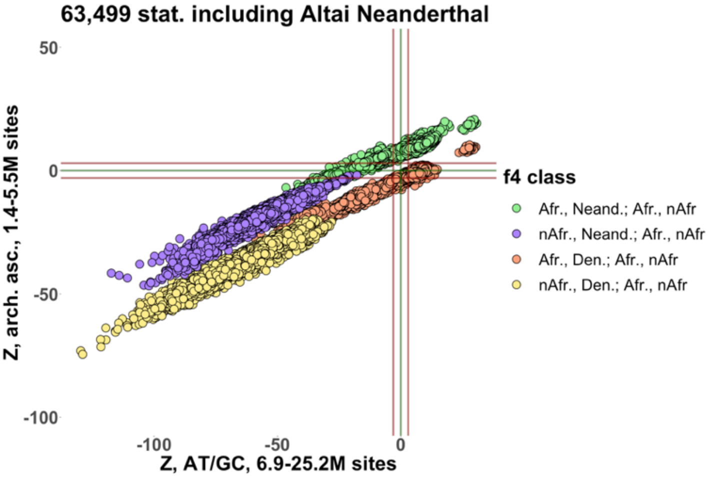

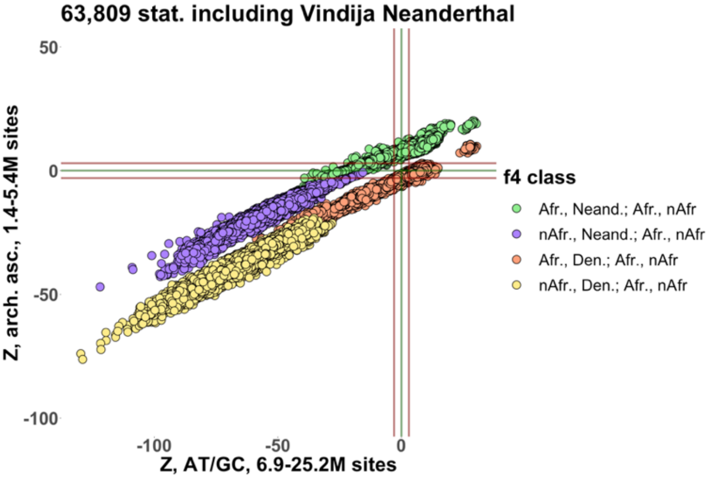

**b**

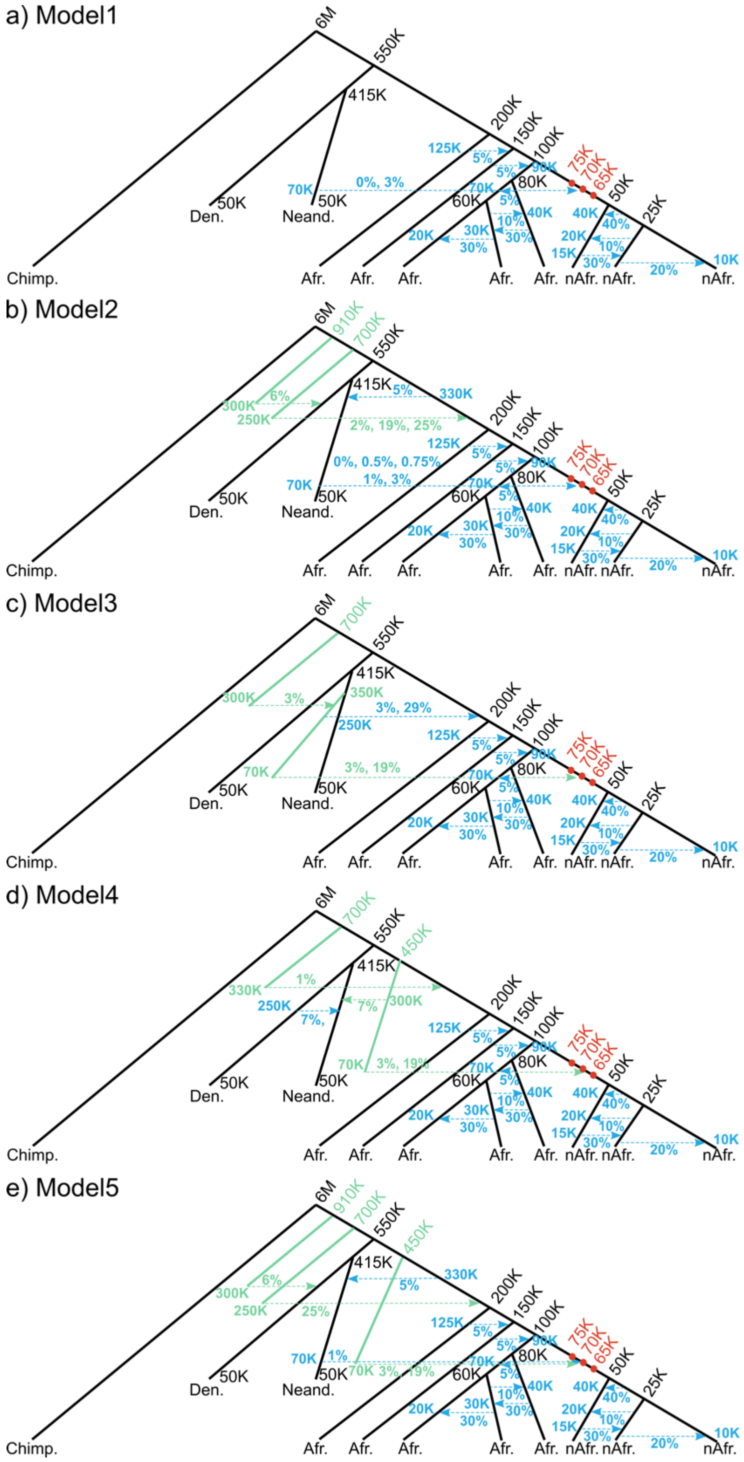
a) Model1
b) Model2
c) Model3
d) Model4
e) Model5
6M
550K
415K
200K
150K
100K
80K
60K
50K
25K
75K
70K
65K
125K
70K
40K
30K
20K
15K
10K
910K
700K
450K
330K
300K
250K
350K
0%, 3%
2%, 19%, 25%
0%, 0.5%, 0.75%
1%, 3%
3%, 29%
3%, 19%
Chimp.
Den.
Neand.
Afr.
nAfr.

c

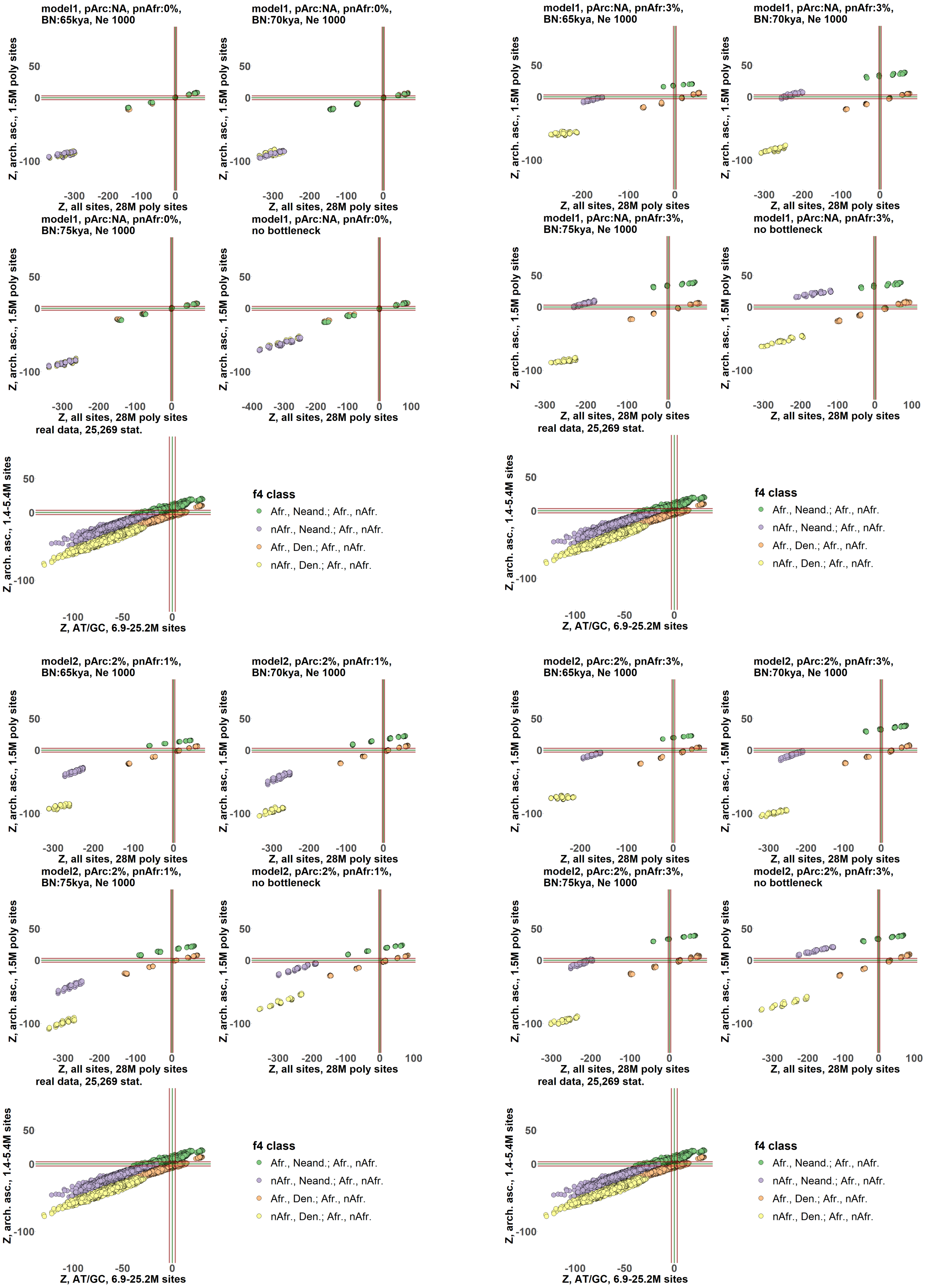

model1, pArc:NA, pnAfr:0%, BN:65kya, Ne 1000
model1, pArc:NA, pnAfr:0%, BN:70kya, Ne 1000
model1, pArc:NA, pnAfr:3%, BN:65kya, Ne 1000
model1, pArc:NA, pnAfr:3%, BN:70kya, Ne 1000
model1, pArc:NA, pnAfr:0%, BN:75kya, Ne 1000
model1, pArc:NA, pnAfr:0%, no bottleneck
model1, pArc:NA, pnAfr:3%, BN:75kya, Ne 1000
model1, pArc:NA, pnAfr:3%, no bottleneck
real data, 25,269 stat.
model2, pArc:2%, pnAfr:1%, BN:65kya, Ne 1000
model2, pArc:2%, pnAfr:1%, BN:70kya, Ne 1000
model2, pArc:2%, pnAfr:3%, BN:65kya, Ne 1000
model2, pArc:2%, pnAfr:3%, BN:70kya, Ne 1000
model2, pArc:2%, pnAfr:1%, BN:75kya, Ne 1000
model2, pArc:2%, pnAfr:1%, no bottleneck
model2, pArc:2%, pnAfr:3%, BN:75kya, Ne 1000
model2, pArc:2%, pnAfr:3%, no bottleneck
Z, arch. asc., 1.5M poly sites
Z, all sites, 28M poly sites
Z, arch. asc., 1.4-5.4M sites
Z, AT/GC, 6.9-25.2M sites
f4 class
Afr., Neand.; Afr., nAfr.
nAfr., Neand.; Afr., nAfr.
Afr., Den.; Afr., nAfr.
nAfr., Den.; Afr., nAfr.

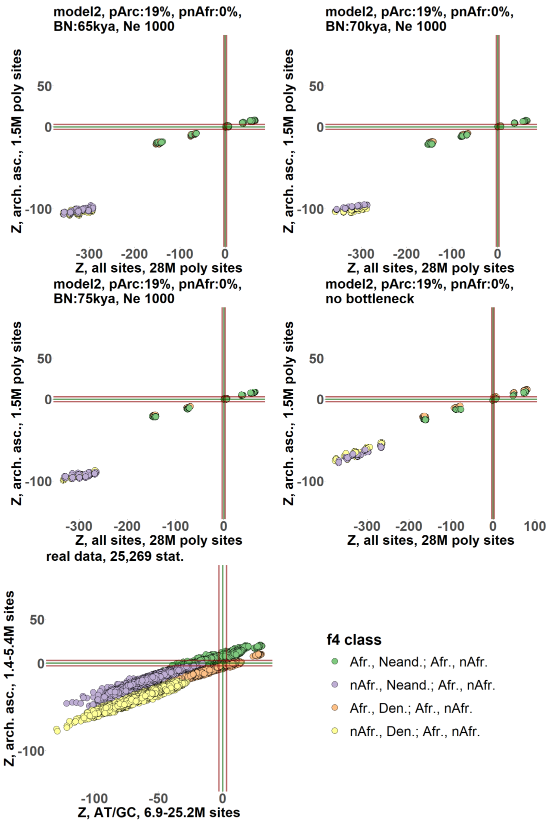
model2, pArc:19%, pnAfr:0%, BN:65kya, Ne 1000
model2, pArc:19%, pnAfr:0%, BN:70kya, Ne 1000
model2, pArc:19%, pnAfr:0%, BN:75kya, Ne 1000
model2, pArc:19%, pnAfr:0%, no bottleneck
real data, 25,269 stat.
Z, arch. asc., 1.5M poly sites
Z, all sites, 28M poly sites
Z, arch. asc., 1.4-5.4M sites
Z, AT/GC, 6.9-25.2M sites
f4 class
Afr., Neand.; Afr., nAfr.
nAfr., Neand.; Afr., nAfr.
Afr., Den.; Afr., nAfr.
nAfr., Den.; Afr., nAfr.

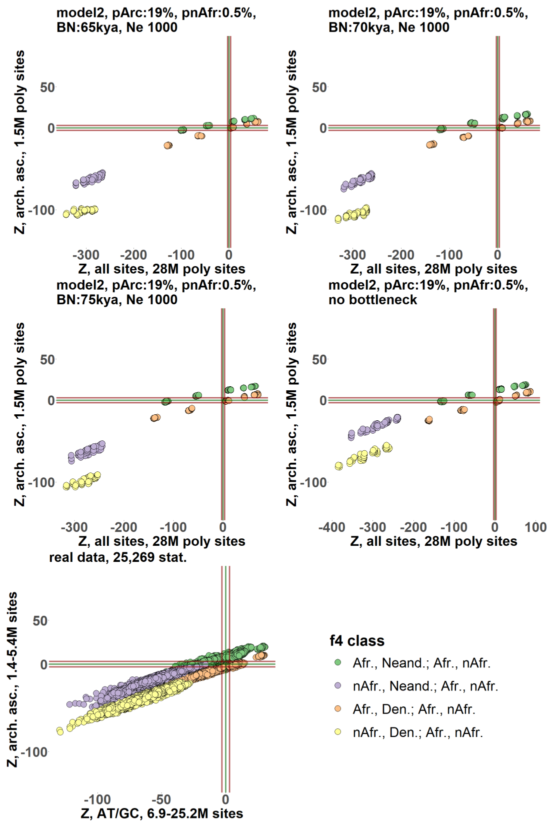
model2, pArc:19%, pnAfr:0.5%, BN:65kya, Ne 1000
model2, pArc:19%, pnAfr:0.5%, BN:70kya, Ne 1000
model2, pArc:19%, pnAfr:0.5%, BN:75kya, Ne 1000
model2, pArc:19%, pnAfr:0.5%, no bottleneck
real data, 25,269 stat.
Z, arch. asc., 1.5M poly sites
Z, all sites, 28M poly sites
Z, arch. asc., 1.4-5.4M sites
Z, AT/GC, 6.9-25.2M sites
f4 class
Afr., Neand.; Afr., nAfr.
nAfr., Neand.; Afr., nAfr.
Afr., Den.; Afr., nAfr.
nAfr., Den.; Afr., nAfr.

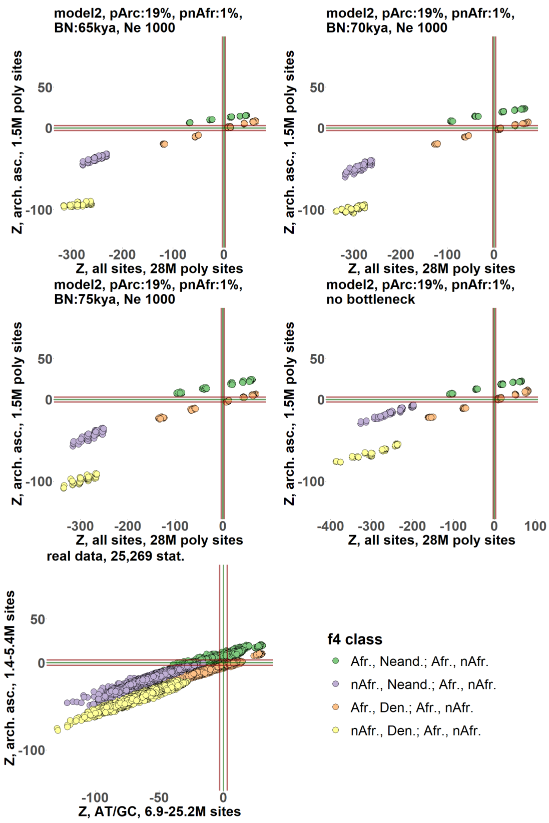
model2, pArc:19%, pnAfr:1%, BN:65kya, Ne 1000
model2, pArc:19%, pnAfr:1%, BN:70kya, Ne 1000
model2, pArc:19%, pnAfr:1%, BN:75kya, Ne 1000
model2, pArc:19%, pnAfr:1%, no bottleneck
real data, 25,269 stat.
Z, arch. asc., 1.5M poly sites
Z, all sites, 28M poly sites
Z, arch. asc., 1.4-5.4M sites
Z, AT/GC, 6.9-25.2M sites
f4 class
Afr., Neand.; Afr., nAfr.
nAfr., Neand.; Afr., nAfr.
Afr., Den.; Afr., nAfr.
nAfr., Den.; Afr., nAfr.

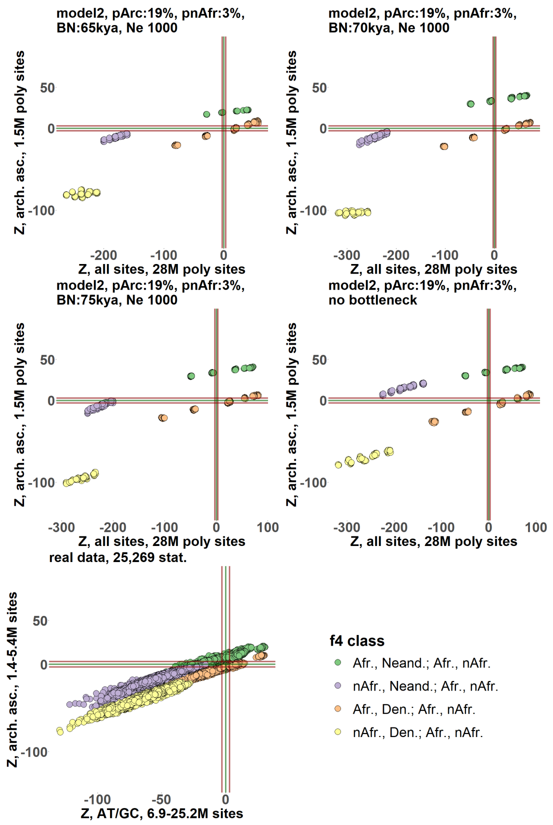
model2, pArc:19%, pnAfr:3%, BN:65kya, Ne 1000
model2, pArc:19%, pnAfr:3%, BN:70kya, Ne 1000
model2, pArc:19%, pnAfr:3%, BN:75kya, Ne 1000
model2, pArc:19%, pnAfr:3%, no bottleneck
real data, 25,269 stat.
Z, arch. asc., 1.5M poly sites
Z, all sites, 28M poly sites
Z, arch. asc., 1.4-5.4M sites
Z, AT/GC, 6.9-25.2M sites
f4 class
Afr., Neand.; Afr., nAfr.
nAfr., Neand.; Afr., nAfr.
Afr., Den.; Afr., nAfr.
nAfr., Den.; Afr., nAfr.

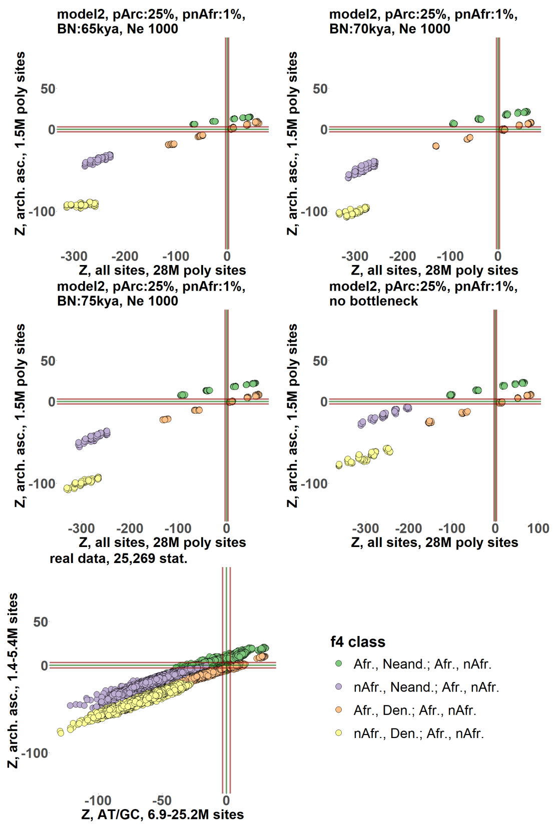
model2, pArc:25%, pnAfr:1%, BN:65kya, Ne 1000
model2, pArc:25%, pnAfr:1%, BN:70kya, Ne 1000
model2, pArc:25%, pnAfr:1%, BN:75kya, Ne 1000
model2, pArc:25%, pnAfr:1%, no bottleneck
Z, arch. asc., 1.5M poly sites
Z, all sites, 28M poly sites
real data, 25,269 stat.
Z, arch. asc., 1.4-5.4M sites
Z, AT/GC, 6.9-25.2M sites
f4 class
Afr., Neand.; Afr., nAfr.
nAfr., Neand.; Afr., nAfr.
Afr., Den.; Afr., nAfr.
nAfr., Den.; Afr., nAfr.

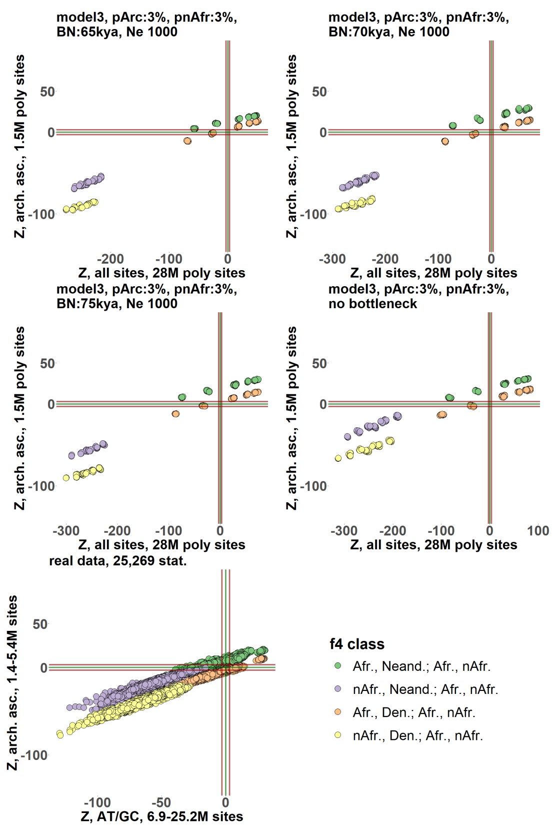
model3, pArc:3%, pnAfr:3%, BN:65kya, Ne 1000
model3, pArc:3%, pnAfr:3%, BN:70kya, Ne 1000
model3, pArc:3%, pnAfr:3%, BN:75kya, Ne 1000
model3, pArc:3%, pnAfr:3%, no bottleneck
Z, arch. asc., 1.5M poly sites
Z, all sites, 28M poly sites
real data, 25,269 stat.
Z, arch. asc., 1.4-5.4M sites
Z, AT/GC, 6.9-25.2M sites
f4 class
Afr., Neand.; Afr., nAfr.
nAfr., Neand.; Afr., nAfr.
Afr., Den.; Afr., nAfr.
nAfr., Den.; Afr., nAfr.

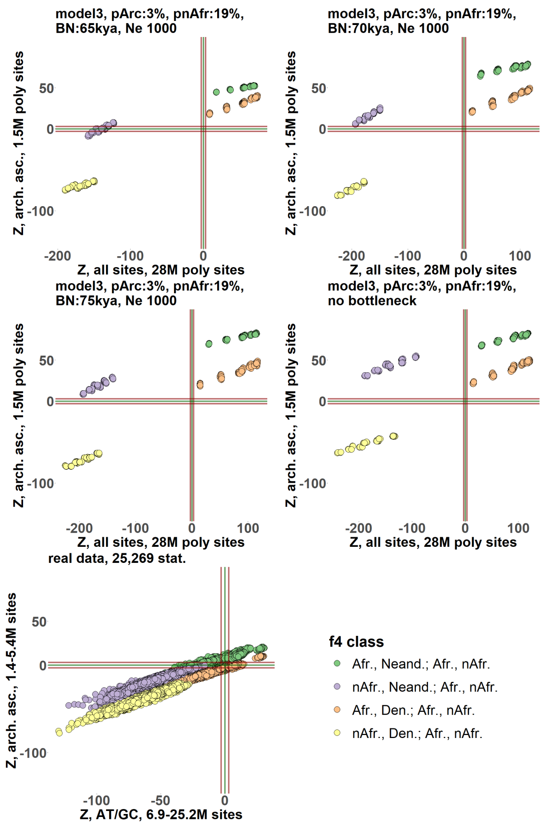
model3, pArc:3%, pnAfr:19%, BN:65kya, Ne 1000
model3, pArc:3%, pnAfr:19%, BN:70kya, Ne 1000
model3, pArc:3%, pnAfr:19%, BN:75kya, Ne 1000
model3, pArc:3%, pnAfr:19%, no bottleneck
Z, arch. asc., 1.5M poly sites
Z, all sites, 28M poly sites
real data, 25,269 stat.
Z, arch. asc., 1.4-5.4M sites
Z, AT/GC, 6.9-25.2M sites
f4 class
Afr., Neand.; Afr., nAfr.
nAfr., Neand.; Afr., nAfr.
Afr., Den.; Afr., nAfr.
nAfr., Den.; Afr., nAfr.

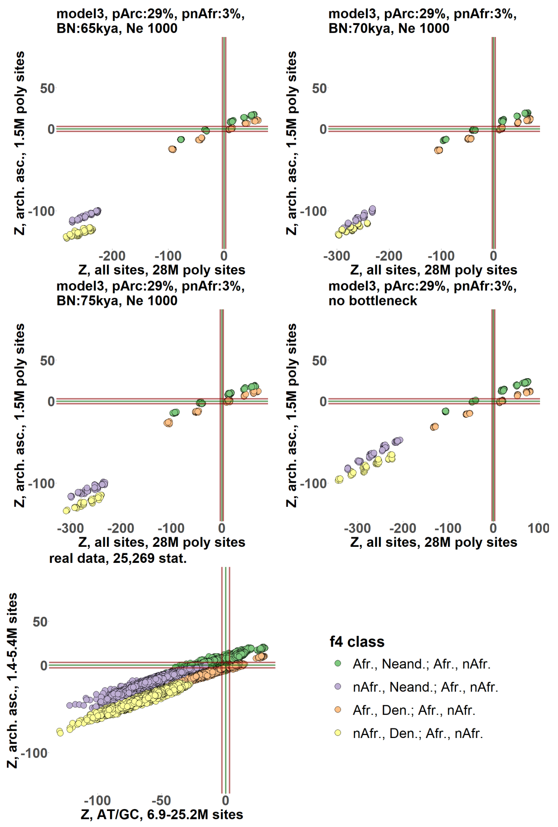
model3, pArc:29%, pnAfr:3%, BN:65kya, Ne 1000
model3, pArc:29%, pnAfr:3%, BN:70kya, Ne 1000
model3, pArc:29%, pnAfr:3%, BN:75kya, Ne 1000
model3, pArc:29%, pnAfr:3%, no bottleneck
Z, arch. asc., 1.5M poly sites
Z, all sites, 28M poly sites
real data, 25,269 stat.
Z, arch. asc., 1.4-5.4M sites
Z, AT/GC, 6.9-25.2M sites
f4 class
Afr., Neand.; Afr., nAfr.
nAfr., Neand.; Afr., nAfr.
Afr., Den.; Afr., nAfr.
nAfr., Den.; Afr., nAfr.

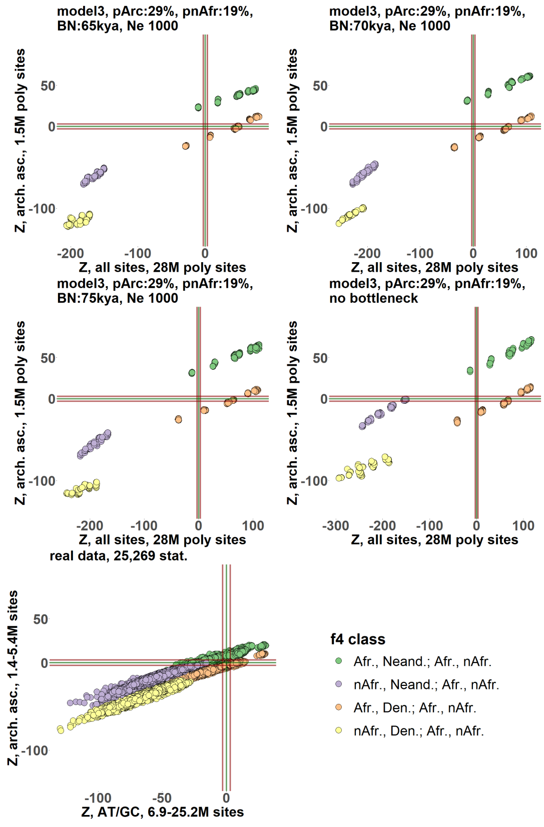
model3, pArc:29%, pnAfr:19%, BN:65kya, Ne 1000
model3, pArc:29%, pnAfr:19%, BN:70kya, Ne 1000
model3, pArc:29%, pnAfr:19%, BN:75kya, Ne 1000
model3, pArc:29%, pnAfr:19%, no bottleneck
Z, arch. asc., 1.5M poly sites
Z, all sites, 28M poly sites
real data, 25,269 stat.
Z, arch. asc., 1.4-5.4M sites
Z, AT/GC, 6.9-25.2M sites
f4 class
Afr., Neand.; Afr., nAfr.
nAfr., Neand.; Afr., nAfr.
Afr., Den.; Afr., nAfr.
nAfr., Den.; Afr., nAfr.

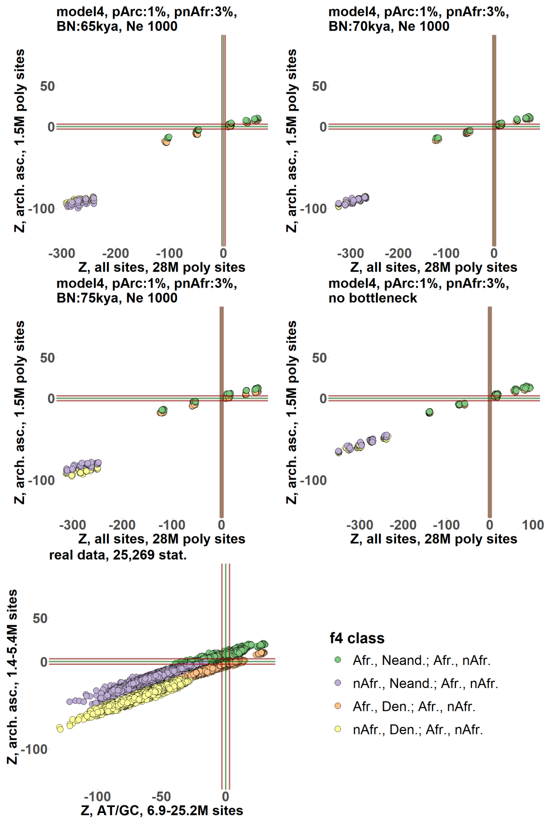
model4, pArc:1%, pnAfr:3%, BN:65kya, Ne 1000
model4, pArc:1%, pnAfr:3%, BN:70kya, Ne 1000
model4, pArc:1%, pnAfr:3%, BN:75kya, Ne 1000
model4, pArc:1%, pnAfr:3%, no bottleneck
Z, arch. asc., 1.5M poly sites
Z, all sites, 28M poly sites
real data, 25,269 stat.
Z, arch. asc., 1.4-5.4M sites
Z, AT/GC, 6.9-25.2M sites
f4 class
Afr., Neand.; Afr., nAfr.
nAfr., Neand.; Afr., nAfr.
Afr., Den.; Afr., nAfr.
nAfr., Den.; Afr., nAfr.

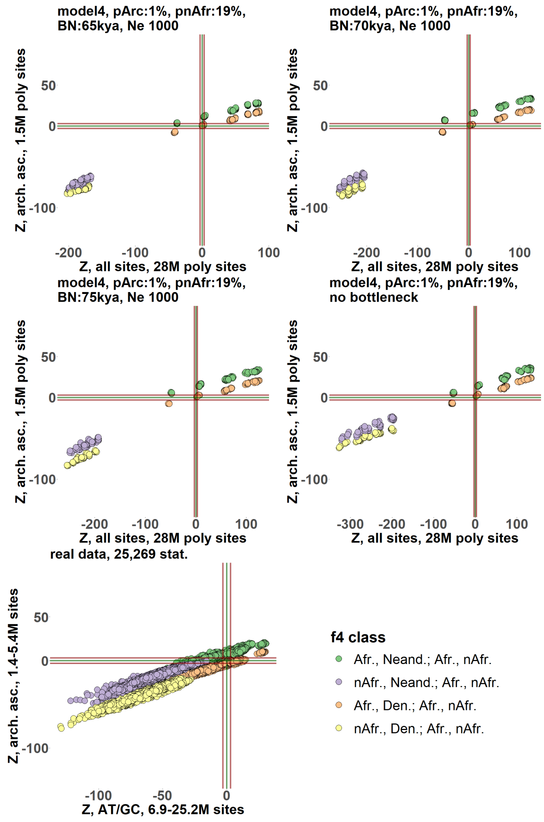
model4, pArc:1%, pnAfr:19%, BN:65kya, Ne 1000
model4, pArc:1%, pnAfr:19%, BN:70kya, Ne 1000
model4, pArc:1%, pnAfr:19%, BN:75kya, Ne 1000
model4, pArc:1%, pnAfr:19%, no bottleneck
Z, arch. asc., 1.5M poly sites
Z, all sites, 28M poly sites
real data, 25,269 stat.
Z, arch. asc., 1.4-5.4M sites
Z, AT/GC, 6.9-25.2M sites
f4 class
Afr., Neand.; Afr., nAfr.
nAfr., Neand.; Afr., nAfr.
Afr., Den.; Afr., nAfr.
nAfr., Den.; Afr., nAfr.

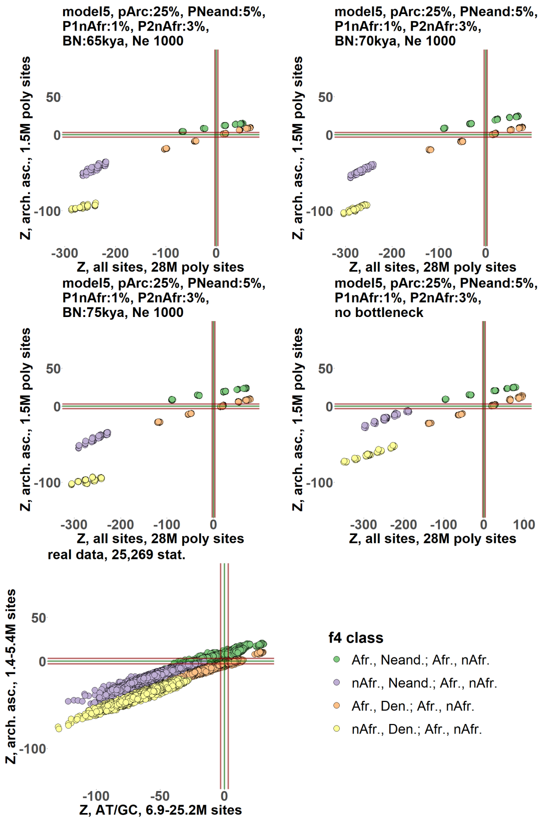
model5, pArc:25%, PNeand:5%, P1nAfr:1%, P2nAfr:3%, BN:65kya, Ne 1000
model5, pArc:25%, PNeand:5%, P1nAfr:1%, P2nAfr:3%, BN:70kya, Ne 1000
model5, pArc:25%, PNeand:5%, P1nAfr:1%, P2nAfr:3%, BN:75kya, Ne 1000
model5, pArc:25%, PNeand:5%, P1nAfr:1%, P2nAfr:3%, no bottleneck
Z, arch. asc., 1.5M poly sites
Z, all sites, 28M poly sites
real data, 25,269 stat.
Z, arch. asc., 1.4-5.4M sites
Z, AT/GC, 6.9-25.2M sites
f4 class
Afr., Neand.; Afr., nAfr.
nAfr., Neand.; Afr., nAfr.
Afr., Den.; Afr., nAfr.
nAfr., Den.; Afr., nAfr.

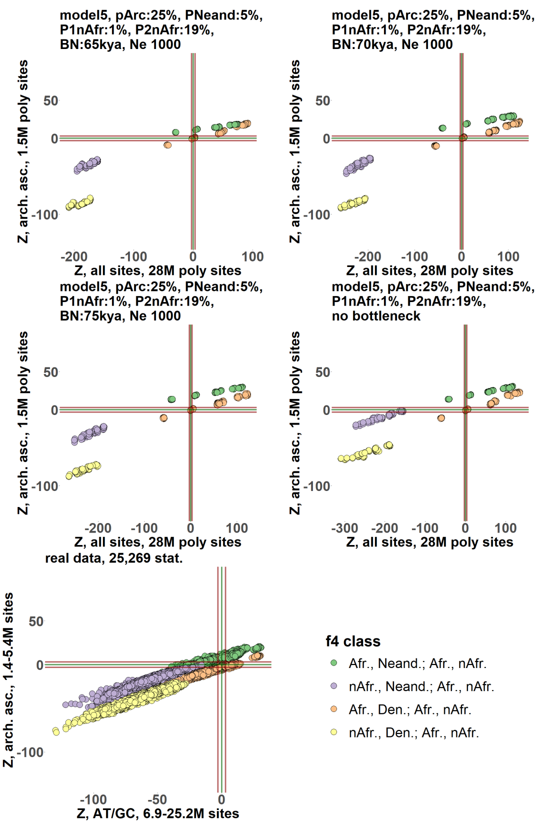

**Suppl. Fig. 17.** The effects of ascertaining SNPs polymorphic in archaic humans on real (**a**) and simulated data (**b, c**). We focused on four $f_4$-statistic classes that are most strongly affected by any type of ascertainment on real data: $f_4$(African X, Neanderthal or Denisovan; African Y, non-African) and $f_4$(non-African X, Neanderthal or Denisovan; African, non-African Y). On real data, statistics from these classes were sampled randomly and were calculated on AT/GC sites and on archaic-ascertained sites (transitions and transversions), using all sites without missing data on the level of each quadruplet (i.e., using the "*allsnps=TRUE*" or "*useallsnps: YES*" setting). Papuans and Australians were excluded from the pool of AMH groups due to their Denisovan ancestry, which was not simulated; and Africans with substantial non-African ancestry (Suppl. Table 1) were also removed to make the distinction between various classes of statistics clearer. Archaic ascertainment was performed either on a group composed of the Altai Neanderthal and Denisovan (panel **a**, left), or Vindija Neanderthal and Denisovan (panel **a**, right). A slightly different protocol was used for archaic ascertainment in other parts of this paper since it was performed on a group composed of both Neanderthal individuals and the Denisovan.

Graphs illustrating five classes of simulated demographic histories are shown in panel **b** and scatterplots illustrating the effects of ascertaining SNPs polymorphic in the group composed of one "Neanderthal" and one "Denisovan" individual on genetic data simulated according to those histories are shown in panel **c**. The same "Neanderthal" and "Denisovan" individuals were used for ascertainment and for calculating $f_4$-statistics. On the graphs, the following abbreviations are used: Afr., Africans; nAfr., non-Africans; Den., Denisovan; Neand., Neanderthal. Alternative positions of the out-of-Africa bottleneck simulated at 65, 70, or 75 kya (generation time = 25 years) are marked with red dots. Gene flows from ghost unsampled lineages are shown in green, and those from sampled lineages are shown in blue. Divergence or split times are shown in kya, and effective population sizes are not shown for clarity. Results for 68 simulated histories are presented, and by "history" we assume a combination of admixture graph topology, admixture proportions, effective population sizes, population divergence and bottleneck times. The following key simulation parameters are shown in plot titles for Models 1 through 4: model name, proportion of "super-archaic" or Neanderthal-related admixture in ancestral AMH ("Parc"), proportion of Neanderthal or ghost AMH admixture in non-Africans ("PnAfr"), out-of-Africa bottleneck date in kya ("BN"), diploid effective population size during the out-of-Africa bottleneck ("Ne"). The following additional simulation parameters are shown in plot titles for Model 5: proportion of AMH admixture in the Neanderthal lineage ("Pneand"), proportion of Neanderthal admixture in non-Africans ("P1nAfr"), proportion of "ghost AMH" admixture in non-Africans ("P2nAfr").

**Suppl. Fig. 18.**

**a**

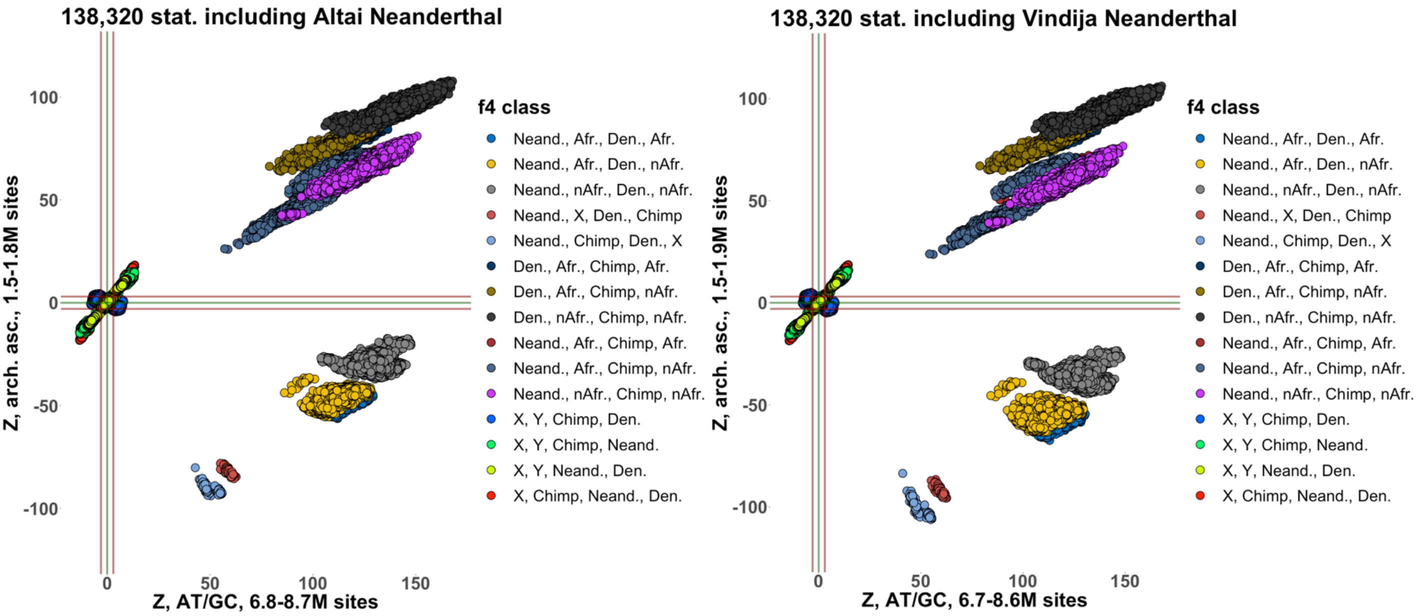

**b**

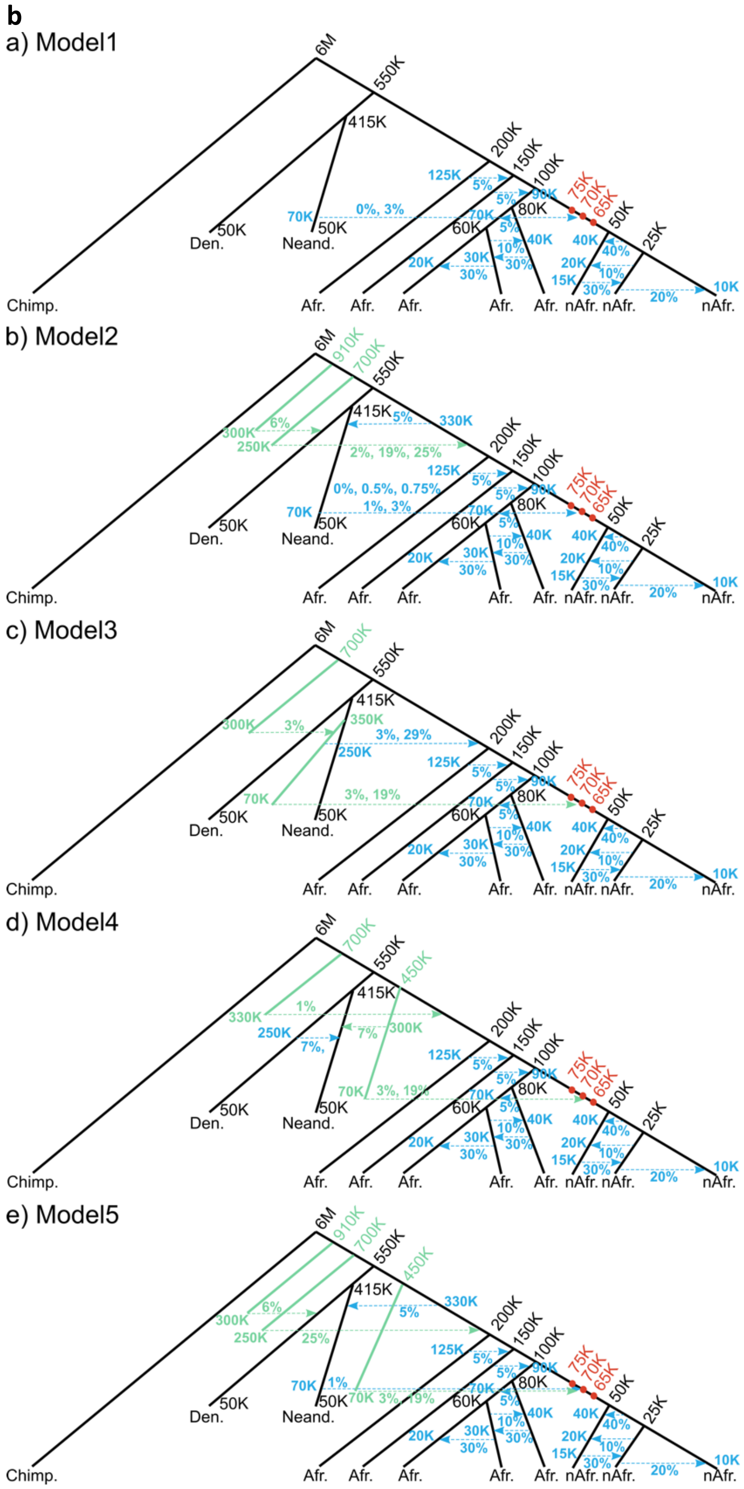

a) Model1
6M
550K
415K
200K
150K
100K
75K
70K
65K
50K
25K
80K
60K
125K
70K
0%, 3%
50K
Den.
Neand.
Chimp.
Afr.
nAfr.
10K
20%
b) Model2
910K
700K
300K
250K
6%
330K
5%
2%, 19%, 25%
0%, 0.5%, 0.75%
1%, 3%
c) Model3
350K
3%
3%, 29%
3%, 19%
d) Model4
450K
330K
1%
7%
300K
3%, 19%
e) Model5
25%
70K 3%, 19%

**C**

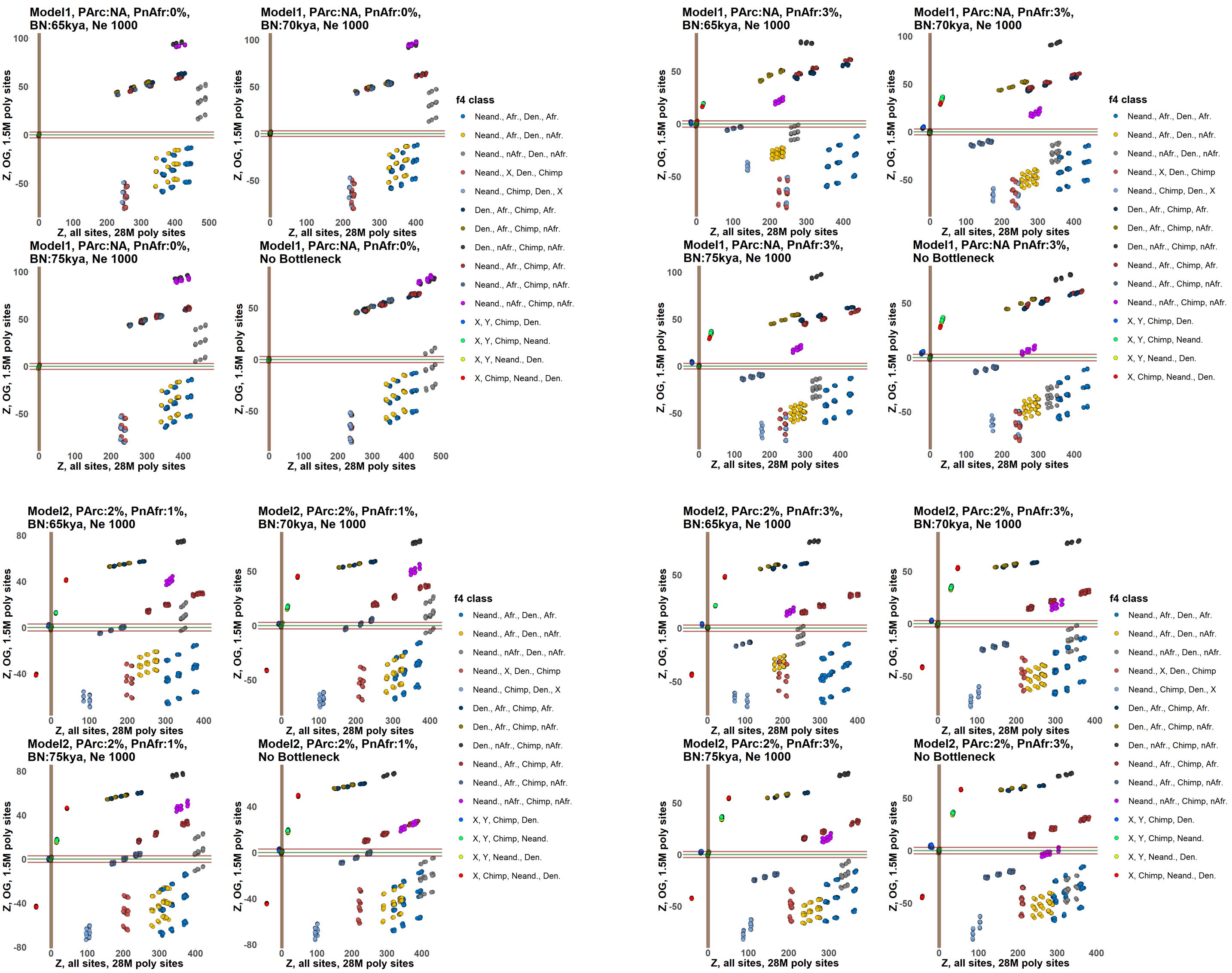

Model1, PArc:NA, PnAfr:0%, BN:65kya, Ne 1000
Model1, PArc:NA, PnAfr:0%, BN:70kya, Ne 1000
Model1, PArc:NA, PnAfr:0%, BN:75kya, Ne 1000
Model1, PArc:NA, PnAfr:0%, No Bottleneck
Model1, PArc:NA, PnAfr:3%, BN:65kya, Ne 1000
Model1, PArc:NA, PnAfr:3%, BN:70kya, Ne 1000
Model1, PArc:NA, PnAfr:3%, BN:75kya, Ne 1000
Model1, PArc:NA PnAfr:3%, No Bottleneck
Model2, PArc:2%, PnAfr:1%, BN:65kya, Ne 1000
Model2, PArc:2%, PnAfr:1%, BN:70kya, Ne 1000
Model2, PArc:2%, PnAfr:1%, BN:75kya, Ne 1000
Model2, PArc:2%, PnAfr:1%, No Bottleneck
Model2, PArc:2%, PnAfr:3%, BN:65kya, Ne 1000
Model2, PArc:2%, PnAfr:3%, BN:70kya, Ne 1000
Model2, PArc:2%, PnAfr:3%, BN:75kya, Ne 1000
Model2, PArc:2%, PnAfr:3%, No Bottleneck
Z, OG, 1.5M poly sites
Z, all sites, 28M poly sites
f4 class
Neand., Afr., Den., Afr.
Neand., Afr., Den., nAfr.
Neand., nAfr., Den., nAfr.
Neand., X, Den., Chimp
Neand., Chimp, Den., X
Den., Afr., Chimp, Afr.
Den., Afr., Chimp, nAfr.
Den., nAfr., Chimp, nAfr.
Neand., Afr., Chimp, Afr.
Neand., Afr., Chimp, nAfr.
Neand., nAfr., Chimp, nAfr.
X, Y, Chimp, Den.
X, Y, Chimp, Neand.
X, Y, Neand., Den.
X, Chimp, Neand., Den.

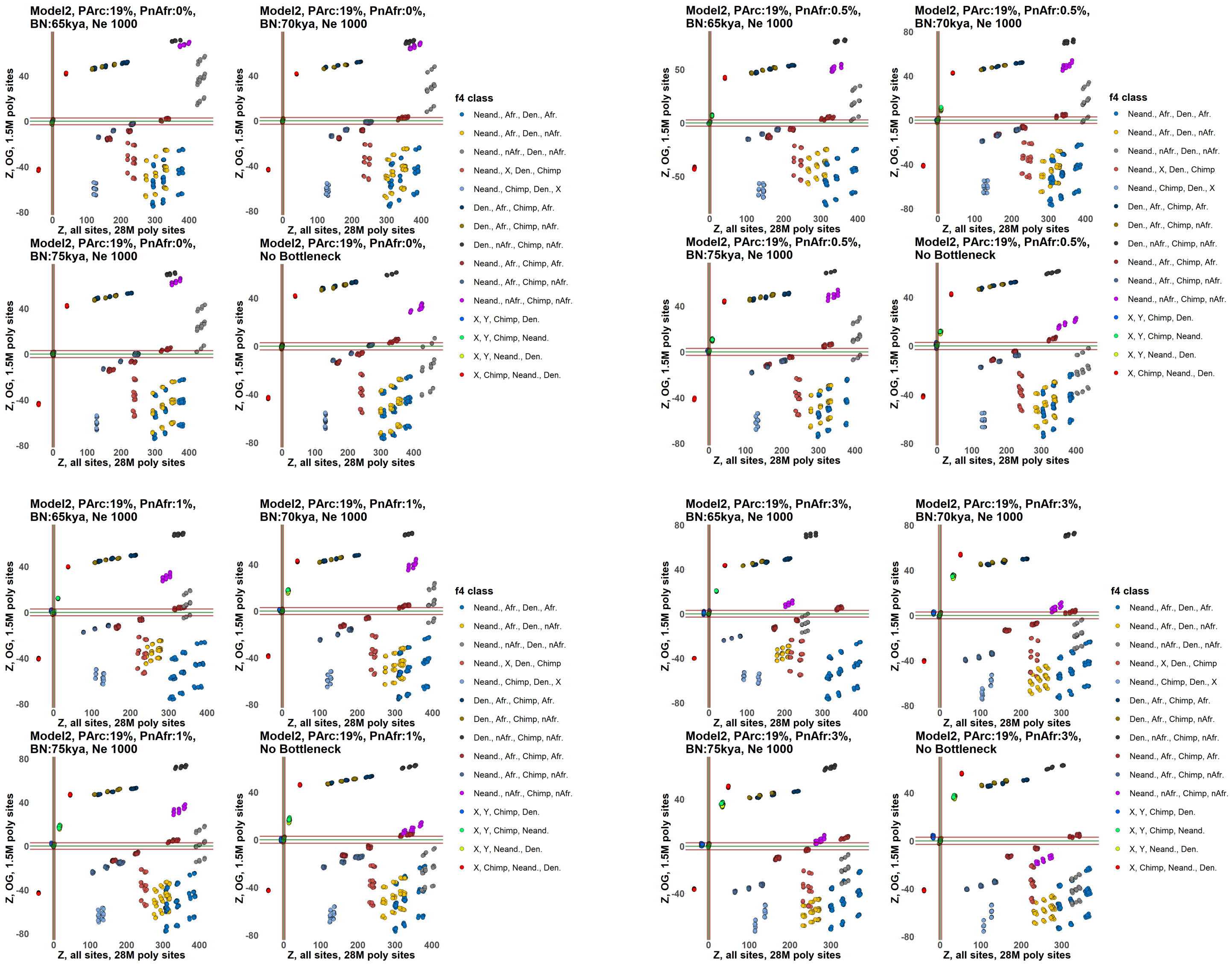
Model2, PArc:19%, PnAfr:0%, BN:65kya, Ne 1000
Model2, PArc:19%, PnAfr:0%, BN:70kya, Ne 1000
Model2, PArc:19%, PnAfr:0%, BN:75kya, Ne 1000
Model2, PArc:19%, PnAfr:0%, No Bottleneck
Model2, PArc:19%, PnAfr:0.5%, BN:65kya, Ne 1000
Model2, PArc:19%, PnAfr:0.5%, BN:70kya, Ne 1000
Model2, PArc:19%, PnAfr:0.5%, BN:75kya, Ne 1000
Model2, PArc:19%, PnAfr:0.5%, No Bottleneck
Model2, PArc:19%, PnAfr:1%, BN:65kya, Ne 1000
Model2, PArc:19%, PnAfr:1%, BN:70kya, Ne 1000
Model2, PArc:19%, PnAfr:1%, BN:75kya, Ne 1000
Model2, PArc:19%, PnAfr:1%, No Bottleneck
Model2, PArc:19%, PnAfr:3%, BN:65kya, Ne 1000
Model2, PArc:19%, PnAfr:3%, BN:70kya, Ne 1000
Model2, PArc:19%, PnAfr:3%, BN:75kya, Ne 1000
Model2, PArc:19%, PnAfr:3%, No Bottleneck
Z, OG, 1.5M poly sites
Z, all sites, 28M poly sites
f4 class
Neand., Afr., Den., Afr.
Neand., Afr., Den., nAfr.
Neand., nAfr., Den., nAfr.
Neand., X, Den., Chimp
Neand., Chimp, Den., X
Den., Afr., Chimp, Afr.
Den., Afr., Chimp, nAfr.
Den., nAfr., Chimp, nAfr.
Neand., Afr., Chimp, Afr.
Neand., Afr., Chimp, nAfr.
Neand., nAfr., Chimp, nAfr.
X, Y, Chimp, Den.
X, Y, Chimp, Neand.
X, Y, Neand., Den.
X, Chimp, Neand., Den.

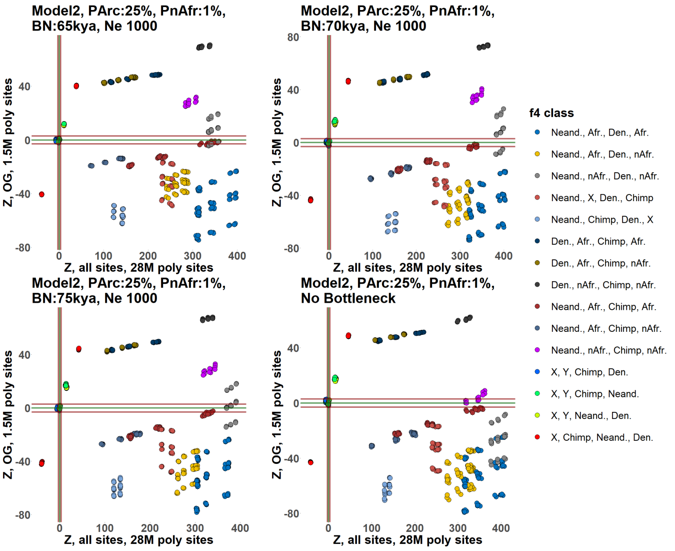
Model2, PArc:25%, PnAfr:1%, BN:65kya, Ne 1000
Model2, PArc:25%, PnAfr:1%, BN:70kya, Ne 1000
Model2, PArc:25%, PnAfr:1%, BN:75kya, Ne 1000
Model2, PArc:25%, PnAfr:1%, No Bottleneck
Z, OG, 1.5M poly sites
Z, all sites, 28M poly sites
f4 class
Neand., Afr., Den., Afr.
Neand., Afr., Den., nAfr.
Neand., nAfr., Den., nAfr.
Neand., X, Den., Chimp
Neand., Chimp, Den., X
Den., Afr., Chimp, Afr.
Den., Afr., Chimp, nAfr.
Den., nAfr., Chimp, nAfr.
Neand., Afr., Chimp, Afr.
Neand., Afr., Chimp, nAfr.
Neand., nAfr., Chimp, nAfr.
X, Y, Chimp, Den.
X, Y, Chimp, Neand.
X, Y, Neand., Den.
X, Chimp, Neand., Den.

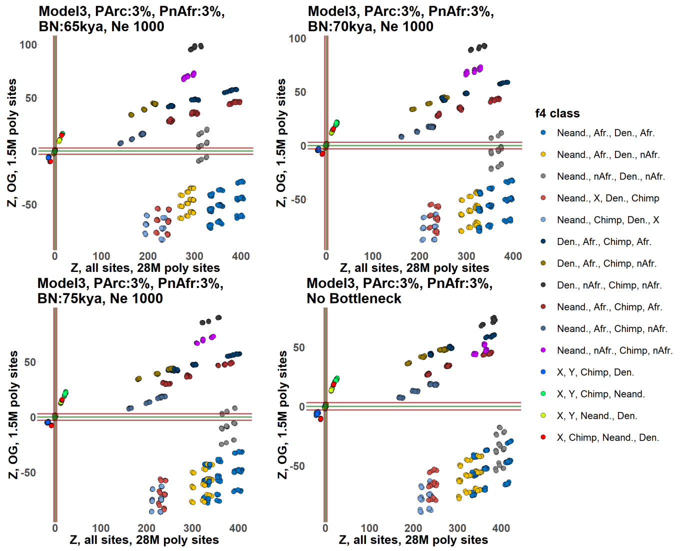
Model3, PArc:3%, PnAfr:3%, BN:65kya, Ne 1000
Model3, PArc:3%, PnAfr:3%, BN:70kya, Ne 1000
Model3, PArc:3%, PnAfr:3%, BN:75kya, Ne 1000
Model3, PArc:3%, PnAfr:3%, No Bottleneck
Z, OG, 1.5M poly sites
Z, all sites, 28M poly sites
f4 class
Neand., Afr., Den., Afr.
Neand., Afr., Den., nAfr.
Neand., nAfr., Den., nAfr.
Neand., X, Den., Chimp
Neand., Chimp, Den., X
Den., Afr., Chimp, Afr.
Den., Afr., Chimp, nAfr.
Den., nAfr., Chimp, nAfr.
Neand., Afr., Chimp, Afr.
Neand., Afr., Chimp, nAfr.
Neand., nAfr., Chimp, nAfr.
X, Y, Chimp, Den.
X, Y, Chimp, Neand.
X, Y, Neand., Den.
X, Chimp, Neand., Den.

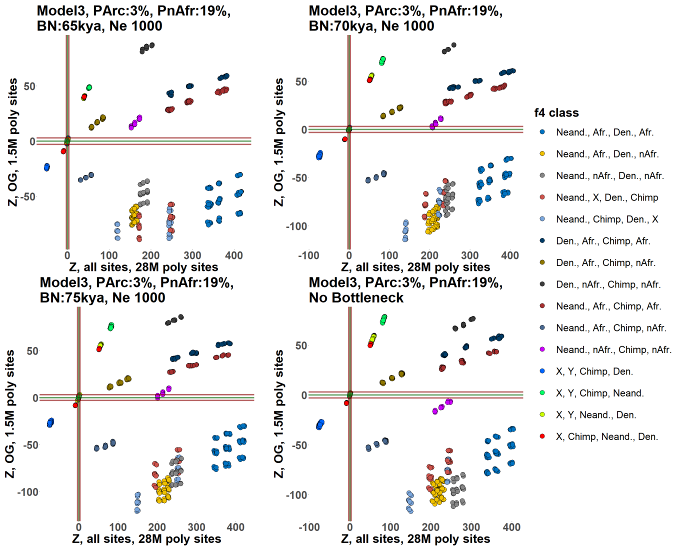
Model3, PArc:3%, PnAfr:19%, BN:65kya, Ne 1000
Model3, PArc:3%, PnAfr:19%, BN:70kya, Ne 1000
Model3, PArc:3%, PnAfr:19%, BN:75kya, Ne 1000
Model3, PArc:3%, PnAfr:19%, No Bottleneck
Z, OG, 1.5M poly sites
Z, all sites, 28M poly sites
f4 class
Neand., Afr., Den., Afr.
Neand., Afr., Den., nAfr.
Neand., nAfr., Den., nAfr.
Neand., X, Den., Chimp
Neand., Chimp, Den., X
Den., Afr., Chimp, Afr.
Den., Afr., Chimp, nAfr.
Den., nAfr., Chimp, nAfr.
Neand., Afr., Chimp, Afr.
Neand., Afr., Chimp, nAfr.
Neand., nAfr., Chimp, nAfr.
X, Y, Chimp, Den.
X, Y, Chimp, Neand.
X, Y, Neand., Den.
X, Chimp, Neand., Den.

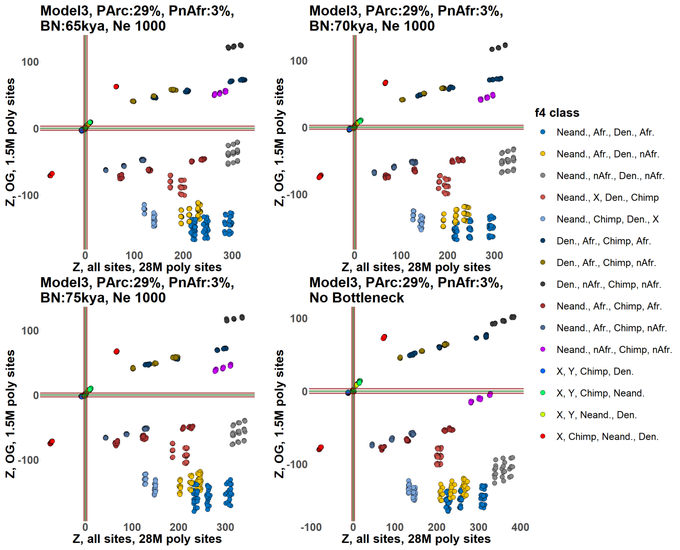
Model3, PArc:29%, PnAfr:3%, BN:65kya, Ne 1000
Model3, PArc:29%, PnAfr:3%, BN:70kya, Ne 1000
Model3, PArc:29%, PnAfr:3%, BN:75kya, Ne 1000
Model3, PArc:29%, PnAfr:3%, No Bottleneck
Z, OG, 1.5M poly sites
Z, all sites, 28M poly sites
f4 class
Neand., Afr., Den., Afr.
Neand., Afr., Den., nAfr.
Neand., nAfr., Den., nAfr.
Neand., X, Den., Chimp
Neand., Chimp, Den., X
Den., Afr., Chimp, Afr.
Den., Afr., Chimp, nAfr.
Den., nAfr., Chimp, nAfr.
Neand., Afr., Chimp, Afr.
Neand., Afr., Chimp, nAfr.
Neand., nAfr., Chimp, nAfr.
X, Y, Chimp, Den.
X, Y, Chimp, Neand.
X, Y, Neand., Den.
X, Chimp, Neand., Den.

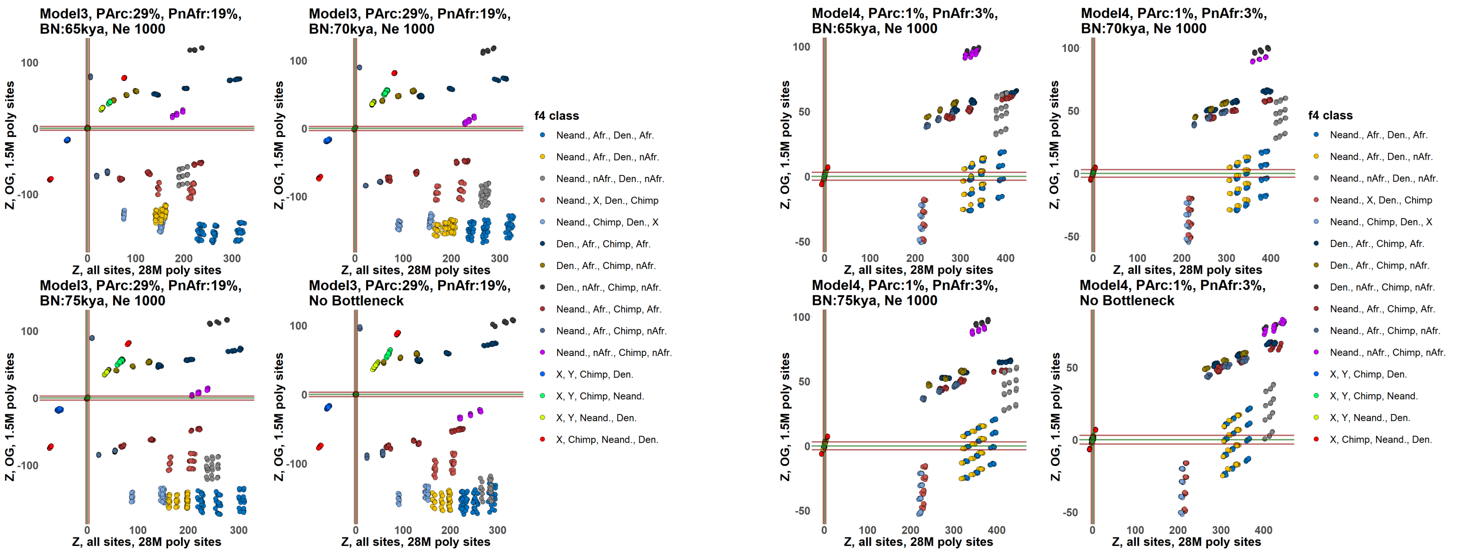

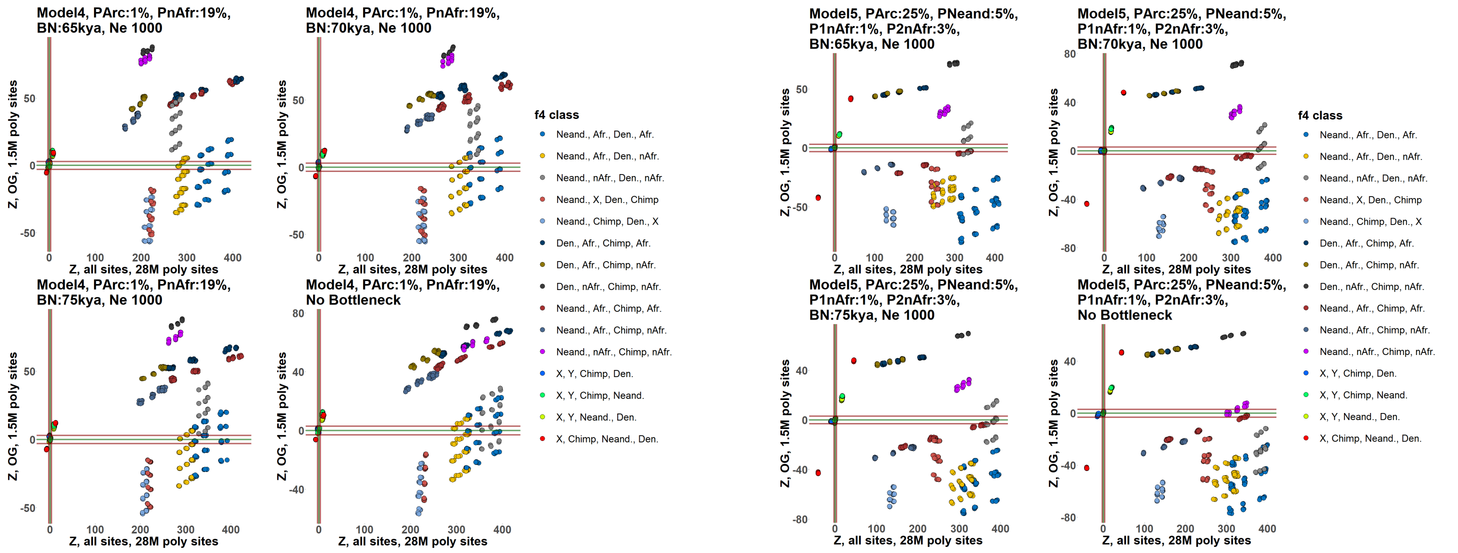

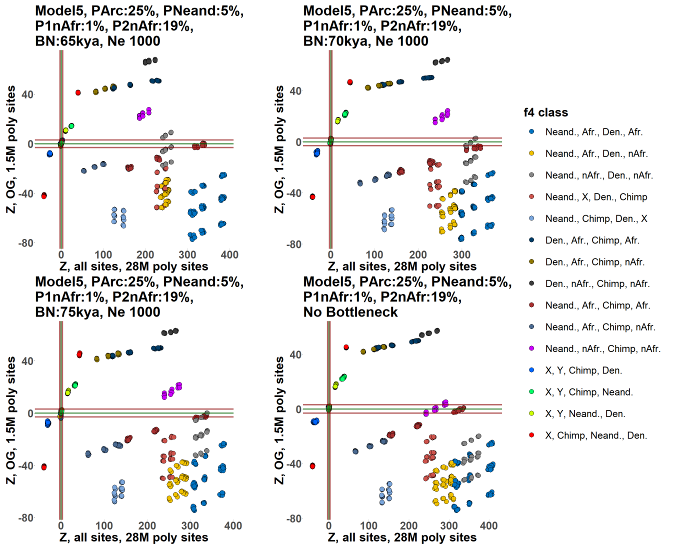

**Suppl. Fig. 18.** The effects of ascertaining SNPs polymorphic in archaic humans on real (**a**) and simulated data (**b, c**). We focused on 15 $f_4$-statistic classes that are most strongly affected by archaic ascertainment on simulated data (see a list of classes in the legend for panel **a**). On real data, statistics from these classes were sampled exhaustively and were calculated on AT/GC sites and on archaic-ascertained sites (transitions and transversions), using all sites without missing data on the level of each quadruplet (i.e., using the "*allsnps=TRUE*" or "*useallsnps: YES*" setting). Papuans and Australians were excluded from the pool of AMH groups due to their Denisovan ancestry, which was not simulated; and Africans with substantial non-African ancestry (Suppl. Table 1) were also removed to make the distinction between various classes of statistics clearer. Archaic ascertainment was performed either on a group composed of the Altai Neanderthal and Denisovan (panel **a**, left), or Vindija Neanderthal and Denisovan (panel **a**, right). A slightly different protocol was used for archaic ascertainment in other parts of this paper since it was performed on a group composed of both Neanderthals and the Denisovan.

Graphs illustrating five classes of simulated demographic histories are shown in panel **b** and scatterplots illustrating the effects of ascertaining SNPs polymorphic in the group composed of one "Neanderthal" and one "Denisovan" individual on genetic data simulated according to those histories are shown in panel **c**. The same "Neanderthal" and "Denisovan" individuals were used for ascertainment and for calculating $f_4$-statistics. On the graphs, the following abbreviations are used: Afr., Africans; nAfr., non-Africans; Den., Denisovan; Neand., Neanderthal. Alternative positions of the out-of-Africa bottleneck simulated at 65, 70, or 75 kya (generation time = 25 years) are marked with red dots. Gene flows from ghost unsampled lineages are shown in green, and those from sampled lineages are shown in blue. Divergence or split times are shown in kya, and effective population sizes are not shown for clarity. We focused on 15 $f_4$-statistic classes that are most affected by archaic ascertainment on simulated data (see a list of classes in the legends for each panel). Results for 68 simulated histories are presented, and by "history" we assume a combination of admixture graph topology, admixture proportions, effective population sizes, population divergence and bottleneck times. The following key simulation parameters are shown in plot titles for Models 1 through 4: model name, proportion of "super-archaic" or Neanderthal-related admixture in ancestral AMH ("Parc"), proportion of Neanderthal or ghost AMH admixture in non-Africans ("PnAfr"), out-of-Africa bottleneck date in kya ("BN"), diploid effective population size during the out-of-Africa bottleneck ("Ne"). The following additional simulation parameters are shown in plot titles for Model 5: proportion of AMH admixture in the Neanderthal lineage ("Pneand"), proportion of Neanderthal admixture in non-Africans ("P1nAfr"), proportion of "ghost AMH" admixture in non-Africans ("P2nAfr").

### percentage of models accepted on asc. but rejected on all sites

| ascertainment type | further details on the ascertainment | min. size of the SNP panel* | max. size of the SNP panel* | *arch 2, afr 3*<br>Denisovan, Altai, Yoruba, Dinka, Bulala | *arch 1, afr 4*<br>Denisovan, Khomani San, Mbuti, Dinka, Mursi | *arch 1, afr 3, nafr 1*<br>Altai, Ju hoan North, Biaka, Yoruba, Agaw | *arch 1, afr 2, nafr 2*<br>Altai, Ju hoan North, Luhya, Palestinian, Spanish | *afr 5*<br>Mbuti, Baka, Laka, Fulani, Bantu Tswana | *afr 4, nafr 1*<br>Khomani San, Bakola, Igbo, Mursi, Aari | *afr 4, nafr 1*<br>Bedzan, Cameroon SMA, Esan, Mozabite, Masai | *afr 3, nafr 2*<br>Mbuti, Biaka, Ngumba, LBK, Iranian | *afr 1, nafr 4*<br>Luo, Bedouin B, Jordanian, Abkhasian, Sardinian | *nafr 5*<br>Australian, Quechua, Mayan, Lezgin, French | *nafr 5*<br>Papuan, Chipewyan, Eskimo Naukan, Finnish, Sardinian | *nafr 5*<br>Karitiana, Cree, Eskimo Sireniki, Hungarian, Icelandic | number of biased pop. sets |
|---|---|---|---|---|---|---|---|---|---|---|---|---|---|---|---|---|
| A<>T and G<>C mutations | | 805,042 | 1,757,840 | 10.679% | 0.015% | 0.000% | 0.000% | 0.003% | 0.000% | 0.018% | 2.144% | 7.051% | 0.000% | 1.469% | 0.000% | 0 |
| 1240K panel | | 501,429 | 663,239 | 0.055% | 0.018% | **1.023%** | 5.525% | 0.003% | 0.000% | 0.018% | 6.016% | 8.643% | 0.000% | 1.191% | 0.000% | 1 |
| components of the 1240K panel | Illumina610-Quad sites | 256,277 | 304,292 | 0.052% | 0.024% | **1.023%** | 5.525% | 0.000% | 0.000% | 0.067% | 17.716% | 0.000% | 0.000% | 1.460% | 0.000% | 1 |
| | sites exclusive to Illumina610-Quad | 183,680 | 216,478 | 10.679% | 0.024% | **1.023%** | 5.528% | 0.003% | 0.000% | 7.140% | 17.716% | 0.916% | 0.000% | 0.443% | 0.000% | 1 |
| | sites included in both Illumina610-Quad and HumanOrigins | 72,597 | 87,814 | 0.049% | 5.341% | **1.047%** | 5.528% | 11.684% | 0.000% | **7.155%** | 17.716% | **9.714%** | 0.000% | **6.792%** | 0.000% | 4 |
| | HumanOrigins sites | 244,922 | 354,460 | 10.683% | 5.341% | **1.020%** | 11.681% | 0.003% | 0.000% | 0.067% | 0.003% | 8.664% | 0.000% | **1.542%** | 0.000% | 2 |
| | sites exclusive to HumanOrigins | 171,249 | 266,646 | 10.683% | 5.567% | **1.041%** | 11.681% | 0.003% | 0.000% | 6.972% | 0.250% | 8.664% | 0.000% | 1.096% | 0.000% | 1 |
| | 1240K, other sites | 67,096 | 92,301 | 10.683% | 6.386% | **1.023%** | 11.681% | 1.115% | **1.063%** | 0.360% | 6.031% | 8.664% | 0.000% | **1.542%** | 0.000% | 3 |
| the largest panels included in the HumanOrigins array | panel 13 based on a San individual and Denisovan | 67,557 | 89,655 | 10.683% | **21.099%** | **8.786%** | 11.681% | 1.081% | **2.071%** | **17.777%** | 17.716% | 8.664% | 0.000% | **6.789%** | 0.000% | 5 |
| | panel 4 based on a San individual | 52,862 | 94,493 | 10.679% | 17.022% | **1.051%** | 11.681% | 1.087% | **1.115%** | **17.777%** | 0.296% | 8.664% | 0.000% | **6.792%** | 0.000% | 4 |
| | panel 5 based on a Yoruba individual | 44,674 | 73,180 | 0.031% | 17.022% | **1.029%** | 11.681% | 11.684% | **2.159%** | **7.219%** | 17.716% | **9.714%** | 0.000% | **2.507%** | **0.092%** | 6 |
| | panels 4 and 5 | 46,701 | 157,126 | 0.064% | 17.022% | **1.038%** | 11.681% | 0.003% | 1.023% | 6.453% | 0.003% | 8.664% | 0.000% | 1.460% | 0.000% | 1 |
| other enrichment panels | 1000K: transversions in 2 Yoruba ind. and in Altai Neand. | 364,079 | 590,775 | 10.683% | 0.098% | 0.000% | 11.681% | 0.003% | 0.000% | 0.092% | 0.296% | 8.664% | 0.000% | **2.495%** | 0.000% | 1 |
| | 2200K = 1000K + 1240K | 814,915 | 1,190,758 | 0.049% | 0.024% | 0.000% | 11.681% | 0.003% | 0.000% | 0.018% | 0.000% | 8.646% | 0.000% | 1.454% | 0.000% | 0 |
| archaic ascertainment | transitions and transversions | 525,014 | 1,555,781 | 2.019% | 17.022% | **0.092%** | 0.000% | 11.684% | **1.063%** | **7.192%** | 7.018% | **11.446%** | 0.000% | **2.562%** | **2.324%** | 6 |
| | transversions | 165,249 | 484,675 | 2.046% | 17.022% | **0.092%** | 0.000% | 11.684% | **2.086%** | **7.192%** | 7.039% | **24.327%** | 0.000% | **2.642%** | **2.361%** | 6 |
| MAF ascertainment | global, >5% MAF | 2,129,201 | 2,511,335 | 10.683% | 0.073% | **1.011%** | 11.681% | 0.003% | 0.000% | 0.018% | 0.293% | 0.000% | 0.000% | 1.011% | 0.000% | 1 |
| | >5% MAF in Africans unadmixed with non-Africans | 2,045,769 | 3,231,875 | 0.021% | 0.024% | 0.000% | 11.681% | 0.003% | 0.000% | 0.000% | 0.006% | 7.051% | 0.000% | 1.448% | 0.000% | 0 |
| | >5% MAF in all Africans | 2,109,808 | 3,120,326 | 0.031% | 0.061% | 0.000% | 11.681% | 0.003% | 0.000% | 0.018% | 0.006% | 7.039% | 0.000% | 1.017% | 0.000% | 0 |
| | >5% MAF in Native Americans | 1,513,207 | 1,764,715 | 0.055% | 5.341% | **1.023%** | 6.016% | 0.003% | 0.000% | 0.000% | 6.315% | 1.139% | 0.000% | 1.469% | 0.000% | 1 |
| | >5% MAF in Central Asians and Siberians | 1,843,262 | 2,150,675 | 0.055% | 5.341% | **1.023%** | 11.681% | 0.003% | 0.000% | 0.000% | 0.290% | 0.950% | 0.000% | 0.434% | 0.000% | 1 |
| | >5% MAF in East Asians | 1,723,831 | 2,020,860 | 10.683% | 5.341% | **1.023%** | 5.378% | 0.003% | 0.000% | 0.000% | 0.293% | 7.051% | 0.000% | 1.454% | 0.000% | 1 |
| | >5% MAF in Europeans | 1,885,336 | 2,192,571 | 10.679% | 5.341% | **1.008%** | 11.681% | 0.003% | 0.000% | 0.018% | 0.263% | 2.055% | 0.000% | 1.454% | 0.000% | 1 |
| | >5% MAF in Middle Eastern groups | 2,018,884 | 2,306,319 | 10.683% | 5.341% | 0.000% | 11.681% | 0.003% | 0.000% | 0.018% | 0.293% | 1.139% | 0.000% | 1.444% | 0.000% | 0 |
| | >5% MAF in Papuans and Aboriginal Australians | 1,515,022 | 1,791,390 | 0.021% | 5.341% | **1.047%** | 5.528% | 0.003% | 0.000% | 1.020% | 6.328% | 1.051% | 0.000% | 1.460% | 0.000% | 1 |
| | >5% MAF in South Asians | 1,908,459 | 2,235,024 | 10.679% | 5.341% | **1.023%** | 11.681% | 0.003% | 0.000% | 0.018% | 0.281% | 0.000% | 0.000% | 1.435% | 0.000% | 1 |
| AT/GC mutation types + MAF ascertainment | global, >>5% MAF | 323,296 | 378,287 | 10.683% | 0.098% | **1.023%** | 11.681% | 0.003% | 0.000% | 0.092% | 17.716% | 7.051% | 0.000% | 1.475% | 0.000% | 1 |
| | >5% MAF in Africans unadmixed with non-Africans | 309,172 | 486,906 | 10.683% | 0.018% | 0.000% | 11.681% | 0.003% | 0.000% | 0.018% | 7.091% | 7.051% | 0.000% | 1.466% | 0.000% | 0 |
| | >5% MAF in all Africans | 319,053 | 470,070 | 10.683% | 0.070% | 0.000% | 11.681% | 0.003% | 0.000% | 0.018% | 17.716% | 7.051% | 0.000% | 1.469% | 0.000% | 0 |
| | >5% MAF in Native Americans | 229,939 | 266,113 | 10.679% | 5.341% | **1.047%** | 7.039% | 11.684% | 0.000% | 6.963% | 7.317% | 8.078% | 0.000% | 1.475% | 0.000% | 1 |
| | >5% MAF in Central Asians and Siberians | 280,103 | 324,245 | 10.683% | 5.341% | **1.023%** | 11.681% | 11.684% | 0.000% | 1.051% | 17.716% | 1.044% | 0.000% | 1.469% | 0.000% | 1 |
| | >5% MAF in East Asians | 261,857 | 304,567 | 10.683% | 0.098% | **1.023%** | 11.681% | 11.684% | 0.000% | 1.047% | 7.277% | 8.026% | 0.000% | 1.506% | 0.000% | 1 |
| | >5% MAF in Europeans | 285,723 | 330,244 | 10.683% | 5.341% | **1.020%** | 11.681% | 1.023% | 0.000% | **7.161%** | 17.713% | 8.099% | 0.000% | 1.472% | 0.000% | 2 |
| | >5% MAF in Middle Eastern groups | 306,450 | 347,536 | 10.683% | 5.341% | **1.014%** | 11.681% | 1.002% | 0.000% | **7.201%** | 17.703% | 7.036% | 0.000% | 1.472% | 0.000% | 2 |
| | >5% MAF in Papuans and Aboriginal Australians | 230,124 | 272,093 | 0.027% | 5.341% | **1.047%** | 5.509% | 11.684% | 1.008% | 6.337% | 7.235% | 8.539% | 0.000% | 1.472% | 0.000% | 1 |
| | >5% MAF in South Asians | 289,739 | 336,996 | 10.683% | 5.341% | **1.044%** | 11.681% | 1.057% | 0.000% | **17.761%** | 17.713% | 8.078% | 0.000% | 1.469% | 0.000% | 2 |
| * the SNP counts correspond to sites polymorphic in larger collections of groups from which the analyzed population quintuplets were taken, see Suppl. Table X. | | number of biased asc. => | | 0 | 1 | 28 | 0 | 0 | 6 | 9 | 0 | 4 | 0 | 9 | 3 | |

**Suppl. Table 3.** Performance of ascertainment schemes explored across 12 population quintuplets and assessed as the fraction of all admixture graph topologies that are accepted under ascertainment (WR < 3 SE) but rejected on all sites (WR > 3 SE). We also applied the binary classifier to determine if the ascertainment produces unbiased or biased results (the latter highlighted in bold and underlined text). The numbers of population quintuplets or ascertainment schemes affected by bias (according to this classifier) are shown in the rightmost column and in the bottom row, respectively. The composition of the population sets is shown above the table in an abbreviated way: *arch*, archaic humans, followed by the number of archaic groups; *afr*, Africans; *nafr*, non-Africans or Africans with substantial non-African admixture (Fan et al. 2019).

## WR, R2

| ascertainment type | further details on the ascertainment | min. size of the SNP panel* | max. size of the SNP panel* | *arch 2, afr 3* | | *arch 1, afr 4* | | *arch 1, afr 3, nafr 1* | | *arch 1, afr 2, nafr 2* | | *afr 5* | | *afr 4, nafr 1* | | *afr 4, nafr 1* | | *afr 3, nafr 2* | | *afr 1, nafr 4* | | *nafr 5* | | *nafr 5* | | *nafr 5* | | med. | number of biased pop. sets |
|---|---|---|---|---|---|---|---|---|---|---|---|---|---|---|---|---|---|---|---|---|---|---|---|---|---|---|---|---|---|
| | | | | Denisovan, Altai, Yoruba, Dinka, Bulala | | Denisovan, Khomani San, Mbuti, Dinka, Mursi | | Altai, Ju hoan North, Biaka, Yoruba, Agaw | | Altai, Ju hoan North, Luhya, Palestinian, Spanish | | Mbuti, Baka, Laka, Fulani, Bantu Tswana | | Khomani San, Bakola, Igbo, Mursi, Aari | | Bedzan, Cameroon SMA, Esan, Mozabite, Masai | | Mbuti, Biaka, Ngumba, LBK, Iranian | | Luo, Bedouin B, Jordanian, Abkhasian, Sardinian | | Australian, Quechua, Mayan, Lezgin, French | | Papuan, Chipewyan, Eskimo Naukan, Finnish, Sardinian | | Karitiana, Cree, Eskimo Sireniki, Hungarian, Icelandic | | | |
| | | | | 5000 | all | 5000 | all | 5000 | all | 5000 | all | 5000 | all | 5000 | all | 5000 | all | 5000 | all | 5000 | all | 5000 | all | 5000 | all | 5000 | all | | |
| A<>T and G<>C mutations | | 805,042 | 1,757,840 | 0.953 | 1.000 | 0.990 | 0.999 | 0.996 | 1.000 | 0.953 | 0.998 | 0.963 | 0.998 | 0.998 | 0.997 | 0.957 | 0.998 | 0.873 | 1.000 | 0.918 | 0.987 | 0.812 | 1.000 | 0.995 | 1.000 | 0.997 | 1.000 | 0.997 | 0 |
| 1240K panel | | 501,429 | 663,239 | 0.858 | 1.000 | 0.652 | 0.995 | **0.093** | **0.971** | **0.000** | 0.996 | 0.987 | 0.998 | **0.915** | 0.992 | 0.996 | 0.995 | 0.983 | 0.999 | 0.908 | 0.975 | 0.180 | 0.999 | 0.957 | **0.997** | 0.986 | 0.999 | 0.987 | 5 |
| components of the 1240K panel | Illumina610-Quad sites | 256,277 | 304,292 | 0.806 | **1.000** | **0.511** | 0.988 | **0.001** | **0.956** | **0.000** | 0.996 | 0.994 | 0.996 | 0.951 | 0.995 | 0.945 | 0.993 | 0.960 | **0.998** | 0.841 | 0.976 | 0.795 | **0.999** | 0.974 | **0.997** | 0.994 | 1.000 | 0.982 | 8 |
| | sites exclusive to Illumina610-Quad | 183,680 | 216,478 | 0.281 | **1.000** | **0.520** | 0.988 | **0.031** | **0.966** | **0.000** | 0.997 | 0.975 | 0.997 | 0.947 | 0.993 | 0.973 | 0.994 | 0.981 | 0.999 | 0.807 | 0.973 | 0.564 | 0.999 | 0.988 | **0.998** | 0.991 | 1.000 | 0.984 | 6 |
| | sites included in both Illumina610-Quad and HumanOrigins | 72,597 | 87,814 | 0.875 | 1.000 | **0.501** | 0.988 | **0.013** | **0.938** | **0.001** | 0.991 | 0.990 | 0.997 | **0.926** | 0.990 | 0.892 | **0.957** | 0.882 | **0.998** | 0.880 | 0.940 | 0.804 | **0.997** | **0.705** | **0.992** | 0.993 | 0.999 | 0.949 | 10 |
| | HumanOrigins sites | 244,922 | 354,460 | 0.944 | 1.000 | 0.689 | **0.982** | **0.170** | **0.965** | **0.000** | 0.995 | 0.995 | 0.997 | **0.919** | 0.991 | 0.975 | **0.988** | 0.968 | 0.999 | 0.754 | 0.953 | 0.040 | **0.999** | **0.844** | **0.995** | **0.983** | 0.998 | 0.979 | 10 |
| | sites exclusive to HumanOrigins | 171,249 | 266,646 | 0.771 | 1.000 | 0.873 | **0.966** | **0.569** | **0.963** | **0.093** | 0.994 | 0.987 | 0.995 | **0.896** | 0.988 | 0.980 | 0.993 | 0.942 | 0.999 | 0.583 | 0.917 | 0.009 | **0.999** | 0.900 | **0.994** | **0.971** | **0.995** | 0.969 | 9 |
| | 1240K, other sites | 67,096 | 92,301 | 0.842 | 1.000 | 0.905 | 0.990 | **0.366** | **0.984** | **0.002** | 0.995 | **0.642** | 0.993 | **0.651** | **0.976** | 0.816 | **0.987** | 0.602 | **0.998** | 0.913 | 0.965 | 0.197 | **0.999** | 0.952 | **0.998** | **0.929** | **0.992** | 0.959 | 12 |
| the largest panels included in the HumanOrigins array | panel 13 based on a San individual and Denisovan | 67,557 | 89,655 | 0.089 | **0.999** | **0.291** | **0.770** | **0.174** | **0.401** | **0.001** | **0.896** | **0.572** | **0.986** | 0.963 | 0.989 | 0.937 | **0.981** | 0.849 | **0.997** | **0.369** | **0.842** | 0.241 | **0.995** | **0.778** | **0.996** | **0.985** | **0.997** | 0.873 | 18 |
| | panel 4 based on a San individual | 52,862 | 94,493 | 0.205 | **0.999** | **0.254** | **0.980** | **0.796** | **0.954** | **0.093** | 0.992 | 0.984 | 0.994 | 0.974 | **0.979** | 0.673 | **0.983** | 0.764 | **0.999** | **0.383** | **0.793** | 0.415 | **0.996** | **0.884** | **0.995** | **0.929** | **0.998** | 0.964 | 16 |
| | panel 5 based on a Yoruba individual | 44,674 | 73,180 | 0.500 | **0.999** | 0.987 | 0.990 | **0.818** | **0.986** | **0.014** | 0.991 | 0.946 | 0.997 | **0.917** | **0.983** | 0.752 | **0.978** | 0.884 | 0.999 | 0.669 | **0.861** | 0.004 | **0.999** | 0.911 | **0.997** | **0.977** | **0.990** | 0.978 | 12 |
| | panels 4 and 5 | 46,701 | 157,126 | 0.921 | **0.999** | 0.642 | 0.986 | **0.901** | **0.972** | **0.006** | 0.997 | 0.999 | 0.996 | 0.944 | **0.980** | 0.996 | **0.976** | 0.851 | 0.999 | **0.549** | **0.875** | 0.005 | **0.999** | 0.994 | **0.994** | **0.952** | **0.997** | 0.978 | 12 |
| other enrichment panels | 1000K: transversions in 2 Yoruba ind. and in Altai Neand. | 364,079 | 590,775 | 0.940 | **0.998** | 0.726 | 0.998 | **0.497** | **0.984** | **0.138** | 0.997 | 0.907 | 0.999 | 0.982 | 0.995 | **0.964** | 0.997 | 0.817 | 0.999 | 0.973 | 0.991 | 0.678 | **0.999** | 0.973 | 0.999 | 0.990 | 0.998 | 0.987 | 5 |
| | 2200K = 1000K + 1240K | 814,915 | 1,190,758 | 0.918 | **0.999** | 0.695 | 0.998 | **0.233** | **0.977** | **0.014** | 0.997 | 0.972 | 0.999 | 0.969 | 0.995 | 0.996 | 0.996 | 0.963 | 0.999 | 0.955 | 0.985 | 0.880 | 0.999 | 0.995 | 0.998 | 0.991 | 0.999 | 0.993 | 4 |
| archaic ascertainment | transitions and transversions | 525,014 | 1,555,781 | **0.040** | **0.001** | **0.225** | **0.891** | **0.570** | **0.899** | 0.912 | **0.833** | 0.996 | 0.997 | 0.937 | 0.987 | 0.489 | 0.993 | **0.162** | 0.999 | 0.585 | **0.876** | 0.011 | **0.999** | 0.986 | 0.999 | **0.907** | **0.994** | 0.903 | 13 |
| | transversions | 165,249 | 484,675 | **0.036** | **0.000** | **0.130** | **0.888** | **0.455** | **0.888** | 0.917 | **0.832** | **0.213** | **0.992** | 0.958 | 0.988 | 0.822 | 0.993 | **0.142** | 0.999 | **0.345** | **0.740** | 0.022 | **0.999** | 0.935 | 0.999 | **0.855** | **0.991** | 0.872 | 15 |
| MAF ascertainment | global, >5% MAF | 2,129,201 | 2,511,335 | 0.933 | 1.000 | **0.531** | 0.995 | **0.295** | **0.984** | **0.123** | 0.998 | 0.990 | 0.999 | 0.976 | 0.995 | 0.990 | 0.998 | 0.829 | 1.000 | 0.987 | 0.997 | 0.951 | 1.000 | 0.997 | 0.999 | 0.999 | 1.000 | 0.995 | 4 |
| | >5% MAF in Africans unadmixed with non-Africans | 2,045,769 | 3,231,875 | 0.883 | 1.000 | 0.976 | 1.000 | **0.768** | 0.992 | **0.069** | 0.996 | 0.997 | 1.000 | 0.998 | 0.998 | 0.998 | 1.000 | 0.990 | 1.000 | 0.946 | 0.990 | 0.929 | 1.000 | 0.993 | 0.999 | 0.999 | 0.999 | 0.998 | 2 |
| | >5% MAF in all Africans | 2,109,808 | 3,120,326 | 0.895 | 1.000 | 0.930 | 1.000 | **0.824** | 0.995 | **0.131** | 0.997 | 0.996 | 1.000 | 0.993 | 0.999 | 0.998 | 0.999 | 0.986 | 1.000 | 0.995 | 0.995 | 0.954 | 1.000 | 0.992 | 0.999 | 0.999 | 1.000 | 0.997 | 2 |
| | >5% MAF in Native Americans | 1,513,207 | 1,764,715 | 0.868 | 1.000 | **0.419** | 0.995 | **0.030** | **0.970** | **0.024** | 0.995 | 0.997 | 0.996 | **0.872** | **0.984** | 0.962 | 0.999 | 0.672 | 1.000 | 0.771 | 0.978 | 0.298 | 1.000 | 0.982 | **0.998** | **0.975** | 0.999 | 0.980 | 8 |
| | >5% MAF in Central Asians and Siberians | 1,843,262 | 2,150,675 | 0.819 | 1.000 | **0.461** | 0.995 | **0.068** | **0.975** | **0.046** | 0.998 | 0.995 | 0.999 | **0.919** | 0.988 | 0.947 | 0.998 | 0.803 | 1.000 | 0.774 | 0.982 | 0.749 | 0.999 | 0.993 | 0.999 | 0.998 | 1.000 | 0.991 | 5 |
| | >5% MAF in East Asians | 1,723,831 | 2,020,860 | 0.822 | 1.000 | **0.447** | 0.994 | **0.052** | **0.974** | **0.018** | 0.995 | 0.974 | 0.998 | 0.943 | 0.990 | 0.968 | 0.998 | 0.868 | 1.000 | 0.975 | 0.994 | 0.929 | 0.999 | 0.986 | **0.998** | 0.996 | 0.999 | 0.988 | 5 |
| | >5% MAF in Europeans | 1,885,336 | 2,192,571 | 0.917 | 1.000 | **0.485** | 0.996 | **0.391** | **0.987** | **0.030** | 0.998 | 0.994 | 0.999 | 0.937 | 0.990 | 0.984 | 0.999 | 0.929 | 1.000 | 0.731 | 0.977 | 0.880 | **0.999** | 0.994 | 0.999 | 0.995 | 1.000 | 0.992 | 5 |
| | >5% MAF in Middle Eastern groups | 2,018,884 | 2,306,319 | 0.808 | 1.000 | **0.443** | 0.995 | **0.468** | **0.988** | **0.006** | 0.999 | 0.997 | 0.999 | **0.926** | 0.991 | 0.988 | 0.999 | 0.887 | 0.999 | 0.941 | 0.984 | 0.920 | 1.000 | 0.994 | 0.999 | 0.997 | 0.999 | 0.993 | 5 |
| | >5% MAF in Papuans and Aboriginal Australians | 1,515,022 | 1,791,390 | 0.813 | 1.000 | **0.414** | 0.990 | **0.001** | **0.965** | **0.001** | 0.992 | 0.996 | 0.997 | **0.889** | 0.987 | 0.875 | 0.997 | 0.655 | 1.000 | 0.669 | 0.961 | 0.103 | 0.999 | 0.964 | **0.998** | 0.995 | 0.999 | 0.976 | 6 |
| | >5% MAF in South Asians | 1,908,459 | 2,235,024 | 0.888 | 1.000 | **0.466** | 0.995 | **0.199** | **0.981** | **0.087** | 0.998 | 0.990 | 0.999 | **0.917** | 0.990 | 0.927 | 0.999 | 0.877 | 1.000 | 0.848 | 0.989 | 0.942 | **0.999** | 0.996 | 0.999 | 0.995 | 1.000 | 0.990 | 6 |
| AT/GC mutation types + MAF ascertainment | global, >>5% MAF | 323,296 | 378,287 | 0.867 | 1.000 | **0.537** | 0.994 | **0.310** | **0.985** | **0.110** | 0.995 | 0.944 | 0.995 | 0.957 | 0.992 | 0.970 | 0.995 | 0.468 | 0.999 | 0.932 | 0.989 | 0.797 | **0.999** | 0.997 | 0.999 | 0.995 | 1.000 | 0.991 | 5 |
| | >5% MAF in Africans unadmixed with non-Africans | 309,172 | 486,906 | 0.867 | 1.000 | 0.982 | 0.998 | **0.764** | 0.992 | **0.107** | 0.998 | 0.955 | 0.997 | 0.989 | 0.994 | 0.925 | 0.998 | 0.833 | 0.999 | 0.823 | 0.969 | 0.906 | **0.999** | 0.996 | 0.999 | 0.996 | 0.999 | 0.993 | 3 |
| | >5% MAF in all Africans | 319,053 | 470,070 | 0.862 | 1.000 | 0.942 | 0.998 | **0.801** | 0.995 | **0.122** | 0.997 | 0.955 | 0.996 | 0.979 | 0.995 | 0.943 | 0.997 | 0.792 | 0.999 | 0.932 | 0.982 | 0.813 | **0.998** | 0.992 | 0.999 | 0.992 | 0.999 | 0.992 | 3 |
| | >5% MAF in Native Americans | 229,939 | 266,113 | 0.883 | 1.000 | **0.495** | 0.995 | **0.045** | **0.975** | **0.069** | 0.996 | 0.959 | **0.990** | **0.850** | **0.982** | 0.857 | 0.997 | 0.344 | 0.999 | 0.784 | 0.976 | 0.929 | **0.999** | 0.965 | 0.999 | **0.974** | 0.999 | 0.975 | 9 |
| | >5% MAF in Central Asians and Siberians | 280,103 | 324,245 | 0.779 | 1.000 | **0.526** | 0.995 | **0.103** | **0.977** | **0.081** | 0.997 | 0.979 | 0.996 | **0.906** | **0.984** | 0.828 | 0.994 | 0.564 | 0.999 | 0.632 | 0.936 | 0.888 | **0.998** | 0.988 | 1.000 | 0.994 | 1.000 | 0.981 | 7 |
| | >5% MAF in East Asians | 261,857 | 304,567 | 0.775 | 1.000 | **0.554** | 0.995 | **0.049** | **0.975** | **0.042** | 0.996 | 0.949 | 0.994 | **0.913** | **0.985** | 0.901 | 0.994 | 0.463 | 0.999 | 0.851 | 0.976 | 0.865 | **0.999** | 0.989 | 0.999 | 0.993 | 0.999 | 0.980 | 7 |
| | >5% MAF in Europeans | 285,723 | 330,244 | 0.755 | 1.000 | **0.539** | 0.995 | **0.413** | 0.990 | **0.019** | 0.998 | 0.955 | 0.998 | **0.921** | 0.988 | 0.918 | 0.996 | 0.741 | 0.999 | 0.683 | 0.954 | 0.777 | **0.998** | 0.988 | 0.999 | 0.989 | 0.999 | 0.988 | 5 |
| | >5% MAF in Middle Eastern groups | 306,450 | 347,536 | 0.793 | 1.000 | **0.460** | 0.993 | **0.478** | 0.992 | **0.019** | 0.999 | 0.964 | 0.996 | **0.863** | 0.987 | 0.923 | 0.996 | 0.557 | 0.999 | 0.965 | 0.995 | 0.788 | **0.998** | 0.985 | 0.999 | 0.990 | 0.999 | 0.989 | 5 |
| | >5% MAF in Papuans and Aboriginal Australians | 230,124 | 272,093 | 0.825 | 1.000 | **0.415** | 0.990 | **0.001** | **0.960** | **0.002** | 0.994 | 0.986 | 0.996 | **0.884** | 0.987 | 0.715 | **0.990** | **0.222** | 0.999 | 0.584 | 0.914 | 0.078 | **0.999** | 0.950 | **0.997** | 0.989 | 0.999 | 0.973 | 9 |
| | >5% MAF in South Asians | 289,739 | 336,996 | 0.848 | 1.000 | **0.544** | 0.995 | **0.240** | **0.984** | **0.102** | 0.998 | 0.950 | 0.995 | **0.892** | 0.987 | 0.931 | 0.996 | 0.626 | 0.999 | 0.829 | 0.976 | 0.783 | **0.998** | 0.991 | 0.999 | 0.992 | 0.999 | 0.986 | 6 |
| * the SNP counts correspond to sites polymorphic in larger collections of groups from which the analyzed population quintuplets were taken, see Suppl. Table X. | | | **median =>** | 0.845 | 1.000 | 0.528 | 0.995 | 0.303 | 0.976 | 0.036 | 0.996 | 0.982 | 0.997 | 0.931 | 0.990 | 0.944 | 0.996 | 0.831 | 0.999 | 0.826 | 0.976 | 0.785 | 0.999 | 0.987 | 0.999 | 0.991 | 0.999 | | |

**Suppl. Table 4.** Performance of ascertainment schemes explored across 12 population quintuplets and assessed as the correlation of admixture graph worst residuals (WR) using $R^2$ of linear trends, based on 5,000 graphs that are best-fitting on all sites, or based on all graphs. We also applied the binary classifier to determine if the ascertainment produces unbiased or biased results (the latter highlighted in bold and underlined text). Median $R^2$ values across all population quintuplets or ascertainment schemes are shown in the second rightmost column and in the bottom row, respectively, and the numbers of population quintuplets affected by bias (according to this classifier) are shown in the rightmost column. The composition of the population sets is shown above the table in an abbreviated way: *arch*, archaic humans, followed by the number of archaic groups; *afr*, Africans; *nafr*, non-Africans or Africans with substantial non-African admixture (Fan et al. 2019). The same results based on log-likelihood scores of admixture graphs (LL) are shown in Suppl. Table 5.

**LL, R2**

| ascertainment type | further details on the ascertainment | min. size of the SNP panel* | max. size of the SNP panel* | *arch 2, afr 3*: Denisovan, Altai, Yoruba, Dinka, Bulala | | *arch 1, afr 4*: Denisovan, Khomani San, Mbuti, Dinka, Mursi | | *arch 1, afr 3, nafr 1*: Altai, Ju hoan North, Biaka, Yoruba, Agaw | | *arch 1, afr 2, nafr 2*: Altai, Ju hoan North, Luhya, Palestinian, Spanish | | *afr 5*: Mbuti, Baka, Laka, Fulani, Bantu Tswana | | *afr 4, nafr 1*: Khomani San, Bakola, Igbo, Mursi, Aari | | *afr 4, nafr 1*: Bedzan, Cameroon SMA, Esan, Mozabite, Masai | | *afr 3, nafr 2*: Mbuti, Biaka, Ngumba, LBK, Iranian | | *afr 1, nafr 4*: Luo, Bedouin B, Jordanian, Abkhasian, Sardinian | | *nafr 5*: Australian, Quechua, Mayan, Lezgin, French | | *nafr 5*: Papuan, Chipewyan, Eskimo Naukan, Finnish, Sardinian | | *nafr 5*: Karitiana, Cree, Eskimo Sireniki, Hungarian, Icelandic | | med. | number of biased pop. sets |
|---|---|---|---|---|---|---|---|---|---|---|---|---|---|---|---|---|---|---|---|---|---|---|---|---|---|---|---|---|---|
| | | | | 5000 | all | 5000 | all | 5000 | all | 5000 | all | 5000 | all | 5000 | all | 5000 | all | 5000 | all | 5000 | all | 5000 | all | 5000 | all | 5000 | all | | |
| A<>T and G<>C mutations | | 805,042 | 1,757,840 | 0.969 | 1.000 | 0.945 | 0.998 | 0.998 | 0.999 | 0.972 | 0.996 | 0.992 | 0.999 | 0.992 | 0.999 | 0.936 | 0.999 | 0.763 | 1.000 | 0.945 | 0.992 | 0.882 | 1.000 | 0.984 | 1.000 | 0.999 | 1.000 | 0.994 | 0 |
| 1240K panel | | 501,429 | 663,239 | 0.903 | 1.000 | 0.521 | 0.992 | **0.154** | 0.989 | **0.000** | 0.995 | 0.994 | 0.998 | **0.844** | 0.995 | 0.997 | 0.994 | 0.981 | 0.999 | 0.919 | 0.988 | 0.128 | 0.999 | 0.917 | **0.998** | **0.989** | 1.000 | 0.991 | 5 |
| components of the 1240K panel | Illumina610-Quad sites | 256,277 | 304,292 | 0.865 | **0.999** | 0.414 | 0.990 | **0.007** | **0.984** | **0.001** | 0.992 | 0.997 | 0.997 | 0.909 | 0.997 | 0.952 | 0.993 | 0.933 | **0.998** | 0.934 | 0.988 | 0.888 | 0.999 | 0.936 | **0.997** | 0.995 | 1.000 | 0.986 | 6 |
| | sites exclusive to Illumina610-Quad | 183,680 | 216,478 | 0.204 | **0.999** | 0.423 | 0.988 | **0.077** | **0.984** | **0.000** | 0.995 | 0.991 | 0.997 | 0.897 | 0.995 | 0.980 | 0.994 | 0.959 | 0.999 | 0.913 | 0.987 | 0.756 | 1.000 | 0.937 | 0.998 | **0.993** | 0.999 | 0.985 | 5 |
| | sites included in both Illumina610-Quad and HumanOrigins | 72,597 | 87,814 | 0.910 | **0.999** | 0.405 | 0.982 | **0.039** | **0.975** | **0.003** | 0.992 | 0.994 | 0.996 | **0.853** | **0.989** | 0.883 | **0.957** | 0.735 | 0.999 | 0.912 | 0.949 | 0.860 | **0.998** | **0.651** | **0.992** | 0.997 | 0.998 | 0.953 | 10 |
| | HumanOrigins sites | 244,922 | 354,460 | 0.950 | 1.000 | 0.543 | **0.974** | **0.287** | **0.977** | **0.000** | 0.996 | 0.999 | 0.998 | **0.857** | 0.993 | 0.960 | **0.985** | 0.952 | 0.999 | 0.797 | 0.980 | 0.065 | 0.999 | 0.798 | **0.996** | **0.986** | 0.997 | 0.979 | 8 |
| | sites exclusive to HumanOrigins | 171,249 | 266,646 | 0.229 | 1.000 | 0.784 | **0.957** | **0.733** | **0.967** | **0.207** | 0.994 | 0.998 | 0.997 | **0.812** | **0.991** | 0.916 | **0.989** | 0.957 | 0.999 | 0.717 | 0.971 | **0.002** | **0.999** | 0.880 | **0.997** | **0.972** | **0.992** | 0.969 | 12 |
| | 1240K, other sites | 67,096 | 92,301 | 0.914 | **0.999** | 0.727 | 0.986 | **0.447** | **0.988** | **0.002** | 0.993 | **0.801** | 0.995 | **0.370** | **0.986** | 0.890 | **0.985** | 0.742 | 0.999 | 0.924 | 0.971 | 0.064 | 0.999 | 0.905 | **0.997** | **0.940** | **0.993** | 0.947 | 11 |
| the largest panels included in the HumanOrigins array | panel 13 based on a San individual and Denisovan | 67,557 | 89,655 | 0.062 | 1.000 | **0.351** | **0.695** | **0.307** | **0.407** | **0.000** | **0.878** | **0.833** | **0.992** | 0.958 | 0.994 | 0.740 | **0.975** | 0.915 | **0.996** | 0.648 | **0.892** | 0.382 | **0.997** | **0.778** | **0.998** | **0.985** | **0.995** | 0.885 | 16 |
| | panel 4 based on a San individual | 52,862 | 94,493 | 0.348 | **0.999** | **0.270** | **0.975** | **0.923** | **0.961** | **0.222** | 0.991 | 0.990 | 0.995 | 0.957 | **0.986** | **0.282** | **0.979** | 0.609 | **0.998** | 0.640 | **0.881** | 0.629 | **0.998** | 0.810 | **0.996** | **0.910** | 0.997 | 0.959 | 14 |
| | panel 5 based on a Yoruba individual | 44,674 | 73,180 | 0.516 | **0.998** | 0.977 | 0.983 | **0.917** | 0.990 | **0.015** | 0.988 | 0.970 | 0.995 | 0.869 | **0.986** | 0.746 | **0.964** | 0.752 | **0.998** | 0.715 | **0.905** | 0.008 | 0.999 | 0.791 | **0.997** | **0.980** | **0.987** | 0.941 | 10 |
| | panels 4 and 5 | 46,701 | 157,126 | 0.936 | **0.999** | 0.475 | 0.981 | **0.955** | **0.974** | **0.007** | 0.997 | 0.999 | 0.998 | 0.895 | **0.987** | 0.979 | **0.965** | 0.705 | 0.999 | 0.671 | **0.918** | 0.025 | **0.999** | 0.983 | **0.995** | **0.945** | **0.996** | 0.977 | 11 |
| other enrichment panels | 1000K: transversions in 2 Yoruba ind. and in Altai Neand. | 364,079 | 590,775 | 0.955 | **0.996** | 0.571 | 0.998 | **0.572** | 0.991 | **0.257** | 0.997 | 0.977 | 0.999 | 0.968 | 0.996 | 0.947 | 0.995 | 0.660 | 0.999 | 0.968 | 0.992 | 0.778 | **0.998** | 0.902 | 1.000 | 0.994 | **0.996** | 0.984 | 5 |
| | 2200K = 1000K + 1240K | 814,915 | 1,190,758 | 0.945 | **0.999** | 0.545 | 0.998 | **0.279** | 0.991 | **0.051** | 0.996 | 0.992 | 0.999 | 0.948 | 0.997 | 0.990 | 0.995 | 0.944 | 1.000 | 0.950 | 0.993 | 0.926 | **0.999** | 0.959 | 0.999 | **0.993** | 0.999 | 0.991 | 5 |
| archaic ascertainment | transitions and transversions | 525,014 | 1,555,781 | 0.040 | **0.006** | **0.317** | **0.836** | **0.793** | **0.891** | **0.979** | **0.770** | 0.994 | 0.998 | 0.946 | 0.995 | **0.183** | 0.994 | **0.018** | 0.999 | 0.425 | **0.910** | 0.006 | **0.999** | 0.949 | 0.999 | **0.975** | **0.993** | 0.948 | 12 |
| | transversions | 165,249 | 484,675 | 0.039 | **0.006** | **0.151** | **0.838** | **0.685** | **0.871** | **0.980** | **0.765** | **0.300** | 0.998 | 0.942 | 0.996 | 0.527 | 0.990 | **0.004** | 0.999 | **0.171** | **0.815** | 0.010 | 0.999 | 0.817 | 0.999 | **0.962** | **0.991** | 0.844 | 12 |
| MAF ascertainment | global, >5% MAF | 2,129,201 | 2,511,335 | 0.951 | 1.000 | 0.428 | 0.997 | **0.401** | 0.995 | **0.241** | 0.997 | 0.999 | 0.999 | 0.956 | 0.997 | 0.980 | 0.997 | 0.608 | 1.000 | 0.994 | 0.998 | 0.957 | 0.999 | 0.970 | 0.999 | 0.999 | 1.000 | 0.996 | 2 |
| | >5% MAF in Africans unadmixed with non-Africans | 2,045,769 | 3,231,875 | 0.919 | **0.999** | 0.939 | 0.999 | **0.869** | 0.997 | **0.176** | 0.995 | 1.000 | 1.000 | 0.995 | 0.999 | 0.997 | 1.000 | 0.981 | 1.000 | 0.969 | 0.993 | 0.957 | 0.999 | 0.952 | 0.999 | 0.999 | 0.999 | 0.996 | 3 |
| | >5% MAF in all Africans | 2,109,808 | 3,120,326 | 0.925 | **0.999** | 0.837 | 0.999 | **0.916** | 0.998 | **0.256** | 0.996 | 1.000 | 1.000 | 0.987 | 0.999 | 0.998 | 0.999 | 0.969 | 1.000 | 0.994 | 0.996 | 0.961 | 0.999 | 0.961 | 0.999 | 0.999 | 0.999 | 0.997 | 3 |
| | >5% MAF in Native Americans | 1,513,207 | 1,764,715 | 0.921 | 1.000 | 0.382 | 0.997 | **0.083** | 0.992 | **0.069** | 0.995 | 0.998 | 0.995 | **0.756** | **0.989** | 0.935 | 0.999 | 0.332 | 1.000 | 0.808 | 0.990 | 0.390 | 0.999 | 0.968 | **0.998** | **0.973** | 0.999 | 0.989 | 6 |
| | >5% MAF in Central Asians and Siberians | 1,843,262 | 2,150,675 | 0.394 | 1.000 | 0.397 | 0.998 | **0.112** | 0.994 | **0.131** | 0.997 | 0.996 | 0.999 | **0.830** | **0.991** | 0.882 | 0.998 | 0.570 | 1.000 | 0.734 | 0.992 | 0.824 | **0.999** | 0.957 | 0.999 | 0.998 | 1.000 | 0.991 | 5 |
| | >5% MAF in East Asians | 1,723,831 | 2,020,860 | 0.914 | 1.000 | 0.387 | 0.997 | **0.125** | 0.993 | **0.051** | 0.996 | 0.983 | 0.998 | 0.885 | 0.992 | 0.929 | 0.998 | 0.719 | 1.000 | 0.984 | 0.995 | 0.953 | 1.000 | 0.963 | 0.999 | 0.997 | 0.999 | 0.988 | 2 |
| | >5% MAF in Europeans | 1,885,336 | 2,192,571 | 0.944 | 1.000 | 0.405 | 0.998 | **0.528** | 0.997 | **0.157** | 0.998 | 0.997 | 0.999 | 0.869 | 0.993 | 0.982 | 0.998 | 0.862 | 1.000 | 0.689 | 0.988 | 0.931 | **0.999** | 0.955 | 0.999 | 0.996 | 1.000 | 0.990 | 3 |
| | >5% MAF in Middle Eastern groups | 2,018,884 | 2,306,319 | 0.150 | 1.000 | 0.384 | 0.998 | **0.594** | 0.997 | **0.089** | 0.999 | 0.999 | 0.999 | **0.858** | 0.994 | 0.986 | 0.999 | 0.761 | 1.000 | 0.953 | 0.992 | 0.950 | 0.999 | 0.959 | 0.999 | 0.998 | 0.999 | 0.993 | 3 |
| | >5% MAF in Papuans and Aboriginal Australians | 1,515,022 | 1,791,390 | 0.894 | 1.000 | 0.374 | 0.993 | **0.019** | 0.991 | **0.001** | 0.991 | 0.997 | 0.998 | **0.808** | **0.990** | 0.613 | 0.996 | 0.313 | 1.000 | 0.533 | 0.983 | 0.009 | 0.999 | 0.878 | **0.997** | 0.999 | 0.999 | 0.987 | 5 |
| | >5% MAF in South Asians | 1,908,459 | 2,235,024 | 0.939 | 1.000 | 0.395 | 0.998 | **0.316** | 0.995 | **0.196** | 0.998 | 0.994 | 0.999 | **0.831** | 0.992 | 0.836 | 0.998 | 0.738 | 1.000 | 0.870 | 0.994 | 0.960 | **0.998** | 0.960 | 0.999 | 0.996 | 1.000 | 0.993 | 4 |
| AT/GC mutation types + MAF ascertainment | global, >>5% MAF | 323,296 | 378,287 | 0.906 | 0.999 | 0.430 | 0.996 | **0.434** | 0.996 | **0.223** | 0.990 | 0.991 | 0.997 | 0.910 | 0.995 | 0.954 | 0.995 | 0.091 | 1.000 | 0.947 | 0.992 | 0.868 | **0.999** | 0.980 | 0.999 | 0.999 | 1.000 | 0.990 | 3 |
| | >5% MAF in Africans unadmixed with non-Africans | 309,172 | 486,906 | 0.296 | **0.999** | 0.961 | 0.996 | **0.868** | 0.996 | **0.223** | 0.997 | 0.994 | 0.998 | 0.977 | 0.996 | 0.925 | 0.999 | 0.667 | 0.999 | 0.939 | 0.982 | 0.933 | **0.999** | 0.979 | 1.000 | 0.999 | 0.999 | 0.988 | 4 |
| | >5% MAF in all Africans | 319,053 | 470,070 | 0.271 | **0.999** | 0.868 | 0.996 | **0.897** | 0.997 | **0.235** | 0.995 | 0.994 | 0.998 | 0.959 | 0.997 | 0.930 | 0.998 | 0.571 | 0.999 | 0.953 | 0.987 | 0.893 | **0.998** | 0.970 | 0.999 | 0.997 | 1.000 | 0.990 | 4 |
| | >5% MAF in Native Americans | 229,939 | 266,113 | 0.933 | 1.000 | 0.423 | 0.995 | **0.107** | 0.990 | **0.171** | 0.997 | 0.993 | **0.991** | **0.696** | **0.986** | 0.702 | 0.998 | 0.031 | 0.999 | 0.702 | 0.986 | 0.946 | **0.999** | 0.959 | 0.999 | **0.980** | 0.999 | 0.986 | 7 |
| | >5% MAF in Central Asians and Siberians | 280,103 | 324,245 | 0.844 | 1.000 | 0.428 | 0.995 | **0.189** | 0.993 | **0.189** | 0.998 | 0.997 | 0.996 | **0.800** | **0.988** | 0.564 | 0.995 | 0.200 | 1.000 | 0.425 | 0.965 | 0.915 | **0.998** | 0.974 | 0.999 | 0.998 | 1.000 | 0.981 | 5 |
| | >5% MAF in East Asians | 261,857 | 304,567 | 0.862 | 1.000 | 0.440 | 0.993 | **0.114** | 0.992 | **0.128** | 0.996 | 0.990 | 0.996 | **0.813** | **0.988** | 0.723 | 0.996 | 0.130 | 1.000 | 0.786 | 0.983 | 0.894 | 0.999 | 0.971 | 0.999 | 0.994 | 0.999 | 0.985 | 4 |
| | >5% MAF in Europeans | 285,723 | 330,244 | 0.833 | 1.000 | 0.429 | 0.994 | **0.574** | 0.997 | **0.160** | 0.999 | 0.995 | 0.998 | **0.838** | **0.992** | 0.851 | 0.996 | 0.487 | 1.000 | 0.494 | 0.975 | 0.862 | **0.998** | 0.966 | 0.999 | 0.996 | 0.999 | 0.984 | 5 |
| | >5% MAF in Middle Eastern groups | 306,450 | 347,536 | 0.210 | 1.000 | 0.390 | 0.993 | **0.624** | 0.998 | **0.181** | 0.999 | 0.991 | 0.996 | **0.724** | **0.991** | 0.864 | 0.997 | 0.188 | 0.999 | 0.978 | 0.996 | 0.852 | **0.999** | 0.962 | 0.999 | 0.997 | 0.999 | 0.991 | 5 |
| | >5% MAF in Papuans and Aboriginal Australians | 230,124 | 272,093 | 0.902 | 1.000 | 0.373 | 0.991 | **0.016** | **0.987** | **0.003** | 0.995 | 0.999 | 0.996 | **0.785** | **0.991** | **0.317** | 0.993 | **0.004** | 1.000 | **0.360** | 0.953 | 0.045 | 0.999 | 0.923 | **0.996** | 0.997 | 1.000 | 0.970 | 9 |
| | >5% MAF in South Asians | 289,739 | 336,996 | 0.895 | 1.000 | 0.433 | 0.995 | **0.395** | 0.994 | **0.218** | 0.998 | 0.994 | 0.996 | **0.776** | **0.991** | 0.792 | 0.995 | 0.268 | 1.000 | 0.743 | 0.984 | 0.865 | **0.998** | 0.972 | 0.999 | 0.997 | 1.000 | 0.987 | 5 |
| * the SNP counts correspond to sites polymorphic in larger collections of groups from which the analyzed population quintuplets were taken, see Suppl. Table X. | | | **median =>** | 0.899 | 1.000 | 0.428 | 0.994 | 0.417 | 0.991 | 0.159 | 0.996 | 0.994 | 0.998 | 0.869 | 0.992 | 0.921 | 0.995 | 0.686 | 1.000 | 0.839 | 0.986 | 0.861 | 0.999 | 0.958 | 0.999 | 0.996 | 0.999 | | |

**Suppl. Table 5.** Performance of ascertainment schemes explored across 12 population quintuplets assessed as the correlation of log-likelihood scores of admixture graphs (LL) using $R^2$ of linear trends, based on 5,000 graphs that are best-fitting on all sites, or based on all graphs. We also applied the binary classifier to determine if the ascertainment produces unbiased or biased results (the latter highlighted in bold and underlined text). Median $R^2$ values across all population quintuplets or ascertainment schemes are shown in the second rightmost column and in the bottom row, respectively, and the numbers of population quintuplets affected by bias (according to this classifier) are shown in the rightmost column. The composition of the population sets is shown above the table in an abbreviated way: *arch*, archaic humans, followed by the number of archaic groups; *afr*, Africans; *nafr*, non-Africans or Africans with substantial non-African admixture (Fan et al. 2019). The same results based on worst residuals of admixture graphs (WR) are shown in Suppl. Table 4.

| | | Neanderthal=>nAFR gene flow | | | | | | no Neanderthal=>nAFR gene flow | | | | | |
|---|---|---|---|---|---|---|---|---|---|---|---|---|---|
| | | percentage of models rejected on asc. data and accepted on all sites | | | percentage of models accepted on asc. data and rejected on all sites | | | percentage of models rejected on asc. data and accepted on all sites | | | percentage of models accepted on asc. data and rejected on all sites | | |
| ascertainment | simulation iteration | d | n1 | n2 | d | n1 | n2 | d | n1 | n2 | d | n1 | n2 |
| archaic | 1 | **11.681%** | 0.000% | 0.000% | 0.003% | 0.000% | 0.000% | 0.006% | 0.006% | 0.006% | 0.000% | 0.000% | 0.000% |
| | 2 | **11.675%** | 0.000% | 0.000% | 0.000% | 0.000% | 0.003% | **0.003%** | 0.000% | 0.000% | 0.000% | 0.000% | 0.003% |
| | 3 | **11.681%** | 0.000% | **0.003%** | 0.000% | 0.003% | 0.000% | **0.061%** | 0.131% | 0.067% | 0.000% | 0.009% | 0.052% |
| | 4 | **11.681%** | 0.000% | 0.000% | 0.000% | 0.006% | 0.000% | 0.000% | 0.000% | 0.000% | 0.006% | 0.000% | 0.000% |
| | 5 | **11.684%** | 0.000% | 0.003% | 0.009% | 0.000% | 0.003% | 0.000% | 0.000% | 0.000% | 0.003% | 0.000% | 0.131% |
| | 6 | **11.706%** | **0.003%** | 0.000% | 0.003% | 0.000% | 0.000% | 0.000% | 0.000% | 0.000% | 0.003% | 0.003% | 0.000% |
| | 7 | **11.681%** | 0.000% | **0.015%** | 0.000% | 0.003% | 0.003% | 0.024% | 0.009% | 0.128% | 0.000% | 0.000% | 0.003% |
| | 8 | **11.681%** | 0.000% | 0.000% | 0.000% | 0.000% | 0.000% | 0.000% | 0.000% | 0.000% | 0.000% | 0.003% | 0.000% |
| | 9 | **11.681%** | **0.018%** | **0.009%** | 0.000% | 0.003% | 0.003% | 0.031% | 5.796% | 0.049% | 0.049% | 0.034% | 0.089% |
| | 10 | **11.681%** | 0.000% | 0.000% | 0.000% | 0.000% | 0.000% | 0.000% | 0.000% | 0.000% | 0.000% | 0.000% | 0.000% |
| HO1 | 1 | **11.684%** | 0.000% | 0.000% | 0.003% | **11.681%** | **11.681%** | 0.061% | 0.150% | 0.104% | 0.000% | 0.000% | 0.000% |
| | 2 | **11.675%** | 0.000% | 0.000% | 0.000% | **11.681%** | **11.684%** | 0.000% | 0.000% | 0.000% | 0.000% | 0.000% | 0.003% |
| | 3 | **11.681%** | 0.000% | 0.000% | 0.000% | **11.684%** | **11.681%** | 0.000% | 0.000% | 0.000% | 0.000% | 0.015% | 0.128% |
| | 4 | **11.681%** | **0.006%** | **0.006%** | 0.000% | **11.678%** | **11.678%** | 0.003% | 0.000% | 0.000% | 0.006% | 0.000% | 0.000% |
| | 5 | **11.779%** | 0.000% | 0.000% | 0.009% | **11.681%** | **11.684%** | 0.000% | 0.000% | 0.000% | 0.003% | 0.000% | 0.131% |
| | 6 | **11.819%** | 0.000% | 0.000% | 0.003% | **11.681%** | **11.681%** | 0.000% | 0.003% | 0.000% | 0.003% | 0.003% | 0.000% |
| | 7 | **11.681%** | 0.000% | 0.000% | 0.000% | **11.684%** | **11.684%** | 0.000% | 0.000% | 0.000% | 0.000% | 0.000% | 0.003% |
| | 8 | **11.681%** | 0.000% | 0.000% | 0.000% | **11.681%** | **11.681%** | 0.000% | 0.000% | 0.000% | 0.000% | 0.003% | 0.000% |
| | 9 | **17.688%** | **5.347%** | **5.353%** | 0.000% | **6.187%** | **6.178%** | 0.034% | 0.024% | 0.024% | 0.067% | 0.504% | 0.113% |
| | 10 | **11.684%** | 0.000% | 0.000% | 0.000% | **11.681%** | **11.681%** | 0.000% | 0.000% | 0.000% | 0.000% | 0.000% | 0.000% |
| HO4 | 1 | 0.000% | 0.000% | **0.003%** | 0.003% | 0.000% | 0.000% | 0.000% | 0.000% | 0.003% | 0.000% | 0.000% | 0.000% |
| | 2 | **0.003%** | 0.000% | 0.003% | 0.006% | 0.000% | 0.003% | 0.000% | 0.000% | **0.003%** | 0.000% | 0.000% | 0.003% |
| | 3 | 0.000% | 0.000% | 0.000% | 0.000% | 0.003% | 0.000% | 0.000% | 0.009% | 0.052% | 0.000% | 0.006% | 0.076% |
| | 4 | **10.872%** | 0.000% | 0.003% | 0.000% | 0.006% | 0.000% | 0.000% | 0.107% | 0.000% | 0.006% | 0.000% | 0.000% |
| | 5 | 0.000% | 0.000% | 0.000% | 0.009% | 0.000% | 0.003% | 0.000% | 0.012% | 0.061% | 0.003% | 0.000% | 0.086% |
| | 6 | 0.000% | 0.000% | 0.000% | 0.003% | 0.000% | 0.000% | 0.000% | 0.000% | 0.000% | 0.003% | 0.003% | 0.000% |
| | 7 | 0.000% | 0.000% | 0.000% | 0.000% | 0.003% | 0.003% | 0.000% | 0.003% | 0.006% | 0.000% | 0.000% | 0.003% |
| | 8 | 0.000% | 0.000% | 0.000% | 0.000% | 0.000% | 0.000% | 0.000% | 0.000% | 0.003% | 0.000% | 0.003% | 0.000% |
| | 9 | 0.000% | 0.000% | 0.000% | 0.000% | 0.003% | 0.003% | **0.110%** | **6.181%** | 0.339% | 0.024% | 0.027% | 0.058% |
| | 10 | 0.000% | 0.000% | 0.000% | 0.000% | 0.000% | 0.000% | 0.000% | 0.006% | 0.000% | 0.000% | 0.000% | 0.000% |
| MAF AFR | 1 | **11.681%** | **0.003%** | 0.000% | 0.003% | 0.000% | **11.486%** | 0.000% | 0.006% | 0.000% | 0.000% | 0.000% | 0.000% |
| | 2 | **11.675%** | 0.000% | 0.000% | 0.000% | **11.681%** | **11.684%** | 0.000% | 0.000% | 0.000% | 0.000% | 0.000% | 0.003% |
| | 3 | **11.681%** | 0.000% | 0.000% | 0.000% | **11.684%** | **11.681%** | 0.000% | 0.000% | 0.000% | 0.000% | 0.015% | 0.128% |
| | 4 | **11.684%** | 0.000% | 0.000% | 0.000% | **11.687%** | **11.654%** | 0.000% | 0.003% | 0.000% | 0.006% | 0.000% | 0.000% |
| | 5 | **11.687%** | 0.000% | **0.241%** | 0.009% | **11.681%** | **1.044%** | 0.000% | 0.000% | 0.000% | 0.003% | 0.000% | 0.131% |
| | 6 | **11.687%** | 0.000% | 0.000% | 0.003% | **11.681%** | **1.072%** | 0.000% | 0.000% | 0.000% | 0.003% | 0.003% | 0.000% |
| | 7 | **11.681%** | 0.000% | 0.000% | 0.000% | **11.681%** | **11.684%** | 0.003% | 0.000% | 0.003% | 0.000% | 0.000% | 0.003% |
| | 8 | **11.681%** | 0.000% | 0.000% | 0.000% | **11.681%** | **11.681%** | 0.000% | 0.000% | 0.000% | 0.000% | 0.003% | 0.000% |
| | 9 | **11.687%** | 0.000% | 0.000% | 0.000% | **11.684%** | **11.681%** | 0.043% | 0.275% | 0.034% | 0.046% | 0.144% | 0.150% |
| | 10 | **11.696%** | **0.049%** | **0.192%** | 0.000% | **1.014%** | **1.063%** | 0.000% | 0.000% | 0.000% | 0.000% | 0.000% | 0.000% |
| MAF AMH | 1 | **0.003%** | 0.000% | 0.000% | 0.003% | 0.000% | 0.000% | 0.000% | **6.239%** | 0.003% | 0.000% | 0.000% | 0.000% |
| | 2 | 0.000% | 0.000% | **6.157%** | 0.006% | 0.000% | 0.000% | 0.000% | 0.000% | 0.000% | 0.000% | 0.000% | 0.003% |
| | 3 | 0.000% | 0.000% | 0.000% | 0.000% | 0.003% | 0.000% | 0.000% | 0.006% | 0.000% | 0.000% | 0.015% | 0.128% |
| | 4 | **0.360%** | 0.003% | 0.000% | 0.000% | 0.006% | 0.000% | 0.000% | 0.150% | **0.137%** | 0.006% | 0.000% | 0.000% |
| | 5 | **0.003%** | 0.000% | 0.000% | 0.009% | 0.000% | 0.003% | **6.215%** | **0.134%** | 0.101% | 0.003% | 0.000% | 0.061% |
| | 6 | 0.000% | 0.000% | 0.000% | 0.003% | 0.000% | 0.000% | **6.202%** | **6.236%** | **6.233%** | 0.003% | 0.003% | 0.000% |
| | 7 | **0.003%** | **0.006%** | 0.000% | 0.000% | 0.003% | 0.003% | 0.000% | 0.003% | 0.003% | 0.000% | 0.000% | 0.003% |
| | 8 | 0.000% | **0.003%** | 0.000% | 0.000% | 0.000% | 0.000% | 0.018% | 0.006% | **0.018%** | 0.000% | 0.003% | 0.000% |
| | 9 | **10.896%** | **5.821%** | **5.454%** | 0.000% | 0.000% | 0.003% | **6.770%** | **6.380%** | **6.731%** | 0.003% | 0.031% | 0.021% |
| | 10 | 0.000% | 0.000% | 0.000% | 0.000% | 0.000% | 0.000% | 0.000% | 0.000% | 0.006% | 0.000% | 0.000% | 0.000% |
| MAF nAFR | 1 | 0.000% | 0.000% | 0.000% | 0.003% | 0.000% | 0.000% | 0.260% | **6.319%** | **6.325%** | 0.000% | **0.947%** | **0.953%** |
| | 2 | 0.000% | 0.000% | 0.000% | 0.006% | 0.000% | 0.003% | **5.302%** | **5.314%** | **5.280%** | 0.000% | 0.000% | 0.000% |
| | 3 | 0.000% | 0.000% | 0.000% | 0.000% | 0.003% | 0.000% | **0.507%** | **0.247%** | **0.290%** | 0.000% | **1.038%** | 11.809% |
| | 4 | **11.925%** | 0.000% | 0.000% | 0.000% | 0.006% | 0.000% | 5.332% | **5.836%** | **5.326%** | 0.006% | 1.090% | 1.093% |
| | 5 | **10.744%** | 0.000% | 0.000% | 0.000% | 0.000% | 0.003% | **6.914%** | **5.323%** | **5.692%** | 0.003% | **0.507%** | **0.278%** |
| | 6 | **10.939%** | 0.000% | 0.000% | 0.000% | 0.000% | 0.000% | **6.911%** | **6.917%** | **6.902%** | **0.498%** | 0.003% | 0.000% |
| | 7 | 0.000% | 0.000% | 0.000% | 0.000% | 0.003% | 0.003% | **0.223%** | **0.238%** | **0.269%** | 0.000% | 0.000% | **0.244%** |
| | 8 | 0.000% | **0.003%** | 0.000% | 0.000% | 0.000% | 0.000% | **5.320%** | **5.323%** | **5.323%** | **0.983%** | 1.057% | 1.054% |
| | 9 | **10.851%** | 0.000% | 0.000% | 0.000% | 0.003% | 0.003% | **6.850%** | **6.401%** | **6.758%** | 0.003% | **1.078%** | 1.078% |
| | 10 | **11.681%** | 0.000% | 0.000% | 0.000% | 0.000% | 0.000% | **6.862%** | **5.317%** | **5.335%** | 0.000% | **1.026%** | **1.017%** |

**Suppl. Table 6.** Performance of ascertainment schemes on simulated data explored across three population quintuplets (including either "d", or "n1", or "n2" "archaic" individuals, in addition to the "a1", "a2", "na1", and "na2" groups) and assessed as the fraction of all topologies that are rejected under ascertainment (WR > 3 SE) but accepted on all sites (WR < 3 SE), or as the fraction of all topologies that are accepted under ascertainment (WR < 3 SE) but rejected on all sites (WR > 3 SE). We also applied the binary classifier (based on a $10^{th}$ percentile threshold and 10 random site subsamples matching the average size of the "Human Origins, one panel" set, 500K sites) to determine if the ascertainment produces unbiased or biased results (the latter highlighted in bold and underlined text). Ten independent simulations with the same parameters were performed, and the following ascertainment schemes were explored on each of them: 1) archaic ascertainment

(1.05M sites on average); 2) Human Origins-like ascertainment, one panel based on the “a2” group (500K sites on average across simulation iterations, abbreviated as “HO1”); 3) Human Origins-like ascertainment, four panes based on randomly selected individuals from four groups (“a1”, “a2”, “na1”, and “na2”, 1.34M sites on average, abbreviated as “HO4”); 4) African MAF ascertainment, that restricting to sites with MAF > 5% in the union of “a1” and “a2” groups (1.85M sites on average, abbreviated as “MAF AFR”); 5) similar MAF ascertainment on the union of “a1”, “a2”, “na1”, “na2” (1.62M sites on average, abbreviated as “MAF AMH”); 6) similar MAF ascertainment on the union of “na1” and “na2” groups (1.48M sites on average, abbreviated as “MAF nAFR”).

| residual SE of a linear trend for *f4*-statistic Z-scores (\|Z\|< 15 SE on all sites) | | no. f4-stat. | | 37,950 | 26,565 | 71,253 | 11,628 | 36,627 | 10,317 | 500,859 | 119,016 | 213,900 | 164,703 | 418,500 | 352,935 | 337,125 | 310,155 | 40,455 | 39,732 | 94,395 | 83,700 | 6,975 | 396 | 69,750 | 4,185 | 8,514 | 20,700 | 2,700 | 75 | 93 | |
|---|---|---|---|---|---|---|---|---|---|---|---|---|---|---|---|---|---|---|---|---|---|---|---|---|---|---|---|---|---|---|
| | | no. f4-stat., \|Z\|<15 SE | | 37,950 | 26,319 | 71,238 | 8,946 | 34,191 | 6,755 | 331,307 | 51,837 | 213,474 | 163,658 | 139,500 | 117,645 | 245,619 | 243,638 | 26,540 | 24,165 | 77,840 | 27,900 | 2,325 | 127 | 59,240 | 1,395 | 2,838 | 20,699 | 900 | 25 | 31 | |
| | | | | non-African 4 | | | | | | | | Afr. 1, non-Afr. 3 | | Afr. 2, non-Afr. 2 | | African 3, X | | | | Afr. 4 | African 1-2, archaic 1-3, X | | | | | | arch. 1-2, non-Afr. 2-3 | | arch. 3, non-Afr. 1 | | |
| ascertainment type | further details on the ascertainment | min. size of the SNP panel* | max. size of the SNP panel* | AFR 0, ARCH 0, ME 4 | AFR 0, EAS 4 | AMRSIB 0, EUR 4, PAP 0 | AMRSIB 4, EUR 0, PAP 0 | AMRSIB 0, EUR X, PAP Y | AMRSIB X, EUR 0, PAP Y | AMRSIB X, EUR Y, PAP 0 | AMRSIB X, EUR Y, PAP Z | AFR 1, ARCH 0, ME 3 | AFR 1, EAS 3 | AFR 2, ARCH 0, ME 2 | AFR 2, EAS 2 | AFR 3, ARCH 0, ME 1 | AFR 3, EAS 1 | AFR 3, ARCH 1, ME 0 | AFR 3, Chimp 1 | AFR 4, ARCH 0, ME 0 | AFR 1, ARCH 1, ME 2 | AFR 1, ARCH 2, ME 1 | AFR 1, ARCH 2, Chimp 1 | AFR 2, ARCH 1, ME 1 ** | AFR 2, ARCH 2, ME 0 | AFR 2, ARCH 1, Chimp 1 | AFR 0, ARCH 1, ME 3 | AFR 0, ARCH 2, ME 2 | AFR 0, ARCH 3, ME 1 | AFR 1, ARCH 3, ME 0 | number of biased f4-stat. classes |
| A<>T and G<>C mutations | | 805,042 | 3,610,461 | 0.36 | 0.32 | 0.34 | 0.33 | 0.32 | 0.30 | 0.34 | 0.33 | 0.38 | 0.34 | 0.49 | 0.43 | 0.46 | 0.37 | 0.48 | 0.59 | 0.46 | 0.42 | 0.51 | 0.31 | 0.40 | 0.59 | 0.66 | 0.30 | 0.54 | 0.12 | 0.13 | 0 |
| 1240K panel | | 501,429 | 763,188 | 0.41 | 0.40 | 0.41 | 0.42 | 0.47 | 0.42 | 0.44 | 0.45 | 0.43 | 0.46 | 0.50 | 0.45 | **1.23** | **1.18** | **2.11** | **2.05** | **0.79** | 0.65 | 0.73 | 1.16 | **6.65** | 0.79 | **1.40** | 0.70 | 0.58 | **0.30** | **0.29** | 9 |
| components of the 1240K panel | Illumina610-Quad sites | 256,277 | 354,539 | 0.54 | **0.58** | **0.53** | **0.56** | **0.63** | **0.55** | **0.58** | **0.62** | 0.59 | **0.62** | 0.67 | 0.63 | **1.30** | **1.29** | **2.51** | **2.25** | **0.95** | **0.84** | 0.80 | 0.75 | **7.89** | 0.93 | **1.49** | **0.75** | 0.86 | **0.38** | **0.38** | 19 |
| | sites exclusive to Illumina610-Quad | 183,680 | 255,216 | 0.57 | **0.61** | **0.59** | **0.58** | **0.64** | **0.53** | **0.61** | **0.63** | 0.63 | **0.69** | 0.72 | **0.72** | **1.29** | **1.34** | **2.29** | **2.05** | **0.99** | **0.89** | 0.89 | 0.56 | **7.06** | **0.97** | **1.19** | **0.83** | 0.86 | **0.35** | **0.37** | 21 |
| | sites included in both Illumina610-Quad and HumanOrigins | 72,597 | 99,323 | **0.81** | **0.78** | **0.73** | **0.75** | **0.80** | **0.92** | **0.80** | **0.87** | **0.82** | **0.74** | **0.88** | **0.81** | **1.22** | **1.06** | **1.97** | **1.74** | **1.01** | **0.88** | 0.79 | **1.21** | **6.11** | **0.98** | **1.36** | **0.81** | 0.87 | **0.38** | **0.38** | 25 |
| | HumanOrigins sites | 244,922 | 399,064 | 0.56 | 0.48 | **0.55** | **0.55** | **0.51** | **0.63** | **0.55** | **0.56** | 0.56 | 0.52 | 0.65 | 0.58 | **1.33** | **1.26** | **1.70** | **1.77** | **0.88** | **0.81** | 0.74 | **1.69** | **5.27** | 0.95 | **1.33** | 0.72 | 0.64 | 0.24 | **0.31** | 16 |
| | sites exclusive to HumanOrigins | 171,249 | 299,741 | 0.58 | 0.52 | **0.59** | **0.62** | **0.56** | **0.62** | **0.62** | **0.58** | 0.63 | 0.60 | 0.69 | 0.63 | **1.32** | **1.33** | **1.32** | **1.53** | **0.84** | **0.85** | 0.82 | **1.92** | **3.80** | 0.92 | **1.22** | **0.76** | 0.69 | 0.24 | **0.30** | 17 |
| | 1240K, other sites | 67,096 | 108,908 | **0.76** | **0.77** | **0.75** | **0.75** | **0.75** | **0.73** | **0.81** | **0.82** | **0.77** | **0.81** | **0.85** | **0.82** | **0.97** | **1.00** | **1.26** | **1.25** | **0.85** | **0.92** | 0.80 | 0.81 | **3.21** | **0.97** | 0.95 | **0.99** | 0.79 | **0.32** | **0.27** | 23 |
| the largest panels included in the HumanOrigins array | panel 13 based on a San individual and Denisovan | 67,557 | 90,136 | **0.85** | **0.76** | **0.79** | **0.80** | **0.75** | **0.77** | **0.79** | **0.79** | **0.86** | **0.78** | **0.88** | **0.84** | **1.19** | **1.16** | **1.85** | **1.81** | **0.87** | **0.93** | **1.75** | **3.86** | **2.64** | **2.79** | **2.54** | **1.02** | **0.92** | 0.22 | 0.24 | 25 |
| | panel 4 based on a San individual | 52,862 | 106,537 | **0.77** | **0.75** | **0.85** | **0.82** | **0.80** | **0.77** | **0.83** | **0.80** | **0.81** | **0.88** | **0.86** | **0.86** | **1.24** | **1.16** | **0.90** | **1.19** | **0.93** | **1.06** | 0.85 | 0.45 | **2.54** | 0.86 | 0.81 | **1.01** | **0.96** | **0.29** | **0.38** | 23 |
| | panel 5 based on a Yoruba individual | 44,674 | 82,895 | **0.83** | **0.83** | **0.84** | **0.74** | **0.78** | **0.78** | **0.86** | **0.81** | **0.86** | **0.84** | **0.89** | **0.80** | **0.80** | **0.84** | **1.14** | **1.05** | **0.92** | **0.89** | 0.85 | 0.48 | **1.86** | **0.99** | 0.99 | **0.89** | **0.93** | **0.33** | **0.34** | 24 |
| | panels 4 and 5 | 46,701 | 177,443 | **0.73** | **0.68** | **0.74** | **0.67** | **0.66** | **0.68** | **0.74** | **0.69** | **0.77** | **0.83** | **0.80** | **0.74** | **0.95** | **0.97** | **1.02** | **1.08** | **0.85** | **0.93** | 0.74 | 0.43 | **2.60** | 0.87 | 0.73 | **0.92** | 0.83 | **0.30** | **0.38** | 22 |
| other enrichment panels | 1000K: transversions in 2 Yoruba ind. and in Altai Neand. | 364,079 | 610,754 | 0.52 | 0.51 | **0.47** | **0.48** | 0.45 | 0.45 | **0.50** | **0.49** | 0.54 | 0.51 | 0.57 | 0.58 | 0.65 | **0.70** | **1.23** | **1.09** | 0.67 | 0.68 | 0.65 | 1.00 | **3.16** | 0.66 | **1.17** | 0.70 | 0.83 | 0.19 | 0.17 | 9 |
| | 2200K = 1000K + 1240K | 814,915 | 1,304,074 | 0.31 | 0.33 | 0.29 | 0.30 | 0.31 | 0.29 | 0.33 | 0.31 | 0.33 | 0.36 | 0.38 | 0.38 | **0.77** | **0.85** | **1.65** | **1.50** | 0.63 | 0.52 | 0.58 | 1.00 | **5.08** | 0.61 | **1.31** | 0.58 | 0.57 | 0.21 | 0.18 | 6 |
| archaic ascertainment | transitions and transversions | 525,014 | 1,555,781 | **0.78** | **0.75** | **0.81** | **0.75** | **0.78** | **0.74** | **0.76** | **0.78** | **0.87** | **0.78** | **0.81** | **0.84** | **1.01** | **0.98** | **1.16** | 0.92 | **0.76** | **0.88** | 0.09 | 0.04 | **6.03** | 0.02 | **1.62** | **1.11** | 0.04 | 0.02 | 0.01 | 20 |
| | transversions | 165,249 | 484,675 | **0.79** | **0.75** | **0.81** | **0.75** | **0.79** | **0.76** | **0.81** | **0.81** | **0.88** | **0.76** | **0.84** | **0.86** | **1.02** | **0.98** | **1.06** | **1.10** | **0.83** | **0.88** | 0.37 | 0.58 | **5.51** | 0.42 | **1.47** | **1.08** | 0.39 | 0.09 | 0.10 | 21 |
| MAF ascertainment | global, >>5% MAF | 2,129,201 | 3,387,302 | 0.09 | 0.08 | 0.10 | 0.10 | 0.08 | 0.12 | 0.10 | 0.09 | 0.11 | 0.12 | 0.08 | 0.04 | **0.86** | **0.77** | **1.76** | **1.09** | 0.67 | 0.25 | 0.47 | 0.64 | **4.14** | 0.50 | **2.25** | 0.23 | 0.29 | 0.24 | **0.28** | 7 |
| | >5% MAF in Africans unadmixed with non-Africans | 2,045,769 | 4,345,234 | 0.17 | 0.18 | 0.19 | 0.18 | 0.17 | 0.18 | 0.18 | 0.22 | 0.20 | 0.23 | 0.09 | 0.08 | 0.27 | 0.25 | 0.62 | 0.74 | 0.30 | 0.35 | 0.63 | 0.93 | **3.82** | 0.32 | 1.11 | 0.43 | 0.61 | 0.18 | 0.15 | 1 |
| | >5% MAF in all Africans | 2,109,808 | 4,195,368 | 0.14 | 0.16 | 0.15 | 0.17 | 0.14 | 0.18 | 0.16 | 0.23 | 0.17 | 0.19 | 0.07 | 0.06 | 0.34 | 0.34 | 0.71 | 0.48 | 0.37 | 0.35 | 0.64 | 0.87 | **3.59** | 0.28 | **1.24** | 0.44 | 0.57 | 0.19 | 0.16 | 2 |
| | >5% MAF in Native Americans | 1,513,207 | 2,434,343 | 0.34 | 0.24 | 0.31 | 0.39 | 0.38 | **0.56** | 0.38 | 0.32 | 0.44 | 0.26 | 0.35 | 0.30 | **1.04** | **0.90** | **1.94** | **1.15** | **0.77** | 0.55 | 0.66 | 0.92 | **5.55** | 0.76 | **2.33** | 0.58 | 0.52 | **0.39** | **0.32** | 10 |
| | >5% MAF in Central Asians and Siberians | 1,843,262 | 2,972,941 | 0.18 | 0.11 | 0.17 | 0.12 | 0.18 | 0.13 | 0.17 | 0.15 | 0.35 | 0.15 | 0.22 | 0.14 | **0.95** | **0.78** | **1.92** | **1.07** | **0.74** | 0.41 | 0.49 | 0.85 | **4.95** | 0.73 | **2.51** | 0.44 | 0.37 | **0.29** | **0.30** | 9 |
| | >5% MAF in East Asians | 1,723,831 | 2,841,709 | 0.26 | 0.06 | 0.26 | 0.19 | 0.24 | 0.21 | 0.23 | 0.38 | 0.32 | 0.26 | 0.28 | 0.11 | **1.01** | **0.81** | **2.00** | **1.15** | **0.78** | 0.50 | 0.65 | 0.93 | **5.95** | 0.73 | **2.53** | 0.48 | 0.46 | **0.37** | **0.37** | 9 |
| | >5% MAF in Europeans | 1,885,336 | 2,899,015 | 0.14 | 0.22 | 0.06 | 0.22 | 0.11 | 0.21 | 0.16 | 0.18 | 0.42 | 0.24 | 0.19 | 0.19 | **0.79** | **0.76** | **1.69** | 0.95 | **0.71** | 0.39 | 0.49 | 0.82 | **3.11** | 0.62 | **2.30** | 0.47 | 0.39 | **0.32** | **0.31** | 8 |
| | >5% MAF in Middle Eastern groups | 2,018,884 | 3,018,442 | 0.27 | 0.22 | 0.13 | 0.20 | 0.14 | 0.20 | 0.17 | 0.22 | 0.42 | 0.23 | 0.15 | 0.19 | **0.76** | **0.73** | **1.69** | 0.96 | **0.71** | **0.83** | 0.64 | 0.82 | **3.05** | 0.54 | **2.31** | **0.95** | **0.96** | **0.36** | **0.29** | 11 |
| | >5% MAF in Papuans and Aboriginal Australians | 1,515,022 | 2,367,290 | 0.39 | 0.34 | 0.38 | 0.33 | 0.40 | 0.37 | 0.35 | **0.60** | 0.41 | 0.33 | 0.39 | 0.33 | **1.20** | **1.21** | **2.38** | **1.79** | **0.86** | 0.51 | **1.07** | 0.58 | **8.13** | 0.65 | **2.85** | 0.44 | 0.57 | **0.30** | **0.30** | 11 |
| | >5% MAF in South Asians | 1,908,459 | 3,008,027 | 0.14 | 0.16 | 0.13 | 0.16 | 0.11 | 0.16 | 0.15 | 0.20 | 0.31 | 0.17 | 0.16 | 0.13 | **0.86** | **0.75** | **1.81** | 0.99 | **0.71** | 0.34 | 0.45 | 0.69 | **4.41** | 0.60 | **2.31** | 0.37 | 0.36 | **0.28** | **0.27** | 8 |
| AT/GC mutation types + MAF ascertainment | global, >5% MAF | 323,296 | 508,728 | 0.37 | 0.34 | 0.35 | 0.34 | 0.33 | 0.34 | 0.35 | 0.33 | 0.41 | 0.35 | 0.50 | 0.43 | **0.86** | **0.81** | **1.68** | **1.42** | **0.78** | 0.54 | 0.65 | 0.51 | **3.82** | 0.70 | **1.20** | 0.38 | 0.66 | **0.27** | 0.25 | 8 |
| | >5% MAF in Africans unadmixed with non-Africans | 309,172 | 653,567 | 0.41 | 0.37 | 0.39 | 0.39 | 0.37 | 0.35 | 0.39 | 0.37 | 0.43 | 0.39 | 0.50 | 0.43 | 0.50 | 0.45 | 0.75 | 0.70 | 0.54 | 0.57 | 0.65 | 0.53 | **3.52** | 0.60 | 0.74 | 0.52 | 0.74 | 0.24 | 0.22 | 1 |
| | >5% MAF in all Africans | 319,053 | 630,818 | 0.40 | 0.35 | 0.38 | 0.38 | 0.36 | 0.35 | 0.37 | 0.37 | 0.44 | 0.37 | 0.50 | 0.43 | 0.54 | 0.50 | 0.84 | 0.73 | 0.58 | 0.59 | 0.65 | 0.54 | **3.38** | 0.60 | 0.79 | 0.52 | 0.73 | 0.25 | 0.22 | 1 |
| | >5% MAF in Native Americans | 229,939 | 366,407 | 0.48 | 0.41 | **0.48** | **0.49** | **0.49** | **0.60** | **0.48** | 0.44 | 0.58 | 0.42 | 0.59 | 0.54 | **1.01** | **0.90** | **1.77** | **1.55** | **0.89** | 0.66 | 0.84 | 0.71 | **4.79** | 0.91 | 1.12 | 0.60 | 0.74 | **0.41** | **0.31** | 13 |
| | >5% MAF in Central Asians and Siberians | 280,103 | 447,394 | 0.40 | 0.34 | 0.39 | 0.35 | 0.35 | 0.34 | 0.37 | 0.35 | 0.52 | 0.36 | 0.54 | 0.45 | **0.93** | **0.80** | **1.75** | **1.49** | **0.84** | 0.58 | 0.72 | 0.86 | **4.54** | 0.89 | **1.18** | 0.48 | 0.67 | **0.32** | **0.27** | 9 |
| | >5% MAF in East Asians | 261,857 | 427,465 | 0.44 | 0.33 | 0.43 | 0.40 | 0.39 | 0.38 | 0.40 | **0.52** | 0.49 | 0.41 | 0.55 | 0.44 | **0.97** | **0.83** | **1.79** | **1.56** | **0.88** | 0.63 | 0.82 | 0.73 | **5.23** | **0.98** | **1.24** | 0.50 | 0.71 | **0.36** | **0.32** | 11 |
| | >5% MAF in Europeans | 285,723 | 435,343 | 0.39 | 0.41 | 0.35 | 0.38 | 0.34 | 0.37 | 0.37 | 0.36 | 0.56 | 0.42 | 0.52 | 0.47 | **0.86** | **0.81** | **1.62** | **1.36** | **0.82** | 0.58 | 0.77 | 0.71 | **3.07** | 0.85 | 1.10 | 0.52 | 0.75 | **0.35** | **0.27** | 8 |
| | >5% MAF in Middle Eastern groups | 306,450 | 453,707 | 0.45 | 0.40 | 0.37 | 0.38 | 0.36 | 0.37 | 0.37 | 0.38 | 0.59 | 0.40 | 0.52 | 0.47 | **0.86** | **0.80** | **1.64** | **1.32** | **0.82** | **0.91** | 0.83 | 0.79 | **3.02** | 0.80 | 1.10 | **0.92** | **0.99** | **0.38** | 0.25 | 10 |
| | >5% MAF in Papuans and Aboriginal Australians | 230,124 | 355,386 | 0.50 | 0.48 | **0.52** | **0.48** | **0.54** | 0.45 | **0.48** | **0.69** | 0.50 | 0.50 | 0.62 | 0.53 | **1.12** | **1.14** | **2.10** | **2.12** | **0.92** | 0.71 | **1.18** | 0.76 | **7.18** | 0.85 | **1.49** | 0.51 | 0.84 | **0.32** | **0.31** | 15 |
| | >5% MAF in South Asians | 289,739 | 452,200 | 0.38 | 0.37 | 0.37 | 0.36 | 0.33 | 0.34 | 0.37 | 0.36 | 0.50 | 0.39 | 0.52 | 0.45 | **0.90** | **0.78** | **1.69** | **1.43** | **0.83** | 0.55 | 0.68 | 0.54 | **4.07** | 0.86 | 1.10 | 0.43 | 0.67 | **0.29** | 0.24 | 7 |
| * the SNP counts correspond to sites polymorphic in larger collections of groups | | number of biased asc. => | | 8 | 10 | 15 | 15 | 14 | 14 | 15 | 16 | 8 | 10 | 8 | 9 | 30 | 31 | 31 | 27 | 28 | 14 | 3 | 4 | 35 | 6 | 24 | 13 | 5 | 23 | 23 | |

** only statistics of the form f4(African X, archaic; African Y, non-African) were considered within this class

**Suppl. Table 7.** Performance of ascertainment schemes explored across 27 exhaustively sampled $f_4$-statistic classes and assessed as estimated standard deviation of residuals from a linear model for $f_4$-statistic Z-scores ("residual SE"). Only $f_4$-statistics having absolute Z-scores < 15 SE (on all sites) were considered. We also applied the binary classifier to determine if the ascertainment produces unbiased or biased results (the latter highlighted in bold and underlined text). The numbers of $f_4$-statistic classes or ascertainment schemes affected by bias are shown in the rightmost column and in the bottom row, respectively. The composition of the $f_4$-statistic classes is shown above the table in an abbreviated way. For each $f_4$-statistic class, total number of distinct statistics and the number of statistics having absolute Z-scores < 15 SE (on all sites) are shown.

***R2* of a linear trend for *f4*-statistics (|Z|< 15 SE on all sites)**

| ascertainment type | further details on the ascertainment | min. size of the SNP panel* | max. size of the SNP panel* | AFR 0, ARCH 0, **ME 4** | AFR 0, **EAS 4** | AMRSIB 0, **EUR 4,** PAP 0 | **AMRSIB 4,** EUR 0, PAP 0 | AMRSIB 0, **EUR X, PAP Y** | **AMRSIB X,** EUR 0, **PAP Y** | **AMRSIB X, EUR Y,** PAP 0 | **AMRSIB X, EUR Y, PAP Z** |
|---|---|---|---|---|---|---|---|---|---|---|---|
| | | | group: | non-African 4 | | | | | | | |
| | | | no. f4-stat. | 37,950 | 26,565 | 71,253 | 11,628 | 36,627 | 10,317 | 500,859 | 119,016 |
| | | | no. f4-stat., \|Z\|<15 SE | 37,950 | 26,319 | 71,238 | 8,946 | 34,191 | 6,755 | 331,307 | 51,837 |
| A<>T and G<>C mutations | | 805,042 | 3,610,461 | 0.98 | 1.00 | 0.99 | 1.00 | 0.99 | 1.00 | 0.99 | 1.00 |
| 1240K panel | | 501,429 | 763,188 | 0.97 | 0.99 | 0.99 | 1.00 | 0.98 | 1.00 | 0.99 | 1.00 |
| components of the 1240K panel | Illumina610-Quad sites | 256,277 | 354,539 | 0.96 | 0.99 | **0.98** | **0.99** | **0.97** | **0.99** | **0.99** | **0.99** |
| | sites exclusive to Illumina610-Quad | 183,680 | 255,216 | 0.95 | 0.98 | **0.97** | **0.99** | **0.96** | **0.99** | **0.98** | **0.99** |
| | sites included in both Illumina610-Quad and HumanOrigins | 72,597 | 99,323 | **0.87** | **0.97** | **0.95** | **0.98** | **0.92** | **0.98** | **0.96** | **0.98** |
| | HumanOrigins sites | 244,922 | 399,064 | 0.94 | 0.99 | **0.97** | **0.99** | **0.97** | **0.99** | **0.98** | **0.99** |
| | sites exclusive to HumanOrigins | 171,249 | 299,741 | **0.92** | **0.98** | **0.96** | **0.99** | **0.96** | **0.99** | **0.98** | **0.99** |
| | 1240K, other sites | 67,096 | 108,908 | **0.84** | **0.96** | **0.92** | **0.98** | **0.90** | **0.98** | **0.94** | **0.98** |
| the largest panels included in the HumanOrigins array | panel 13 based on a San individual and Denisovan | 67,557 | 90,136 | **0.76** | **0.94** | **0.89** | **0.97** | **0.89** | **0.97** | **0.93** | **0.97** |
| | panel 4 based on a San individual | 52,862 | 106,537 | **0.81** | **0.94** | **0.88** | **0.97** | **0.85** | **0.97** | **0.93** | **0.97** |
| | panel 5 based on a Yoruba individual | 44,674 | 82,895 | **0.75** | **0.94** | **0.86** | **0.97** | **0.84** | **0.97** | **0.92** | **0.97** |
| | panels 4 and 5 | 46,701 | 177,443 | **0.86** | **0.97** | **0.93** | **0.98** | **0.92** | **0.98** | **0.96** | **0.98** |
| other enrichment panels | 1000K: transversions in 2 Yoruba ind. and in Altai Neand. | 364,079 | 610,754 | 0.95 | 0.98 | **0.98** | **0.99** | **0.97** | **0.99** | **0.98** | **0.99** |
| | 2200K = 1000K + 1240K | 814,915 | 1,304,074 | 0.99 | 1.00 | 0.99 | 1.00 | 0.99 | 1.00 | 0.99 | 1.00 |
| archaic ascertainment | transitions and transversions | 525,014 | 1,555,781 | **0.83** | **0.95** | **0.90** | **0.97** | **0.89** | **0.98** | **0.95** | **0.98** |
| | transversions | 165,249 | 484,675 | **0.80** | **0.94** | **0.89** | **0.97** | **0.87** | **0.97** | **0.93** | **0.97** |
| MAF ascertainment | global, >>5% MAF | 2,129,201 | 3,387,302 | 1.00 | 1.00 | 1.00 | 1.00 | 1.00 | 1.00 | 1.00 | 1.00 |
| | >5% MAF in Africans unadmixed with non-Africans | 2,045,769 | 4,345,234 | 1.00 | 1.00 | 1.00 | 1.00 | 1.00 | 1.00 | 1.00 | 1.00 |
| | >5% MAF in all Africans | 2,109,808 | 4,195,368 | 1.00 | 1.00 | 1.00 | 1.00 | 1.00 | 1.00 | 1.00 | 1.00 |
| | >5% MAF in Native Americans | 1,513,207 | 2,434,343 | 0.98 | 1.00 | 0.99 | **0.99** | 0.98 | **0.99** | **0.99** | 1.00 |
| | >5% MAF in Central Asians and Siberians | 1,843,262 | 2,972,941 | 0.99 | 1.00 | 1.00 | 1.00 | 1.00 | 1.00 | 1.00 | 1.00 |
| | >5% MAF in East Asians | 1,723,831 | 2,841,709 | 0.99 | 1.00 | 0.99 | 1.00 | 0.99 | 1.00 | 1.00 | **0.99** |
| | >5% MAF in Europeans | 1,885,336 | 2,899,015 | 1.00 | 1.00 | 1.00 | 1.00 | 1.00 | 1.00 | 1.00 | 1.00 |
| | >5% MAF in Middle Eastern groups | 2,018,884 | 3,018,442 | 0.99 | 1.00 | 1.00 | 1.00 | 1.00 | 1.00 | 1.00 | 1.00 |
| | >5% MAF in Papuans and Aboriginal Australians | 1,515,022 | 2,367,290 | 0.97 | 0.99 | 0.99 | 1.00 | 0.98 | 1.00 | 0.99 | **0.99** |
| | >5% MAF in South Asians | 1,908,459 | 3,008,027 | 1.00 | 1.00 | 1.00 | 1.00 | 1.00 | 1.00 | 1.00 | 1.00 |
| AT/GC mutation types + MAF ascertainment | global, >5% MAF | 323,296 | 508,728 | 0.98 | 0.99 | 0.99 | 1.00 | 0.99 | 1.00 | 0.99 | 1.00 |
| | >5% MAF in Africans unadmixed with non-Africans | 309,172 | 653,567 | 0.97 | 0.99 | 0.98 | 1.00 | 0.98 | 1.00 | 0.99 | 1.00 |
| | >5% MAF in all Africans | 319,053 | 630,818 | 0.97 | 0.99 | 0.99 | 1.00 | 0.98 | 1.00 | 0.99 | 1.00 |
| | >5% MAF in Native Americans | 229,939 | 366,407 | 0.95 | 0.99 | **0.98** | **0.99** | **0.97** | **0.99** | **0.98** | **0.99** |
| | >5% MAF in Central Asians and Siberians | 280,103 | 447,394 | 0.97 | 0.99 | 0.99 | 1.00 | 0.98 | 1.00 | 0.99 | 1.00 |
| | >5% MAF in East Asians | 261,857 | 427,465 | 0.96 | 0.99 | 0.98 | 1.00 | 0.98 | 1.00 | 0.99 | **0.99** |
| | >5% MAF in Europeans | 285,723 | 435,343 | 0.97 | 0.99 | 0.99 | 1.00 | 0.99 | 1.00 | 0.99 | 1.00 |
| | >5% MAF in Middle Eastern groups | 306,450 | 453,707 | 0.96 | 0.99 | 0.99 | 1.00 | 0.98 | 1.00 | 0.99 | 1.00 |
| | >5% MAF in Papuans and Aboriginal Australians | 230,124 | 355,386 | 0.95 | 0.98 | **0.97** | **0.99** | **0.96** | **0.99** | **0.99** | **0.98** |
| | >5% MAF in South Asians | 289,739 | 452,200 | 0.97 | 0.99 | 0.99 | 1.00 | 0.99 | 1.00 | 0.99 | 1.00 |
| * the SNP counts correspond to sites polymorphic in larger collections of groups | | | **number of biased asc. =>** | 9 | 9 | 15 | 16 | 15 | 16 | 16 | 18 |

| ascertainment type | further details on the ascertainment | **AFR 1,** ARCH 0, **ME 3** | **AFR 1, EAS 3** | **AFR 2,** ARCH 0, **ME 2** | **AFR 2, EAS 2** | **AFR 3,** ARCH 0, **ME 1** | **AFR 3, EAS 1** | **AFR 3, ARCH 1,** ME 0 | **AFR 3, Chimp 1** | **AFR 4,** ARCH 0, ME 0 |
|---|---|---|---|---|---|---|---|---|---|---|
| | group: | Afr. 1, non-Afr. 3 | | Afr. 2, non-Afr. 2 | | African 3, X | | | | Afr. 4 |
| | no. f4-stat. | 213,900 | 164,703 | 418,500 | 352,935 | 337,125 | 310,155 | 40,455 | 39,732 | 94,395 |
| | no. f4-stat., \|Z\|<15 SE | 213,474 | 163,658 | 139,500 | 117,645 | 245,619 | 243,638 | 26,540 | 24,165 | 77,840 |
| A<>T and G<>C mutations | | 0.99 | 1.00 | 0.78 | 0.83 | 0.99 | 1.00 | 0.99 | 0.99 | 0.99 |
| 1240K panel | | 0.99 | 0.99 | 0.78 | 0.79 | **0.96** | **0.97** | **0.87** | **0.90** | **0.97** |
| components of the 1240K panel | Illumina610-Quad sites | 0.99 | 0.99 | 0.60 | 0.61 | **0.95** | **0.96** | **0.81** | **0.87** | **0.95** |
| | sites exclusive to Illumina610-Quad | 0.98 | **0.98** | 0.52 | **0.51** | **0.94** | **0.95** | **0.81** | **0.87** | **0.94** |
| | sites included in both Illumina610-Quad and HumanOrigins | **0.96** | **0.97** | **0.31** | **0.34** | **0.92** | **0.95** | **0.76** | **0.83** | **0.90** |
| | HumanOrigins sites | 0.98 | 0.99 | 0.63 | 0.65 | **0.94** | **0.95** | **0.87** | **0.89** | **0.95** |
| | sites exclusive to HumanOrigins | 0.97 | **0.98** | 0.56 | 0.58 | **0.93** | **0.94** | **0.89** | **0.89** | **0.95** |
| | 1240K, other sites | **0.95** | **0.96** | **0.34** | **0.33** | **0.95** | **0.95** | **0.87** | **0.89** | **0.92** |
| the largest panels included in the HumanOrigins array | panel 13 based on a San individual and Denisovan | **0.92** | **0.95** | **0.28** | **0.26** | **0.89** | **0.92** | **0.54** | **0.61** | **0.90** |
| | panel 4 based on a San individual | **0.93** | **0.94** | **0.32** | **0.23** | **0.88** | **0.92** | **0.90** | **0.86** | **0.88** |
| | panel 5 based on a Yoruba individual | **0.91** | **0.94** | **0.27** | **0.22** | **0.94** | **0.94** | **0.84** | **0.89** | **0.88** |
| | panels 4 and 5 | **0.95** | **0.96** | **0.44** | **0.34** | **0.95** | **0.96** | **0.92** | 0.93 | **0.93** |
| other enrichment panels | 1000K: transversions in 2 Yoruba ind. and in Altai Neand. | 0.98 | 0.99 | 0.70 | 0.68 | 0.98 | **0.98** | **0.93** | 0.96 | **0.97** |
| | 2200K = 1000K + 1240K | 1.00 | 1.00 | 0.87 | 0.85 | 0.98 | **0.98** | **0.91** | 0.94 | 0.98 |
| archaic ascertainment | transitions and transversions | **0.93** | **0.95** | **0.40** | **0.38** | **0.94** | **0.94** | **0.76** | 0.94 | **0.94** |
| | transversions | **0.91** | **0.94** | **0.33** | **0.34** | **0.92** | **0.93** | **0.76** | **0.87** | **0.91** |
| MAF ascertainment | global, >>5% MAF | 1.00 | 1.00 | 0.99 | 1.00 | **0.98** | **0.98** | **0.90** | 0.97 | 0.97 |
| | >5% MAF in Africans unadmixed with non-Africans | 1.00 | 1.00 | 0.99 | 0.99 | 1.00 | 1.00 | 0.99 | 0.99 | 0.99 |
| | >5% MAF in all Africans | 1.00 | 1.00 | 1.00 | 1.00 | 1.00 | 1.00 | 0.98 | 0.99 | 0.99 |
| | >5% MAF in Native Americans | 0.99 | 1.00 | 0.88 | 0.91 | **0.96** | **0.97** | **0.86** | 0.96 | **0.96** |
| | >5% MAF in Central Asians and Siberians | 0.99 | 1.00 | 0.95 | 0.98 | **0.96** | **0.98** | **0.87** | 0.97 | **0.97** |
| | >5% MAF in East Asians | 0.99 | 1.00 | 0.92 | 0.99 | **0.96** | **0.98** | **0.86** | 0.96 | **0.96** |
| | >5% MAF in Europeans | 0.99 | 1.00 | 0.97 | 0.96 | **0.97** | **0.98** | **0.90** | 0.97 | **0.97** |
| | >5% MAF in Middle Eastern groups | 0.99 | 1.00 | 0.98 | 0.96 | **0.98** | **0.98** | **0.90** | 0.97 | **0.97** |
| | >5% MAF in Papuans and Aboriginal Australians | 0.99 | 1.00 | 0.85 | 0.88 | **0.94** | **0.95** | **0.80** | **0.90** | **0.95** |
| | >5% MAF in South Asians | 0.99 | 1.00 | 0.98 | 0.98 | **0.97** | **0.98** | **0.89** | 0.97 | **0.97** |
| AT/GC mutation types + MAF ascertainment | global, >5% MAF | 0.99 | 1.00 | 0.77 | 0.82 | **0.97** | **0.98** | **0.88** | 0.93 | **0.96** |
| | >5% MAF in Africans unadmixed with non-Africans | 0.99 | 0.99 | 0.77 | 0.82 | 0.99 | 0.99 | 0.98 | 0.99 | 0.98 |
| | >5% MAF in all Africans | 0.99 | 0.99 | 0.77 | 0.82 | 0.99 | 0.99 | 0.97 | 0.98 | 0.98 |
| | >5% MAF in Native Americans | 0.98 | 0.99 | 0.68 | 0.73 | **0.95** | **0.96** | **0.84** | **0.90** | **0.94** |
| | >5% MAF in Central Asians and Siberians | 0.98 | 0.99 | 0.73 | 0.80 | **0.96** | **0.97** | **0.85** | **0.92** | **0.95** |
| | >5% MAF in East Asians | 0.98 | 0.99 | 0.71 | 0.81 | **0.96** | **0.97** | **0.84** | **0.90** | **0.94** |
| | >5% MAF in Europeans | 0.98 | 0.99 | 0.75 | 0.78 | **0.97** | **0.97** | **0.88** | 0.93 | **0.95** |
| | >5% MAF in Middle Eastern groups | 0.98 | 0.99 | 0.76 | 0.79 | **0.97** | **0.97** | **0.88** | 0.93 | **0.95** |
| | >5% MAF in Papuans and Aboriginal Australians | 0.98 | 0.99 | 0.64 | 0.72 | **0.94** | **0.94** | **0.78** | **0.83** | **0.93** |
| | >5% MAF in South Asians | 0.98 | 0.99 | 0.75 | 0.80 | **0.96** | **0.98** | **0.87** | **0.92** | **0.95** |
| | **number of biased asc. =>** | 8 | 10 | 8 | 9 | 29 | 31 | 31 | 17 | 29 |

| ascertainment type | further details on the ascertainment | **AFR 1, ARCH 1, ME 2** | **AFR 1, ARCH 2, ME 1** | **AFR 1, ARCH 2, Chimp 1** | **AFR 2, ARCH 1, ME 1 **** | **AFR 2, ARCH 2,** ME 0 | **AFR 2, ARCH 1, Chimp 1** | AFR 0, **ARCH 1, ME 3** | AFR 0, **ARCH 2, ME 2** | AFR 0, **ARCH 3, ME 1** | **AFR 1, ARCH 3,** ME 0 | number of biased f4-stat. classes |
|---|---|---|---|---|---|---|---|---|---|---|---|---|
| | group: | African 1-2, archaic 1-3, X | | | | | | arch. 1-2, non-Afr. 2-3 | | arch. 3, non-Afr. 1 | | |
| | no. f4-stat. | 83,700 | 6,975 | 396 | 69,750 | 4,185 | 8,514 | 20,700 | 2,700 | 75 | 93 | |
| | no. f4-stat., \|Z\|<15 SE | 27,900 | 2,325 | 127 | 59,240 | 1,395 | 2,838 | 20,699 | 900 | 25 | 31 | |
| A<>T and G<>C mutations | | 0.91 | 1.00 | 1.00 | 1.00 | 0.62 | 0.96 | 0.99 | 0.85 | 1.00 | 1.00 | 0 |
| 1240K panel | | 0.69 | 0.99 | 1.00 | **0.36** | 0.45 | **0.40** | 0.97 | 0.77 | **0.99** | **0.97** | 9 |
| components of the 1240K panel | Illumina610-Quad sites | **0.53** | **0.98** | **0.97** | **0.23** | 0.20 | **0.00** | 0.96 | 0.60 | **0.97** | **0.90** | 18 |
| | sites exclusive to Illumina610-Quad | **0.33** | **0.97** | **0.03** | **0.25** | 0.30 | **0.05** | 0.95 | 0.49 | **0.96** | **0.85** | 20 |
| | sites included in both Illumina610-Quad and HumanOrigins | **0.53** | **0.91** | 0.98 | **0.18** | **0.01** | **0.04** | **0.92** | 0.46 | **0.96** | **0.85** | 26 |
| | HumanOrigins sites | **0.57** | **0.98** | 1.00 | **0.41** | 0.29 | **0.38** | 0.96 | 0.55 | **0.99** | **0.97** | 17 |
| | sites exclusive to HumanOrigins | **0.44** | **0.97** | 1.00 | **0.55** | 0.43 | **0.56** | 0.94 | **0.35** | **0.99** | **0.96** | 21 |
| | 1240K, other sites | **0.30** | **0.89** | 1.00 | **0.56** | **0.00** | **0.56** | **0.89** | 0.46 | **0.97** | **0.97** | 25 |
| the largest panels included in the HumanOrigins array | panel 13 based on a San individual and Denisovan | **0.40** | **0.57** | 1.00 | **0.47** | **0.01** | **0.08** | **0.70** | **0.23** | **0.99** | **0.97** | 26 |
| | panel 4 based on a San individual | **0.19** | **0.85** | **0.98** | **0.56** | 0.25 | **0.62** | **0.80** | **0.18** | **0.46** | **0.39** | 26 |
| | panel 5 based on a Yoruba individual | **0.30** | **0.87** | **0.91** | **0.76** | 0.17 | **0.44** | **0.83** | **0.25** | **0.98** | **0.96** | 26 |
| | panels 4 and 5 | **0.32** | **0.93** | **0.98** | **0.70** | 0.39 | 0.73 | **0.88** | **0.39** | **0.95** | **0.86** | 24 |
| other enrichment panels | 1000K: transversions in 2 Yoruba ind. and in Altai Neand. | **0.62** | **0.96** | **0.98** | **0.72** | 0.48 | **0.62** | 0.96 | 0.46 | 1.00 | 1.00 | 14 |
| | 2200K = 1000K + 1240K | 0.78 | 0.99 | 1.00 | **0.54** | 0.62 | **0.57** | 0.98 | 0.77 | 1.00 | 1.00 | 4 |
| archaic ascertainment | transitions and transversions | 0.71 | 1.00 | 1.00 | **0.43** | 1.00 | 0.85 | **0.60** | 1.00 | 1.00 | 1.00 | 18 |
| | transversions | 0.66 | 1.00 | 1.00 | **0.43** | 0.87 | 0.84 | **0.57** | 0.92 | 1.00 | 1.00 | 19 |
| MAF ascertainment | global, >>5% MAF | 0.96 | 0.99 | **0.95** | **0.64** | 0.77 | **0.08** | 1.00 | 0.95 | 0.99 | **0.98** | 7 |
| | >5% MAF in Africans unadmixed with non-Africans | 0.90 | 0.98 | 1.00 | **0.73** | 0.91 | 0.76 | 0.99 | 0.71 | 1.00 | 1.00 | 1 |
| | >5% MAF in all Africans | 0.90 | 0.99 | 1.00 | **0.75** | 0.93 | 0.67 | 0.99 | 0.74 | 1.00 | 1.00 | 1 |
| | >5% MAF in Native Americans | 0.76 | **0.98** | **0.74** | **0.37** | 0.34 | **0.07** | 0.97 | 0.84 | **0.97** | **0.96** | 13 |
| | >5% MAF in Central Asians and Siberians | 0.90 | 0.99 | **0.65** | **0.48** | 0.47 | **0.02** | 0.98 | 0.89 | **0.99** | **0.95** | 9 |
| | >5% MAF in East Asians | 0.79 | **0.98** | **0.07** | **0.34** | 0.42 | **0.07** | 0.98 | 0.82 | **0.93** | **0.81** | 11 |
| | >5% MAF in Europeans | 0.94 | 0.99 | **0.89** | **0.76** | 0.64 | **0.01** | 0.97 | 0.92 | **0.99** | **0.96** | 9 |
| | >5% MAF in Middle Eastern groups | **0.43** | 0.99 | **0.93** | **0.78** | 0.76 | **0.03** | 0.95 | **0.36** | **0.98** | **0.95** | 11 |
| | >5% MAF in Papuans and Aboriginal Australians | 0.77 | 0.98 | 1.00 | **0.14** | 0.73 | **0.12** | 0.98 | 0.76 | **0.97** | **0.95** | 10 |
| | >5% MAF in South Asians | 0.92 | 0.99 | **0.90** | **0.56** | 0.58 | **0.00** | 0.99 | 0.91 | **0.99** | **0.98** | 9 |
| AT/GC mutation types + MAF ascertainment | global, >5% MAF | 0.80 | 0.98 | **0.93** | **0.62** | 0.25 | **0.54** | 0.99 | 0.75 | **0.99** | 0.98 | 8 |
| | >5% MAF in Africans unadmixed with non-Africans | 0.71 | **0.96** | 1.00 | **0.71** | 0.42 | 0.81 | 0.98 | 0.60 | 1.00 | 1.00 | 2 |
| | >5% MAF in all Africans | 0.70 | **0.97** | 0.99 | **0.73** | 0.41 | 0.80 | 0.98 | 0.59 | 1.00 | 1.00 | 2 |
| | >5% MAF in Native Americans | **0.61** | **0.94** | **0.44** | **0.37** | **0.05** | **0.27** | 0.96 | 0.70 | **0.98** | 0.98 | 18 |
| | >5% MAF in Central Asians and Siberians | 0.78 | **0.97** | **0.03** | **0.45** | **0.11** | **0.33** | 0.98 | 0.72 | **0.99** | **0.98** | 12 |
| | >5% MAF in East Asians | 0.67 | **0.94** | **0.06** | **0.34** | **0.05** | **0.18** | 0.98 | 0.68 | **0.96** | **0.94** | 13 |
| | >5% MAF in Europeans | 0.84 | 0.98 | **0.83** | **0.71** | 0.20 | **0.61** | 0.96 | 0.74 | **0.99** | 0.98 | 8 |
| | >5% MAF in Middle Eastern groups | **0.32** | **0.98** | **0.88** | **0.72** | 0.33 | **0.63** | 0.94 | **0.32** | **0.98** | **0.98** | 12 |
| | >5% MAF in Papuans and Aboriginal Australians | **0.59** | **0.95** | 0.99 | **0.13** | 0.34 | **0.00** | 0.97 | 0.57 | **0.94** | **0.94** | 17 |
| | >5% MAF in South Asians | 0.78 | **0.97** | **0.68** | **0.53** | **0.10** | **0.47** | 0.98 | 0.72 | **0.99** | 0.98 | 11 |
| | **number of biased asc. =>** | 15 | 21 | 21 | 35 | 7 | 28 | 8 | 7 | 26 | 23 | |

* the SNP counts correspond to sites polymorphic in larger collections of groups

** only statistics of the form f4(African X, archaic; African Y, non-African) were considered within this class

**Suppl. Table 8.** Performance of ascertainment schemes explored across 27 exhaustively sampled $f_4$-statistic classes and assessed as $R^2$ of a linear model for $f_4$-statistics having absolute Z-scores < 15 SE (on all sites). We also applied the binary classifier to determine if the ascertainment produces unbiased or biased results (the latter highlighted in bold and underlined text). The numbers of $f_4$-statistic classes or ascertainment schemes affected by bias are shown in the rightmost column and in the bottom row, respectively. The composition of the $f_4$-statistic classes is shown above the table in an abbreviated way. For each $f_4$-statistic class, total number of distinct statistics and the number of statistics having absolute Z-scores < 15 SE (on all sites) are shown.

| | *f4*-statistic | Altai<br>Biaka<br>Mbuti<br>Saharawi | | Fulani<br>Ju\|'hoan North<br>Igbo<br>Ogiek | | Burmese<br>Dinka<br>Ju\|'hoan North<br>Sengwer | |
|---|---|---|---|---|---|---|---|
| **SNP set** | **ascertainment type** | ***f4*** | **Z** | ***f4*** | **Z** | ***f4*** | **Z** |
| all sites | all sites | 2.4E-05 | 0.81 | -5.6E-05 | -2.44 | -4.5E-05 | -1.88 |
| **AT GC** | AT GC | 3.6E-05 | 1.03 | -3.6E-05 | -1.33 | -4.2E-05 | -1.62 |
| thinned 5 | thinned | 4.8E-05 | 1.25 | -6.0E-05 | -2.16 | -2.5E-05 | -0.93 |
| **1240K** | 1240K | 4.3E-03 | 11.16 | -1.9E-03 | -6.14 | -1.9E-03 | -5.56 |
| **global** | MAF | 3.7E-03 | 8.29 | -1.8E-03 | -4.94 | -1.3E-03 | -3.41 |
| **AFR2** | MAF | 2.0E-03 | 5.39 | -1.1E-03 | -3.31 | -7.5E-04 | -2.30 |
| EAS | MAF | 4.5E-03 | 10.30 | -1.9E-03 | -5.12 | -9.0E-04 | -2.29 |
| panel 4 | HO panels | 2.2E-03 | 3.25 | -6.8E-04 | -1.36 | -1.5E-03 | -3.14 |
| **archaic** | archaic asc. | -2.4E-03 | -6.79 | -1.8E-04 | -1.09 | -2.0E-04 | -1.24 |
| archaic, transv. | archaic asc. | -2.5E-03 | -6.36 | -9.6E-05 | -0.54 | -2.4E-04 | -1.37 |

**Suppl. Table 9.** Values of three selected $f_4$-statistics and corresponding Z-scores across 8 ascertainment schemes and on all sites. Five ascertainment schemes featured in Suppl. Tables 10-12 are highlighted in bold.

| ascertainment scheme | meta-population | DAF bin | no. sites | average derived allele frequency in: Altai | Biaka | Mbuti | Saharawi | av. (a-b) | av. (c-d) | av. (a-b) (c-d) = f4 | Z-score | no. sites used for calculating f4 | % sites used for calculating f4 |
|---|---|---|---|---|---|---|---|---|---|---|---|---|---|
| all sites | AFR | <=5% | 38,883,326 | 0.015 | 0.004 | 0.004 | 0.003 | 1.1E-02 | 1.2E-03 | -1.8E-04 | -14.8 | 38,883,326 | 64.4% |
| | | 5-95% | 4,422,900 | 0.186 | 0.285 | 0.282 | 0.292 | -9.8E-02 | -1.1E-02 | 2.1E-03 | 6.0 | 4,422,900 | 7.3% |
| | | >=95% | 17,081,929 | 0.995 | 1.000 | 1.000 | 1.000 | -5.0E-03 | -1.2E-04 | -4.2E-05 | -5.1 | 17,081,929 | 28.3% |
| | EUR | <=5% | 39,946,373 | 0.016 | 0.009 | 0.010 | 0.003 | 7.0E-03 | 6.8E-03 | -4.4E-04 | -38.7 | 39,946,373 | 66.1% |
| | | 5-95% | 3,114,167 | 0.202 | 0.278 | 0.274 | 0.344 | -7.6E-02 | -7.0E-02 | 3.2E-03 | 7.2 | 3,114,167 | 5.2% |
| | | >=95% | 17,327,607 | 0.989 | 0.996 | 0.996 | 0.999 | -6.9E-03 | -3.3E-03 | 5.3E-04 | 26.3 | 17,327,607 | 28.7% |
| AT/GC | AFR | <=5% | 5,237,869 | 0.017 | 0.005 | 0.005 | 0.004 | 1.3E-02 | 1.4E-03 | -2.0E-04 | -14.1 | 5,237,869 | 61.6% |
| | | 5-95% | 666,914 | 0.172 | 0.275 | 0.272 | 0.283 | -1.0E-01 | -1.1E-02 | 2.3E-03 | 5.9 | 666,914 | 7.8% |
| | | >=95% | 2,601,903 | 0.995 | 1.000 | 1.000 | 1.000 | -4.4E-03 | -8.6E-05 | -2.7E-05 | -3.5 | 2,601,903 | 30.6% |
| | EUR | <=5% | 5,401,054 | 0.018 | 0.010 | 0.011 | 0.003 | 7.8E-03 | 7.7E-03 | -5.1E-04 | -34.6 | 5,401,054 | 63.5% |
| | | 5-95% | 469,526 | 0.189 | 0.269 | 0.264 | 0.338 | -8.0E-02 | -7.3E-02 | 3.5E-03 | 7.0 | 469,526 | 5.5% |
| | | >=95% | 2,636,106 | 0.990 | 0.996 | 0.996 | 0.999 | -6.6E-03 | -3.0E-03 | 5.6E-04 | 23.9 | 2,636,106 | 31.0% |
| 1240K | AFR | <=5% | 165,728 | 0.061 | 0.015 | 0.017 | 0.079 | 4.6E-02 | -6.3E-02 | -2.8E-03 | -9.7 | 165,728 | 18.2% |
| | | 5-95% | 725,952 | 0.202 | 0.331 | 0.326 | 0.377 | -1.3E-01 | -5.1E-02 | 6.4E-03 | 14.0 | 725,952 | 79.6% |
| | | >=95% | 20,096 | 0.623 | 0.983 | 0.983 | 0.961 | -3.6E-01 | 2.2E-02 | -7.2E-03 | -8.6 | 20,096 | 2.2% |
| | EUR | <=5% | 198,368 | 0.076 | 0.103 | 0.104 | 0.033 | -2.7E-02 | 7.1E-02 | -7.8E-03 | -27.8 | 198,368 | 21.8% |
| | | 5-95% | 666,343 | 0.195 | 0.307 | 0.302 | 0.382 | -1.1E-01 | -8.0E-02 | 5.9E-03 | 12.7 | 666,343 | 73.1% |
| | | >=95% | 47,065 | 0.520 | 0.806 | 0.799 | 0.951 | -2.9E-01 | -1.5E-01 | 3.5E-02 | 29.6 | 47,065 | 5.2% |
| global MAF 5% | AFR | <=5% | 513,122 | 0.118 | 0.016 | 0.017 | 0.120 | 1.0E-01 | -1.0E-01 | -6.3E-03 | -10.5 | 513,122 | 14.6% |
| | | 5-95% | 2,973,079 | 0.225 | 0.338 | 0.333 | 0.381 | -1.1E-01 | -4.8E-02 | 5.9E-03 | 12.3 | 2,973,079 | 84.4% |
| | | >=95% | 36,944 | 0.585 | 0.977 | 0.973 | 0.894 | -3.9E-01 | 7.9E-02 | -2.6E-02 | -14.4 | 36,944 | 1.0% |
| | EUR | <=5% | 498,801 | 0.167 | 0.173 | 0.173 | 0.063 | -6.2E-03 | 1.1E-01 | -1.1E-02 | -20.4 | 498,801 | 14.2% |
| | | 5-95% | 2,866,222 | 0.208 | 0.297 | 0.293 | 0.366 | -8.9E-02 | -7.3E-02 | 3.9E-03 | 8.1 | 2,866,222 | 81.4% |
| | | >=95% | 158,119 | 0.455 | 0.704 | 0.689 | 0.923 | -2.5E-01 | -2.3E-01 | 4.7E-02 | 29.0 | 158,119 | 4.5% |
| AFR MAF 5% | AFR | <=5% | 53,286 | 0.038 | 0.055 | 0.057 | 0.048 | -1.7E-02 | 9.2E-03 | -6.8E-04 | -2.0 | 53,286 | 1.2% |
| | | 5-95% | 4,413,474 | 0.187 | 0.285 | 0.282 | 0.293 | -9.9E-02 | -1.1E-02 | 2.1E-03 | 6.0 | 4,413,474 | 98.7% |
| | | >=95% | 4,625 | 0.735 | 0.935 | 0.933 | 0.952 | -2.0E-01 | -1.9E-02 | 1.0E-03 | 0.5 | 4,625 | 0.1% |
| | EUR | <=5% | 1,710,056 | 0.064 | 0.128 | 0.130 | 0.032 | -6.4E-02 | 9.7E-02 | -8.7E-03 | -40.3 | 1,710,056 | 38.2% |
| | | 5-95% | 2,476,700 | 0.221 | 0.333 | 0.328 | 0.393 | -1.1E-01 | -6.6E-02 | 6.2E-03 | 12.5 | 2,476,700 | 55.4% |
| | | >=95% | 284,627 | 0.608 | 0.779 | 0.767 | 0.949 | -1.7E-01 | -1.8E-01 | 3.1E-02 | 28.7 | 284,627 | 6.4% |
| archaic | AFR | <=5% | 1,364,490 | 0.343 | 0.002 | 0.003 | 0.006 | 3.4E-01 | -3.1E-03 | -2.7E-03 | -11.0 | 1,364,490 | 66.3% |
| | | 5-95% | 580,958 | 0.499 | 0.431 | 0.430 | 0.435 | 6.8E-02 | -4.5E-03 | -1.7E-03 | -1.9 | 580,958 | 28.2% |
| | | >=95% | 113,316 | 0.598 | 0.990 | 0.988 | 0.988 | -3.9E-01 | 4.0E-04 | -2.3E-03 | -3.6 | 113,316 | 5.5% |
| | EUR | <=5% | 1,421,966 | 0.337 | 0.018 | 0.018 | 0.005 | 3.2E-01 | 1.3E-02 | 2.3E-03 | 15.6 | 1,421,966 | 69.1% |
| | | 5-95% | 463,636 | 0.525 | 0.397 | 0.396 | 0.422 | 1.3E-01 | -2.6E-02 | -2.2E-02 | -18.3 | 463,636 | 22.5% |
| | | >=95% | 173,160 | 0.589 | 0.906 | 0.901 | 0.978 | -3.2E-01 | -7.7E-02 | 1.1E-02 | 15.7 | 173,160 | 8.4% |

**Suppl. Table 10.** Dissecting the statistic $f_4$(Altai Neanderthal, Biaka; Mbuti, Saharawi) belonging to the (archaic, African X; African Y, non-African) class. Sites were stratified by DAF in Africans or DAF in Europeans into three bins: nearly fixed ancestral (DAF <=5%), non-fixed (DAF 5-95%), and nearly fixed derived (DAF >=95%). This was done for all sites, for random ascertainment (AT/GC sites), and for four non-random ascertainment schemes as indicated in the leftmost column. The number and proportion of $f_4$-informative sites falling into each bin are shown. Mean DAF in four populations, mean differences in DAF between populations 1 and 2, populations 3 and 4, mean products of the DAF differences (i.e., $f_4$-statistics) and their Z-scores are shown for these frequency bins.

| ascertainment scheme | meta-population | DAF bin | no. sites | average derived allele frequency in: Fulani | Ju\|'hoan North | Igbo | Ogiek | av. (a-b) | av. (c-d) | av. (a-b) (c-d) = f4 | Z-score | no. sites used for calculating f4 | % sites used for calculating f4 |
|---|---|---|---|---|---|---|---|---|---|---|---|---|---|
| all sites | AFR | <=5% | 51,867,384 | 0.003 | 0.006 | 0.003 | 0.004 | -2.9E-03 | -6.4E-04 | -1.8E-06 | -0.6 | 51,867,384 | 63.1% |
| | | 5-95% | 6,149,286 | 0.290 | 0.268 | 0.293 | 0.288 | 2.3E-02 | 5.1E-03 | -8.5E-04 | -2.8 | 6,149,286 | 7.5% |
| | | >=95% | 24,190,036 | 1.000 | 0.999 | 1.000 | 1.000 | 3.7E-04 | 1.0E-04 | 2.4E-06 | 2.0 | 24,190,036 | 29.4% |
| | EUR | <=5% | 53,357,917 | 0.006 | 0.011 | 0.008 | 0.006 | -5.3E-03 | 1.8E-03 | 7.9E-05 | 8.8 | 53,357,917 | 64.9% |
| | | 5-95% | 4,326,314 | 0.314 | 0.262 | 0.291 | 0.308 | 5.2E-02 | -1.7E-02 | -1.8E-03 | -5.1 | 4,326,314 | 5.3% |
| | | >=95% | 24,522,364 | 0.998 | 0.995 | 0.997 | 0.998 | 2.4E-03 | -7.9E-04 | -6.2E-05 | -5.9 | 24,522,364 | 29.8% |
| AT/GC | AFR | <=5% | 7,258,497 | 0.004 | 0.007 | 0.003 | 0.004 | -3.2E-03 | -6.7E-04 | -2.9E-08 | 0.0 | 7,258,497 | 61.3% |
| | | 5-95% | 921,846 | 0.280 | 0.257 | 0.283 | 0.277 | 2.3E-02 | 5.4E-03 | -8.0E-04 | -2.5 | 921,846 | 7.8% |
| | | >=95% | 3,668,575 | 1.000 | 1.000 | 1.000 | 1.000 | 2.6E-04 | 7.0E-05 | 2.3E-06 | 1.8 | 3,668,575 | 31.0% |
| | EUR | <=5% | 7,487,196 | 0.006 | 0.012 | 0.009 | 0.007 | -5.9E-03 | 2.0E-03 | 9.0E-05 | 8.4 | 7,487,196 | 63.2% |
| | | 5-95% | 647,754 | 0.305 | 0.251 | 0.281 | 0.299 | 5.4E-02 | -1.8E-02 | -1.8E-03 | -4.8 | 647,754 | 5.5% |
| | | >=95% | 3,713,941 | 0.998 | 0.996 | 0.997 | 0.998 | 2.2E-03 | -7.4E-04 | -6.3E-05 | -5.4 | 3,713,941 | 31.3% |
| 1240K | AFR | <=5% | 185,808 | 0.044 | 0.040 | 0.015 | 0.038 | 4.2E-03 | -2.3E-02 | -1.7E-03 | 14.6 | 185,808 | 18.1% |
| | | 5-95% | 816,649 | 0.362 | 0.316 | 0.349 | 0.357 | 4.6E-02 | -7.5E-03 | -2.0E-03 | -5.3 | 816,649 | 79.7% |
| | | >=95% | 22,212 | 0.976 | 0.971 | 0.987 | 0.978 | 5.0E-03 | 9.6E-03 | -3.4E-04 | -2.2 | 22,212 | 2.2% |
| | EUR | <=5% | 221,295 | 0.073 | 0.127 | 0.104 | 0.075 | -5.5E-02 | 2.9E-02 | 5.1E-05 | 0.2 | 221,295 | 21.6% |
| | | 5-95% | 751,296 | 0.351 | 0.291 | 0.324 | 0.343 | 6.0E-02 | -1.9E-02 | -2.2E-03 | -5.9 | 751,296 | 73.3% |
| | | >=95% | 52,078 | 0.888 | 0.782 | 0.831 | 0.875 | 1.1E-01 | -4.5E-02 | -5.3E-03 | -8.0 | 52,078 | 5.1% |
| global MAF 5% | AFR | <=5% | 716,787 | 0.062 | 0.020 | 0.015 | 0.054 | 4.2E-02 | -3.9E-02 | -2.6E-03 | 16.6 | 716,787 | 14.7% |
| | | 5-95% | 4,111,766 | 0.365 | 0.317 | 0.353 | 0.362 | 4.8E-02 | -8.3E-03 | -1.8E-03 | -4.4 | 4,111,766 | 84.3% |
| | | >=95% | 51,056 | 0.942 | 0.965 | 0.981 | 0.944 | -2.3E-02 | 3.7E-02 | -1.1E-03 | -3.1 | 51,056 | 1.0% |
| | EUR | <=5% | 697,286 | 0.127 | 0.163 | 0.174 | 0.139 | -3.6E-02 | 3.5E-02 | 1.4E-04 | 0.3 | 697,286 | 14.3% |
| | | 5-95% | 3,970,679 | 0.335 | 0.280 | 0.311 | 0.329 | 5.5E-02 | -1.8E-02 | -2.0E-03 | -5.1 | 3,970,679 | 81.4% |
| | | >=95% | 211,622 | 0.825 | 0.672 | 0.742 | 0.803 | 1.5E-01 | -6.1E-02 | -7.4E-03 | -7.0 | 211,622 | 4.3% |
| AFR MAF 5% | AFR | <=5% | 74,613 | 0.047 | 0.048 | 0.043 | 0.049 | -1.3E-03 | -5.2E-03 | -4.1E-04 | -2.3 | 74,613 | 1.2% |
| | | 5-95% | 6,132,565 | 0.291 | 0.268 | 0.293 | 0.288 | 2.3E-02 | 5.1E-03 | -8.5E-04 | -2.8 | 6,132,565 | 98.7% |
| | | >=95% | 6,709 | 0.955 | 0.937 | 0.956 | 0.946 | 1.8E-02 | 1.0E-02 | -5.0E-04 | -0.8 | 6,709 | 0.1% |
| | EUR | <=5% | 2,399,542 | 0.079 | 0.121 | 0.123 | 0.081 | -4.3E-02 | 4.1E-02 | 1.0E-03 | 5.5 | 2,399,542 | 38.6% |
| | | 5-95% | 3,426,567 | 0.369 | 0.313 | 0.350 | 0.364 | 5.6E-02 | -1.4E-02 | -1.8E-03 | -4.0 | 3,426,567 | 55.1% |
| | | >=95% | 387,759 | 0.879 | 0.752 | 0.814 | 0.865 | 1.3E-01 | -5.1E-02 | -4.2E-03 | -6.5 | 387,759 | 6.2% |
| archaic | AFR | <=5% | 1,422,921 | 0.004 | 0.004 | 0.002 | 0.003 | -5.5E-04 | -1.4E-03 | -2.3E-06 | -0.2 | 1,422,921 | 66.3% |
| | | 5-95% | 605,944 | 0.434 | 0.429 | 0.433 | 0.432 | 4.7E-03 | 1.1E-03 | -6.2E-04 | -1.1 | 605,944 | 28.2% |
| | | >=95% | 118,551 | 0.990 | 0.982 | 0.993 | 0.991 | 8.0E-03 | 2.8E-03 | 9.2E-05 | 1.5 | 118,551 | 5.5% |
| | EUR | <=5% | 1,482,687 | 0.010 | 0.021 | 0.015 | 0.011 | -1.0E-02 | 3.2E-03 | -5.1E-05 | -1.0 | 1,482,687 | 69.0% |
| | | 5-95% | 483,906 | 0.410 | 0.394 | 0.401 | 0.407 | 1.6E-02 | -5.5E-03 | -1.7E-04 | -0.3 | 483,906 | 22.5% |
| | | >=95% | 180,823 | 0.948 | 0.889 | 0.922 | 0.940 | 5.9E-02 | -1.8E-02 | -1.2E-03 | -3.3 | 180,823 | 8.4% |

**Suppl. Table 11.** Dissecting the statistic $f_4$(Fulani, Ju|'hoan North; Igbo, Ogiek) composed of four African groups. Sites were stratified by DAF in Africans or DAF in Europeans into three bins: nearly fixed ancestral (DAF <=5%), non-fixed (DAF 5-95%), and nearly fixed derived (DAF >=95%). This was done for "all sites", for random ascertainment (AT/GC sites), and for four non-random ascertainment schemes as indicated in the leftmost column The number and proportion of $f_4$-informative sites falling into each bin are shown. Mean DAF in four populations, mean differences in DAF between populations 1 and 2, populations 3 and 4, mean products of the DAF differences (i.e., $f_4$-statistics) and their Z-scores are shown for these frequency bins.

| ascertainment scheme | meta-population | DAF bin | no. sites | Burmese | Dinka | Ju\|'hoan North | Sengwer | av. (a-b) | av. (c-d) | av. (a-b)(c-d) = f4 | Z-score | no. sites used for calculating f4 | % sites used for calculating f4 |
|---|---|---|---|---|---|---|---|---|---|---|---|---|---|
| | | | | average derived allele frequency in: | | | | | | | | | |
| all sites | AFR | <=5% | 51,604,923 | 0.004 | 0.003 | 0.006 | 0.003 | 8.7E-04 | 3.0E-03 | -1.3E-05 | -4.4 | 51,604,923 | 63.1% |
| | | 5-95% | 6,113,283 | 0.287 | 0.294 | 0.268 | 0.291 | -6.6E-03 | -2.3E-02 | -5.6E-04 | -1.7 | 6,113,283 | 7.5% |
| | | >=95% | 24,012,588 | 1.000 | 1.000 | 0.999 | 1.000 | -1.6E-04 | -4.0E-04 | 5.2E-07 | 1.0 | 24,012,588 | 29.4% |
| | EUR | <=5% | 53,087,943 | 0.002 | 0.007 | 0.011 | 0.007 | -4.9E-03 | 4.5E-03 | 8.4E-05 | 10.2 | 53,087,943 | 65.0% |
| | | 5-95% | 4,298,814 | 0.347 | 0.298 | 0.262 | 0.305 | 4.8E-02 | -4.3E-02 | -9.5E-04 | -2.3 | 4,298,814 | 5.3% |
| | | >=95% | 24,344,015 | 0.999 | 0.997 | 0.995 | 0.997 | 2.1E-03 | -2.0E-03 | -1.9E-04 | -19.6 | 24,344,015 | 29.8% |
| AT/GC | AFR | <=5% | 7,220,965 | 0.004 | 0.003 | 0.007 | 0.003 | 9.8E-04 | 3.3E-03 | -1.8E-05 | -4.7 | 7,220,965 | 61.3% |
| | | 5-95% | 917,284 | 0.277 | 0.284 | 0.257 | 0.281 | -7.2E-03 | -2.4E-02 | -4.9E-04 | -1.4 | 917,284 | 7.8% |
| | | >=95% | 3,643,149 | 1.000 | 1.000 | 1.000 | 1.000 | -1.3E-04 | -2.9E-04 | 2.5E-06 | 2.7 | 3,643,149 | 30.9% |
| | EUR | <=5% | 7,448,705 | 0.003 | 0.008 | 0.012 | 0.007 | -5.3E-03 | 5.0E-03 | 8.9E-05 | 9.2 | 7,448,705 | 63.2% |
| | | 5-95% | 644,315 | 0.340 | 0.289 | 0.251 | 0.296 | 5.0E-02 | -4.5E-02 | -8.9E-04 | -1.9 | 644,315 | 5.5% |
| | | >=95% | 3,688,376 | 0.999 | 0.998 | 0.996 | 0.998 | 1.8E-03 | -1.8E-03 | -1.8E-04 | -17.8 | 3,688,376 | 31.3% |
| 1240K | AFR | <=5% | 185,776 | 0.102 | 0.015 | 0.040 | 0.027 | 8.7E-02 | 1.3E-02 | -2.3E-03 | -16.6 | 185,776 | 18.1% |
| | | 5-95% | 816,268 | 0.381 | 0.355 | 0.316 | 0.358 | 2.6E-02 | -4.1E-02 | -1.9E-03 | -4.6 | 816,268 | 79.7% |
| | | >=95% | 22,190 | 0.948 | 0.989 | 0.971 | 0.984 | -4.1E-02 | -1.3E-02 | -5.1E-04 | -2.3 | 22,190 | 2.2% |
| | EUR | <=5% | 221,282 | 0.035 | 0.091 | 0.127 | 0.079 | -5.7E-02 | 4.8E-02 | -1.6E-04 | -0.7 | 221,282 | 21.6% |
| | | 5-95% | 750,868 | 0.391 | 0.333 | 0.291 | 0.341 | 5.8E-02 | -5.0E-02 | -1.8E-03 | -4.3 | 750,868 | 73.3% |
| | | >=95% | 52,084 | 0.957 | 0.854 | 0.782 | 0.869 | 1.0E-01 | -8.7E-02 | -1.2E-02 | -18.8 | 52,084 | 5.1% |
| global MAF 5% | AFR | <=5% | 711,440 | 0.173 | 0.019 | 0.020 | 0.037 | 1.5E-01 | -1.7E-02 | -3.1E-03 | -16.5 | 711,440 | 14.7% |
| | | 5-95% | 4,086,812 | 0.383 | 0.361 | 0.317 | 0.363 | 2.2E-02 | -4.6E-02 | -1.1E-03 | -2.4 | 4,086,812 | 84.3% |
| | | >=95% | 50,592 | 0.850 | 0.980 | 0.965 | 0.965 | -1.3E-01 | 8.0E-04 | -7.4E-04 | -1.5 | 50,592 | 1.0% |
| | EUR | <=5% | 692,601 | 0.093 | 0.168 | 0.163 | 0.147 | -7.5E-02 | 1.5E-02 | 2.0E-03 | 4.2 | 692,601 | 14.3% |
| | | 5-95% | 3,945,569 | 0.373 | 0.319 | 0.280 | 0.327 | 5.3E-02 | -4.7E-02 | -1.1E-03 | -2.3 | 3,945,569 | 81.4% |
| | | >=95% | 210,671 | 0.931 | 0.771 | 0.671 | 0.794 | 1.6E-01 | -1.2E-01 | -1.9E-02 | -20.0 | 210,671 | 4.3% |
| AFR MAF 5% | AFR | <=5% | 74,186 | 0.047 | 0.043 | 0.048 | 0.081 | 4.3E-03 | -3.3E-02 | -1.1E-03 | -4.1 | 74,186 | 1.2% |
| | | 5-95% | 6,096,746 | 0.288 | 0.294 | 0.268 | 0.292 | -6.6E-03 | -2.4E-02 | -5.7E-04 | -1.7 | 6,096,746 | 98.7% |
| | | >=95% | 6,664 | 0.951 | 0.964 | 0.936 | 0.932 | -1.3E-02 | 4.4E-03 | -1.2E-03 | -2.6 | 6,664 | 0.1% |
| | EUR | <=5% | 2,385,634 | 0.019 | 0.109 | 0.122 | 0.093 | -9.0E-02 | 2.9E-02 | 1.3E-03 | 7.5 | 2,385,634 | 38.6% |
| | | 5-95% | 3,405,679 | 0.395 | 0.358 | 0.313 | 0.364 | 3.7E-02 | -5.1E-02 | -6.1E-04 | -1.1 | 3,405,679 | 55.1% |
| | | >=95% | 386,279 | 0.965 | 0.839 | 0.752 | 0.856 | 1.3E-01 | -1.0E-01 | -1.2E-02 | -21.0 | 386,279 | 6.3% |
| archaic | AFR | <=5% | 1,421,754 | 0.009 | 0.001 | 0.004 | 0.002 | 7.3E-03 | 1.8E-03 | -1.1E-05 | -0.8 | 1,421,754 | 66.3% |
| | | 5-95% | 605,174 | 0.436 | 0.434 | 0.429 | 0.433 | 2.2E-03 | -4.2E-03 | -8.6E-04 | -1.3 | 605,174 | 28.2% |
| | | >=95% | 118,504 | 0.985 | 0.994 | 0.982 | 0.993 | -9.3E-03 | -1.0E-02 | 7.8E-05 | 1.3 | 118,504 | 5.5% |
| | EUR | <=5% | 1,481,463 | 0.008 | 0.013 | 0.021 | 0.012 | -5.0E-03 | 9.2E-03 | -4.6E-04 | -10.1 | 1,481,463 | 69.1% |
| | | 5-95% | 483,239 | 0.422 | 0.403 | 0.394 | 0.406 | 1.9E-02 | -1.2E-02 | 2.0E-03 | 3.3 | 483,239 | 22.5% |
| | | >=95% | 180,730 | 0.981 | 0.932 | 0.889 | 0.939 | 4.9E-02 | -5.0E-02 | -4.5E-03 | -13.9 | 180,730 | 8.4% |

**Suppl. Table 12.** Dissecting the statistic $f_4$(Burmese, Dinka; Ju|'hoan North, Sengwer) composed of three African groups and one East Asian group. Sites were stratified by DAF in Africans or DAF in Europeans into three bins: nearly fixed ancestral (DAF <=5%), non-fixed (DAF 5-95%), and nearly fixed derived (DAF >=95%). This was done for "all sites", for random ascertainment (AT/GC sites,) and for 4 non-random ascertainment schemes as indicated in the leftmost column. The number and proportion of $f_4$-informative sites falling into each bin are shown. Mean DAF in four populations, mean differences in DAF between populations 1 and 2, populations 3 and 4, mean products of the DAF differences (i.e., $f_4$-statistics) and their Z-scores are shown for these frequency bins.

| Simulated topology | Group name | Effective population size, diploid individuals | Sampling date, generations |
|---|---|---|---|
| Suppl. Fig. 17b | Chimpanzee | 1,000 | 0 |
| | Denisovan | 3,000 | 2,000 |
| | Neanderthal | 3,000 | 2,000 |
| | African 1 | 22,500 | 0 |
| | African 2 | 22,500 | 0 |
| | African 3 | 22,500 | 0 |
| | African 4 | 22,500 | 0 |
| | African 5 | 22,500 | 0 |
| | Non-African 1 | 5,000 | 0 |
| | Non-African 2 | 5,000 | 0 |
| | Non-African 3 | 5,000 | 0 |
| Fig. 2a | a0 (ancestor of a2 + na1 + na2) | 16,914 | N/A |
| | African 1 (a1) | 44,541 | 0 |
| | African 2 (a2) | 46,139 | 0 |
| | eff. pop. size after the out-of-Africa bottleneck | 1,506 | N/A |
| | Denisovan (d) | 16,758 | 1,700 |
| | Neanderthal 1 (n1) | 14,399 | 3,790 |
| | Neanderthal 2 (n2) | 14,399 | 1,700 |
| | AMH (a1 + a2 + na1 + na2) | 222,379 | N/A |
| | ancestral Neanderthal population | 8,145 | N/A |
| | ancestral non-African population (na1 + na2) | 8,821 | N/A |
| | non-African 1 (na1) | 35,744 | 0 |
| | non-African 2 (na2) | 14,763 | 0 |
| | Neanderthal + AMH ancestral population | 86,161 | N/A |
| | super-archaic population | 35,414 | N/A |
| | chimpanzee (outgroup) | 1,000 | N/A |
| | root population (chimpanzee + archaic + AMH) | 13,858 | N/A |

**Suppl. Table 13.** Parameters of the simulated demographic histories that are not shown on the respective graphs in Fig. 2a and Suppl. Figs. 17b and 18b: effective population sizes and sampling dates.
